# Domain-specificity in phonological and acoustic characteristics of pseudoword iconicity across multiple meaning domains

**DOI:** 10.1101/2024.09.03.610973

**Authors:** Simon Lacey, Kaitlyn L. Matthews, Ahaana Shrivastava, K. Sathian, Lynne C. Nygaard

**Author notes:** Corresponding authors: Lynne C. Nygaard Department of Psychology Emory University, College of Arts and Sciences Atlanta, GA 30322, USA; K. Sathian, Department of Neurology, Penn State College of Medicine Hershey, PA 17033-0859, USA.

## Abstract

In spoken language, iconic (sound-symbolic) words are those whose sounds convey their meaning. Iconicity is widespread in natural languages, whether signed or spoken, but its instantiation across different domains of meaning has not been systematically studied. Here, participants rated a set of 537 auditory pseudowords on opposing dimensions of eight different sound-symbolic domains: shape (rounded-pointed), roughness (smooth-rough), hardness (hard-soft), weight (light-heavy), size (small-big), brightness (bright-dark), arousal (calming-exciting), and valence (good-bad). Ratings showed cross-domain relationships, some mirroring those between corresponding physical domains, e.g. size and weight ratings were associated, reflecting a physical size-weight relationship, while others involved figurative relationships, e.g. bright/dark mapped onto good/bad, respectively. Using four separate multiple regression analyses, we found that the phonetic categories, phonemic associations, and acoustic associations at both the whole-item and segmental levels, formed unique sets with characteristic feature weightings for each meaning domain studied. For the majority (12 of 16 dimensions), the phonemic regression accounted for the most variance in the ratings, followed by the phonetic category, segmental acoustic and whole-item acoustic regressions. We conclude that pseudoword iconicity is present across a range of meaning domains, with domain-specific patterns of linguistic and acoustic properties. Linguistic characterizations best captured judgments of iconicity across domains, suggesting that these properties represent the bundles of acoustic, articulatory, and abstract linguistic factors that may underlie iconicity in spoken language.

## INTRODUCTION

Iconicity in language refers to the idea that the form of a sign in some way resembles its meaning (e.g., Winter et al., in press). Thus, in spoken language, the sound of a word may resemble its meaning, for example, in onomatopoeic words like ‘thud’ and ‘splash’, that imitate the sound they describe (e.g., Catricalà & Guidi, 2015). Iconic words extend beyond the auditory modality to encompass, for example, the visuo-haptic property of shape: ‘balloon’ and ‘spike’ sound like, and refer to, rounded and pointed objects respectively (Sučević et al., 2015)^1^. But iconicity also occurs in non-spoken language, such as sign language (e.g. Emmorey, 2023) and common gestures, like thumbs up or down to indicate good or bad. The opposing idea is that the connection between form and meaning is arbitrary, that words are associated with their meaning purely by convention (e.g., de Saussure, 1916/2009; Hockett, 1959), and that instances of iconicity are too few to be functionally meaningful (see Lev-Ari & McKay, 2023). Alongside this, however, is a large body of work showing that humans are sensitive to iconic mappings, the classic being the ‘kiki-bouba’ effect, in which pseudowords such as ‘kiki’ or ‘takete’ are reliably associated with pointed shapes, while ‘bouba’ or ‘maluma’ are associated with rounded shapes (Köhler, 1929, 1947; Ramachandran & Hubbard, 2001). Moreover, recent work shows that, far from occurring sporadically, sound iconicity regularly occurs in thousands of spoken languages, across different language families (Blasi et al., 2016).

### Phonetic aspects of iconicity

Most studies of sound iconicity investigate a single mapping and by far the most extensively researched domain is shape. Some studies of the sound-to-shape mapping have investigated its phonetic underpinnings, examining the roles of voiced and unvoiced consonants (Shinohara & Kawahara, 2010; D’Onofrio, 2014; Cuskley et al., 2017), obstruents and sonorants (McCormick et al., 2015), consonants and vowels (Fort et al., 2015; Nielsen & Rendall, 2011), vowel formants (Knoeferle et al., 2017), and rounded and unrounded vowels (Maurer et al., 2006; McCormick et al., 2015). More recent studies have explored the influence of acoustic parameters derived from spectro-temporal and voice characteristics (Knoeferle et al., 2017; Lacey et al., 2020; Akita, 2021; Villegas et al., 2023). However, investigating iconicity by reference to a single domain has obvious limitations in that the relative importance of these factors, phonetic or acoustic, might change depending on the particular sound-meaning mapping. Furthermore, examining multiple domains enables us to consider whether sound-meaning mappings that differ with respect to phonological structure can nonetheless be grouped by reference to higher-order factors such as intensity or magnitude (Sidhu & Pexman, 2018), arousal (Aryani et al., 2020), or categories such as activity, valence, potency, and novelty (Osgood et al, 1957; Sidhu et al., 2022), although these higher-order factors are not the focus of the present study.

Few studies have investigated more than a single domain of meaning and, in some cases, the number of domains comes at the expense of the number of items. For example, Miron (1961) examined 15 domains but obtained ratings for only 50 pseudowords, while Johansson et al. (2020) investigated 344 basic concepts in representative spoken languages from 245 language families but using only a single word for each concept. Most recently, Sidhu et al. (2022) obtained ratings for 24 domains using a set of only 40 pseudowords. By contrast, Westbury et al. (2018) used a set of 7996 pseudowords to investigate 18 semantic categories grouped into 5 domains^2^. Tzeng et al. (2017) found that participants could select the correct iconic meaning for 80 foreign real words covering only 4 domains. Finally, Winter et al. (2017) obtained iconicity ratings for 3001 real words, finding that ratings were higher for words relating to the five sensory domains than for words belonging to more abstract domains. For completeness, Greenberg and Jenkins (1966) investigated 26 domains, but only in relation to individual consonants and vowels (though this is important for comparison to the phonetic make-up of both real words and pseudowords). We should also note that Winter et al. (2023) obtained iconicity ratings for more than 14000 real words, but these were not organized into domains.

Thus, studies of sound iconicity have generally either examined relatively few semantic domains but employed large pseudoword sets, or many semantic domains but with relatively smaller pseudoword sets. Here, we collected ratings of a relatively large set of 537 consonant-vowel-consonant-vowel (CVCV) pseudowords (McCormick et al., 2015) for eight meaning domains as a reasonable compromise to this trade-off between items and domains. The pseudowords were created using phonemes with reliable, but not exclusive, associations to the shape domain (McCormick et al., 2015; Lacey et al., 2020). We extended our previous work on the shape domain by also obtaining ratings for other perceptual properties of physical objects, i.e., roughness, hardness, weight, size, and brightness, as well as for the more abstract domains of arousal and valence. This choice of domains enabled us to compare concrete and abstract domains, and different sensory modalities; for example, brightness is perceived visually, roughness is primarily salient to touch (Klatzky et al., 1987), assessing hardness requires active touch (Srinivasan & LaMotte, 1995), and weight judgments rely on active lifting and associated somatosensory processing (e.g., Valchev et al., 2017). More importantly, examining multiple domains enables us to test whether iconic sound-meaning mappings are domain-specific. If so, the phonetic profile (i.e., the relative contributions of different phonetic categories, and indeed particular phonemes, to any particular sound-meaning mapping) would vary depending on the domain. Alternatively, sound-to-meaning correspondences may reflect mappings to a single, domain-general, abstract factor (such as arousal, magnitude, or valence [see Aryani et al., 2020; Sidhu & Pexman, 2018; Spence, 2011]). In the latter case, the phonetic and phonemic profile would be similar across domains and reflect that of the hypothesized domain-general factor. For example, if the domain-general factor were arousal then the iconic phonetic profile for arousal should be preserved across all other domains.

### Acoustic aspects of iconicity

While the phonetic categories involved in such iconic mappings have been extensively investigated, whether and how acoustic properties, of either real words or pseudowords, contribute to their sound-meaning mappings is largely unexplored (Knoeferle et al., 2017). Acoustic properties may be important because they relate to the listener’s end of the ‘speech chain’ (Denes & Pinson, 1993) with the spectro-temporal characteristics of speech mapping to phonetic categories and individual phonemes. For example, intracranial recordings show that, during listening to natural speech, the superior temporal gyrus (STG) shows phonetic selectivity arising from neuronal populations tuned to the specific spectro-temporal profiles of each phonetic category (Mesgarani et al., 2014; see also Oganian et al., 2023; Hamilton et al., 2020). Accordingly, cortical responses to phonetic features of speech could be decoded from low-level acoustic parameters (Daube et al., 2019).

However, investigation of the role of such acoustic properties in iconicity has been intermittent, largely confined to pseudowords and a limited range of acoustic features and meaning domains. For example, in the study of Knoeferle et al. (2017), participants rated auditory pseudowords containing one of five vowels (back rounded /ɒ/ and /u/, back unrounded /ɑ/, and front unrounded /ε/ and /i/): shape ratings were related to the frequencies of the vowel formants^3^ F2 and F3, while size ratings were related to F1 and F2. When participants vocalized a single vowel sound (back unrounded /ɑ/) in response to visual shapes (dodecagon vs. triangle), the frequency of F3 was higher for triangles, the more pointed of the two (Parise & Pavani, 2011). However, Parise & Pavani (2011) did not find any associations between the fundamental frequency (F0) or formant frequencies of vocalizations in response to large and small visual stimuli, perhaps as a result of the restriction to a single vowel. Nonetheless, vocalizations were louder for complex compared to simple shapes (dodecagon vs. triangle) and were also louder for brighter than darker stimuli (Parise & Pavani, 2011). This latter result is consistent with a later finding that speakers produced pseudowords referring to bright colors with greater amplitude than those for darker colors (Tzeng et al., 2018). Speakers also produced pseudowords for brighter colors with higher fundamental frequency, and shorter duration, than those for darker colors, and listeners could use these prosodic cues to reliably assign pseudowords to their target color (Tzeng et al., 2018).

In the first such systematic exploration of its kind, we previously investigated the contributions of a range of spectro-temporal and vocal parameters of a large set of pseudowords to iconicity of the rounded-pointed dimensions^4^ of shape (Lacey et al., 2020). In a novel application of representational similarity analysis (RSA: Kriegeskorte et al., 2008), we showed that representational dissimilarity matrices (RDMs) for ratings of pseudowords as rounded or pointed were significantly correlated with RDMs for three spectro-temporal parameters: spectral tilt, the fast Fourier transform (FFT) and the speech envelope (Lacey et al., 2020). Spectral tilt, i.e. the slope of the power spectrum over the frequency range, was steeper for pseudowords rated as more rounded with power concentrated at lower frequencies, but flatter for pseudowords rated as more pointed as spectral power shifted to higher frequencies. The FFT, representing the power spectrum of the pseudoword across its duration, showed smoother distributions of spectral power over time, mainly at lower frequencies, for pseudowords with higher roundedness ratings, compared to more irregular distributions spreading to higher frequencies for pseudowords with higher pointedness ratings. The speech envelope, measuring changes in the amplitude profile over time, was more continuous and smoother for pseudowords rated as more rounded, but more uneven and discontinuous for pseudowords rated as more pointed. (In relation to the FFT and speech envelope, Fort & Schwartz (2022) have suggested that the iconic sound-shape mapping may derive from audiovisual regularities that can be observed for physical objects. For example, rolling a round object on a surface produces smooth, continuous sounds whereas a spiky object produces irregular, discontinuous sounds (Fort & Schwartz, 2022)). The vocal parameters could be broadly divided into those that reflect the relative periodicity of the speech signal, which largely results from the balance between voiced and unvoiced segments, (i.e., the fraction of unvoiced frames [FUF], the mean autocorrelation [MAC], the harmonics-to-noise ratio [HNR], and pulse number), and those that reflect vocal variability (i.e., jitter [variability in frequency], shimmer [variability in amplitude], and the standard deviation of F0 [F0_SD_: variability in the fundamental frequency of the speech signal]). Conventional correlational analyses showed that periodicity (FUF, MAC, HNR, and pulse number) decreased, and vocal variability (shimmer and jitter) increased as ratings of pseudowords transitioned from rounded to pointed (Lacey et al., 2020). However, F0_SD_, while also a measure of vocal variability, was unrelated to shape ratings (Lacey et al., 2020). A full description of all the acoustic parameters used in Lacey et al. (2020) and in the current work is provided in the acoustic analysis section below.

Subsequently, in the shape domain, pulse phonation (in which the vocal folds open and close – i.e., pulse – less frequently than normal, resulting in a ‘creaky voice’ [Ishi et al., 2008]) has been associated with iconic pointedness (Akita, 2021), replicating our finding that lower pulse numbers, i.e. a less periodic, more uneven voice pattern, indicated pointedness (Lacey et al., 2020). In addition, pseudowords rated as pointed were associated with increased vocal variability, defined as rapid variation in amplitude (Villegas et al., 2023), consistent with our finding that shimmer increased as ratings changed from rounded to pointed (Lacey et al., 2020). Roundedness ratings have also been associated with higher pitch (Villegas et al., 2023), and with falsetto phonation, which is characterized by a higher F0 than the speaker’s normal voice (Akita, 2021). However, both of these findings seem to contrast with the crossmodal correspondence between low/high pitch and obtuse/acute, i.e., less/more pointed, angles respectively (Parise & Spence, 2012). In a different domain, pulse phonation was associated with large size (Akita, 2021), and pseudowords rated as bigger were associated with increasing loudness while those rated as smaller were associated with increasing sharpness^5^ (Villegas et al., 2023).

### Outline of present study

Here, we evaluate the relationships of a range of phonological and acoustic variables to iconic mappings across eight domains of meaning: shape (rounded/pointed), size (small/big), the roughness (/rough) and hardness (hard/soft) aspects of the texture domain, weight (light/heavy), brightness (bright/dark), arousal (calming/exciting), and valence (good/bad). Participants listened to a large set of spoken pseudowords that sampled from the phonetic space of American English and rated how rounded, smooth, bright, etc., each item sounded on a unipolar seven-point Likert scale (i.e., 1 = ‘not rounded’ while 7 = ‘very rounded’, for example). We then compared these ratings to sets of phonological and acoustic parameters using four independent multiple regressions. The parameter sets comprised (1) phonetic categories; (2) the specific phonemes constituting the pseudowords; (3) whole-item acoustic parameters reflecting spectro-temporal characteristics and voice quality as used in our previous study of the shape domain (Lacey et al., 2020); and (4) acoustic parameters for the phonemic segments comprising each pseudoword. This use of multiple characterizations of pseudoword properties (i.e., in terms of phonetic categories, individual phonemes, and both whole-item and segmental acoustic profiles) allowed us to evaluate which representation of pseudoword form best accounts for perception of sound iconicity in spoken language.

As a secondary aim, the use of multiple domains of meaning allowed us to consider some of the mechanistic accounts of sound iconicity and the extent to which these might generalize across multiple, or even all, domains, or are specific to particular domains. A strict domain-general account might suggest that all sound-symbolic mappings reflect associations with a single dimension such as arousal (see Aryani et al., 2020) or magnitude (see Sidhu & Pexman, 2018). On this account, phonetic-acoustic profiles (i.e., the pattern of the relative importance of the various phonetic, phonemic, and acoustic parameters) should be similar across all domains, reflecting the mapping to the index domain. For example, if the index domain were arousal and HNR were the most important parameter, then HNR would be the most important parameter across all other domains. By contrast, on a domain-specific account, the size domain might be governed by statistical regularities in the environment in that small/big items tend to resonate at higher/lower frequencies, respectively (see Sidhu & Pexman, 2018; Spence, 2011), and thus phonemes with high/low F0 would be prominent in the phonetic-acoustic profile for size. However, such environmental regularities might be harder to reconcile with more abstract domains like valence (since many things that are good/bad make no noise) which might be governed by a different mechanism, resulting in a different phonetic-acoustic profile. Thus, in the current study, we both evaluate which types of representation may underlie sensitivity to sound iconicity and begin to examine the extent of putative domain-general and domain-specific accounts.

## METHODS

### Participants

Participants were recruited, and compensated for their time, online via the Prolific participant pool (https://prolific.co: see Peer et al., 2017). The eligibility criteria were that participants be aged between 18 and 35, be speakers of American English as their first language, and have no language or hearing disorders. A total of 646 people took part, but data from 257 were excluded: 73 because they failed a headphone screen (see General Procedures below) and a further 184 because post-task questionnaires (see General Procedures below) indicated that they were bilingual or spoke a second language (120); they used less than 5 values on the full 7-point scale (47); there was a malfunction in stimulus presentation (8); they experienced synesthesia (3), which has been associated with increased sensitivity to sound-symbolic crossmodal correspondences (Lacey et al., 2016); and 6 for other reasons. Thus, the final sample comprised 389 participants (171 male, 208 female, 6 non-binary, 2 agender, 1 gender-fluid, and 1 declined to state their gender; mean age 27 years, 1 month (standard deviation = 8 months). The rating experiments were hosted on the Gorilla platform (https://gorilla.sc: Anwyl-Irvine et al., 2020) where participants also gave informed consent. All procedures were approved by the Emory University Institutional Review Board. For the roughness aspect of the texture domain (see below), we include data from Nayak (2024) for an additional 58 participants (26 male, 31 female and 1 non-binary; mean age: 25 years, 9 months, SD = 3 years, 7 months) collected separately from the above but following the same procedures. This study was approved by the Penn State University Institutional Review Board.

### Auditory Pseudowords

We used the 537 two-syllable CVCV pseudoword set created by McCormick et al. (2015), including only phonemes and combinations of phonemes that occur in spoken English, and excluding items that were considered to be homophones of real words. Because of the high number of possible phoneme combinations, the pseudoword set was restricted to a subset of phonemes and phoneme combinations that were hypothesized, based on prior studies, to reliably map to the rounded/pointed dimension of shape, which was the focus of the original study (see McCormick et al., 2015, for details). Within a given pseudoword, consonants were either both unvoiced (as in /kupo/) or both voiced (as in /gubo/) but could vary in place of articulation. Pseudowords contained either back/rounded or front/unrounded vowels and did not contain repeated syllables, so possible pseudowords such as /kiki/ and /lolo/ were excluded. For a complete description of the stimulus set, see McCormick et al. (2015); the complete set of pseudowords is available at https://osf.io/ekpgh/.

The pseudowords were recorded in random order, and with neutral intonation, by a female speaker whose first language was American English. Recordings took place in a sound-attenuated room, using a Zoom 2 Cardioid microphone, and were digitized at a 44.1 kHz sampling rate. Two independent judges assessed whether each item was spoken with neutral intonation, e.g., that they avoided any kind of ‘list intonation’ (see Cauldwell & Hewings, 1996); accurately produced the intended phonemic content; and sounded consistent with other items (e.g., did not sound faster/slower or louder/softer than other items). For those pseudowords where one of the judges noted that the recording did not conform to any one of these requirements, that item was re-recorded and judged again. Each pseudoword was then down-sampled at 22.05 kHz, which is a standard sampling rate for speech, and amplitude-normalized using Praat speech analysis software (Boersma & Weenink, 2012). The pseudowords had a mean (± standard deviation) duration of 457 ± 62 ms.

### Sample size

In order to get a reliable estimate of the mean rating for each pseudoword, we expected that a sample size of 25-30 participants per scale would be sufficient. McCormick et al. (2015), whose ratings data we used in our previous study (Lacey et al., 2020), had sample sizes of 15 and 16 for their rounded and pointed scales respectively, while other studies included as many as 40+ or 70+ participants (e.g., Miron, 1961). Power analysis based on Equation 7 from Bonett (2002) suggests that a sample size of 30 would have power of .8 to detect a Cronbach alpha of .5 for inter-rater reliability. The sample size for the correlations between the ratings scales is fixed by the number of pseudowords at 537 and this gives statistical power of >80% to detect a Pearson correlation of .3 when α = .05 (regression power analysis was carried out in SPSS v29.0 (IBM Corp, Armonk NY).

### General procedures

Participants recruited via Prolific followed a link that took them to the experiment on the Gorilla platform. At the landing page, they could read the consent form and a short description of the task, before clicking on ‘yes’ to continue and complete the rating task, or ‘no’ to exit. Having consented, and before continuing with the rating task, participants completed a headphone check designed to ensure compliance with headphone use for web-based experiments (Woods et al., 2017). Participants who failed the headphone check could take it a second time but if they failed again, while they could still participate, in order to avoid discriminating against those who did not have access to headphones, their data were excluded from analysis (except that for the roughness data, which were collected separately, participants who failed the headphones at both attempts were excluded immediately and did not progress to the ratings task). The randomizer function in Gorilla then assigned participants (including those who failed the headphone check) to one of the two opposing rating scales for the relevant domain. Once the rating task had been completed, participants were directed to a series of short questionnaires that asked about demographic information, and language experience and ability. Participants could then read a debriefing statement and exit the experiment. Once participants had completed any one scale, they were excluded from further participation in the study. Thus, the fourteen scales were completed by independent groups of participants.

### Perceptual rating tasks

Participants were randomly assigned to one of fourteen 7-point Likert-type scales and rated all 537 pseudowords on the scale to which they were assigned. The scales represented categorical opposites across seven different meaning domains: shape (rounded/pointed, N =30/30), two aspects of texture, hardness (hard/soft, N = 27/27) and roughness (smooth/rough, N = 29/29; these data were collected in Nayak, 2024), weight (light/heavy, N = 29/26), size (small/big, N = 32/31), brightness (bright/dark, N = 24/26), arousal (calming/exciting, N = 30/28), and valence (good/bad, N = 26/23). The sample sizes vary slightly due to the exclusions noted above; we opted not to recruit replacement participants because the sample sizes largely fell within our desired range of 25-30. In order to avoid response bias, one of the scales rated one end of the meaning domain, for example, roundedness, from 1 (not rounded) to 7 (very rounded) and the other rated the other end, i.e., pointedness, from 1 (not pointed) to 7 (very pointed). This also avoided assumptions about contrastive meaning for each domain, i.e., that ‘rounded’ inherently also means ‘not pointed’, or that ‘not exciting’ also means ‘calming’ as opposed to ‘dull’. The instructions included several related terms for the categorical opposites in each domain (see Supplementary Table 1). The 7-point rating scale appeared in the center of the screen on each trial and always listed 1-7 from left to right. Each pseudoword was presented only once, in random order.

### General statistical approach

Since our goal was to characterize the properties of pseudowords that contribute to perception along a particular dimension of meaning, rather than to characterize the sources of variation in those ratings, all analyses are across items based on per-item ratings averaged across participants, and that participant variability is not considered. In addition, we analyzed each unipolar scale (i.e. ‘not rounded’ to ‘very rounded’) separately instead of combining them into bipolar scales (i.e. ‘rounded’ to ‘pointed’) because there is no guarantee that antonym pairs, e.g. rounded-pointed, will necessarily result in opposing significant effects for the predictors, positive for one and negative for the other (see Westbury et al., 2018; Monaghan & Fletcher, 2019). The advantage of this is that we obtain ratings for items that are clearly regarded as ‘very rounded’ etc. rather than combining them with those that are regarded as merely ‘not pointed’ (see also Connell & Lynott [2012] who found that people applied different decision criteria at each end of a bipolar ‘concrete’ to ‘abstract’ scale). All statistical analyses were carried out in SPSS v29.0 (IBM Corp, Armonk NY).

## RATINGS ANALYSIS

Within each domain, the two scales were intended to be in opposition to each other, e.g. good vs bad, and therefore were predicted to be strongly negatively correlated. This was, in fact, the case (see the bolded values on the diagonal in Table 2). Thus, for each domain, the two scales adequately reflected the categorical opposites and the meaning domains likely consisted of continuous, contrastive, dimensions. The strongest correlation, however, was only -.69 (between the rough and smooth scales), vindicating the point made in the General Statistical Analysis section above, that ‘very rounded’ etc. does not necessarily equate to ‘not pointed’ etc., otherwise these negative correlations would likely have been stronger. In relation to the shape domain, note that of the 10 most highly-rated pseudowords for each scale, 5 (2 rounded and 3 pointed) were common to both the current results and those of McCormick et al. (2015: see Table 3).

Across domains, there were some ‘double dissociation’ effects in that ratings on the two scales of a given dimension showed opposing correlations with those of another dimension. Interestingly, these seemed to cluster into cross-domain relationships that reflect either physical or metaphorical associations (Lacey et al., 2022). For example, for the size and weight domains, pseudowords rated as small were also rated as light but not heavy, while pseudowords rated as big were also rated as heavy but not light (Lacey et al., 2022). Other instances illustrate more metaphorical relationships. For the valence and weight domains, for example, pseudowords rated good were also positively correlated with light but negatively with heavy, while those rated as bad were positively correlated with heavy but negatively with light (Lacey et al., 2022).

## PHONETIC CATEGORY ANALYSIS

### Introduction

Full details of the consonants and vowels used in the pseudoword set, and their manner, voicing, and place of articulation, are provided in Table 1. Consonants were sampled from sonorants, affricates/fricatives (henceforth ‘af/fricatives’), and stops; for the af/fricatives and stops, half were voiced and half were unvoiced. Vowels were either back/rounded or front/unrounded. The place of articulation for consonants was grouped into bilabial/labiodental, alveolar, and post-alveolar/velar; for vowels, place of articulation was divided into middle and high vowel height. Note that this set is not exhaustive of English phonemes or phonetic categories, for example it does not include other fricatives like /h/ and /∫/, the glides /w/ and /y/, nor low vowels). Our predictions for the phonetic categories involved in the sound-iconic mapping for each domain are grounded in previous findings, as follows.

**Table 1:**
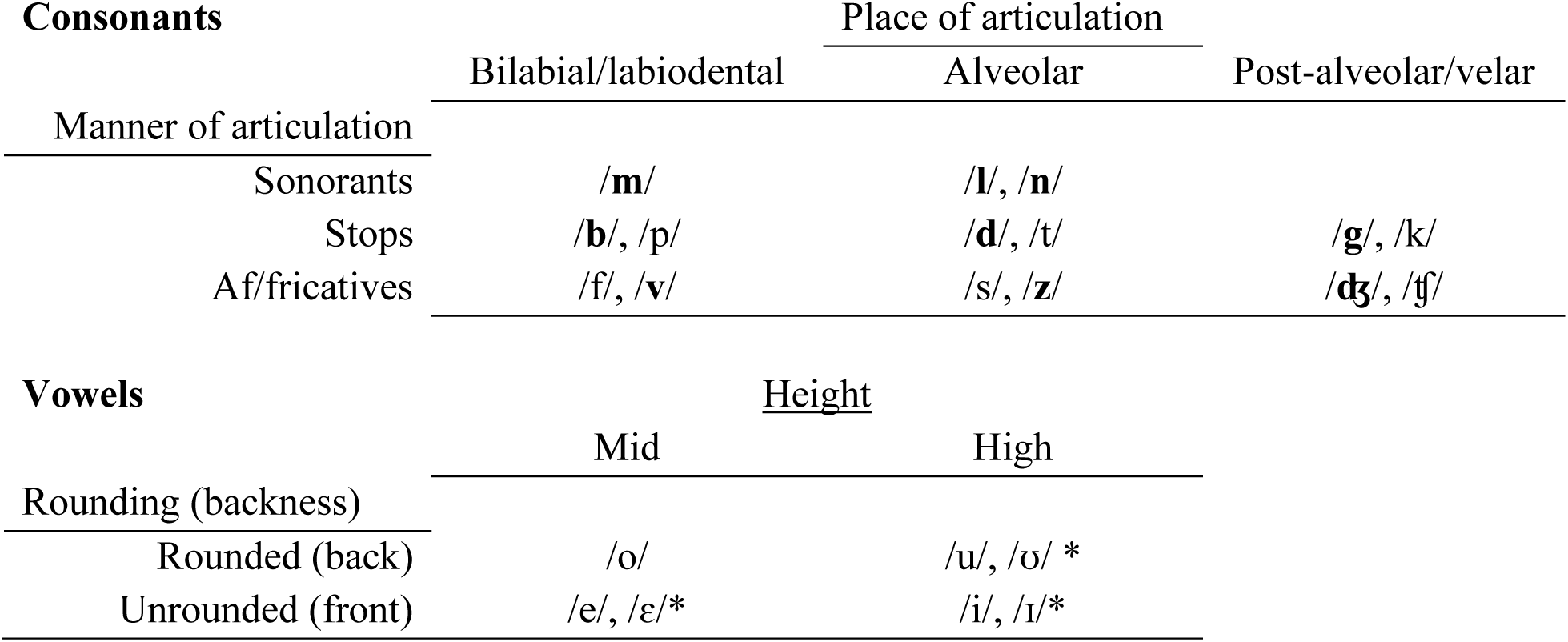
Phonetic and articulatory categories of consonants and vowels used in creating the CVCV pseudowords. Voiced consonants are in **bold** type. Vowels marked * only occur in the first vowel position as they do not occur in word-final positions in English.

#### Shape

For the shape domain, we predicted that roundedness would be associated with voiced (e.g., /b/, /d/) rather than unvoiced (e.g., /t/, /k/) consonants, and with back/rounded vowels (e.g., /u/ or /o/) (Köhler, 1929, 1947; Ramachandran & Hubbard, 2001; D’Onofrio, 2014; McCormick et al., 2015), while pointedness would be associated with stops (e.g., /p/ and /t/) rather than sonorants (e.g., /m/ or /l/), and with front/unrounded vowels (e.g., /i/ or /e/: Köhler, 1929, 1947; Ramachandran & Hubbard, 2001; McCormick et al., 2015). For the place of articulation, we predicted that pointedness would be associated with velar consonants, e.g., /g/ or /k/, following Westbury et al. (2018: note that this study reported on sharpness, which we take to be synonymous with pointedness since it was paired antonymically with roundness rather than bluntness). We predicted that roundedness would be associated with nasal consonants (Westbury et al., 2018): of the three nasal consonants in English, the present study uses /m/ and /n/; the other is /ŋ/ as in ‘sing’ (Reetz & Jongman, 2020). Additionally, roundedness should be associated with labial consonants like /m/ and /b/ (D’Onofrio, 2014). A more recent study generally supports these predictions for consonants, with roundedness associated with sonorants and voiced stops, and pointedness associated with unvoiced stops and voiced fricatives (Sidhu et al., 2022).

#### Size

Following the classic finding for sound-iconic size associations, we predicted that that front/unrounded and back/rounded vowels would reflect small and large size, respectively (Sapir, 1929; replications can be found in Auracher, 2017; Sidhu et al., 2022), and that mid-height vowels would be associated with larger size than high vowels (Shinohara & Kawahara, 2010: note that our phoneme inventory did not include low vowels). Among the consonants, we could predict that fricatives would be associated with small size, and stops with large size (Preziosi & Coane, 2017). The associations for fricatives may depend on voicing, with unvoiced/voiced fricatives associated with small and big, respectively (Sidhu et al., 2022). Consistent with this, Shinohara & Kawahara (2010) found that large size was associated with voiced, rather than unvoiced, obstruents. Similarly, while Westbury et al. (2018) found that large size was associated with voiced consonants, Sidhu et al. (2022) reported that this only related to voiced stops. Moreover, sonorants – which are always voiced – have also been associated with small size (Sidhu et al., 2022).

#### Roughness

For the roughness aspect of texture, we predicted that roughness would be associated with stops (Greenberg & Junkins, 1966) and voiced consonants generally (Sakamoto & Watanabe, 2018), and the voiced velar stop /g/ in particular, across different language speakers (Wong et al., 2022). For completeness, though not included in the phonemes here, roughness has also been associated with the trilled /r/ (Winter et al., 2022) and perhaps ‘r-like’ sounds generally (Anselme et al., 2025). By contrast, we predicted that smoothness would be reflected in the sonorants /l/ and /m/, and the unvoiced alveolar fricative /s/ (Greenberg & Jenkins, 1966) and unvoiced consonants generally (Sakamoto & Watanabe, 2018).

#### Hardness

For the hardness aspect of texture, we predicted that the hardness dimension would be associated with the alveolar affricate /ʧ/ and the velar stop /k/, which was found when textures were directly perceived via touch (Sakamoto & Watanabe, 2018) and with voiced fricatives and unvoiced stops generally (Sidhu et al., 2022; see also Greenberg & Jenkins, 1966). By contrast, the softness dimension would be associated with the bilabial stops /b/ and /p/, the alveolar sonorant /n/ (again, for textures directly perceived via touch, Sakamoto & Watanabe, 2018) and with sonorants generally (Sidhu et al., 2022; Greenberg & Jenkins, 1966). However, there appear to be no specific associations to vowels (Sidhu et al., 2022).

#### Weight

For the weight domain, predictions were somewhat complicated by differing results in prior studies. For example, stops have been described as sounding heavy, regardless of voicing (Greenberg & Jenkins, 1966). However, in another study heaviness was associated with voiced stops only, and also with voiced fricatives (Sidhu et al., 2022). We could predict that lightness would be associated with sonorants and unvoiced fricatives (Sidhu et al., 2022). For the vowels, we predicted that front/unrounded vowels would be associated with lightness, and back vowels with heaviness (Walker & Parameswaran, 2019; Sidhu et al., 2022).

#### Brightness

We predicted that brightness would be associated with unvoiced stops /k/ and fricatives /s/, and front vowels /i/, /ɪ/, while darkness would be associated with voiced stops /b/ /g/and sonorants /m/, and back vowels /u/ /o/ (Newman, 1933). Note that these findings were only partially replicated by Greenberg & Jenkins (1966) where brightness was associated with both voiced and unvoiced fricatives, and sonorants, while darkness was reflected in both voiced and unvoiced stops.

#### Arousal

For the arousal domain, we predicted that sonorants and back vowels would be associated with pseudowords perceived as calming while those perceived as exciting would be associated with voiced fricatives, unvoiced stops, and front vowels (Sidhu et al., 2022). These predictions are consistent with studies that found voiced consonants and long vowels were rated as sounding less arousing (Aryani et al., 2018), while unvoiced consonants, sibilants, and short vowels were rated as sounding more arousing (Aryani et al., 2020).

#### Valence

Sound-iconic mappings for valence might depend on the definition: moral good/bad vs emotional happy/sad, for example; here, we chose the good/bad dimension. For real words across several spoken languages, longer/shorter initial phonemes predicted positive/negative valence, respectively (Adelman et al., 2018), which suggests that, in our pseudoword set, sonorants and obstruents might be associated with good/bad, respectively. Supporting this prediction, a study of names for the complex concept of villains (Uno et al., 2020) found that voiced obstruents (stops, affricates, and fricatives) were associated with negative valence. Additionally, a recent study of constructed, fictional, spoken languages, intended by their creators to sound either harsh and evil (e.g., Klingon, Orkish) or pleasant and good (e.g., Sindarin, Quenya), suggests that voiced speech sounds are perceived more positively than unvoiced speech sounds (Mooshammer et al., 2023). Note, however, that unvoiced consonants and sibilants have been associated with negative valence, while voiced consonants have been associated with positive valence (Aryani et al., 2018). We predicted that the front vowel /i/ and the back vowel /o/ would be associated with positive and negative valence respectively (Körner & Rummer, 2022; Schmidtke et al., 2025). Finally, we should note that Sidhu et al. (2022) found no significant effect on good/bad valence ratings for either consonants or vowels in their pseudowords.

### Statistical analysis

We used linear regression to assess the extent to which each phonetic category was associated with judgments for each different meaning domain. A regression model was run separately for each of the two scales for all eight domains, in which each phonetic predictor was evaluated for its influence on perceptual ratings, relative to a reference predictor. For the manner of articulation of consonants, sonorants were the reference predictor for comparison with stops and af/fricatives; voiced consonants were the reference for unvoiced consonants. For the place of articulation of consonants, bilabial/labiodental consonants were the reference for comparison with alveolar and post-alveolar/velar consonants (each sub-divided into first or second consonant position). Note that grouping the consonants into bilabial/labiodental and post-alveolar/velar groups avoids multicollinearity issues arising from treating similar places of articulation separately. For the vowels, front unrounded vowels were the referent for back rounded vowels, and middle height vowels were the referent for high vowels (also sub-divided into first or second vowel position). These analyses were conducted in SPSS. All predictors were entered simultaneously, i.e., forced-entry (Field, 2018), meaning that there were no *a priori* assumptions about the relative importance of each of the phonetic categories. This approach allowed for predictors to be differently weighted depending on the domain. In addition, we computed the change in model fit (R^2^) attributable to each predictor.

The standardized beta coefficients express the change in the outcome variable (ratings) on each scale in standard deviation units. Taking the rounded scale (1 = ‘not rounded’ to 7 = ‘very rounded’) as an example, a positive β indicates that, compared to the reference, this predictor is rated towards the positive end of the scale, i.e. very rounded, and therefore associated with roundedness. A negative β indicates that, compared to the reference, this predictor is rated towards the negative end of the scale, i.e. as not rounded, and therefore that the reference predictor is associated with roundedness.

To evaluate the robustness of the regression results, we firstly addressed the problem of multicollinearity, the extent to which two or more predictors are correlated with each other: the stronger such correlations are, the more difficult it becomes to estimate the importance of any predictor on its own. However, the variance inflation factors (VIF) ranged from 1.02 to 2.55, well below the threshold value of 10 recommended by Myers (1990: Supplementary Table 2) and the mean VIF of 1.5 was close to 1, as recommended by Bowerman & O’Connell (1990). In addition, following Menard (1995), there were no tolerance values (the reciprocal of the VIF) below .2, with observed values ranging from .39 to .98, Thus, we concluded that multicollinearity was not a problem. Secondly, we tested the assumption of independent errors, i.e. the extent to which residual errors are correlated, via the Durbin-Watson test. Field (2018) recommends a conservative approach in which values < 1 or > 3 would be cause for concern. Since our observed values ranged from 1.79 to 2.23 (Supplementary Table 3), we concluded that the assumption of independent errors was met. Visual inspection of histograms and Q-Q plots for standardized residuals indicated that these were normally distributed for all scales. Plots of standardized residuals against standardized predicted values indicated that the assumption of homoscedasticity was also met for all scales.

### Results

Given the number of domains and predictors, we report in Table 4 only the R^2^, adjusted R^2^, and standardized beta weights; we report the change in R^2^ attributable to a predictor only where this is significant. The domains are reported in approximately descending order of the overall model fit, by reference to the highest R^2^ value for either of the two scales involved; we retain this order for subsequent analyses. Wherever possible, the examples given below of the associated phonetic categories are drawn from the ten most highly rated pseudowords on each of the two scales for each domain (Table 3); where they are not, the ordinal position of the item is specified.

#### Shape

Unsurprisingly, since the pseudoword set was originally designed to investigate sound-to-shape mappings, the shape domain had the best model fit between phonetic categories and ratings, with R^2^ of .88 for the rounded dimension and .76 for the pointed dimension (Table 4). Inter-rater reliability (Cronbach alpha) was .87 for the roundedness scale and .86 for the pointedness scale.

For roundedness, negative β values for the obstruents (stops and af/fricatives) indicated that, compared to sonorants, these attracted ratings towards the ‘not rounded’ end of the scale and that therefore, sonorants contributed more to ratings of pseudowords as rounded. Accordingly, obstruents were associated with only small changes to model fit as shown by the R^2^ change values (Table 4). Thus, consistent with previous studies (e.g., Köhler, 1929, 1947; Ramachandran & Hubbard, 2011; McCormick et al., 2015), roundedness was predicted by sonorants, for example, /m/ and /n/ in /mʊmo/ and /nonu/ (Table 3). However, there was also a small, but significant, negative β for unvoiced consonants generally, indicating that roundedness could also be associated with voiced consonants, exemplified by /b/ and /g/ in /gubo/ (Table 3) and, in fact, 5 of the 10 pseudowords with the highest rounded ratings contained voiced stops while only 3 contained sonorants. The change in model fit attributable to unvoiced consonants was minimal (Table 4).

In terms of place of articulation, negative β values for alveolar and post-alveolar/velar consonants meant that, by comparison, bilabial and labiodental consonants were more likely to attract ratings of roundedness (Table 4), with /b/, /m/, and /p/ all appearing in the most highly rated rounded pseudowords (Table 3). The β values for alveolar and post-alveolar/velar consonants were either small or non-significant, resulting in minimal or small R^2^ changes and thus model fit was relatively unaffected.

Back rounded vowels were by far the strongest predictor of roundedness ratings with a large positive β and a large change in the R^2^ value (in fact the largest values for these across all domains: Table 4), for example, /o/ and /ʊ/ in /mʊmo/ (Table 3) and consistent with previous studies (e.g., Köhler, 1929, 1947; Ramachandran & Hubbard, 2011; McCormick et al., 2015; Sidhu et al., 2022). There were also small, but significant, negative β values for high vowels in both vowel positions (Table 4), indicating that mid-height vowels, such as /o/ in /lolu/ (Table 3), were more likely to be rated as rounded than high vowels. As would be expected then, the addition of high vowels to the model brought minimal change to model fit.

By contrast, for pointedness, large and positive β values for obstruents indicated that these attracted ratings towards the ‘very pointed’ end of the scale, although stops were associated with a greater change in model fit than af/fricatives (Table 4). Thus, consistent with Sidhu et al. (2022), we observed, for example, /t/ and /ʧ/ respectively, in /tike/ and /ʧiʧe/ (Table 3). Unvoiced consonants, such as /t/ and /k/ in /tʊtu/ and /kike/ (Table 3) also produced a large positive β, indicating that these were more predictive of pointedness than voiced consonants (replicating McCormick et al., 2015) and producing a moderate improvement in model fit (Table 4). Together with stops, unvoiced consonants were the largest contributors to the pointedness model fit.

Alveolar and post-alveolar/velar consonants, exemplified by /t/ and /k/ respectively, were associated with significant positive β values indicating that they attracted ratings towards the pointed end of the scale. This was so in either consonant position and produced improvements to model fit ranging from minimal to small (Table 4).

A negative β value for back rounded vowels, indicated that pointedness was predominantly associated with front unrounded vowels, for example /ɪ/, /i/ and /ɛ/ in /kɪke/ (Table 3) and /tɛti/, the 30^th^ most pointed pseudoword, (consistent with Köhler, 1929, 1947; Ramachandran & Hubbard, 2001; McCormick et al., 2015; Sidhu et al., 2022). Accordingly, back rounded vowels made only a small change to the pointedness model (Table 4). A small, but significant, positive β value, and associated change to R^2^, for high vowels, like /i/ and /ɪ/, suggested that these made a very small improvement to the pointedness model (Table 4) but only in the first vowel position of the CVCV pseudowords, for example, /tike/ and /tɪke/ (Table 4).

#### Roughness

The next best model fit between phonetic categories and ratings was for the roughness aspect of the texture domain, with R^2^ of .57 for the smooth dimension, and .77 for the rough dimension (Table 4). Inter-rater reliability (Cronbach alpha) was .86 for the smoothness scale and .61 for the roughness scale.

For the smoothness dimension, large negative β values for the obstruents indicated that, compared to sonorants, these attracted ratings very much towards the ‘not smooth’ end of the scale (Table 4) and that therefore, sonorants contributed more to ratings of pseudowords as smooth as, for example, /l/ and /m/ in /mɛle/ and /leme/ (Table 3), consistent with Greenberg & Jenkins (1966). Obstruents were associated with a large change to R^2^ (Table 4) and improved model fit by substantially defining what was “not smooth’ and therefore perceived as sounding rough. The effect of voicing on smoothness ratings was not significant, in contrast to Sakamoto & Watanabe (2018).

For the place of articulation, alveolar consonants had no significant effect on smoothness ratings in either position, but there were significant small negative β values for the post-alveolar/velar consonants in both positions (Table 4). This suggested that, compared to post-alveolar/velar consonants, smoothness ratings were more associated with bilabial and labiodental consonants, such as /m/ in /mɪle/ (Table 3) or /f/ in /sife/ (the 42^nd^ smoothest item). The predominance of bilabial consonants meant that post-alveolar/velar consonants made minimal difference to the smoothness model (Table 4).

Vowels had little effect on smoothness ratings. There was no effect of vowel rounding, and although there was a small significant negative β for high vowels in the second position, the accompanying change to the smoothness model was minimal (Table 4).

As might be expected from the smoothness results, ratings on the roughness scale were predicted by obstruents (see Greenberg & Jenkins, 1966); these were associated with large positive β values that made some of the greatest improvements to model fit across all domains (Table 4).

The effect was greater for af/fricatives than stops, reflected in the fact that only one of the ten most highly-rated pseudowords contained a stop, /k/ in /kike/; whereas all the rest contained af/fricatives, for example, /z/ and /dʒ/ in /zɪdʒe/ (Table 3). In contrast to the smoothness dimension, roughness ratings were associated with unvoiced consonants (again in contrast to Sakamoto & Watanabe, 2018), for example, /k/ as above, and also /tʃ/ and /f/ in /tʃɪfe/ (Table 3), although the improvement to the roughness model was small (Table 4).

There were significant positive β values for both alveolar and post-alveolar/velar consonants, in either position indicating that they attracted ratings towards the ‘very rough’ end of the scale and resulting in improvements to model fit ranging from minimal to small (Table 4). Alveolar and post-alveolar/velar consonants contributing to perceptions of roughness are exemplified by /s/ and /z/, and by /tʃ/ and /dʒ/, respectively, for example in /tʃɪsi / and /zɪdʒe/ (Table 3).

Another contrast with the smoothness dimension was the effect of vowel rounding – there was a significant negative β for back rounded vowels indicating that, compared to front unrounded vowels, these were rated towards the ‘not rough’ end of the scale although the change in R^2^ was small (Table 4). Accordingly, front unrounded vowels, like /e/ and /i/ in / tʃetʃi/ account for most of the vowels in the most highly-rated rough pseudowords (Table 3). There was also a small but significant effect of vowel height with a positive β for high vowels in the first position, such as /ɪ/ in /zɪdʒe/, although the change to R^2^ was minimal (Table 4).

#### Hardness

For the hardness aspect of the texture domain, although R^2^ for the hard dimension was .68 (the fourth highest value overall), the model fit was also one of the most asymmetric for any domain, in that R^2^ for the soft dimension was only .33 (the fourth lowest value overall: Table 4). Inter-rater reliability (Cronbach alpha) was .72 for the hardness scale and .45 for the softness scale.

For the hardness dimension, relatively large and positive β values for the obstruents indicated that, compared to sonorants, these attracted ratings towards the ‘very hard’ end of the scale, although the effect on model fit was greater for stops than af/fricatives (Table 4). Consistent with previous findings (Greenberg & Jenkins, 1966; Sakamoto & Watanabe, 2018; Sidhu et al., 2022), we observed, for example the stops /p/ and /k/ in /peke/ and /toko/ (Table 3). Although af/fricatives did not occur in the 30 hardest-rated items, this likely reflects the relative weighting of stops and af/fricatives, .81 and .36 respectively. There was a small, but significant, positive β for unvoiced consonants indicating that these tended to be rated as hard (Table 4); for example, /t/ and /p/ in /teke/ and /pɪke/ (Table 3), consistent with Sidhu et al. (2022). The effect on model fit was minimal, however.

In terms of the place of articulation, there were significant positive β values (Table 4) for alveolar consonants in the second position, such as /t/ in /toko/, and with the post-alveolar consonants in either position, for example, /k/ and /g/ in /kike/ (Table 3) and /degi/, the 13^th^ hardest pseudoword.

There was no significant influence of vowel roundedness, but there was a small effect for vowel height. A small, but significant, negative β for high vowels in the second position of the CVCV pseudowords indicated that, compared to mid-height vowels, these tended to be rated towards the ‘not hard’ end of the scale (Table 4). Accordingly, second position high vowels made only a minimal difference to the hardness model.

Pseudoword softness, meanwhile, was associated with sonorants (see also Sidhu et al., 2022; Greenberg & Jenkins, 1966), such as /m/, /n/, and /l/ in /mumo/ and /nʊlu/ (Table 3). This is supported by the negative β values for obstruents, meaning that these attracted ratings towards the ‘not soft’ end of the scale compared to sonorants, which therefore contributed more to softness ratings (Table 4). The effect of obstruents on the softness model, in terms of the change in R^2^, was smaller than that on the hardness model (Table 4).

In terms of place of articulation, there were negative β values, and only small effects on the softness model, for both alveolar and post-alveolar/velar consonants, in either consonant position (Table 4). This indicated that, by comparison, labiodental consonants, such as /f/ in /sefi/ (Table 3) predicted softness.

Back rounded vowels, for example, in /mulu/ (Table 3), were a significant predictor of softness and were associated with a small improvement to model fit (Table 4). A small, but significant, negative β for high vowels in the first position indicated that, compared to mid-height vowels, these tended to be rated towards the ‘not soft’ end of the scale and made only a minimal difference to the softness model (Table 4); note that /e/ in /sefi/, is a mid-height vowel in a highly-rated ‘soft’ pseudoword (Table 3).

#### Weight

The weight domain had the next best model fit, with R^2^ of .43 for the light dimension and .53 for the heavy dimension (Table 4). Inter-rater reliability (Cronbach alpha) was .44 for the lightness scale and .62 for the heaviness scale.

For the lightness dimension, significant, and large, negative β values for the obstruents indicated that, compared to sonorants, these attracted ratings at the ‘not light’ end of the scale; once again, the effect on model fit was greater for stops than af/fricatives (Table 4) suggesting that stops played a greater role in defining the ‘not light’ end of the scale. Items rated as light were thus more likely to be associated with sonorants (as in Sidhu et al., 2022) such as /l/ and /n/ in /lonu/ (Table 3). Lightness was also associated with a preference for unvoiced consonants, these produced a positive β and a small improvement in the lightness model fit (Table 4), as illustrated by, for example, /f/ and /s/ in /fɛfi/ and /sεse/ (Table 3), consistent with the association with unvoiced fricatives in Sidhu et al. (2022).

For the place of articulation, there were significant negative β values for alveolar and post-alveolar/velar consonants in both consonant positions, indicating that these attracted ratings towards the ‘not light’ end of the scale, but resulting only in small changes to R^2^ (Table 4) Accordingly, lightness was more reflected in bilabial and labiodental consonants, such as those in /monu/ and /vevi/, respectively (Table 3).

The negative β for back rounded vowels indicated that these attracted ratings towards the ‘not light’ end of the scale and thus resulted in only a small change in R^2^ (Table 4). Thus, front unrounded vowels were more predictive of lightness (as in Walker & Parameswaran, 2019, and Sidhu et al., 2022) as shown in /fɪfe/ and /sεse/ (Table 3).

Pseudowords rated as heavy were more likely to contain obstruents, as shown by the large positive β values, although stops produced a greater improvement to the heaviness model fit than af/fricatives (Table 4). For voicing, unvoiced consonants produced a negative β in the heaviness scale, reflecting that, as reported above, these were predictive of lightness; consequently, these made only a small change in model fit (Table 4). Heaviness was thus associated with voiced consonants (consistent with Sidhu et al., 2022; Greenberg & Jenkins, 1966); these, together with the obstruents, are exemplified by, for example /b/, /g/, and /ʤ/in /bεge/ and /ʤoʤu/ (Table 3).

For the place of articulation, only post-alveolar/velar consonants, in both consonant positions, such as /ʤ/ above, and /g/ and /k/ in /gebi/ and /pɪke/ (Table 3), produced significant positive β values and small improvements to the heaviness model fit (Table 4).

Heaviness was also associated with back rounded vowels, for example /o/, /u/, and /ʊ/, in /ʤoʤu/ and /pʊpo/ (Table 3), rather than front unrounded vowels (as in Walker & Parameswaran, 2019, and Sidhu et al., 2022). Only high vowels in the second position produced a significant β but, since this was negative, it indicated that these were associated with the ‘not heavy’ end of the scale and resulted in only a minimal effect on model fit (Table 4).

### Arousal

For the arousal domain, R^2^ was .48 for the calming dimension and .36 for the exciting dimension (Table 4). Inter-rater reliability (Cronbach alpha) was .64 for the calming scale and .52 for the exciting scale.

For the calming dimension, there were large negative β values for obstruents, indicating that, compared to sonorants, these attracted ratings towards the ‘not calming’ end of the scale and resulted in moderate changes in R^2^ (Table 4). Accordingly, the sonorants, as in /mulu/ and /lonu/, were more associated with calming ratings (Table 3), consistent with Sidhu et al. (2022, and see also Aryani et al., 2020). There was a small, but significant, negative β for unvoiced consonants suggesting that, in addition to sonorants, voiced obstruents might also be predictive of calming ratings (Aryani et al., 2018), though none appeared in the ten most highly-rated pseudowords for the calming dimension (Table 3).

In terms of the place of articulation, there were significant negative β values for alveolar consonants in the second position only (with a minimal change to R^2^) and for post-alveolar/velar consonants in either position (with small changes to R^2^) indicating that these attracted ratings towards the ‘not calming’ end of the scale (Table 4). Thus, compared to the alveolar and post-alveolar/velar categories, bilabial/labiodental consonants were considered calming and, in keeping with the preference for sonorants, six of the ten highest-rated calming pseudowords contain the bilabial sonorant /m/ at least once (Table 3).

The back rounded vowels produced a significant positive β and a small improvement to the calming model fit (Table 4); thus, these were predictive of calming ratings, and all three of the rounded vowels in our phoneme set (Table 1) were present in /mumo/ and /mʊmo/ (Table 3), consistent with Sidhu et al. (2022). However, vowel height had no significant effect.

For the exciting dimension, there were significant positive β values for the obstruents, both stops and af/fricatives, each associated with small improvements to the exciting model fit (Table 4), and consistent with Sidhu et al. (2022, and see Aryani et al., 2020). These were exemplified by /t/ or /k/, and /z/ or /ʤ/, respectively, in /tɪke/ and /zɛʤi/ (Table 3). Consistent with Aryani et al. (2018), unvoiced, rather than voiced, consonants were predictive of exciting ratings, producing a significant positive β and a small improvement to R^2^ (Table 4), for example /t/ and /k/ as above. However, there was no significant effect of the place of articulation (Table 4), with the ten highest-rated pseudowords on the exciting scale containing bilabial /p/, alveolar /t/ and /z/, and post-alveolar/velar /ʤ/ and /k/ (Table 3).

Back rounded vowels showed a significant negative β, indicating that these were associated with the ‘not exciting’ end of the scale and a moderate improvement in the exciting model fit (Table 4). Accordingly, front unrounded vowels, as seen in /tɪke/ and /zɛʤi/ (Table 3) predicted perception of a pseudoword as sounding exciting (Sidhu et al., 2022). Note that these are also high vowels and, where these occurred in the first position, they produced a significant β and a small improvement in model fit (Table 4).

#### Brightness

For the brightness domain, R^2^ was .47 for the brightness dimension and .31 for the darkness dimension (Table 4). Inter-rater reliability (Cronbach alpha) was .63 for the brightness scale and .52 for the darkness scale.

For pseudowords rated as sounding bright, the manner of articulation for consonants was less important than their voicing, consistent with Newman (1933). The β values for the obstruents were not significant, but that for unvoiced consonants was significant and positive, producing a small improvement to the brightness model fit (Table 4). Accordingly, the unvoiced stops /t/ and /k/, and unvoiced af/fricatives /f/, /s/, and /ʧ/, occurred in nine of the ten brightest pseudowords, while the voiced /v/ and /z/ occurred only once and in the same item: /vizi/ (Table 3).

For place of articulation, there were small but significant effects for alveolar and post-alveolar/velar consonants in the first and second consonant positions respectively, for example, /s/ and /ʧ/ in /siʧi/ (Table 3). However, these made only minimal changes to R^2^ (Table 4).

Vowels were by far the largest influence on brightness ratings: back rounded vowels produced a large significant negative β indicating that they attracted ratings very much towards the ‘not bright’ end of the scale, and resulted in a large improvement to model fit (in fact, the second largest improvement across all domains) by defining what was considered ‘not bright’ and therefore perceived as sounding dark (Table 4). Consistent with this, all ten of the most highly-rated bright pseudowords contained only front unrounded vowels (see also Newman, 1933), for example /e/ and /i/, in /kike/ and /pite/ (Table 3). Note that /vizi/, although an outlier in containing voiced consonants (see above), also contains two front unrounded vowels. The effect of vowel height was not significant.

In contrast to the brightness dimension, both manner of articulation and voicing of consonants were important to the darkness dimension. There were significant positive β values for the obstruents although the resulting changes to R^2^ were small, albeit rather better for the af/fricatives than for stops (Table 4). Nonetheless, there was only one occurrence of af/fricatives in the ten highest-rated pseudowords – /vuʤo/ – although there were a further eight in the next ten items – while voiced stops occurred in seven of the top ten pseudowords (Table 3). The significant negative β for unvoiced consonants (Table 4) indicated that voiced consonants were more predictive of darkness ratings (consistent with Newman, 1933) and these are the only consonants to appear in the ten highest-rated items (Table 3).

The place of articulation had a limited influence in that the only significant effect was for post-alveolar/velar consonants in the first position and the resulting change to model fit was minimal (Table 4).

As would be expected from the brightness scale results, back rounded vowels were strongly predictive of darkness ratings (consistent with Newman, 1933) and 8 of the 10 most highly-rated dark pseudowords contained these, for example in /gubo/ (Table 3). Back rounded vowels produced a significant positive β and a moderate improvement to the darkness model fit; as with the brightness scale, vowel height had no influence on darkness ratings (Table 4).

#### Valence

The valence domain had the second worst fit between phonetic categories and ratings, with R^2^ of .4 for the good dimension and .28 for the bad dimension (Table 4). Inter-rater reliability (Cronbach alpha) was .54 for the goodness scale and .28 for the badness scale.

For goodness, there were significant negative β values for the obstruents indicating that, compared to sonorants, these attracted ratings towards the ‘not good’ end of the scale; both stops and af/fricatives produced moderate changes in R^2^ (Table 4). Pseudowords rated as good were thus more likely to be associated with sonorants than obstruents, as suggested by Adelman et al. (2018), for example, in /meni/ and /lunu/ (Table 3). There was also a small but significant influence of unvoiced consonants, although the effect on model fit was minimal (Table 4) and there were only 3 occurrences of unvoiced consonants in the first 30 items, all outside the top 10 most highly-rated items. This finding is not consistent with Aryani et al. (2018), who found that unvoiced consonants were rated as negatively valenced.

The place of articulation was not a significant predictor, with no significant β values; accordingly, both bilabial /m/ and /b/, and alveolar /l/ and /n/, appeared in the ten most highly-rated good pseudowords (Table 4).

The significant negative β for back rounded vowels indicated that these were rated as ‘not good’ compared to front unrounded vowels, and this was associated with a small improvement to the goodness model fit (Table 4). This was consistent with Körner & Rummer (2022), and exemplified by, for example, /lɛme/ and /bɪbi/ (Table 3).

For ratings on the badness scale, there were significant positive β values for the obstruents, with small and moderate improvements in model fit for stops and af/fricatives, respectively (Table 4). This is consistent with previous studies (Adelman et al., 2018; Uno et al., 2020) and exemplified by, for example, /d/ and /ʧ/ in /dudo/ and /foʧu/ (Table 3). The importance of fricatives for badness ratings is in line with Aryani et al. (2018) who reported that hissing sibilants (of which the fricatives /s/ and /z/ are examples) were associated with negative valence. There was no significant influence of voicing (Table 4): /dudo/and /foʧu/ contain voiced stops and unvoiced af/fricatives respectively (Table 3). As with the goodness ratings above, this is inconsistent with Aryani et al. (2018) who reported that positive and negative valence were reflected in voiced and unvoiced consonants, respectively.

In terms of the place of articulation, there were no significant effects other than for a negative β for alveolar consonants in the first position indicating that these contributed to ratings towards the ‘not bad’ end of the scale, and resulting in a small change to R^2^ (Table 4). This suggested that badness was more likely to be associated with the referent predictor, bilabial and labiodental consonants which did appear in the most highly-rated bad pseudowords, as for example /f/, /p/, and /v/ in /foʧu/, /pupo/, and /vʊvu/ (Table 3).

These examples also illustrate the significant influence of back rounded vowels (Table 4) on the perception of pseudowords as sounding bad, consistent with Körner & Rummer (2022); the effect on model fit was small.

#### Size

The size domain had the worst model fit, with R^2^ of .12 for the small dimension and .29 for the big dimension (Table 4). Nonetheless, there were some significant phonetic predictors. Inter-rater reliability (Cronbach alpha) was .48 for the smallness scale and .46 for the bigness scale.

In contrast to Sidhu et al. (2022), stops were preferred over sonorants for smallness ratings but these made only a small improvement to the model fit (Table 4). There was a similar result for unvoiced consonants (Table 4), illustrated by, for example /p/ and /t/ in /pete/ and /pɪte/ (Table 3; consistent with Winter & Perlman, 2021). There was no significant effect for the af/fricative category, contrasting with Preziosi & Coane (2017), who reported that fricatives were associated with small size.

For the place of articulation, there were significant negative β values for post-alveolar/velar consonants in either position indicating that these were rated towards the ‘not small’ end of the scale (Table 4). By comparison then, bilabial and labiodental consonants were more likely to be associated with smallness as, for example, /p/ and /f/ in /pete/ and /fɪse/ (Table 3).

Vowels were not a significant predictor of smallness, again in contrast to previous studies (Sapir, 1929; Auracher, 2017; Sidhu et al., 2022), where smallness was predicted by front vowels.

Obstruents were again preferred over sonorants for pseudowords rated as sounding big, but here there were significant positive β values for both stops and af/fricatives, each producing small improvements to the model fit (Table 4). This is consistent with previous studies (Sidhu et al., 2022; Preziosi & Coane, 2017) and illustrated by, for example, /ʤ/ and /b/ in /ʤɪʤe/ and /bɛge/ (Table 3).

For the place of articulation, there were significant positive β values for post-alveolar/velar consonants (Table 4) which were thus reasonable predictors of bigness ratings and in either consonant position: /ʤ/, for example appears in both positions in /ʤɪʤe/ (Table 3).

Consistent with prior work (Sapir, 1929; Auracher, 2017; Sidhu et al., 2022), bigness was significantly associated with back rounded vowels (Table 4), for example /ʊ/ and /o/ in /gʊdo/ (Table 3). There was a small effect of vowel height in that high vowels in the second position produced a significant negative β and a small change to R^2^ (Table 4), suggesting that, compared to high vowels, mid-height vowels were preferred in this position, as is also illustrated by /gʊdo/.

### Discussion

Sonorants were associated with roundedness (Köhler, 1929, 1947; Ramachandran & Hubbard, 2011; McCormick et al., 2015; Sidhu et al., 2022), softness (Sidhu et al., 2022; note that this contrasts with Sakamoto & Watanabe, 2018, who found that bilabial stops also indicated softness), lightness (Sidhu et al., 2022), calmness (Sidhu et al., 2022), and goodness (Adelman et al., 2018). Obstruents were associated with pointedness (Köhler, 1929, 1947; Ramachandran & Hubbard, 2011; McCormick et al., 2015; Sidhu et al., 2022), hardness (Sakamoto & Watanabe, 2018; Sidhu et al., 2022), heaviness (Greenberg & Jenkins, 1966), exciting-ness (Aryani et al., 2020; Sidhu et al., 2022), and badness (Adelman et al., 2018; Uno et al., 2020).

Voiced consonants were primarily associated with heaviness (Sidhu et al., 2022), darkness (Newman, 1933), and large size (Westbury et al., 2018) but there were also significant, if weaker, associations with roundedness (Köhler, 1929, 1947; Ramachandran & Hubbard, 2001; McCormick et al., 2015; Sidhu et al., 2022), and calmness (Aryani et al., 2020). Unvoiced consonants were most strongly associated with pointedness (McCormick et al., 2015; Sidhu et al., 2022), lightness (Sidhu et al., 2022), exciting-ness (Aryani et al., 2018), brightness (Newman, 1933) and small size (Winter & Perlman, 2021), with weaker associations to hardness (Sakamoto & Watanabe, 2018), and goodness (see Mooshammer et al., 2023).

For vowels, consistent with previous studies, back/rounded and front/unrounded vowels were respectively associated with roundedness and pointedness (Köhler, 1929, 1947; Ramachandran & Hubbard, 2001; McCormick et al., 2015), heaviness and lightness (Walker & Parameswaran, 2019; Sidhu et al., 2022), calming and exciting (Sidhu et al., 2022), darkness and brightness (Newman, 1933), and badness and goodness (Körner & Rummer, 2022). Back/rounded vowels were also associated with softness, a new finding and in contrast to Sidhu et al. (2022) who found no association between vowels and either hardness or softness. Consistent with earlier findings (Sapir, 1929; Auracher 2017; Sidhu et al., 2022), back/rounded vowels were associated with large size but we did not find the corresponding association reported by these authors between front/unrounded vowels and small size.

In terms of the place of articulation, bilabial/labiodental consonants were associated with roundedness (see also Westbury et al., 2018), softness, lightness, and calmness. Both alveolar and postalveolar/velar consonants were associated with pointedness (see also Westbury et al., 2018), hardness, and brightness, while post-alveolar/velar consonants were associated with heaviness and also large size. The functional importance of the place of articulation is that it alters the length of the vocal tract involved in producing the sound and thus has acoustic consequences (Reetz & Jongman, 2020). In brief, bilabial/labiodental consonants result in low-frequency sounds while alveolar consonants result in high-frequency sounds, and intermediate frequencies arise from post-alveolar/velar consonants (for a more detailed treatment of the acoustic consequences of the place of articulation, see the acoustic analysis below). In this respect, it is interesting that roundedness was associated with the low-frequency bilabial/labiodental consonants while pointedness was associated with the relatively higher-frequency alveolar and post-alveolar/velar consonants. These frequency differences match those for vowel sounds associated with rounded/pointed pseudowords (Knoeferle et al., 2017) and visual shapes (Parise & Pavani, 2011). Similarly, the association of brightness with the relatively higher-frequency alveolar and post-alveolar/velar consonants is consistent with a previous finding that pseudowords are produced with a higher fundamental frequency for brighter, compared to darker, colors (Tzeng et al., 2018). Other associations, reported here for the first time, make intuitive sense; for example, that low-frequency bilabial/labiodental consonants sound calming and soft, while the relatively higher-frequency alveolar and post-alveolar/velar consonants sound hard.

For vowel articulation, backness/rounding was an important contributor to almost every domain as described above. Vowel height, however, was relatively unimportant, being non-significant for 4 of the 16 scales, and with predominantly small beta coefficients of ≤ .1 for the rest (Table 4), although it must be noted that there were no low vowels in the phoneme set, which contained only high and mid-height vowels (Table 1).

Comparing across domains, ratings were best predicted by collections of phonetic categories that were relatively specific to each domain. For example, in the shape and weight domains, one scale (rounded or light) was associated with sonorants while the other (pointed or heavy) was associated with obstruents. But, while they were broadly similar in terms of consonants, these scales differed in their vowel associations: back vowels for the rounded and heavy scales, and front vowels for the pointed and light scales (see Table 4). However, the hardness and size domains showed more differences: hardness was indicated by obstruents and unvoiced consonants but not vowels, while softness was indicated by sonorants (and so unvoiced consonants were not relevant), and by back rounded vowels. In the size domain, bigness was associated with obstruents, voiced consonants, and back rounded vowels, while smallness was indicated only by stops and unvoiced consonants, not by af/fricatives or vowels. Even positive correlations in ratings across domains were no guarantee of similarity in the related phonetic categories. On the one hand, the positively correlated rounded/calming and pointed/exciting scales were reflected in common phonetic categories: sonorants, voiced consonants, and back rounded vowels for rounded/calming, and obstruents, unvoiced consonants, and front unrounded vowels for pointed/exciting. By contrast, although the soft/light and hard/heavy scales were both positively correlated and both relied on the same consonants, the ratings combined with vowels differently: back rounded for softness and front unrounded for lightness, while hardness was not reflected in vowels at all and heaviness was associated with back rounded vowels. Finally, we should note that the graded nature of the ratings suggests that, in addition to the individual phonetic components of each pseudoword, participants’ judgments likely involved processing/analysis at the segment or even whole-item level (McCormick et al., 2015; see also Thompson & Estes, 2011).

Adjusted R^2^ values represent an estimate of the reduction in predictive power had the model been derived from the population from which the sample was taken (Field, 2018); as such, the smaller the difference between the adjusted R^2^ value and R^2^, the more generalizable the model is likely to be. Across all sixteen scales, the maximum such difference was .02 (for the softness, badness, and bigness scales, see Table 4), however, while this is worth noting for those scales that had a good model fit (e.g. rounded, pointed, roughness, and hardness), it is less informative for those scales where the model fit was poor to start with, e.g. smallness (Table 4).

Finally, in many cases, predictors had opposing effects on the opposing dimensions of a particular domain. For example, obstruents were significantly positively weighted on the pointed, roughness, hardness, heaviness, exciting, and badness scales, but significantly negatively weighted on the opposing rounded, smoothness, softness, lightness, calming, and goodness scales (Table 4). However, this was not always the case: for example, while voicing was positively weighted on the roughness scale, it was non-significant for the smoothness scale, and while back rounded vowels were positively weighted for softness, there was no significant effect on the hardness scale (Table 4). Thus, it was not always the case that antonym pairs and their associated phonetic categories had symmetric relationships to each other (see Westbury et al., 2018; Monaghan & Fletcher, 2019), i.e., a rating of ‘not rough’ did not necessarily equate to ‘smooth’. This is a caution against using bipolar scales where one cannot be sure that the same decision criteria underlie ratings at each end of such scales (see Connell & Lynott, 2012).

## PHONEME ANALYSIS

### Introduction

Although the phonetic category analysis is useful in establishing which class(es) of phonemes may be iconically related to roundedness, pointedness, darkness, etc., it is not necessarily the case that all members of a class contribute equally. Therefore, analysis at the level of individual phonemes, rather than effectively averaging across a whole class, might result in better models of iconicity (e.g., Monaghan & Fletcher, 2019). Additionally, the phonetic category analysis above hints at positional effects, i.e., it might matter where a phoneme occurs within a word. For example, on the hardness and brightness scales, alveolar consonants only influenced ratings in the second consonant position, and vowels only influenced ratings on the pointed and roughness scales in the first vowel position (Table 4), although a caveat in relation to vowels is that 3 of the 7 used here cannot appear in the second vowel position anyway because they do not appear in word-final positions in English (see Table 1).

Westbury et al. (2018) analyzed 32 consonants and 17 vowels (a complete list was not provided but see their Table 2) that could occur in any ‘legal’ position within 7996 pseudowords of between 3 and 8 letters. However, while phonetic category and phoneme analyses were carried out, positional effects were not considered. Monaghan & Fletcher (2019) report a more restricted study of 40 pseudowords that investigated only initial consonants and only stops (voiced /b/, /d/, /g/; unvoiced /p/, /t/, /k/) and fricatives (voiced /v/, /z/; unvoiced /f/, /s/); thus sonorants, affricates, and vowels were not considered. Winter & Perlman (2021) investigated real words and 36 phonemes occurring in 52 English size adjectives. Finally, Greenberg & Jenkins (1966) rated individual phonemes – selected consonants and vowels with no word/pseudoword context.

**Table 2:**
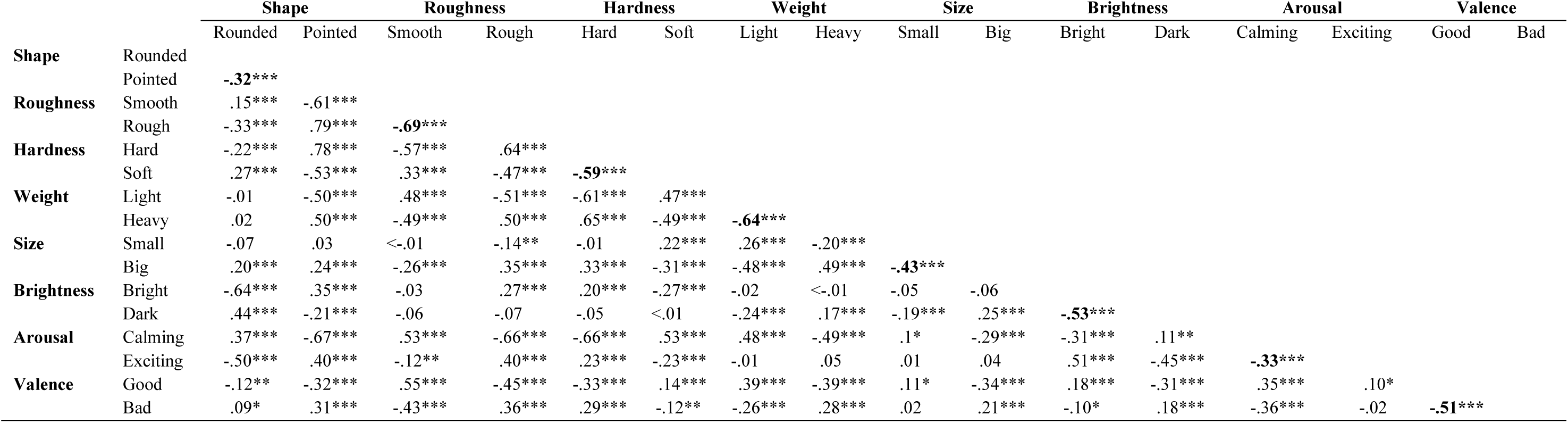
Ratings correlations within and between domains. Within each domain, the two scales were significantly negatively correlated (shown in **bold**), indicating that they adequately reflected the opposing dimensions. Degrees of freedom = 535 in all cases. * p < .05, ** p < .01, *** p < .001.

|  |  | Shape |  | Roughness |  | Hardness |  | Weight |  | Size |  | Brightness |  | Arousal |  | Valence |  |
| --- | --- | --- | --- | --- | --- | --- | --- | --- | --- | --- | --- | --- | --- | --- | --- | --- | --- |
|  |  | Rounded | Pointed | Smooth | Rough | Hard | Soft | Light | Heavy | Small | Big | Bright | Dark | Calming | Exciting | Good | Bad |
| Shape | Rounded |  |  |  |  |  |  |  |  |  |  |  |  |  |  |  |  |
|  | Pointed | **-.32\*\*\*** |  |  |  |  |  |  |  |  |  |  |  |  |  |  |  |
| Roughness | Smooth | .15*** | -.61*** |  |  |  |  |  |  |  |  |  |  |  |  |  |  |
|  | Rough | -.33*** | .79*** | **-.69\*\*\*** |  |  |  |  |  |  |  |  |  |  |  |  |  |
| Hardness | Hard | -.22*** | .78*** | -.57*** | .64*** |  |  |  |  |  |  |  |  |  |  |  |  |
|  | Soft | .27*** | -.53*** | .33*** | -.47*** | **-.59\*\*\*** |  |  |  |  |  |  |  |  |  |  |  |
| Weight | Light | -.01 | -.50*** | .48*** | -.51*** | -.61*** | .47*** |  |  |  |  |  |  |  |  |  |  |
|  | Heavy | .02 | .50*** | -.49*** | .50*** | .65*** | -.49*** | **-.64\*\*\*** |  |  |  |  |  |  |  |  |  |
| Size | Small | -.07 | .03 | <-.01 | -.14** | -.01 | .22*** | .26*** | -.20*** |  |  |  |  |  |  |  |  |
|  | Big | .20*** | .24*** | -.26*** | .35*** | .33*** | -.31*** | -.48*** | .49*** | **-.43\*\*\*** |  |  |  |  |  |  |  |
| Brightness | Bright | -.64*** | .35*** | -.03 | .27*** | .20*** | -.27*** | -.02 | <-.01 | -.05 | -.06 |  |  |  |  |  |  |
|  | Dark | .44*** | -.21*** | -.06 | -.07 | -.05 | <.01 | -.24*** | .17*** | -.19*** | .25*** | **-.53\*\*\*** |  |  |  |  |  |
| Arousal | Calming | .37*** | -.67*** | .53*** | -.66*** | -.66*** | .53*** | .48*** | -.49*** | .1* | -.29*** | -.31*** | .11** |  |  |  |  |
|  | Exciting | -.50*** | .40*** | -.12** | .40*** | .23*** | -.23*** | -.01 | .05 | .01 | .04 | .51*** | -.45*** | **-.33\*\*\*** |  |  |  |
| Valence | Good | -.12** | -.32*** | .55*** | -.45*** | -.33*** | .14*** | .39*** | -.39*** | .11* | -.34*** | .18*** | -.31*** | .35*** | .10* |  |  |
|  | Bad | .09* | .31*** | -.43*** | .36*** | .29*** | -.12** | -.26*** | .28*** | .02 | .21*** | -.10* | .18*** | -.36*** | -.02 | **-.51\*\*\*** |  |

Our predictions for the phoneme analysis are based on these prior phoneme analyses with positional effects inferred from our phonetic category analysis as mentioned above. Note that for Greenberg & Jenkins (1966) we used ratings from their Table 3 for consonants and Table 13 for vowels as these were drawn from substantially larger sample sizes than their initial reports (note too, that the domains differed slightly between consonants and vowels in any case). There was no statistical comparison between phonemes, so we took only those phonemes rated above or below, but not at, the mid-point of 3.5 on their 7-point scale.

**Table 3:**
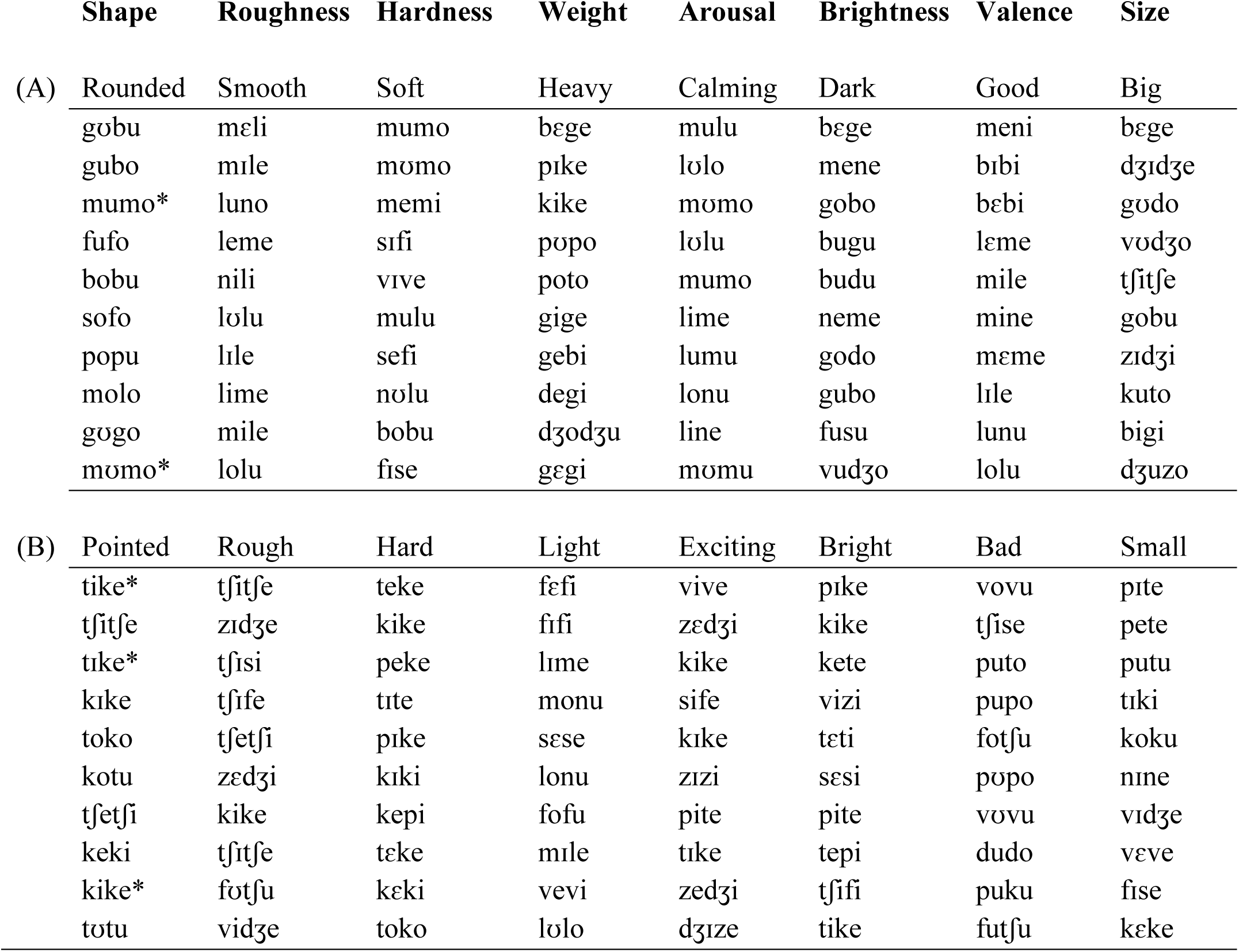
The 10 highest-rated pseudowords for each scale in each domain. Scales are grouped by approximate phonetic similarity: A, predominantly voiced consonants and both rounded and unrounded vowels; B, predominantly unvoiced consonants and unrounded vowels. Most items were unique in each group – A: 45 (64.3%); B: 47 (67.1%). Shape pseudowords marked * were also in the top 10 for the shape ratings obtained by McCormick et al. (2015).

#### Shape

We predicted that roundedness would be associated with /b/ and /m/ (Westbury et al., 2018; Greenberg & Jenkins, 1966) together with /l/, /n/, /d/, /p/, and /s/ (Greenberg & Jenkins, 1966); however, only the sonorants /m/, /l/, and /n/ would be consistent with our phonetic category analysis (Fig. 1A; Supplementary Table 4). We would expect positional effects for bilabial consonants like /m/ at both C1 and C2, and for mid-height vowels, such as /o/, only at V2 (Fig. 1A; Supplementary Table 4). We expected pointedness to be associated with /k/ (Westbury et al., 2018; Greenberg & Jenkins, 1966) and /z/ (Greenberg & Jenkins, 1966), both of which would be consistent with the phonetic category analysis (Fig. 1A; Supplementary Table 4). There should also be positional effects for alveolar and velar consonants like /z/ and /k/, respectively, at both C1 and C2, while high vowels, such as /i/, would only have a significant effect at V1 (Fig. 1A; Supplementary Table 4).

**Figure 1A:**
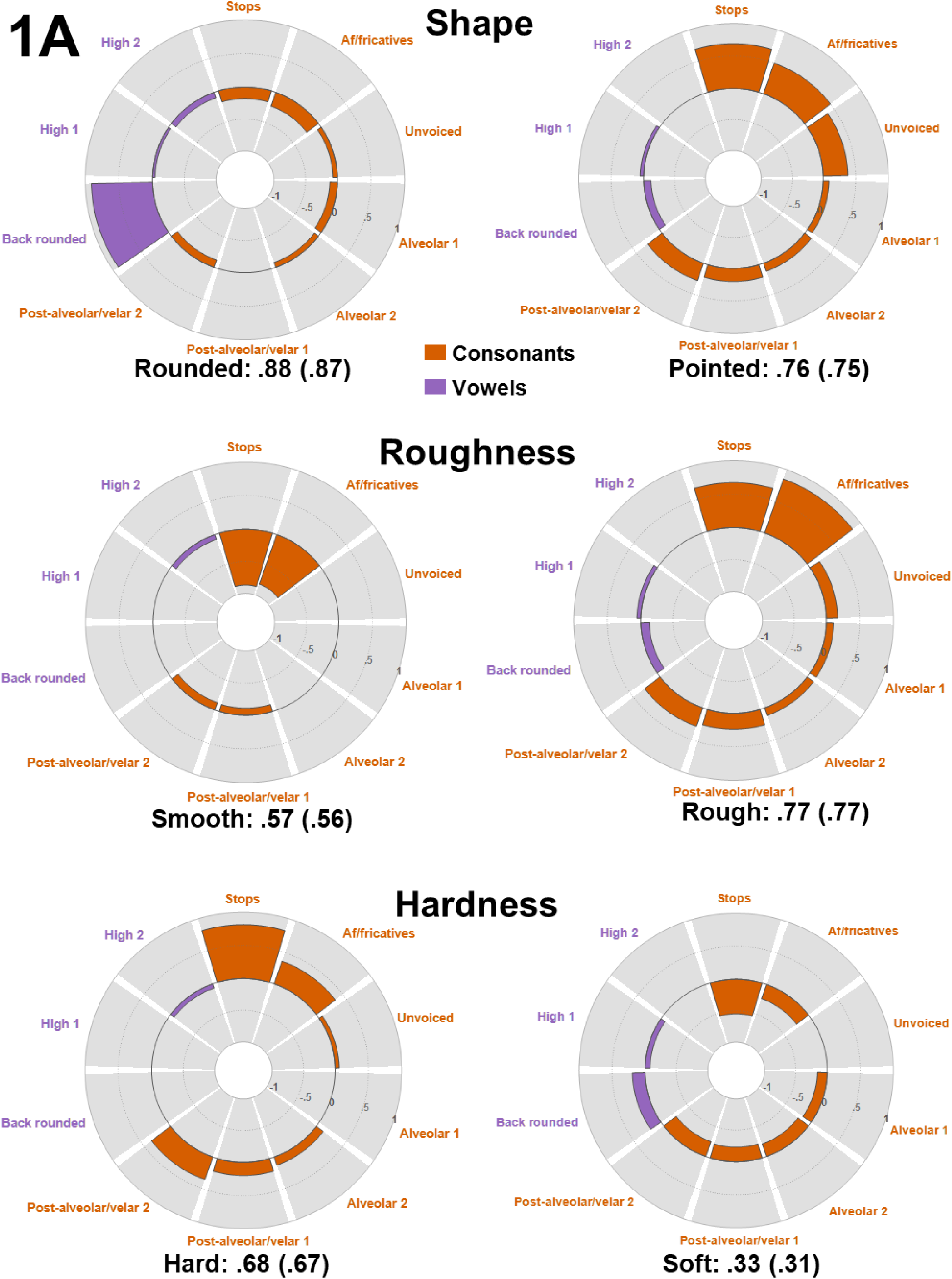
Phonetic category regression results for the opposing scales of the shape, roughness, and hardness domains. The inner and rings show negative and positive standardized beta coefficients, respectively, represented as percentage area of the ‘wedge’; only significant coefficients are shown. R^2^ and (adjusted R^2^) values are provided for each scale. 1 and 2 refer to the first and second consonant or vowel position in the CVCV pseudowords. The reference categories are not graphed but, for stops and af/fricatives, the reference is sonorants; for unvoiced consonants, the reference is voiced consonants; for place of articulation, the reference is bilabial/labiodental; for back rounded vowels, the reference is front unrounded vowels; and for high vowels, the reference is mid-height vowels. For the complete results, see Supplementary Table 4.

#### Roughness

We predicted that smoothness would be associated with the sonorants /m/ and /l/, the fricative /s/, the front unrounded vowel /ɪ/ and the back rounded vowel /u/ (Greenberg & Jenkins, 1966); however, only the sonorants would be consistent with our phonetic category analysis (Fig. 1A; Supplementary Table 4). Additionally, we predicted that bilabial consonants like /m/ would show effects at both C1 and C2, while mid-height vowels would so only at V2 (Fig. 1A; Supplementary Table 4). By contrast, roughness would be associated with the stops /b/, /p/, /d/, and /k/, together with the fricative /z/, and the front unrounded vowels /ε/ and /i/ (Greenberg & Jenkins, 1966), all of which would be consistent with the phonetic category analysis (Fig. 1A; Supplementary Table 4). Additionally, we expected that alveolar consonants like /d/ and /z/ and the velar consonant /k/ would show significant effects at both C1 and C2, while high vowels /i/ would do so only at V1.

#### Hardness

Based on the prior phoneme analyses, we expected hardness to be associated with the consonants/b/, /d/, (Greenberg & Jenkins, 1966), /g/, /z/ (Monaghan & Fletcher, 2019), and /k/ (Monaghan & Fletcher, 2019; Greenberg & Jenkins, 1966), and the front unrounded vowels /i/, /ɪ/, and /ε/ (Greenberg & Jenkins, 1966). However, while all the above consonants would be consistent with the phonetic category analysis, we did not find any effect for vowel frontness (Fig. 1A; Supplementary Table 4). For positional effects, we could expect that the alveolar consonants /d/ and /z/ would show significant effects at C2, the postalveolar /g/ and velar /k/ would show significant effects at both consonant positions. (Fig. 1A; Supplementary Table 4). Furthermore, although we found no overall effect of vowel frontness, we did find an effect for mid-height vowels at V2 that might implicate /e/ and /ε/ (Fig. 1A; Supplementary Table 4). We expected softness to be associated with /m/, /n/, /l/, /p/, /z/ (Greenberg & Jenkins, 1966), /f/ (Monaghan & Fletcher, 2019), and /s/ (Monaghan & Fletcher, 2019; Greenberg & Jenkins, 1966) for the consonants, and with /u/ for the vowels (Greenberg & Jenkins, 1966). However, among these, only the sonorants /m/, /n/, /l/ and the back rounded vowel /u/ would be consistent with the phonetic category analysis (Fig. 1A; Supplementary Table 4). For positional effects, the phonetic category analysis would only predict an effect for bilabial consonants, at both C1 and C2, together with an effect for mid-height vowels at V1 only (Fig. 1A; Supplementary Table 4).

#### Weight

Among the consonants, we predicted associations with lightness for the sonorants /m/ and /l/, and the fricatives /s/ and /z/, and or the vowels /i/, /ɪ/, /ε/, and /u/ (Greenberg & Jenkins, 1966).; however, only the sonorants and the front unrounded vowels would be consistent with the phonetic category findings (Fig. 1B; Supplementary Table 4). Additionally, we could expect positional effects for bilabial consonants only, albeit at both C1 and C2; and for high vowels, like /i/ and /ɪ/, only and restricted to V2 (Fig. 1B; Supplementary Table 4). For heaviness, we could predict strong associations with /n/, /b/, /p/, /d/ and /k/ (Greenberg & Jenkins, 1966) but, among these, only the stops would be consistent with the phonetic category findings (Fig. 1B; Supplementary Table 4). For positional effects, our phonetic analysis would only predict effects for the velar consonant /k/, albeit at both C1 and C2, and for the mid-height vowel /o/ only and restricted to V2. (Fig. 1B; Supplementary Table 4).

**Figure 1B:**
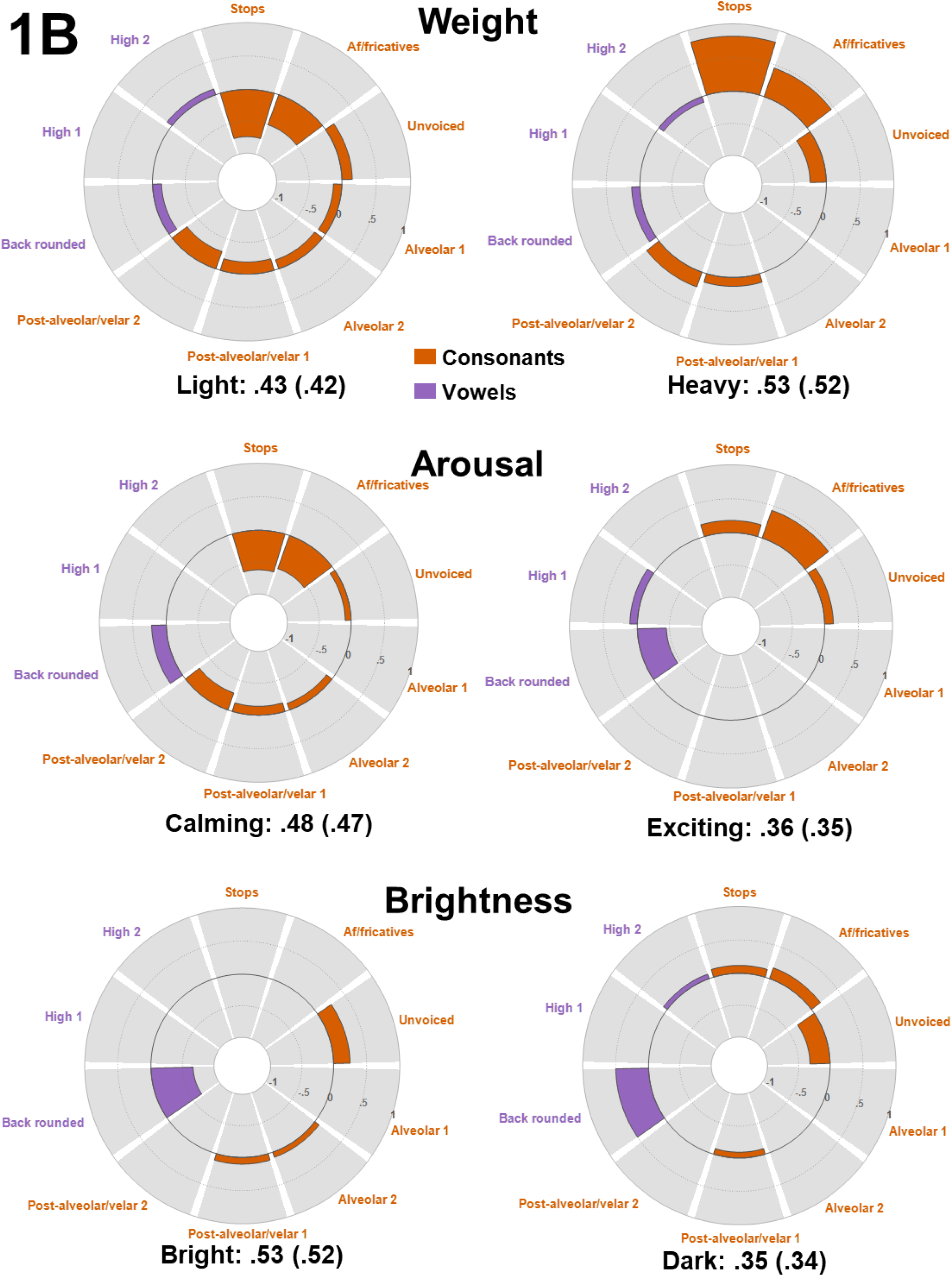
Phonetic category regression results for the opposing scales of the weight, arousal, and brightness domains. Interpretation as for Fig. 1A.

#### Arousal

For positional effects, for the calming scale, we would expect that bilabial/labiodental consonants would have significant effects at both C1 and C2, but we have no specific prediction for vowels (Fig. 1B; Supplementary Table 4). For the exciting scale, by contrast, we have no specific prediction for consonants, but we would expect significant effects for high vowels at V1 only (Fig. 1B; Supplementary Table 4).

#### Brightness

Following Greenberg & Jenkins (1966), we predicted that brightness would be associated with the consonants /m/, /l/, /b/, /p/, /k/, /s/, /z/, and the vowels /ɪ/, /i/, and /ε/. However, only /p/, /k/, and /s/ would be consistent with the phonetic category findings, because these are the only unvoiced stops and fricatives; by contrast, /ɪ/, /i/, and /ε/ would all be consistent because they are all front unrounded vowels (Fig. 1B; Supplementary Table 4). For positional effects, the phonetic category analysis suggests that velar /k/ would be significant at C1, while at C2 the alveolar consonants /l/, /s/, and /z/ would be significant (Fig. 1B; Supplementary Table 4); we had no predictions for the vowels. We predicted that darkness would be associated with /d/ (Greenberg & Jenkins, 1966) and, as a voiced stop, this would be consistent with the phonetic category analysis (Fig. 1B; Supplementary Table 4). For positional effects, the phonetic category analysis suggested an effect for bilabial/labiodental consonants at C1 only (although this would exclude the alveolar /d/) and an effect for high vowels at V2 only (Fig. 1B; Supplementary Table 4).

#### Valence

Again following Greenberg & Jenkins (1966), we predicted that goodness would be associated with the consonants /m/, /b/,/p/, /d/, /n/, /l/, and /s/, together with the vowels /i/, /ε/, and /u/. The sonorants /m/, /n/, /l/ and the front vowels /i/ and /ε/ would be consistent with the phonetic category analysis (Fig. 1C; Supplementary Table 4). This analysis also suggested that positional effects would only occur for mid-height vowels such as /ε/, and only at V2. We expected that badness would be associated with /ɪ/ (Greenberg & Jenkins, 1966) but that there would be no positional effects vowels; rather, the phonetic category analysis suggested an effect for bilabial/labiodental consonants at C1 only (Fig. 1C; Supplementary Table 4).

**Figure 1C:**
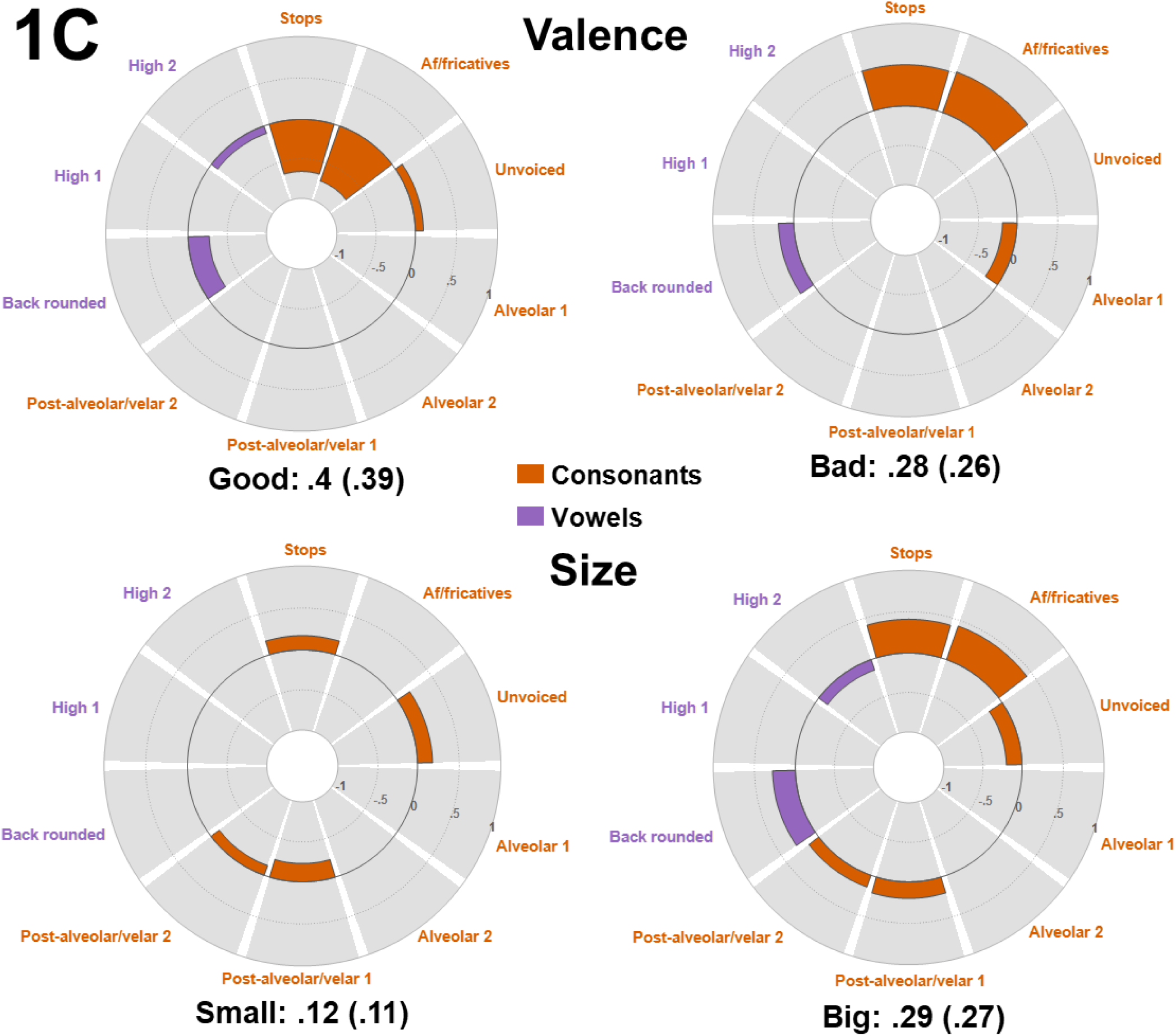
Phonetic category regression results for the opposing scales of the valence and size domains. Interpretation as for Fig. 1A.

#### Size

For the consonants, we predicted that smallness would be associated with /z/, /m/, /n/, /l/, and /ε/ (Greenberg & Jenkins, 1966), /p/ and /s/ (Monaghan & Fletcher, 2019; Greenberg & Jenkins, 1966), /t/ (Monaghan & Fletcher, 2019; Winter & Perlman, 2021), and /z/ (Greenberg & Jenkins, 1966). We predicted that it would also be associated with the front unrounded vowels /ε/ (Greenberg & Jenkins, 1966), /i/, and /ɪ/ (Winter & Perlman, 2021; Greenberg & Jenkins, 1966). However, only the unvoiced stops /p/ and /t/ would be consistent with the phonetic category analysis (Fig. 1C; Supplementary Table 4). For positional effects, we expected significant effects for bilabial/labiodental consonants, like /p/ and /t/ respectively, at both C1 and C2 (Fig. 1C; Supplementary Table 4). Following prior work, we predicted that bigness would be associated with the consonants /b/ (Westbury et al., 2018; Monaghan & Fletcher, 2019; Greenberg & Jenkins, 1966), /d/, /k/ (Greenberg & Jenkins, 1966), /g/ (Westbury et al., 2018; Monaghan & Fletcher, 2019), and /z/ (Monaghan & Fletcher, 2019), but only /u/ of the vowels (Greenberg & Jenkins, 1966). The voiced stops /b/, /d/, /g/ and fricative /z/, together with the back rounded vowel /u/ would be consistent with the phonetic category analysis (Fig. 1C; Supplementary Table 4). For positional effects, the phonetic category analysis suggested an effect for post-alveolar/velar consonants, e.g., /g/ and /k/ respectively, at both C1 and C2, but an effect for mid-height vowels at V2 only (Fig. 1C; Supplementary Table 4).

### Statistical analysis

As for the phonetic category regression above, we ran a forced-entry linear regression for each of the 2 scales in each of the 8 domains. There were four categories: the first and second consonant (C1, C2) and the first and second vowel (V1, V2). C1 and C2 each had 15 levels: b (the reference predictor), the sonorants /l/, /m/, /n/; the stops /d/, /g/ (voiced), and /k/, /p/, /t/ (unvoiced); the af/fricatives /v/, /z/, /ʤ/ (voiced), and /f/, /s/, /ʧ/ (unvoiced). V1 had 7 levels: /o/ (the reference predictor), /u/, /ʊ/ (rounded) and /e/, /ɛ/, /i/, /ɪ/ (unrounded). V2 had only 4 levels: /o/ (the reference predictor), /u/, /e/, and /i/, because the other vowels do not appear in word-final positions.

As for the phonetic category regression above, positive β values indicate that, compared to the reference predictor, the phoneme attracted ratings towards the positive end of the scale, e.g., ‘very rounded’, while negative β values indicate that the phoneme attracted ratings towards the negative end of the scale, e.g., ‘not rounded’ and therefore that the reference predictor was positively related instead.

SPSS automatically excluded /f/, /k/, and /n/ in the first, and /v/ in the second, consonant position together with /e/ in the second vowel position as the tolerance values for these were zero. Multicollinearity testing on the remaining phonemes showed that there were no tolerance values below .2 (Menard, 1995), with observed values ranging from .28 to .7, and that VIF values ranged from 1.42 to 3.56, well below the threshold value of 10 recommended by Myers (1990: Supplementary Table 4). In addition, the mean VIF of 2.44 was close to 1, as recommended by Bowerman & O’Connell (1990), although slightly greater than observed in the phonetic category regression. Thus, we concluded that multicollinearity was not a problem. There were no Durbin-Watson test values < 1 or > 3 (Field, 2018), ranging from 1.82 to 2.35 (Supplementary Table 5), thus the assumption of independent errors was met. Visual inspection of histograms and Q-Q plots for standardized residuals indicated that these were normally distributed for all scales. Plots of standardized residuals against standardized predicted values indicated that the assumption of homoscedasticity was also met for all scales.

### Results

#### Shape

Consistent with the phonetic category analysis, the shape domain had the best model fit between individual phonemes and ratings for both shape dimensions with R^2^ values of .83 for roundedness and .9 for pointedness (Figure 2A; Supplementary Table 7).

**Figure 2A:**
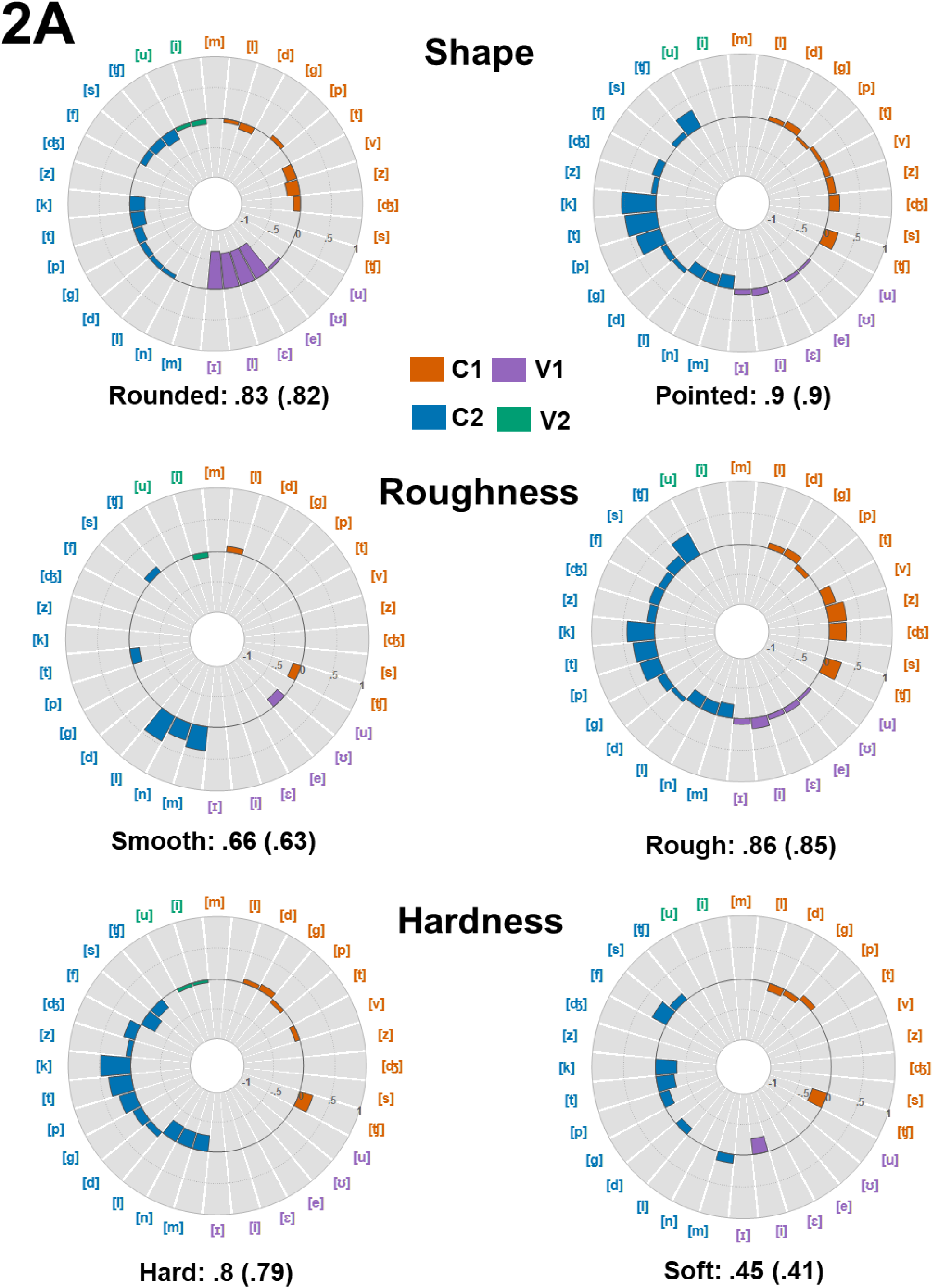
Phonemic regression results for the opposing scales of the shape, roughness, and hardness domains. C1, 2 and V1, 2 refer to consonants and vowels in the first or second position in the CVCV pseudowords. The reference phonemes are not graphed but, for C1 and C2, the reference phoneme is /b/; for V1 and V2, the reference phoneme is /o/. Otherwise, interpretation is as for Figure 1A. For the complete results, see Supplementary Table 7.

Vowels were by far the largest influence on roundedness ratings. Compared to the referent predictor, the back rounded vowel /o/, all the front unrounded vowels in the V1 position were associated with ratings decreasing towards ‘not rounded’ (Figure 2A; Supplementary Table 7), thus rounded vowels were associated with roundedness and, indeed, /o/ appears in 9 of the 10 highest-rated rounded pseudowords (Table 3). Collectively and individually, the addition of front unrounded vowels to the model produced the largest changes in R^2^, defining roundedness by its opposite. Interestingly, these effects were much more limited in the V2 position (Figure 2A; Supplementary Table 7). The only consonant phoneme that was positively associated with roundedness was the unvoiced stop /p/, but the effect was small, the change to R^2^ was minimal, and it appears in only one highly rated pseudoword /popu/ (Table 3). There were negative βs for all other significant consonants (Figure 2A; Supplementary Table 7). Compared to the referent predictor, the voiced stop /b/, there was a small negative effect for /l/ in both C1 and C2 consonant positions, suggesting that /b/ was likely associated with roundedness: this appears in 3 of the 10 highest-rated rounded pseudowords, for example, /bobu/ (Table 3). The af/fricatives were either non-significant or strongly associated with decreases in ratings towards ‘not rounded’ in both consonant positions, and the same was largely true for stops especially in the C2 position.

By contrast, vowels had little influence on pointedness ratings. In the V1 position, there were small positive effects for the back rounded vowel /ʊ/ and the front unrounded vowels /e/, /i/, and /ɪ/ (Figure 2A; Supplementary Table 7), indicating that these were rated as pointed, for example, /tʊtu/, keki, and //tike/ (Table 3) but there were no significant vowel effects in the V2 position. Instead, pointedness was indicated more by consonants. The strongest drivers of pointedness ratings were the unvoiced stops /p/, /t/, and /k/ in the C2 position and these produced the largest improvements to the model fit (Figure 2A; Supplementary Table 7 and see examples above). In both C1 and C2 positions, the voiced stops /d/ and /g/ were positively associated with pointedness, as were voiced af/fricatives, although the gains in model fit were relatively modest (Figure 2A; Supplementary Table 7). The unvoiced affricate /ʧ/ also increased ratings towards ‘very pointed’ in both consonant positions, as in /ʧeʧi/. There were significant negative effects for the sonorants in the C2 position, and these did not appear in any of the pseudowords rated as most pointed.

#### Roughness

The roughness domain had the next best fit between phonemes and ratings with R^2^ of .66 for the smoothness dimension and .86 for the roughness dimension (Figure 2A; Supplementary Table 7).

Smoothness ratings were driven almost entirely by the sonorants in the C2 position (Figure 2A; Supplementary Table 7), although they occur in both consonant positions in the ten highest-rated smooth pseudowords (Table 3). There were only two other, smaller, positive influences, for the sonorant /l/ at C1 and the unvoiced fricative /s/ at C2. There were small negative effects for the unvoiced affricate /ʧ/ at C1 and the unvoiced stop /t/ at C2, indicating that both were associated with ratings of ‘not smooth’. There were only two significant effects for vowels and both were negative: the rounded vowel /ʊ/ at V1 and the unrounded vowel /i/ at V2 both contributed to ‘not smooth’ ratings (Figure 2A; Supplementary Table 7), although front unrounded vowels predominate in the pseudowords with the highest smoothness ratings (see Table 3).

By contrast, roughness ratings were almost entirely driven by the af/fricatives in both consonant positions (Figure 2A; Supplementary Table 7), as exemplified by /ʧiʧe/ and /zεʤi/ (Table 3), with additional effects for voiced stops at C1 and both voiced and unvoiced stops at C2. There were relatively large negative effects for the sonorants at C2, indicating that these influenced ‘not rough’ ratings, consistent with their prominence for the smoothness dimension (Figure 2A; Supplementary Table 7). There was also a small negative effect for /p/ at C1, but it does not appear in the ten highest-rated items on either the rough or the smooth scales. In a further contrast to the smoothness dimension, vowels featured prominently in roughness ratings: all four unrounded vowels and the rounded vowel /ʊ/ contributed positively to roughness ratings, as can be seen from Figure 1A and Supplementary Table 4, but only at V1 – there were no significant effects for any vowel at V2 (Figure 2A; Supplementary Table 7).

#### Hardness

The hardness domain had the third best model fit with R^2^ of .8 for the hardness dimension but .45 for the softness dimension (Figure 2A; Supplementary Table 7).

The largest positive effects for hardness were for the affricate /ʧ/ in the C1 position and all three unvoiced stops in the C2 position (Figure 2A; Supplementary Table 7); the latter appear in all ten of the pseudowords with the highest hardness ratings for example, /peke/ and /kepi/ (Table 3). There were generally smaller positive effects for the voiced stops in both consonant positions, but these did not appear in the ten highest-rated items. Similarly, the voiced af/fricatives /z/ and /ʤ/ were positively associated with hardness, but only when they occurred in the C2 position (Figure 2A; Supplementary Table 7) and, again, these did not occur in the ten highest-rated items (Table 3). There were large negative effects for the sonorants in the C2 position (the C1 position was non-significant), indicating that compared to the referent, the voiced stop /b/, these tended to lead to ratings towards the ‘not hard’ end of the scale. There were smaller negative effects for the unvoiced stop /p/ and the voiced fricative /v/ in the C1 position, and the unvoiced fricatives in the C2 position. Hardness was represented almost exclusively by consonants since there were no significant effects for any V1 vowel and very small negative effects, for both rounded and unrounded V2 vowels (Figure 2A; Supplementary Table 7).

For the softness dimension, the largest positive effects were for the sonorant /m/ and the unvoiced fricatives /f/ and /s/ in the C2 position (Figure 2A; Supplementary Table 7) as in, for example, /mʊmo/, /sefi/, and /fɪse/ (Table 3). The unvoiced stop /p/ had a small positive effect in the C1 position. The largest negative effects were for the C1 africate /ʧ/ and C2 unvoiced stops, indicating that these favored ratings towards the ‘not soft’ end of the scale. Voiced stops in either consonant position were also linked to ratings as ‘not soft’. As for the hardness dimension, vowels had little influence: the V1 unrounded /i/ was significantly negative in its influence, contributing to ‘not soft’ ratings, while the V2 vowels had non-significant effects (Figure 2A; Supplementary Table 7).

#### Weight

The next best model fits were for the weight domain with R^2^ of .52 for the lightness dimension and .61 for the heaviness dimension (Figure 2B; Supplementary Table 7).

**Figure 2B:**
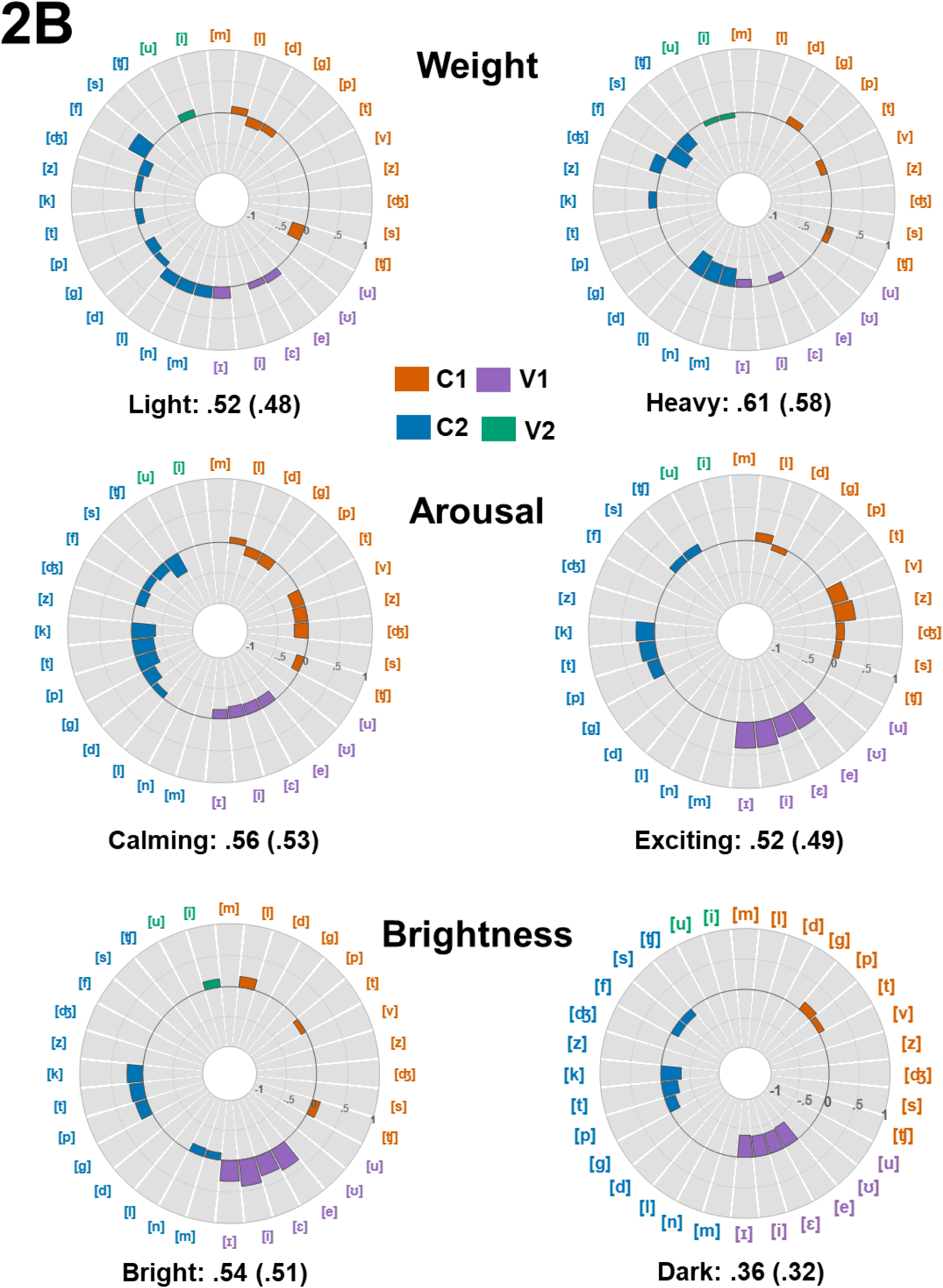
Phonemic regression results for the opposing scales of the weight, arousal, and brightness domains. Interpretation as for Figure 2A. For the complete results, see Supplementary Table 7.

The largest positive driver for lightness ratings was the unvoiced fricative /f/ in the C2 position, as seen in /fεfi/ (Table 3); this increased ratings towards ‘very light’ and produced the largest improvement to model fit in terms of R^2^Δ (Figure 2B; Supplementary Table 7). There were also significant positive effects for the sonorants: /l/ in both C1 and C2, and /m/ and /n/ in C2 only, for example in /lonu/ (Table 3). Lightness ratings were also driven by the front unrounded vowels /e/, /ε/, and /ɪ/ but only in the V1 position. Back rounded vowels were only associated with lightness in the V2 position and only for /u/ (Figure 2B; Supplementary Table 7). There were also significant negative effects of which the largest was for the unvoiced affricate /ʧ/ in the C1 position, rated as ‘not light’ and producing a large change in R^2^. The voiced af/fricatives /z/ and /ʤ/ in the C2 position were also associated with items rated as ‘not light’, as were the unvoiced stop /t/ and, in both C1 and C2 positions, the voiced stops /d/ and /g/ (Figure 2B; Supplementary Table 7).

For the heaviness dimension, the largest positive effect was for the voiced affricate /ʤ/ in the C2 position (Figure 2B; Supplementary Table 7), for example /ʤoʤu/ is among the pseudowords with the highest heaviness ratings (Table 3). The other phonemes in items rated as heavy were the voiced stop /g/ and the unvoiced affricate /ʧ/, both in the C1 position, for example /gebi/ (Table 3), and the unvoiced stop /k/ in the C2 position. The largest negative influence on heaviness ratings were the negative βs for the sonorants in the C2 position, leading to ratings as ‘not heavy’ compared to the referent predictor, the voiced stop /b/, as in /gebi/(Figure 2B; Supplementary Table 7; and see Table 3). The other negative influences were /v/ in the C1 position, /f/ and /s/ in the C2 position. Unlike the lightness scale, none of the vowels produced positive associations to heaviness in either vowel position; instead, these were associated with decrements in heaviness ratings (Figure 2B; Supplementary Table 7).

#### Arousal

The R^2^ for the calming scale was .56 and .52 for the exciting scale (Figure 2B; Supplementary Table 7).

Remarkably, ratings on the calming scale were almost completely defined by negative effects, i.e., phonemes that were rated towards the ‘not calming’ end of the scale (Figure 2B; Supplementary Table 7). The sole positive result was for the sonorant /l/ at C1, which occurred in six of the ten highest-rated items on the calming scale, e.g., /lʊlo/ (Table 3). Otherwise, voiced stops at C1 and C2, unvoiced stops at C2, and af/fricatives generally at both C1 and C2, whether voiced or not, all tended to evoke ‘not calming’ ratings; this only leaves the sonorants, all of which appear in the highest-rated items in Table 3. Compared to the rounded vowel /o/ that was the referent predictor, unrounded vowels at V1 were also significantly associated with ratings that decreased towards the ‘not calming’ end of the scale (Figure 2B; Supplementary Table 7) and, accordingly, rounded vowels appear in 8 of the 10 highest-rated calming pseudowords (Table 3). There were no significant V2 vowel effects.

For the exciting dimension, V1 unrounded vowels were, collectively and individually, the largest influence on ratings (Figure 2B; Supplementary Table 7) and appear in all the ten highest-rated items on this scale, e.g., /vive/ and /ʤɪze/ (Table 3); but there were no significant V2 vowel effects. For the consonants, there were relatively large positive effects for the voiced af/fricatives at C1 but not C2, while there were positive effects for unvoiced stops at C2 but not C1 (Figure 2B; Supplementary Table 7). There were small effects for the sonorant /l/ at C1, the unvoiced fricative /s/ at both C1 and C2, and for the unvoiced affricate /ʧ/ at C2, although only /s/ appears in Table 3: /sife/.

#### Brightness

The R^2^ for the brightness dimension was .54 and .36 for the darkness dimension (Figure 2B; Supplementary Table 7).

Brightness ratings were primarily driven by front unrounded vowels at both V1 and V2, as well as unvoiced stops at C2 (Figure 2B; Supplementary Table 7), both exemplified by /tεti/ (Table 3). There were also small positive effects for the sonorant /l/ and the unvoiced affricate /ʧ/ at C1, although only the latter appeared in the ten highest-rated items, in /ʧifi/ (Table 3). There were also small negative effects for the C1 unvoiced stop /t/ and the C2 sonorants /m/ and /n/.

Complementing the brightness results, darkness ratings were also primarily driven by vowels with large negative effects for V1 unrounded vowels (Figure 2B; Supplementary Table 7), indicating that V1 rounded vowels were important: these occur in seven of the ten pseudowords with the highest ratings on this scale, e.g., /bugu/ and /gobo/ (Table 3). There were no significant V2 vowel effects. There were also positive effects for C1 unvoiced stops, although these did not appear in the ten highest-rated dark pseudowords. There were large negative effects for unvoiced stops and fricatives at C2 (Figure 2B; Supplementary Table 7), indicating that these attracted ratings towards the ‘not dark’ end of the scale.

#### Valence

The valence domain had the second worst model fits with R^2^ of .49 for the goodness dimension and .38 for the badness dimension (Figure 2C; Supplementary Table 7).

**Figure 2C:**
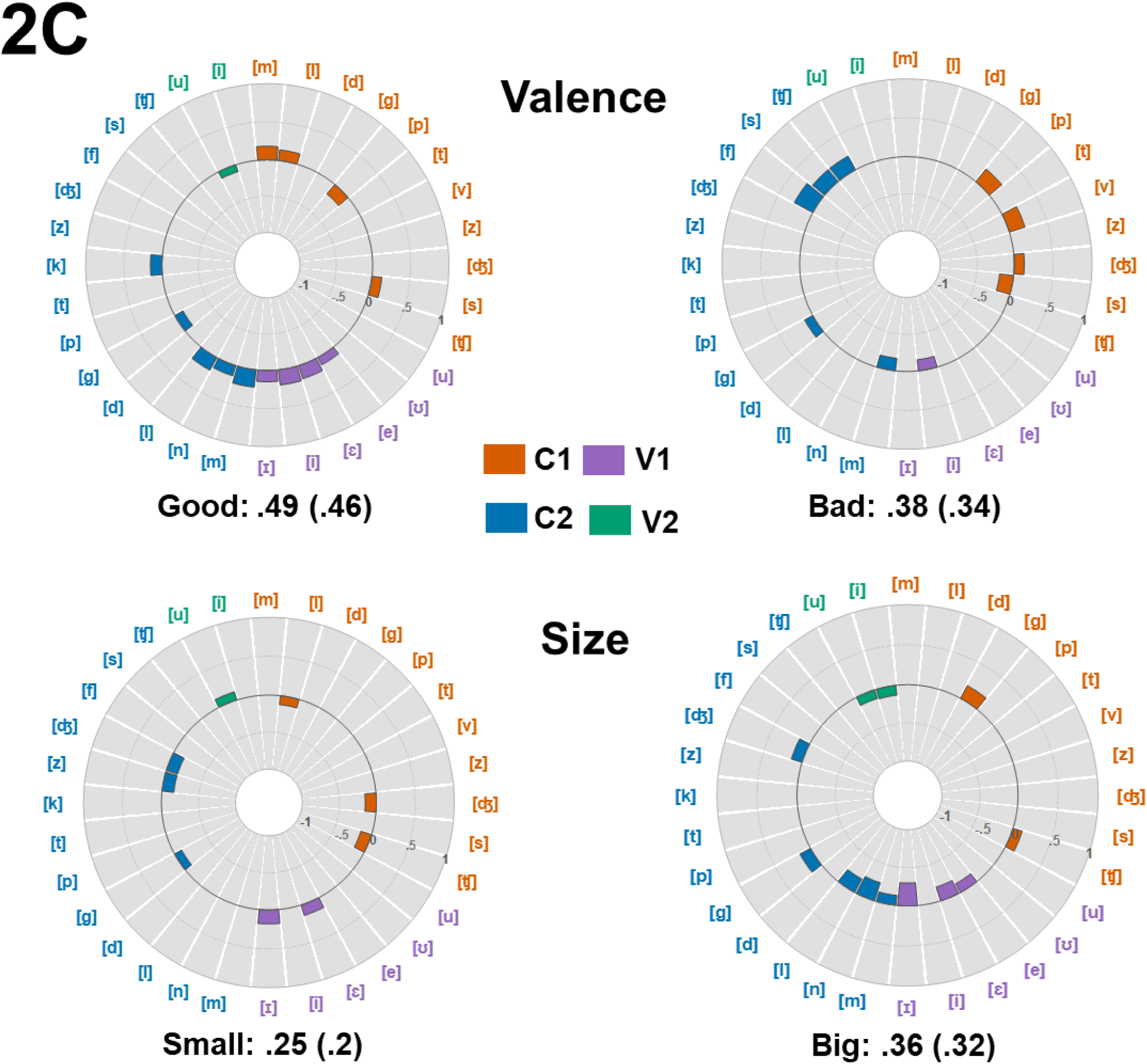
Phonemic regression results for the opposing scales of the valence and size domains. Interpretation as for Figure 2A. For the complete results, see Supplementary Table 7.

**Figure 3:**
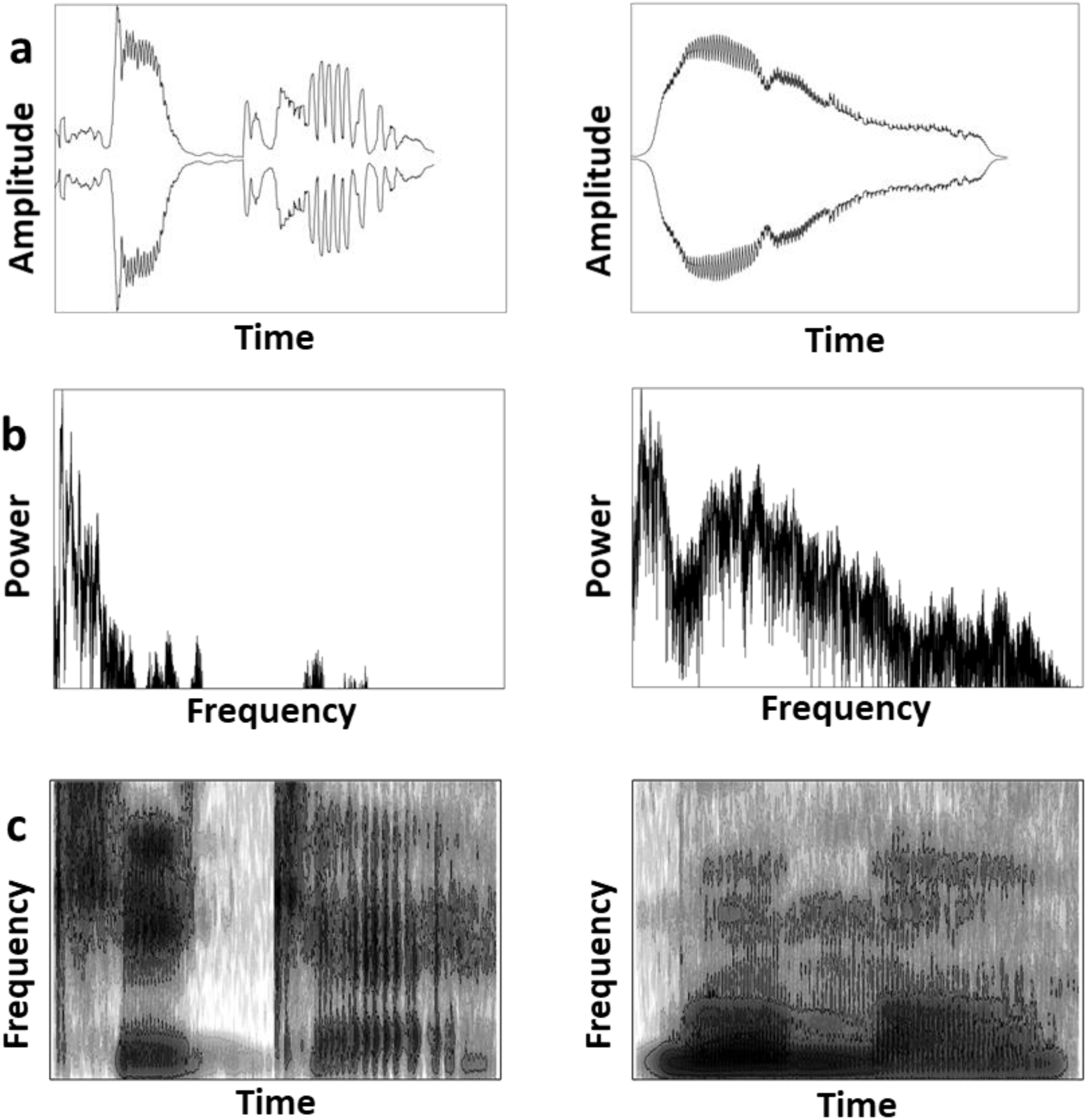
Visual depictions of two pseudowords characterized by stops and high frequency vowels (/kike/, left column) and sonorants and low frequency vowels (/mumo/, right column) as reflected in (a) the speech envelope, (b) spectral tilt: frequency is low to high, left to right, and (c) spectrogram: frequency is low to high, bottom to top; shading = power, low power is lighter and high power is darker.

Goodness ratings were primarily driven by sonorants at both C1 and C2 and front unrounded vowels at V1 (Figure 2C; Supplementary Table 7), for example, /lεme/ and /bɪbi// (Table 3); collectively, these produced the largest improvement to R^2^. There were also small effects for the unvoiced fricative /s/ at C1 and the unvoiced stop /k/ at C2, although neither of these appeared in the highest-rated good pseudowords (Table 3). There were negative effects for the unvoiced stop /p/ at C1, the voiced stop /g/ at C2, and the rounded vowel /u/ at V2, indicating that these were rated as ‘not good’ (Figure 2C; Supplementary Table 7).

For the badness dimension, the results were more complicated: the largest positive effects were for the unvoiced stop /p/ and the voiced fricative /v/ at C1 and, at C2, the unvoiced fricatives /f/ and /s/ and affricate /ʧ/ (Figure 2C; Supplementary Table 7), examples being /pupo/ and /foʧu/ (Table 3). There were smaller positive effects for the voiced affricate /ʤ/ at C1 and the voiced stop /g/ at C2. The only negative effects for consonants were for C1 /s/ and C2 /m/, indicating that these were linked to ratings as ‘not bad’ (Figure 2C; Supplementary Table 7). The only result for vowels on the badness scale was that the front unrounded vowel /i/ at V1 tended to be rated as ‘not bad’; this is consistent with the presence of unrounded vowels in eight of the highest-rated good pseudowords, and rounded vowels in nine of the highest-rated bad pseudowords (Table 3).

#### Size

The size domain had the worst model fit between phonemes and ratings with R^2^ of .25 for the smallness dimension and .36 for the bigness dimension (Figure 2C; Supplementary Table 7).

For the smallness dimension, the only positive results were for vowels: the front unrounded vowels /ε/ and /ɪ/ at V1 and the back rounded vowel /u/ at V2 (Figure 2C; Supplementary Table 7), the former appearing in seven, and the latter in two, of the ten highest-rated small pseudowords (Table 3), e.g., /pɪte/ and /putu/, respectively. There were negative effects for C1 /l/ and /ʧ/, C2 /g/ and /z/, and /ʤ/ at both C1 and C2 (Figure 2C; Supplementary Table 7), indicating that these tended to feature in items rated towards the ‘not small’ end of the scale (and, in fact, with the exception of /l/, all of these appear in the highest-rated big pseudowords, see Table 3).

For the bigness dimension, the only positive effects were for the voiced stop /g/ in both C1 and C2, and for the unvoiced and voiced affricates /ʧ/ and /ʤ/ at C1 and C2, respectively (Figure 2C; Supplementary Table 7), as exemplified by /gʊdo/ and /zɪʤi/, for example (Table 3). The largest improvements to R^2^ were brought about by negative effects for V1 front unrounded vowels, indicating that these were associated with ratings as ‘not big’ and that, in comparison, rounded vowels were associated with ratings as ‘very big’ (Figure 2C; Supplementary Table 7). But, as can be seen from Table 3, both vowel types were equally represented in the highest-rated big pseudowords. There were also negative effects for all the sonorants at C2 and for /u/ and /i/ at V2, indicating that these contributed to ratings as ‘not big’.

### Discussion

In this analysis we attempted to answer two questions: (1) Having identified a phonetic category, e.g., sonorants, as important to a particular iconic mapping, is it the case that any sonorant can contribute to that mapping or do some carry more weight than others? (2) Does any phoneme’s contribution depend on where it occurs in the utterance? As discussed below, the answers are (1) that some iconic mappings involved all members of a phonetic category (e.g. roughness ratings involved all the obstruents at C2 and goodness ratings involved all the unrounded vowels at V1: (Figure 2A; Supplementary Table 7), while other mappings depended on specific phonemes (e.g. at C1, hardness ratings depended only on the unvoiced affricate /ʧ/, and at V1 smallness ratings depended only on /ε/ and /ɪ/ out of the four unrounded vowels: (Figures 2A and 2C, respectively; Supplementary Table 7). As these examples suggest, the answer to (2) is that phoneme positions were also important: for some mappings both first and second positions were important (e.g. pointedness ratings depended on almost all the obstruents at C1 and C2 alike: (Figure 2A; Supplementary Table 7), while for others there were specific positional effects (e.g. obstruents had a greater impact on hardness ratings at C2 than C1: (Figure 2A; Supplementary Table 7).

Consistent with the phonetic category analysis, the answers to these questions tend to be domain-specific. For the shape domain, for example, the category analysis suggests that roundedness is driven almost entirely by vowels (Figure 1A: Supplementary Table 4); the phoneme analysis bears this out but also shows that the vowel effect is almost exclusively at V1, not V2 (Figure 2A; Supplementary Table 7). Pointedness, by contrast, was driven almost entirely by consonants, especially unvoiced obstruents (Figure 1A; Supplementary Table 4); but the phoneme analysis shows that, at C1, this is really only the unvoiced /t/ and /ʧ/, occurring in 5 out of 10 items in Table 3. The main effect is at C2, where it is the unvoiced obstruents (except for /f/) that produce the largest improvements in model fit (Figure 2A; Supplementary Table 7) and these occur at C2 in all 10 items in Table 3.

A different pattern occurred for the roughness aspect of texture. The phonetic category analysis suggested that sonorants were important for the smoothness dimension (Fig 1A; Supplementary Table 4), but the phoneme analysis showed that only /l/ was significant at C1, occurring in 6 of the 10 items in Table 3, and that the main effect was at C2 when all the sonorants contributed, occurring in all 10 items in Table 3 and producing the largest changes to R^2^ only at that position (Figure 2A; Supplementary Table 7). By contrast, vowels were not positively associated with smoothness at either position. The phonetic category analysis suggested that roughness ratings were driven by obstruents, perhaps especially if unvoiced (Fig 1A; Supplementary Table 4); by contrast, the phoneme analysis shows that at C1 only the unvoiced affricate /ʧ/ contributes significantly, whereas all the voiced obstruents contribute (Figure 2A; Supplementary Table 7). At C2, however, all the unvoiced obstruents contributed and produced the largest change to R^2^ with smaller contributions from the voiced obstruents (Figure 2A; Supplementary Table 7).

For the hardness aspect of texture, hardness itself was associated with obstruents, especially if unvoiced (Fig 1A; Supplementary Table 4), whereas the phoneme analysis showed that at C1 this was restricted to /ʧ/ with smaller contributions from voiced stops (Figure 2A; Supplementary Table 7). At C2, all the stops, voiced and unvoiced, together with voiced af/fricatives were positively associated with hardness – although note that Table 3 shows that all 10 of the highest-rated items had unvoiced stops at both C1 and C2. This is consistent with Monaghan & Fletcher (2019) who found that the initial consonants /g/, /k/, and /z/ predicted hardness ratings. For the softness dimension, the phonetic category analysis suggested that sonorants were the main driver but in the phoneme analysis these were not significant at C1, despite appearing in 5 out of the 10 items in Table 3. Consistent with Monaghan & Fletcher (2019), the unvoiced fricatives /f/ and /s/ occurred at C1 in 3 of the 10 highest-rated items (Table 3). At C2, only the sonorant /m/ was positively associated with softness, along with the unvoiced fricatives /f/ and /s/. The latter did not appear in the phonetic category analysis, indicative perhaps of problems inherent in collapsing phonemes across categories and position, although we should note that the phonetic analysis did suggest that bilabial/labiodental consonants were important to softness ratings at both C1 and C2, occurring in 7 items at C1 or C2 (Table 3).

In the weight domain, the phonetic category analysis suggested that lightness ratings were driven by sonorants (Fig 1B; Supplementary Table 4) but the phoneme analysis showed that this effect was concentrated at C2 where all the sonorants were significant, occurring in 5 out of the 10 highest-rated items (Table 3), and less so at C1 where only /l/ was significant (Fig 2B; Supplementary Table 7). Similarly, the smaller influence of unvoiced consonants (Fig 1B; Supplementary Table 4) was only positively associated with lightness ratings at C2 and only for the fricative /f/, accounting for a further 3 of the highest-rated items (Table 3). As for shape, the influence of front unrounded vowels predicted by the phonetic category analysis (Fig 1B; Supplementary Table 4) was concentrated at V1, although this likely also reflects that half of them did not occur at V2 anyway (Table 1) because they do not appear in word-final positions in English. For the heaviness dimension, the phonetic category analysis indicated a strong positive association with stops, together with the af/fricatives and voiced consonants (Fig 1B; Supplementary Table 4). However, in the phoneme analysis, positive associations for stops were limited to the voiced /g/ at C1 (3 instances in Table 3) and the unvoiced /k/ at C2, and for af/fricatives to /ʧ/ at C1 and the voiced /ʤ/ at C2 (Figure 2B; Supplementary Table 7). While this might seem to be a further illustration of the problems involved in collapsing individual phonemes into broad categories reflecting the *manner* of articulation, we should note that the phonetic category analysis also indicated positive associations with heaviness ratings for the post-alveolar/velar consonants at both C1 and C2 – i.e., *place* of articulation – and that these are exactly the four consonants mentioned above (see Table 3 and Fig 1B; Supplementary Table 4).

Turning to the arousal domain, we would expect sonorants and back rounded vowels to have been associated with ratings for the calming dimension from the phonetic category analysis (Fig 1B; Supplementary Table 4) but only /l/ at C1 was significantly positively weighted in the phoneme analysis and there were no significant effects for any sonorant at C2 (Figure 2B; Supplementary Table 7), despite appearing in both positions in all 10 of the highest-rated items (Table 3). Significant negative weights for all unrounded vowels at V1 suggest that the referent rounded vowel /o/ was associated with calming ratings, but it occurred in only one item at this position in the ten highest-rated items (Table 3). However, rounded vowels occurred at both V1 and V2 in 8 of these items, so this is an instance where the phonetic category analysis was likely better than the phoneme analysis. For the exciting dimension of arousal, the phonetic category analysis would suggest obstruents, perhaps especially unvoiced, and front unrounded vowels (Fig 1B; Supplementary Table 4). For the obstruents, the phoneme analysis clarifies that the effect of voicing is largely split between C1 and C2, with the voiced /v/, /z/, and /ʤ/ at C1 together with the unvoiced /d/, while at C2 we find all the unvoiced obstruents except /f/ (Figure 2B; Supplementary Table 7). The unrounded vowels are indeed all positively weighted at V1, but there are no significant vowels at V2 (Figure 2B; Supplementary Table 7). Thus, the phonetic category and phoneme analyses were in broad agreement, with the additional clarification of a positional effect for voicing.

For the brightness domain, we would expect unvoiced consonants to be associated with brightness ratings (Figure 1B; Supplementary Table 4). This was true for 9 of the 10 highest-rated items (Table 3), but the phoneme analysis showed that the main impact of these was at C2: only /ʧ/ was significantly positively weighted at C1 while all three unvoiced stops were significant at C2 (Figure 2B; Supplementary Table 7). The phonetic category analysis also suggested that front unrounded vowels connoted brightness (Figure 1B; Supplementary Table 4), and this was confirmed in the phoneme analysis at both V1 and V2 (see Figure 2B; Supplementary Table 7; and Table 3 for examples). By contrast, the phonetic category analysis suggested that darkness was associated with voiced obstruents and back rounded vowels (Figure 1B; Supplementary Table 4). However, for consonants, the phoneme analysis showed that the only significant results at V1 were positive weights for the unvoiced stops /p/ and /t/ at V1 while, at C2, there were significant negative weights for all unvoiced stops and fricatives, indicating that voiced consonants were more important at this position (Figure 2B; Supplementary Table 7). Thus, the phoneme analysis again suggested positional effects that were not apparent from the phonetic category analysis. Both phonetic category and phoneme analyses converged on the association of back rounded vowels with darkness ratings even if, in the phoneme analysis, this was due to negative weights for the unrounded vowels rather than positive weights for the rounded vowels and only at V1 (Figures 1B and 2B; Supplementary Tables 4 and 7).

The phonetic category analysis suggested that, in the valence domain, goodness was represented by sonorants and front unrounded vowels (Figure 1C; Supplementary Table 4). This was confirmed in the phoneme analysis which showed significant effects for all the sonorants at both C1 and C2, and for all unrounded vowels at V1 (Figure 2C; Supplementary Table 7). Consistent with this, 7 of the 10 highest-rated items have sonorants at both C1 and C2, and also unrounded vowels at V1 (Table 3). Badness was represented by obstruents and back rounded vowels in the phonetic category analysis (Figure 1C; Supplementary Table 4), but the phoneme analysis showed a more nuanced picture (Figure 2C; Supplementary Table 7): at C1 there were significant positive weights for only the unvoiced /p/ and the voiced /v/ and /ʤ/, occurring in 6 of the 10 highest-rated items (Table 3), whilst at C2 there were positive weights for the voiced stop /g/ and the unvoiced af/fricatives /f/, /s/, and /ʧ/, although these only accounted for 3 of the C2 obstruents in Table 3. Moreover, there were no significant positively weighted vowels for badness, although the rounded vowels suggested by the phonetic category analysis accounted for 9 of 10 vowels at V1 and V2 in Table 3.

Finally, in the size domain, the phonetic category analysis suggested that smallness was associated with unvoiced stops (and no vowel effects) while bigness was associated with obstruents in general, especially if voiced, and back rounded vowels (Figure 1C; Supplementary Table 4). The phoneme analysis was quite different, however. For smallness, the unvoiced stops were non-significant at both C1 and C2, and there were no significant positive weights for any other consonant (Figure 2C; Supplementary Table 7). This contrasted with Monaghan & Fletcher (2019) who found that the initial unvoiced consonants /p/, /t/, and /s/ predicted smallness. Another departure from the phonetic category analysis, which showed no vowel effects, was that the front unrounded vowels /ε/ and /ɪ/ were positively weighted at V1 and the rounded vowel /u/ was positively weighted at V2 (Figure 2C; Supplementary Table 7). For bigness ratings, the only positively weighted voiced obstruents were /g/ at both C1 and C2, and /ʤ/ at C2 (Figure 2C; Supplementary Table 7). This was somewhat consistent with Monaghan & Fletcher (2019) who found that an initial /g/, but also initial /b/ and /z/, predicted bigness. Thus, the phonetic category and phoneme analyses diverged substantially for both dimensions of size, but we should recall that size had the worst model fit in both analyses.

## WHOLE-ITEM ACOUSTIC ANALYSIS

### Introduction

Our predictions for the acoustic characteristics associated with each domain flow from the preceding phonetic analysis (see above) of the current pseudoword set. Since the spectro-temporal parameters take multiple samples across the waveform of each pseudoword, these are the most easily connected to the phonetic components of the pseudowords. This is less easily done for the voice quality parameters, for which we derived only one value per pseudoword: while the FUF, mean HNR, MAC, and pulse number all reflect the relative voicing/periodicity of each item – and therefore connect to phonetic features in a broad sense – it is less easy to pinpoint the phonetic features contributing to vocal variability.

Consonants characterized as stops, af/fricatives, and sonorants for example, each have a different manner of articulation and thus different acoustic characteristics. Stops involve a constriction in the vocal tract that temporarily blocks the flow of air, followed by a release as airflow resumes (Ladefoged & Johnson, 2011). The blockage results in a short period of low energy as the constriction is formed, followed by a ‘burst’ of energy (see Chodroff & Wilson, 2014) as the blockage is released; these can be seen in the spectrogram and speech envelope as abrupt variations in power and amplitude, respectively (see Supplementary Figures 1 and 2 for unvoiced and voiced stops in /tike/ and /gobo/, respectively). Fricatives (/f/, /v/, /s/, and /z/ in Table 1) involve a partial obstruction of the vocal tract that results in a turbulent airflow and also involve higher frequencies than almost any other phoneme (Ladefoged & Johnson, 2011; see also Jongman et al., 2000). Pseudowords including fricatives should therefore have a relatively flatter spectral tilt since there will be more power at the higher frequencies (see Supplementary Figure 7 for /vuʤo/). Affricative consonants (/ʤ/ and /ʧ/ in Table 1) combine a stop and a fricative (Ladefoged & Johnson, 2011) and their acoustic consequences therefore manifest as a combination of abrupt variations in power and amplitude, as demonstrated in the spectrogram and speech envelope, respectively, together with concentrations of power at the high frequencies, reflected in both the spectrogram and spectral tilt (see Supplementary Figures 4 and 7 for /ʧiʧe/ and /vuʤo/ respectively). Sonorants do not involve an obstruction of the airflow: although there is a constriction in the vocal tract, the airflow continues through the mouth for /l/, and through the nose for /m/ and /n/ (Ladefoged & Johnson, 2011). Since the airflow is also unobstructed for vowels, the transitions between sonorants and vowels in the CVCV pseudowords should result in more gradual variations in power in the spectrogram and a smoother speech envelope (compare, for instance, /mʊmo/ to /tike/ in Supplementary Figure 1).

Different places of articulation also result in varying acoustic consequences because they alter the length of the vocal tract depending on where the constriction that produces the sound is located (Table 1). Bilabial consonants like /b/, for example, involve resonances of the whole vocal tract because the constriction is at the lips, and therefore are associated with high energy at the lower frequencies (Reetz & Jongman, 2020). By contrast, alveolar consonants like /d/ produce most energy at high frequencies because the constriction is produced by the tongue against the alveolar ridge, just behind the teeth, and therefore the vocal tract in front of the constriction is short (Reetz & Jongman, 2020). Velar consonants like /g/ produce energy in the middle range of frequencies, the constriction being produced by the tongue against the velum, or soft palate, at the back of the mouth, and an intermediate anterior vocal tract length (Reetz & Jongman, 2020). These considerations enable us to make predictions about spectral tilt and the FFT in particular. Pseudowords consisting of bilabial consonants will tend to show a steeper spectral tilt than those consisting of alveolar consonants because of the concentration of energy at low and high frequencies respectively. For the FFT, there will be differences in the distribution of energy at each frequency band over the duration of the pseudoword; these will be apparent from the spectrogram.

#### Shape

In the shape domain, we expected to replicate, in an independent sample of participants, our previous findings for spectro-temporal and voice quality parameters of the auditory speech signal (Lacey et al., 2020), as summarized here. In this earlier study, we found spectral tilt to be steeper for rounded pseudowords where power is concentrated in the low-frequency bands (Lacey et al., 2020), reflecting the sonorants and back rounded vowels that are associated with roundedness (McCormick et al., 2015; see also Table 4). However, spectral tilt flattened out for pointed pseudowords as power migrated to the higher frequencies associated with the obstruents and/or front unrounded vowels that these pseudowords contain (Lacey et al., 2020). This also reflects differences in the place of articulation with power being concentrated at lower frequencies for the bilabial/labiodental consonants associated with roundedness, but dispersing to the higher frequencies for the alveolar and post-alveolar/velar consonants associated with pointedness (see Table 4). We also found that the FFT reflected power variations that were gradual for rounded pseudowords and abrupt for pointed pseudowords (Lacey et al., 2020), again reflecting the presence of sonorants and obstruents, and rounded/unrounded vowels respectively (McCormick et al., 2015; see Table 4). The speech envelope was smoother and more continuous for rounded, compared to pointed, pseudowords (Lacey et al., 2020).

**Table 4:** Summary of the phonetic category, phoneme, and acoustic – whole item and segmental – regression analyses. R^2^ and adjusted R^2^ (Adj R^2^).

|  |  | Phonetic category |  | Phonemes |  | Acoustic: whole item |  | Acoustic: segmental |  |
| --- | --- | --- | --- | --- | --- | --- | --- | --- | --- |
| | | $R^2$ | Adj $R^2$ | $R^2$ | Adj $R^2$ | $R^2$ | Adj $R^2$ | $R^2$ | Adj $R^2$ |
| **Shape** | Rounded | 0.88 | 0.87 | 0.83 | 0.82 | 0.30 | 0.28 | 0.77 | 0.76 |
|  | Pointed | 0.76 | 0.75 | 0.90 | 0.90 | 0.56 | 0.55 | 0.66 | 0.65 |
| **Roughness** | Smooth | 0.57 | 0.56 | 0.66 | 0.63 | 0.35 | 0.34 | 0.43 | 0.41 |
|  | Rough | 0.77 | 0.77 | 0.86 | 0.85 | 0.42 | 0.41 | 0.52 | 0.51 |
| **Hardness** | Hard | 0.68 | 0.67 | 0.80 | 0.79 | 0.45 | 0.43 | 0.54 | 0.52 |
|  | Soft | 0.33 | 0.31 | 0.45 | 0.41 | 0.18 | 0.16 | 0.29 | 0.26 |
| **Weight** | Light | 0.43 | 0.42 | 0.52 | 0.48 | 0.24 | 0.23 | 0.31 | 0.29 |
|  | Heavy | 0.53 | 0.52 | 0.61 | 0.58 | 0.26 | 0.25 | 0.40 | 0.38 |
| **Arousal** | Calming | 0.48 | 0.47 | 0.56 | 0.53 | 0.24 | 0.23 | 0.40 | 0.38 |
|  | Exciting | 0.36 | 0.35 | 0.52 | 0.49 | 0.17 | 0.15 | 0.31 | 0.29 |
| **Brightness** | Bright | 0.53 | 0.52 | 0.54 | 0.51 | 0.23 | 0.21 | 0.48 | 0.47 |
|  | Dark | 0.35 | 0.34 | 0.36 | 0.32 | 0.13 | 0.11 | 0.30 | 0.28 |
| **Valence** | Good | 0.40 | 0.39 | 0.49 | 0.46 | 0.18 | 0.16 | 0.33 | 0.31 |
|  | Bad | 0.28 | 0.26 | 0.38 | 0.34 | 0.16 | 0.14 | 0.24 | 0.22 |
| **Size** | Small | 0.12 | 0.11 | 0.25 | 0.20 | 0.19 | 0.17 | 0.21 | 0.19 |
|  | Big | 0.29 | 0.27 | 0.36 | 0.32 | 0.12 | 0.10 | 0.28 | 0.26 |

For the previously studied voice quality parameters, mean HNR decreased as the speech pattern became progressively less smooth and more uneven, reflecting the change from rounded to pointed (Lacey et al., 2020). The MAC decreased in the same way: higher autocorrelation values indicate a smoother voice pattern and/or more voiced segments, associated with roundedness ratings, while lower values indicate an uneven pattern and/or fewer voiced or periodic segments, associated with ratings of pointedness (Lacey et al., 2020). Similarly, higher pulse numbers indicate a smoother voice pattern, and/or more voiced segments, associated with rounded pseudowords, while lower pulse numbers indicate a more uneven voice pattern, and/or fewer voiced segments, associated with pointed pseudowords (Lacey et al., 2020). In keeping with this, the pulse number decreased from the rounded to the pointed pseudowords (Lacey et al., 2020); note that this effect has been independently replicated (Akita, 2021). The FUF increased as ratings of pseudowords transitioned from rounded to pointed (Lacey et al., 2020), because auditory roundedness and pointedness are more associated with voiced and unvoiced elements, respectively (McCormick et al., 2015; Lacey et al., 2024).

Variation in amplitude and frequency as measured by shimmer and jitter, respectively, also increased, reflecting increasing vocal variability and unevenness in the speech pattern, as pseudoword ratings progressed from rounded to pointed (Lacey et al., 2020). Similarly, lesser and greater F0 variability, as measured by F0_SD_, indicated roundedness and pointedness, respectively (Lacey et al., 2020).

The present study also introduced two parameters that were not included in the prior study of Lacey et al. (2020): mean F0 and pseudoword duration. We expect that mean F0 will increase as ratings move from rounded to pointed (Parise & Pavani, 2011; Knoeferle et al., 2017). Finally, an intuitive prediction is that roundedness and pointedness should be associated with longer and shorter pseudoword duration, respectively.

#### Hardness

Iconic hardness ratings were associated with obstruents at the alveolar and post-alveolar/velar places of articulation, with little influence of vowels, while softness was associated with sonorants and the bilabial/labiodental place of articulation, and with back rounded vowels. We therefore expected that there would be more energy at higher frequencies for hard, compared to soft, pseudowords, which should be reflected in a flatter spectral tilt for the former. The spectrograms should similarly show that hard pseudowords have more energy at the higher frequencies and more abrupt variations in energy, related to the presence of obstruents, compared to the more gradual variations involved in the sonorants that are associated with softness. These differences should also be reflected in the speech envelope, with clear discontinuities for the hard pseudowords but a smoother, more continuous envelope for the soft pseudowords.

For the voice quality parameters, we would expect that softness would be associated with pseudowords that sound smoother, i.e. that have greater periodicity and less variability, while those that indicate hardness would involve less periodicity and more variability, i.e. relatively greater vocal roughness. While we are cognizant of describing one aspect of texture in terms of another (hard/soft and rough/smooth are independent dimensions of tactile texture [Hollins et al., 2000]), this is supported by the fact that we found that hardness was associated with af/fricatives (which are inherently noisy and aperiodic [Reetz & Jongman, 2020]), while softness was associated with sonorants (Table 4). Thus, we would expect that the mean HNR, MAC, and pulse number would all increase, and that FUF would decrease, as increasing periodicity accompanies the transition of ratings from hard to soft. This transition would also be marked by a reduction in vocal variability and so jitter, shimmer, and F0_SD_ would reduce from hard to soft pseudowords. We did not make specific predictions for mean F0 or duration.

#### Weight

Phonetic analysis of the weight domain showed that lightness ratings were associated with sonorants and the bilabial/labiodental place of articulation, together with unrounded vowels; heaviness ratings were associated with voiced obstruents and the post-alveolar/velar place of articulation, together with rounded vowels (Table 4). We do not expect a major association of ratings with spectral tilt because both light and heavy pseudowords involve energy across the frequency range: sonorants and bilabial/labiodental articulation involve low frequencies while unrounded vowels involve high frequencies; by contrast, obstruents with post-alveolar/velar articulation involve somewhat higher frequencies while rounded vowels are generally of lower frequency. Pseudowords rated as light or heavy should, however, be distinguishable by their spectrograms which should show more gradual variations in energy for light pseudowords and their associated sonorants, compared to more abrupt variations for heavy pseudowords and their associated obstruents. The difference between sonorants and obstruents for light and heavy pseudowords, respectively, should also be apparent in more discontinuous speech envelopes for the latter compared to the former.

Since there is a real-world relationship in which size and weight are generally positively correlated, some predictions about the voice quality parameters can be derived by comparison to the size domain. High and low F0 are associated with small and big size, respectively (reviewed by Spence, 2011) and so pseudowords rated as light/heavy should also exhibit high/low mean F0 respectively. We expect that measures of periodicity – pulse number, mean HNR, MAC, and FUF – will show that periodicity decreases as ratings change from light to heavy, tracking the change from sonorants and other voiced consonants, in pseudowords rated as light, to unvoiced consonants in those rated as heavy (Table 4). But it is an open question whether vocal variability – as measured by jitter, shimmer, and F0_SD_ – is related to weight. By analogy to size, however, we would predict that shorter/longer duration would reflect light/heavy pseudowords, respectively.

#### Arousal

Sonorants and back rounded vowels are associated with pseudowords rated as calming (Sidhu et al., 2022; see also Table 4) while unvoiced obstruents and front unrounded vowels are associated with pseudowords rated as exciting (Sidhu et al., 2022; see Table 4). The relationship between arousal and place of articulation is asymmetric: while calming ratings are strongly associated with bilabial/labiodental articulation, exciting ratings are not associated with a specific place of articulation (Table 4). Accordingly, we expect a relatively steep spectral tilt for calming pseudowords with energy concentrated at the lower frequencies, reflecting sonorants and rounded vowels, but a flatter spectral tilt for exciting pseudowords reflecting the higher-frequency energy associated with unvoiced obstruents. In addition, calming/exciting pseudowords should show smoother/more abrupt changes in energy in the spectrogram, together with continuous/discontinuous speech envelopes, respectively.

For the voice quality parameters, since calming/exciting were associated with voiced and unvoiced consonants respectively, the FUF should increase as ratings change from calming to exciting. We would expect that pulse number, MAC, and mean HNR should decrease as ratings transition from calming to exciting, reflecting decreasing periodicity. Although voice stress analysis suggests that jitter and shimmer decrease with arousal (reviewed by Van Puyvelde et al., 2018), this was observed largely in relation to real-life emergency communications, a very different context to the present study. However, this review also notes that general, i.e., non-emergency, arousal produces increases in F0 range, i.e., variability, so we might expect jitter and F0_SD_ to increase with arousal. If increasing arousal produced a general increase in vocal variability, then we would also expect shimmer to increase. Intuitively, mean F0 should increase as ratings progress from calming to exciting, and shorter pseudowords should be rated as more exciting than longer pseudowords.

#### Brightness

Our phonetic analysis of this domain showed that consonant voicing and vowel rounding were the main predictors of sound-symbolic brightness ratings (Table 4). Thus, we expected that pseudowords rated as dark would have energy concentrated in the lower-frequency bands, given their association with voiced consonants and back rounded vowels (Table 4) and thus would show a steeper spectral tilt than those rated as bright, which are associated with unvoiced consonants and front unrounded vowels (Table 4) involving energy at higher frequencies. The spectrogram should therefore show more power at high frequencies for the bright, compared to the dark, pseudowords. The crossmodal correspondence in which louder/quieter sounds are associated with bright/dark stimuli, respectively (reviewed in Spence, 2011; see also Tzeng et al., 2018), suggests that the speech envelope would reveal greater amplitude for bright, compared to dark, pseudowords. Note that place of articulation was not a particular predictor of bright/dark ratings and thus we offer no related acoustic hypotheses.

For the voice quality parameters, there is also a crossmodal correspondence between high/low F0 and bright/dark stimuli, respectively (reviewed in Spence, 2011; see also Tzeng et al., 2018). We could therefore expect pseudowords reflecting the brightness/darkness dimension to have high/low mean F0, following Tzeng et al. (2018) who found that, on average, participants produced pseudowords with higher/lower pitch in response to brighter/darker colors, respectively. Other predictions follow from the association between brightness/darkness and unvoiced/voiced consonants respectively (Newman, 1933; Hirata et al., 2011; Lacey et al., 2024): the FUF should decrease from bright to dark whereas the pulse number, MAC, and mean HNR should increase (see Lacey et al., 2020, for an example in the shape domain of how these parameters co-vary). Finally, brightness/darkness should correspond to shorter/longer duration (Tzeng et al., 2018). Note that the pitch and amplitude crossmodal associations with brightness were initially established with auditory stimuli that, unlike speech utterances, did not intrinsically vary in frequency or intensity (i.e., steady state pure tones, see Wicker, 1968). Thus, we have no specific prediction as to whether parameters that capture variability in pitch (jitter and F0_SD_) and amplitude (shimmer) will be important.

#### Valence

Our phonetic analysis of the valence domain indicated that the main predictors of pseudowords rated as good were sonorants and front unrounded vowels, while pseudowords rated as bad were associated with obstruents and back rounded vowels (Table 4). Place of articulation was not a strong predictor of good/bad ratings (Table 4). Pseudowords rated as sounding good or bad both involve energy at high and low frequencies: unrounded vowels and sonorants, respectively, for good pseudowords, and obstruents and rounded vowels, respectively, for bad pseudowords. Therefore, as for the weight domain, we did not expect major differences in spectral tilt. Good and bad pseudowords should, however, be distinguishable by their spectrograms which should show more gradual and diffuse changes in energy for the good pseudowords and their constituent sonorants, compared to more abrupt changes for the bad pseudowords and their constituent obstruents. The speech envelope should also distinguish between good and bad pseudowords in being smoother and more continuous for the former and their associated sonorants, but more discontinuous for the latter and their associated obstruents.

For the voice quality parameters, it seems reasonable to assume increasing vocal variability as pseudowords change from good ratings and sonorants to bad ratings and obstruents. We therefore expected that jitter, shimmer, and F0_SD_, would increase as ratings transitioned from good to bad and, accordingly, that HNR, MAC, and pulse number would decrease, and FUF would increase, in tandem with this change (since decreasing periodicity implies a noisier signal). However, the effect of voicing was small (Table 4) in general, so we might expect measures of periodicity to be correspondingly weakly associated with valence ratings. Intuitively, badness would be associated with low, rather than high, F0 (Belyk & Brown, 2014), but there is no obvious prediction for pseudoword duration in this domain.

#### Size

For the size domain, iconic smallness ratings were associated with unvoiced, bilabial/labiodental stops but not af/fricatives or vowels, while large size was associated with voiced, post-alveolar/velar stops and af/fricatives, and back rounded vowels (Table 4). We expected that spectral tilt and the spectrogram would be broadly similar for small and big pseudowords since both might involve low and high frequencies. For pseudowords judged as small, although unvoiced consonants are generally produced with higher frequency than voiced consonants, the bilabial/labiodental place of articulation results in lower frequencies. For pseudowords judged as big, fricatives exhibit high frequencies, whereas the post-alveolar/velar place of articulation and back rounded vowels should involve lower frequencies. These concentrations of energy at both low and high frequencies should be apparent from the frequency distributions shown in the spectral tilt and the spectrogram for both small and big pseudowords. Since small and big ratings were both associated with stops, we would expect the speech envelope for both to show a discontinuous profile but, given the difference in voicing, the envelope for voiced, big pseudowords might exhibit greater amplitude compared to the unvoiced, small pseudowords.

For the voice quality parameters, we can make predictions based on the well-established crossmodal correspondence in which high and low pitch are associated with small and big size, respectively (reviewed by Spence, 2011). Pseudowords with high/low mean F0 should therefore be rated as small/big respectively. Although pulse phonation, or ‘creaky voice’ has recently been associated with large size (Akita, 2021), the speaker in that study deliberately employed pulse phonation whereas ours spoke in their typical voice. Nonetheless, it is likely that measures of periodicity will increase (HNR, MAC, pulse number) or decrease (FUF) following the change from unvoiced to voiced consonants for pseudowords judged as small or big, respectively (Table 4). However, it is unclear how measures of vocal variability (jitter, shimmer, F0_SD_) relate to size iconicity. Pseudowords with shorter/longer durations should be rated as smaller/bigger respectively (Knoeferle et al., 2017).

#### Summary of domain-specific predictions

We can summarize the predictions as follows. For the spectro-temporal parameters, we expected that pseudowords rated as rounded, soft, dark, and calming would be associated with a steeper spectral tilt than those rated as pointed, hard, bright, and exciting. Pseudowords rated as rounded, light, soft, calming, and good should exhibit a more continuous speech envelope and more gradual variations in power across frequency in the FFT, whereas those rated as pointed, heavy, hard, exciting, and bad should result in a discontinuous speech envelope and more abrupt variations in power. For the voice quality parameters, we expected that increased periodicity (as measured by the HNR, MAC, pulse number, and FUF) would generally reflect ratings of pseudowords as rounded, small, soft, light, bright, soft, calming, and good. Conversely, decreased periodicity – i.e., more noise in the signal – would reflect pseudowords rated as pointed, big, hard, heavy, dark, exciting, and bad. We expected higher vocal variability (as measured by jitter, shimmer, and F0_SD_) to reflect pseudowords rated as pointed, hard, exciting, and bad, while lower variability would be associated with pseudowords rated as rounded, soft, calming, and good.

### Methods

#### Acoustic and voice quality parameters

##### Spectro-temporal parameters

As in Lacey et al. (2020), we chose to measure the speech envelope, spectral tilt, and the FFT. For the detailed calculation of each parameter, please see the Supplementary Material.

###### Speech envelope

The speech envelope measures changes in the amplitude profile over time, largely corresponding to changes in phonemic properties and syllabic transitions (Aiken & Picton, 2008). To the extent that these transitions are abrupt, reflecting stops, affricates, or fricatives (collectively known as obstruents), the speech envelope is discontinuous and uneven (Figure 1A, left panel); but where they are more gradual, reflecting sonorants, the envelope appears more continuous and smoother (Figure 1A, right panel).

###### Spectral tilt

Spectral tilt reflects differences in power across frequencies and is an estimate of the overall slope of the power spectrum, with sampling across the complete utterance. When high frequencies have less power than low frequencies, the power spectrum slopes steeply downward from low to high frequencies (Figure 1B, right panel), flattening out when power is more concentrated in the high frequencies (Figure 1B, left panel).

###### Fast Fourier Transform (FFT)

The FFT derives the frequency components of the speech signal and the variation in their energy over time, thus reflecting the power spectrum of the frequency composition across the duration of the spoken pseudowords. The FFT is illustrated by the spectrogram which shows how power is distributed across frequencies over time; for example, obstruents tend to be reflected in abrupt changes in power as a function of frequency (Figure 1C, left panel) while for sonorants, power varies more gradually with frequency (Figure 1C, right panel).

##### Voice quality parameters

We chose the same voice quality parameters as in our previous study (Lacey et al., 2020): FUF, MAC, mean HNR, and pulse number (these parameters reflect the relative amount of periodicity of the speech signal), together with jitter and shimmer (which reflect the variability of voicing or voice quality during the production of each pseudoword), and F0_SD_. In the present study, we also included the mean F0 as a voice quality parameter, and the duration of the pseudoword. Note that, although extreme values for some of these parameters can indicate vocal pathology (e.g., Brockmann et al., 2011; Ferrand, 2002; Teixeira & Fernandes, 2014), they also vary naturally in a healthy voice as employed here (see Brockmann et al., 2011).

###### Mean harmonics-to-noise ratio (HNR)

This is the ratio between the periodic, or harmonic, portions of the speech signal and the aperiodic, or noise, portions. The mean HNR is thus an estimate of the overall periodicity of the sound expressed in dB (Teixeira & Fernandes, 2014). The noise element arises from turbulent airflow at the glottis when the vocal cords do not close properly (Ferrand, 2002). As noise increases, and therefore, mean HNR decreases, the voice becomes increasingly hoarse or quavery and the speech pattern becomes progressively more uneven (Ferrand, 2002).

###### Mean autocorrelation (MAC)

This is a measure of the periodicity of a signal (Boersma & Weenink, 2012). Periodicity should be high for a long vowel like /u:/, or consonant like /m/, and each successive segment should sound very similar to the one before, i.e. they should be highly correlated. Higher autocorrelation values indicate a smoother voice pattern and/or more voiced segments, while lower values indicate an uneven pattern and/or fewer voiced or periodic segments.

###### Pulse number

This is the number of glottal pulses, i.e. opening and closing of the vocal folds, during production of vowels or voiced consonants measured across the whole utterance (Boersma & Weenink, 2012). An extreme form of phonation, known as pulse register phonation, will help to understand how the pulse number manifests in the voice. In pulse register phonation, rapid glottal pulses are followed by a long, closed phase (Hollien et al., 1977; Whitehead et al., 1984). This results in an audibly uneven speech pattern described as a ‘creaky voice’ (Ishi et al., 2008) or – onomatopoeically – as a ‘glottal rattle’ (Hornibrook et al., 2018). A lower pulse number indicates a more uneven voice pattern, and/or fewer voiced segments, while higher pulse numbers indicate a smoother voice pattern, and/or more voiced segments.

###### Fraction of unvoiced frames (FUF)

The FUF represents the number of unvoiced elements, expressed as the percentage of measurement windows that do not engage the vocal folds (Boersma & Weenink, 2012). The FUF depends on the phonemic content, increasing for those pseudowords that include unvoiced elements, like obstruents, and decreasing for those containing voiced (i.e., periodic) elements, typically long vowels.

###### Shimmer

Shimmer indexes peak-to-peak variation in the amplitude of the glottal waveform (Brockmann et al., 2011). Shimmer reflects vocal variation or instability: low shimmer results in a smooth speech pattern whereas high shimmer results in an uneven speech pattern and manifests as a hoarse voice. We measured local shimmer, defined as the average absolute difference between the amplitudes of consecutive periods, divided by the average amplitude (Boersma & Weenink, 2012).

###### Jitter

Jitter is defined as the frequency variation between consecutive periods and is a measure of voice quality in that it measures variation in the vibration of the vocal cords (Teixeira & Fernandes, 2014). Vocally, high values of jitter manifest as a ‘breaking’ or rough voice. Jitter is typically measured for long vowel sounds, where little frequency variation would be expected. In the production of the pseudowords, increasing jitter reflects increased vocal variation or instability, and perceived vocal roughness. We measured local jitter, defined as the average absolute difference between consecutive periods, divided by the average period (Boersma & Weenink, 2012)^6^.

###### Mean & standard deviation of F0 (mean/F0_SD_)

Mean F0 is the mean fundamental frequency of the speech object while F0_SD_ indicates the variation in the fundamental frequency present in the speech signal (Boersma & Weenink, 2012). F0_SD_ is a measure of vocal inflection, with low values resulting in a flat, monotone voice and high values in a ‘lively’ voice (Kliper et al., 2016).

###### Duration

This is the duration of the speech signal in milliseconds (ms). Although not strictly a voice quality parameter (at least, not here, where there is only one speaker), it is likely influenced by differences between long and short vowels (Kluender et al., 1988; Hillenbrand et al., 1995), phonetic context (Kluender et al., 1988), and speaking rate (Miller & Volaitis, 1989).

### Statistical analysis

As for the articulatory phonetic analyses above, we again used a forced-entry linear regression for each of the 2 scales in each of the 8 domains. The three spectro-temporal parameters (FFT, SE, and ST) consist of multiple measurements per item (see Supplementary Material) therefore we calculated their coefficient of variation (CV) in order to enter these into the regression on the same basis as the other parameters, which consisted of a single measurement per item. For each spectro-temporal parameter, the CV was computed as the standard deviation of the measurements divided by their mean (see Supplementary Material), thus reducing the spectro-temporal parameters to a single value. The other parameters were: eight voice quality predictors: the FUF, MAC, mean HNR, and pulse number (reflecting periodicity), jitter, shimmer, mean and F0_SD_ (reflecting vocal variability), and finally, item duration.

Since all the predictors are continuous variables, positive β values indicate that, as predictor values increase, the outcome values, i.e. ratings, also increase; negative β values indicate an inverse relationship in which rating values decrease as predictor values increase. In this regard, it is important to note that the FUF and MAC relate differently to periodicity: an *increase* in FUF reflects a greater proportion of unvoiced segments and therefore a *decrease* in periodicity. By contrast, an *increase* in MAC primarily reflects a greater proportion of voiced segments and therefore an *increase* in periodicity.

Multicollinearity testing for the full model described above showed that pulse number, FUF, and mean HNR had tolerance values < .2 and also VIF values >10; additionally, the tolerance value for MAC was <.2, although the VIF value was < 10. Accordingly, we re-ran the regression, omitting pulse number and HNR since these had the lowest tolerance values; multicollinearity testing for this revised model showed that there were no tolerance values below .2 (Menard, 1995), with observed values ranging from .23 to .8, and that VIF values ranged from 1.25 to 4.33, well below the threshold value of 10 recommended by Myers (1990: Supplementary Table 6). In addition, the mean VIF of 2.3 was close to 1, as recommended by Bowerman & O’Connell (1990). Thus, we concluded that multicollinearity was not a problem in the revised model. There were no Durbin-Watson test values < 1 or > 3 (Field, 2018), ranging from 1.85 to 2.13 (Supplementary Table 7), thus the assumption of independent errors was also met in the revised model. Visual inspection of histograms and Q-Q plots for standardized residuals indicated that these were normally distributed for all scales. Plots of standardized residuals against standardized predicted values indicated that the assumption of homoscedasticity was also met for all scales.

### Results

#### Shape

Overall, the shape domain had the best model fit between whole-item acoustic parameters and ratings with R^2^ of .30 for the rounded dimension and .56 for the pointed dimension (Table 6).

For the spectro-temporal parameters, the largest effect was for FFT, which contributed to both roundedness and pointedness, and produced the largest change to model fit (Table 6). As the FFT CV increased, ratings increased towards both ‘very rounded’ and ‘very pointed’ (Table 6). This is likely because, although roundedness and pointedness were associated with sonorants and unvoiced obstruents respectively, there was also a general effect of voicing in which roundedness could also be associated with voiced obstruents as in /gobu/ (see the phonetic and phonemic analyses and Tables 4 and 5). Thus, both roundedness and pointedness were associated with obstruents, primarily stops, voiced for roundedness and unvoiced for pointedness (see Table 4) and therefore both were associated with increased variation in the FFT. For roundedness, compare the spectrograms for /mʊmo/ for sonorants and /gubo/ for voiced stops in Supplementary Figures 1a and 6a, respectively; for pointedness, see the spectrogram for the unvoiced stops in /tike/ (Supplementary Figure 1a). ST contributed to the shape domain only in the sense that as its CV increased, indicating that power spread across both low and high frequencies, ratings reduced towards ‘not rounded’. This is consistent with predictions (and see Supplementary Figure 2b), although ST had no significant effect on the pointed scale itself. There was no significant effect of SE for either scale.

For the other parameters, the largest contributor to perception of roundedness was duration: increasing duration attracted ratings towards the ‘rounded’ end of the scale (Table 6). For the voice quality parameters in particular, as periodicity (MAC) increased, ratings tended to increase towards the ‘rounded’ end of the scale. As mean F0 increased, ratings tended to decrease towards the ‘not rounded’ end of the scale. Thus, roundedness was associated with lower F0 as well as greater periodicity and longer duration (Table 6). Against predictions, increasing vocal variability in frequency, as measured by jitter and F0_SD_, was also associated with roundedness; the reason for this is unclear but these parameters were also the smallest contributors to model fit in terms of R^2^Δ. There were no significant effects for either FUF or shimmer.

In terms of voice quality parameters, while FUF had no impact on roundedness ratings, it had the largest effect on pointedness ratings: as FUF increased, reflecting the greater presence of unvoiced segments (and therefore a decrease in periodicity), so ratings increased towards the ‘very pointed’ end of the scale (Table 6). Increasing vocal variability, as measured by shimmer, mean and F0_SD_, was also associated with pointedness (see Parise & Pavani, 2011; Knoeferle et al., 2017). There were no significant effects for either MAC or jitter. Additionally, as duration increased, ratings decreased towards the ‘not pointed’ end of the scale, the inverse of the relation to roundedness; this likely reflects the phonetic content of the items (most rounded: /mʊmo/, 669 ms – most pointed: /tike/, 505 ms) as well as their vocal realization. Thus, pointedness was associated with lower periodicity, increased variability, and shorter duration (Table 6).

#### Roughness

Roughness had the next best model fit with R^2^ of .35 for the smoothness dimension and .42 for the roughness dimension (Table 6).

For the spectro-temporal parameters, as the FFT CV increased ratings on the smoothness scale decreased and ratings on the roughness scale increased; thus, roughness and smoothness were associated with more and less variation in the spectrogram, respectively (Table 6). This reflected the abrupt transitions in the spectrogram for obstruents associated with roughness and the more gradual transitions for sonorants associated with smoothness (see the phonetic and phonemic analyses and Tables 4 and 5: see the spectrograms for /tʃitʃe/ and /mɛli/ in Supplementary Figure 2a). SE contributed to the roughness domain only in the sense that as its CV increased, indicating the discontinuities associated with obstruents, smoothness ratings reduced towards ‘not smooth’. This was consistent with predictions (and see Supplementary Figure 2c), although SE had no significant effect on the roughness scale itself. There was no significant effect of ST for either scale.

For the voice quality parameters, smoothness ratings were affected by periodicity in that as FUF increased, ratings decreased towards the ‘not smooth’ end of the scale; thus, smoothness itself was associated with more, not less, periodicity (Table 6). As variability in amplitude (shimmer) increased, ratings again decreased towards the ‘not smooth’ end of the scale; thus, smoothness itself was associated with less, not more, variability. Smoothness was also reflected in higher mean F0 although the associated β was relatively small. The accompanying changes in R^2^ were small for all these predictors (Table 6). There were no significant effects of jitter, MAC, or F0_SD_. Higher smoothness ratings were also predicted by longer duration.

Periodicity was also important for roughness ratings: as FUF increased (and therefore periodicity decreased) ratings increased towards the ‘very rough’ end of the scale. By contrast, as MAC increased (and therefore periodicity increased), ratings decreased towards the ‘not rough’ end of the scale. Thus, by either measure, roughness was reflected in lower periodicity (Table 6). Consistent with the results for smoothness, as shimmer increased, so roughness ratings increased. While jitter had no significant relationship to smoothness, roughness ratings were significantly associated with increasing jitter (Table 6). There were no significant effects of mean or F0_SD_, or duration.

#### Hardness

Overall, the model fit for hardness was lower than for roughness with R^2^ of .45 for the hardness dimension and .18 for the softness dimension (Table 6).

For the spectro-temporal parameters, as the FFT CV increased, ratings on the hardness scale increased and ratings on the softness scale decreased; thus, hardness and softness were associated with less and more variation in the spectrogram, respectively (Table 6). This reflected the abrupt transitions in the spectrogram for obstruents that were associated with hardness and the more gradual transitions for sonorants that were associated with softness (see the phonetic and phonemic analyses and Tables 4 and 5: see the spectrograms for /kike/ and /mumo/ in Supplementary Figure 3a). Increasing SE CV was associated with softness ratings rather than roughness ratings. At first glance, it is not clear why this was so since the phonetic category analysis above showed that softness was associated with sonorants (see also Table 3 which shows that 5 of the 10 pseudowords with the highest softness ratings contain sonorants), and therefore decreasing CV due to the more continuous speech envelopes. However, the phonetic category analysis ignored positional effects for consonants while the phonemic analysis showed that there were strong effects for the unvoiced fricatives /f/ and /s/ in the second consonant position (Table 5). These appear in /sefi/ and /fɪse/ among the pseudowords with the highest softness ratings (Table 3: in fact, 4 of the 10 highest-rated soft pseudowords contain fricatives, including voiced /v/) and involve increased SE CV. Again, there was no significant effect of ST for either scale.

As for roughness, periodicity was important to hardness: as FUF increased (and therefore periodicity decreased), hardness ratings increased and as MAC increased (and therefore periodicity increased), ratings decreased towards the ‘not hard’ end of the scale. Thus, by either measure, hardness was reflected in lower periodicity (Table 6). Only variability in frequency as measured by jitter was associated with hardness: jitter increased, ratings tended to decrease towards the ‘not hard’ end of the scale. There were no significant results for shimmer, or for either mean or F0_SD_. In addition, as pseudoword duration increased, ratings also decreased towards ‘not hard’, indicating that longer and shorter durations were associated with softness and hardness, respectively.

The only significant voice quality contributors to perception of softness were FUF and duration, which showed complementary effects to their relationship to hardness. As FUF increased (and therefore periodicity decreased, ratings decreased towards the ‘not soft’ end of the scale and, as duration increased ratings increased towards the ‘very soft’ end of the scale (Table 6). Thus, softness was associated with higher periodicity and longer duration.

#### Weight

The model fit values for the weight domain were R^2^ of .24 for the lightness dimension and .26 for the heaviness dimension (Table 6).

For the spectro-temporal parameters, as the FFT CV increased, ratings on the lightness scale decreased and ratings on the heaviness scale increased; thus, lightness and heaviness were associated with less and more variation in the spectrogram, respectively. Again, this reflected more gradual transitions for sonorants that were associated with lightness compared to the more abrupt transitions for obstruents that were associated with heaviness (see the phonetic and phonemic analyses and Tables 4 and 5: see the spectrograms for /mɪle/ and /bεge/ in Supplementary Figure 4a). Increasing SE CV was associated with an increase in lightness ratings, but not heaviness ratings. Lightness was associated with both sonorants (low SE CV) and unvoiced consonants (high SE CV), primarily fricatives (see the phonemic analysis and Table 3 in which sonorants and fricatives are equally represented in the ten highest-rated soft pseudowords; see Supplementary Figure 4c for the sonorants in /mɪle/). Again, there was no significant effect of ST for either scale.

For voice quality parameters in relation to lightness, the largest effect was for periodicity as measured by MAC and FUF. As MAC (and therefore periodicity) increased, ratings also increased towards the ‘very light’ end of the scale; consistent with this, ratings tended towards the ‘not light’ end of the scale as FUF increased (and periodicity decreased: Table 6). There were also small effects in which ratings increased as mean F0 and duration increased. There were no significant effects for jitter, shimmer, or F0_SD_. Thus, lightness ratings were related to more periodicity, higher mean F0, and longer duration.

These results were mirrored in the heaviness dimension: increasing FUF, indicating less periodicity, was accompanied by an increase in ratings towards the ‘very heavy’ end of the scale; as MAC (and therefore periodicity) increased, ratings decreased towards the ‘not heavy’ end of the scale (Table 6). There were also small effects in which higher mean F0 and longer duration were associated with ratings decreasing towards ‘not heavy’, i.e., heaviness itself was associated with lower mean F0 and shorter duration. We had no specific predictions for shimmer, jitter and F0_SD_, and these turned out not to have any significant influence.

The relationship between measures of periodicity and weight ratings was consistent with our phonetic category analysis (see above), in which lightness ratings were associated with sonorants, which are always voiced; by contrast, obstruents, which can be either voiced or unvoiced, were strongly associated with heaviness ratings and the contribution of voicing was relatively small. Mean F0 was lower for heavy, compared to light, pseudowords, in line with prediction. Against our prediction, shorter items were rated as heavier than longer items. It could perhaps be argued that shorter words are ‘denser’ – since density is mass per unit volume, for two-syllable (mass) words, the shorter (unit volume) they are, the denser and therefore apparently heavier, they are. This would need to be confirmed by reference to ratings for an appropriate density scale.

#### Arousal

The next smallest overall model fit was for arousal with R^2^ of .24 for the calming dimension and .17 for the exciting dimension (Table 6).

For the spectro-temporal parameters, FFT contributed to both calming and exciting ratings: as the FFT CV increased, ratings decreased towards both ‘not calming’ and ‘not exciting’ (Table 6). The reason for this is not clear, given that the calming and exciting dimensions were associated with sonorants and, mostly unvoiced, obstruents respectively (see the phonetic and phonemic analyses and Tables 4 and 5; also compare the spectrograms for /mumo/ and /zεʤi/ in Supplementary Figure 5a). ST contributed to the arousal domain only in the sense that as its CV increased, indicating that power spread across both low and high frequencies, ratings increased towards ‘very exciting’. This is consistent with predictions (and see Supplementary Figure 5b), although ST had no significant effect on the calming scale. There was no significant contribution for SE to either scale.

The largest effect of voice quality on calming ratings was that related to periodicity: increasing FUF (less periodicity) reduced ratings towards ‘not calming’, while increasing MAC (more periodicity) increased ratings towards ‘very calming’ (Table 6). Duration produced the next largest effect with ratings increasing towards ‘very calming’ as duration increased. There was also a small, counter-intuitive, effect for jitter in which increasing variability in frequency led to increased perception of calmness. There were no significant effects for shimmer, mean F0, or F0_SD_.

In contrast to the calming dimension, neither FUF nor MAC had significant associations with the exciting dimension. Instead, vocal variability contributed to perception of pseudowords as exciting: increasing variability in amplitude (shimmer) was associated with ratings increasing towards ‘very exciting’, while increasing variability in frequency (jitter) gave ratings towards ‘not exciting’, again, counter-intuitively (Table 6). Higher mean F0 also produced ratings increasing towards ‘very exciting’ and the single largest R^2^Δ. There were no significant effects for F0_SD_ or duration.

The measures of periodicity presumably reflect the change from voiced to unvoiced consonants for pseudowords going from calming to exciting. Increasing variability in frequency and amplitude from cycle to cycle of the speech waveform, as measured by jitter and shimmer respectively, reflected the change in ratings from calming to exciting.

#### Brightness

The model fit values for the brightness and darkness dimensions were R^2^ of .23 and .13, respectively (Table 6).

The most important spectro-temporal parameter for the brightness domain was ST: as its CV increased, ratings increased towards ‘very bright’ and decreased towards ‘not dark’, consistent with the prediction that brightness and darkness would be associated with higher and lower frequencies, respectively (Table 6; see also Supplementary Figure 6b). The FFT and SE CVs only had significant effects on the brightness scale and only in the sense that as variability in these parameters increased, ratings decreased towards ‘not bright’, i.e., dark. Obstruents were associated with both brightness and darkness, the only difference (as for the shape domain) being voicing – unvoiced for brightness and voiced for darkness (see the phonetic and phonemic analyses together with Tables 4 and 5; and compare the spectrograms and speech envelopes for /kike/ and /gobo/ in Supplementary Figure 6). Thus, although decreasing brightness ratings in response to increasing FFT and SE variation make sense, it is not clear why increasing FFT and SE variation did not positively predict both scales.

Mean F0 and FUF were the best predictors of brightness rating among the voice quality parameters: increasing FUF, i.e. decreasing periodicity, predicted ratings increasing towards ‘very bright’, as did higher F0 (Table 6). There were smaller, contrasting, effects of vocal variability in which ratings increased towards ‘very bright’ as shimmer increased but decreased towards ‘not bright’ as jitter increased (Table 6). There were no significant effects of MAC or F0_SD_. Duration was also a significant predictor in that longer duration was associated with ratings towards the ‘not bright’ end of the scale.

Mean F0 and FUF had the largest effect on darkness ratings: but higher mean F0 and increasing FUF, i.e. decreasing periodicity, predicted ratings decreasing towards ‘not dark’ (Table 6), i.e. darkness itself was associated with lower mean F0. As jitter (variability in frequency) increased, ratings increased towards ‘very dark’. There were no significant effects for shimmer, MAC, F0_SD_, or duration.

In line with predictions, high/low mean F0 were associated with bright/dark pseudowords, respectively (see Tzeng et al., 2018). There was a small effect of duration in which longer pseudowords were rated as less bright, i.e., darker, which is consistent with Tzeng et al. (2018).

#### Valence

The valence domain again had the second worst model fit with R^2^ of .18 for the goodness dimension and .16 for the badness dimension (Table 6) and very few parameters contributed to the model.

The spectro-temporal parameters had only small effects in the valence domain. As the FFT CV increased, ratings decreased towards ‘not good’ and increased towards ‘very bad’. This reflected more gradual transitions for sonorants that were associated with goodness compared to the more abrupt transitions for obstruents that were associated with badness (see the phonetic and phonemic analyses and Tables 4 and 5: see the spectrograms for /mine/ and /pupo/ in Supplementary Figure 7). As the SE CV increased, ratings increased towards ‘very bad’, again reflecting the associated obstruents, but there was no significant effect for SE on the goodness scale. Increased ST CV was associated with ratings increasing towards ‘very good’, but there was no significant effect for ST on the badness scale.

For the goodness dimension there were small effects in which increasing shimmer was associated with ratings that decreased towards ‘not good’ while increasing mean F0 meant that ratings increased towards ‘very good’. There were no significant effects for FUF, jitter, MAC, F0_SD_, or duration for the goodness dimension and none of the voice quality parameters contributed significantly to the badness dimension.

#### Size

The size domain again had the worst model fit with R^2^ of .19 for the smallness dimension and .12 for the bigness dimension (Table 6).

The most important spectro-temporal parameter for the size domain was SE: as the SE CV increased, ratings increased towards ‘very small’ and decreased towards ‘not big’. The same pattern was observed for ST (Table 6 and compare the speech envelops for /pete/ and /vʊʤo/ in Supplementary Figure 8). Bigness was primarily associated with voiced stops while smallness was primarily associated with unvoiced obstruents, both stops and af/fricatives (see the phonetic and phonemic analyses and Tables 4 and 5). Bigness was also associated with back rounded vowels, while vowel rounding was non-significant for smallness.

For the smallness dimension, increasing FUF (less periodicity) reduced ratings towards ‘not small’, while increasing MAC (more periodicity) increased ratings towards ‘very small’ (Table 6). There were also small effects in which higher F0 increased ratings towards ‘very small’, while increasing duration reduced ratings towards ‘not small’. There were no significant effects for jitter, shimmer, or F0_SD_.

For the bigness dimension, FUF and MAC had the opposite effect to that for the smallness scale. Increasing FUF (less periodicity) increased ratings towards ‘very big’, while increasing MAC (more periodicity) reduced ratings towards ‘not big’ (Table 6). There were also small effects in which increasing jitter reduced ratings towards ‘not big’ and increasing duration increased ratings towards ‘very big’. We therefore replicated the finding that short and long pseudoword duration reflect small and large size respectively (Knoeferle et al., 2017). There were no significant effects for shimmer, nor for the mean or F0_SD_.

### Discussion

As with the phonetic category and phoneme analyses, the patterns of acoustic parameters that predicted ratings were largely domain-specific and occasionally scale-specific.

FFT was the most influential of the spectro-temporal parameters since it discriminated between the opposing dimensions of four of the eight domains: roughness, hardness, weight, and valence (Table 6). The FFT CV was significantly positively weighted for the roughness, hardness, heaviness, and badness dimensions – all characterized by obstruents (Tables 4 and 5) – and negatively weighted for the smoothness, softness, lightness, and goodness dimensions, which were mostly characterized by sonorants (Tables 4 and 5). These distinct phonetic features resulted in more vs. less variation in the FFT, as indexed by its CV, for obstruents vs. sonorants, respectively. For other domains, the FFT CV was only related to one dimension, being positively weighted for bigness (characterized by voiced obstruents, Tables 4 and 5) and negatively weighted for brightness (largely characterized by unvoiced consonants, Tables 4 and 5) but not significantly associated with either smallness or darkness. However, FFT did not discriminate between the dimensions of the shape and arousal domains being positively weighted for both roundedness and pointedness and negatively weighted for both the calming and exciting scales: we return to this as a limitation in the General Discussion below.

ST discriminated between dimensions for two domains, brightness and size, being positively weighted for brightness and smallness and negatively weighted for darkness and bigness (Table 6). This is consistent with the fact that brightness and smallness are both associated with higher F0 while darkness and bigness are associated with lower F0 (Table 6 and see below), and so spectral power spreads from lower to higher frequencies as ratings change from dark to bright and big to small. For other domains, ST was only related to one dimension, being positively weighted for the lightness, exciting, and goodness dimensions, all associated with front unrounded vowels (Tables 4 and 5) and with higher F0 (Table 6 and see below) and negatively weighted for the roundedness dimension (associated with back rounded vowels and lower F0), but was not significantly associated with heaviness, calmness, or badness. However, ST was not significantly related to either dimension of the roughness or hardness domains.

SE only discriminated between dimensions for the size domain, being positively weighted for smallness, which was largely associated with unvoiced stops, and negatively for bigness, which was characterized by voiced obstruents and back rounded vowels (Tables 4 and 5). For other domains, SE was only related to one dimension, being positively weighted for the softness, lightness, and badness dimensions and negatively weighted for smoothness and brightness. However, these dimensions had varying phonetic profiles (Tables 4 and 5): softness was mostly associated with sonorants and rounded vowels, lightness with both sonorants and unvoiced affricates but unrounded vowels, and badness with obstruents and rounded vowels. SE was not significantly related to either dimension of the shape or arousal domains.

In terms of voice quality parameters, FUF was the most influential in that it discriminated between smooth/rough, hard/soft, light/heavy, bright/dark and small/big. For these five domains, FUF was significantly positively weighted for one dimension and negatively weighted for its opposite. For example, as FUF increased, and therefore periodicity decreased, ratings increased on the roughness scale but decreased on the smoothness scale (Table 6); thus, smoothness and roughness were associated with more and less periodicity, respectively. For other scales, however, FUF was significantly positively associated with pointedness, and negatively associated with calmness, but was not significantly associated with either the roundedness or exciting scales. Thus, higher FUF (decreasing periodicity) positively defined pointedness, but not roundedness. FUF defined arousal only in that higher ratings on the calming scale were associated with lower FUF (increased periodicity) but FUF had no influence on the exciting scale. FUF was not significantly related to the good/bad dimensions of the valence domain.

By the same token, duration was the next most influential parameter since it significantly discriminated between the opposing dimensions of four domains: shape, hardness, weight, and size (Table 6). For other domains, however, the predictive power of duration was limited to a single dimension: it was significantly positively weighted for smoothness and calmness but unrelated to either the roughness or exciting dimensions. Additionally, duration was negatively weighted for brightness ratings such that, as duration increased, brightness ratings decreased, but duration was not significantly associated with darkness ratings. Like FUF, duration was not significantly related to either the good or bad dimensions of the valence domain.

Mean F0 discriminated between rounded/pointed, light/heavy and bright/dark (Table 6) but for other domains, mean F0 was positively associated with the smoothness, exciting, goodness, and smallness dimensions without being significantly associated with the roughness, calmness, badness, or big dimensions. Mean F0 was not significantly related to either dimension of the hardness domain.

MAC discriminated between light/heavy and small/big (Table 6). For other domains, MAC was positively associated with the roundedness and calmness dimensions, i.e., as MAC (and therefore periodicity) increased so ratings increased towards either ‘very rounded’ or ‘very calming’. Additionally, MAC was negatively associated with the roughness and hardness dimensions so that, as MAC and periodicity increased, ratings decreased towards either ‘not rough’ or ‘not hard’. However, MAC was not significantly associated with the opposing dimension in each case, i.e., the pointedness, exciting, smoothness, or softness dimensions. MAC was not significantly related to either dimension of the brightness or valence domains.

Jitter was one of the least influential voice quality parameters, discriminating between the dimensions of only two domains: calming/exciting and bright/dark (Table 6). For other domains, jitter was positively associated with roundedness without being significantly associated with pointedness. Additionally, jitter was negatively associated with the roughness, hardness, and big dimensions so that, as frequency variability increased, ratings decreased towards ‘not rough/hard/big’. However, jitter was not significantly related to the opposing dimension in each case, i.e., the smoothness, softness, or smallness dimensions. Jitter was not significantly related to either dimension of the weight or valence domains.

Shimmer discriminated only between the smooth/rough dimensions of the roughness domain (Table 6). For other domains, increasing shimmer was positively associated with increasing ratings on the pointedness, exciting, and brightness scales, without being significantly associated with roundedness, calmness, or darkness. Additionally, shimmer was negatively associated with the goodness scale so that, as amplitude variability increased, ratings decreased towards ‘not good’ but was not significantly related to badness. Shimmer was not significantly related to either dimension of the hardness, weight or size domains.

Finally, F0_SD_ was the least influential parameter, discriminating only between rounded/pointed and not being significantly associated with either dimension of any other domain.

We can summarize each domain in terms of those parameters that discriminated between the opposing dimensions of each domain. Here, shape ratings were mainly driven by mean F0, F0_SD_ and duration. Ratings for the roughness aspect pf texture were driven by FFT, FUF and shimmer, and for the hardness aspect by FFT, FUF and duration. In the weight domain, ratings were predicted by FFT, FUF, MAC, mean F0, and duration. For the arousal domain, only jitter appeared to discriminate between the calming and exciting dimensions. For the brightness domain, ratings were driven by ST, FUF, jitter, and mean F0. The FFT was the sole parameter to discriminate between the good and bad dimensions of the valence domain. Finally, in the size domain ratings were driven by SE, ST, FUF, MAC, and duration.

Finally, some domains raised interesting questions. For example, the reason for the association of hardness and softness with shorter and longer duration respectively, is not clear. Perhaps, as with the shape domain (see above) it is simply the preponderance of inherently shorter stop sounds in the harder pseudowords (hardest: /kike/, 533 ms) as compared with the concentration of longer sonorant sounds for the softer pseudowords (softest: /mumo/, 583 ms). Alternatively, it may be that hardness and softness are associated with percussive events that are more or less auditorily distinct respectively (for example, a hammer striking a nail compared to hands plumping up a cushion), and that similarly vary in duration. Aside from roughness and hardness, the other aspect of texture that could potentially be iconically mapped is stickiness, with dimensions of sticky versus slippery, although this aspect has not been robustly established from a perceptual standpoint (Hollins et al., 2000).

Although some predictions for the arousal domain could have been made from voice stress analyses (e.g., Van Puyvelde et al., 2018), a cautionary note is that this previous work was carried out with real-life emergency communications where there is a need to speak more clearly in order to avoid miscommunication and error and also presumably more significant levels of arousal in those contexts, regardless of phonetic content. We found that jitter and shimmer were associated with calmness and excitement, respectively, while general arousal produces an increase in both these voice quality measures (Van Puyvelde et al., 2018). However, in the context of emergency communications, speaking more clearly and loudly might result in a decrease in such vocal variability (see Brockmann-Bauser et al., 2018). An additional point is that our arousal scale was positively valenced as a whole: both ‘calm’ and ‘exciting’ are in the top 10% of items at the positive end of the valence scale in Warriner et al. (2013). A restful/stressful scale might have produced a clearer delineation between positive and negative dimensions.

Like the arousal domain, the valence domain could have been interpreted in several different ways other than the good/bad dimension (see the Limitations section in the General Discussion). Whether different aspects of the same domain share common phonetic and/or acoustic characteristics would be an interesting area for further research. Emotional state is another context in which both valence and arousal are relevant and both may increase from, for example, sad to happy. In this case, F0_SD_ might mediate an iconic relationship. F0_SD_ is a measure of the variation in the fundamental frequency and thus of vocal inflection (Kliper et al., 2016): low F0_SD_ manifests as a monotone voice and higher F0_SD_ as a livelier voice, each reflecting the speaker’s emotional state (Kliper et al., 2016).

## SEGMENTAL ACOUSTIC ANALYSIS

### Introduction

Our predictions for which segmental acoustic characteristics would be associated with each meaning domain were informed in part by the preceding analyses. We both combined findings from the whole-word acoustic and phonetic category analyses and examined the effects of segment position (e.g., C1, V1, C2, V2) by reference to the phoneme analysis.

Measures of mean F0, intensity, and duration for each consonant segment and the formant frequencies for the vowels were hypothesized to depend primarily on the phoneme content of each pseudoword and the influence of adjacent phonemes on acoustic realization. In general, consonants varying in manner and place of articulation have acoustic consequences resulting in breaks in intensity (e.g., obstruents, particularly voiceless stops; Ladefoged & Johnson, 2011), differences in characteristic durations (e.g., longer duration for fricatives [Umeda, 1977; van Son & van Santen, 2005] or longer durations for tense vowels and back vowels [Hillenbrand et al., 1995; Leung et al., 2016; Peterson & Lehiste, 1960; Umeda, 1975]), and in effects on F0 for adjacent vowels (e.g., vowels following unvoiced stops have slightly higher F0; Xu & Xu, 2021). Vowel quality is reflected in formant structure, with F1 and F2 formant frequencies reflecting height and backness and F3 being one indicator of lip rounding (Ladefoged & Johnson, 2011). In addition, English vowels vary in characteristic F0 and duration (Umeda, 1975; Whalen & Levitt, 1995). All of these factors were expected to contribute in meaning-specific ways to listeners’ judgments of iconicity. Moreover, the ability to evaluate positional effects added information to the whole word acoustic analyses and served as a test of a possible intermediary representation between linguistic predictors (phonemes and phonetic categories) and acoustic and vocal properties of the spoken pseudowords.

#### Shape

In the shape domain, we expected associations between lower F0 values and roundedness ratings and higher F0 values and pointedness ratings across positions, reflecting the back rounded vowels and voiced consonants that are associated with roundedness and the voiceless consonants and high front vowels associated with pointedness (McCormick et al., 2015; Shadle, 1985; see also Table 4). For duration, we expected either that roundedness and pointedness would be associated with segments having shorter and longer durations, respectively, reflecting findings for the characteristic durations of the respective consonant segments (Crystal & House, 1982, 1988; Umeda, 1977) or the reverse, reflecting findings for durations of vowel segments (Hillenbrand, et al., 1995; Umeda, 1975). The voiceless stops and fricatives associated with pointedness ratings have been shown to have characteristically longer durations relative to the sonorants and voiced consonants associated with roundedness ratings (Umeda, 1977; van Son & van Santen, 2005; see Figure 1A; Supplementary Table 4). However, the back vowels associated with roundedness ratings are characteristically longer in duration than the front vowels associated with pointedness judgments (see Figure 1A; Supplementary Table 4; Umeda, 1975). Because pointedness ratings are associated with voiceless stops (see Figure 2A; Supplementary Table 7), we expected intensity values for the consonants to be lower for pointedness ratings relative to roundedness ratings, reflecting lower intensity values for stops and fricatives relative to sonorants (Ladefoged & Johnson, 2011; Ohala, 1992). We also expected higher F1 and lower F2 (lower vowel height and backness) to be associated with roundedness ratings and lower F1 and higher F2 (higher vowel height and frontness) to be associated with higher pointedness ratings. Lower F3 reflecting lip rounding was expected to be associated with higher roundedness ratings (Hillebrand et al, 1995; Ladefoged & Johnson, 2011; see Table 4). With respect to positional effects, we expected vowel formants in V1 position to affect ratings to a greater extent than in V2 position, assuming vowel reduction in utterance-final positions, and acoustic correlates of consonant properties for both C1 and C2 positions to influence shape ratings given these segments are syllable-initial in both positions (see Figure 2A; Supplementary Table 7; Ladefoged & Johnson, 2011).

#### Roughness

For the roughness domain, we did not expect strong associations between F0 values and smoothness or roughness ratings (see Figure 4A; Supplementary Table 10). For duration, the presence of stops and fricatives in pseudowords with higher roughness ratings should result in longer segment durations and the presence of sonorants in pseudowords rated as smooth should result in shorter segment durations (Umeda, 1977; van Son & van Santen, 2005). However, as for shape, the vowel composition of pseudowords would result in shorter durations for the front, unrounded vowels associated with roughness ratings (see Figure 2A; Supplementary Table 7; Umeda, 1975). Intensity values were expected to be higher in pseudowords rated as more smooth reflecting higher intensity sonorants and lower in pseudowords rated as rough reflecting lower intensity fricatives and stops (Ladefoged & Johnson, 2011; Ohala, 1992). To the extent that vowels will be associated with roughness ratings, (see Figures 1A, 2A; Supplementary Tables 4 and 7), we expected lower F2 (back vowels) to be associated with smoothness ratings and higher F2 (front vowels) to be associated with higher roughness ratings. Lower F3 reflecting lip rounding was expected to be associated with higher smoothness ratings. As above, with respect to positional effects, in general, we expected contributions to roughness ratings from consonants at both C1 and C2, and stronger contributions from vowels at V1 compared to V2 (see Figure 2A; Supplementary Table 7; Ladefoged & Johnson, 2011).

**Figure 4A:**
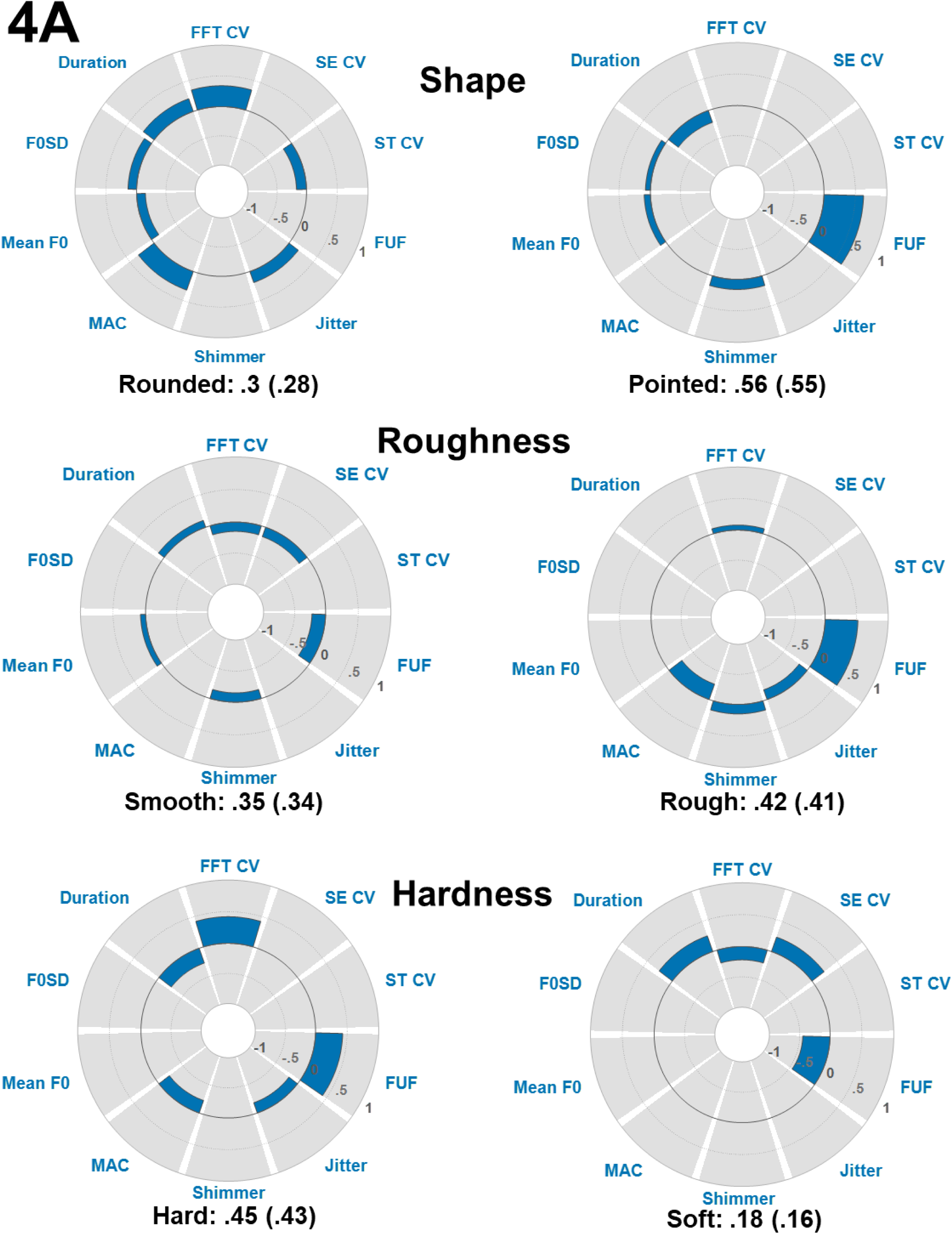
Whole-item acoustic regression results for the opposing scales of the shape, roughness, and hardness domains. Interpretation as for Figure 1A. For the complete results, see Supplementary Table 10.

#### Hardness

Hardness ratings were not expected to be related to overall F0 (see Figure 4A; Supplementary Table 10). We did expect an association of hardness and softness with longer and shorter duration segments respectively, reflecting the phonemic composition of the hard and soft pseudowords, with longer duration stops and af/fricatives associated with hardness and shorter duration sonorants associated with softness (Figure 1A; Supplementary Table 4). Again, however, we would expect the opposite pattern for the vowels based on associations of back vowels with ratings of softness. Intensity values were expected to be lower in pseudowords rated as more hard reflecting low intensity voiceless stops and af/fricatives and higher in pseudowords rated as soft reflecting higher intensity sonorants. We also expected lower F2 (back vowels) to be associated with softness ratings and higher F1/F2 (high front vowels) to be associated with higher hardness ratings. Lower F3, reflecting lip rounding, was expected to be associated with higher softness ratings. With respect to positional effects, in general, we expected contributions to hardness and softness ratings from both consonant positions and stronger contributions to softness ratings for V1 position (see Figure 2A; Supplementary Table 7).

#### Weight

For the weight domain, we expected associations between higher F0 values and lightness ratings and lower F0 values for heaviness ratings across positions, reflecting the high front vowels and voiceless consonants that are associated with lightness and the lower back vowels and voiced consonants associated with heaviness (see Figure 1B; Supplementary Table 4). For duration, from the whole word analysis (Figure 4A; Supplementary Table 10), we expected that lightness and heaviness should be associated with longer and shorter pseudoword durations, respectively. However, based on the phonetic category analysis (see Figure 1B; Supplementary Table 4), we would expect that lightness (sonorants and front vowels) and heaviness (stops, fricatives, and back vowels) should be associated with shorter and longer pseudoword durations, respectively. Similarly, because lightness ratings were associated with sonorants and heaviness with stops and fricatives, we expected intensity values to be higher for lightness ratings relative to heaviness ratings. We also expected higher F1 and lower F2 (lower vowel height and backness) to be associated with heaviness ratings and lower F1 and higher F2 (higher vowel height and frontness) to be associated with higher lightness ratings. Lower F3 reflecting lip rounding was expected to be associated with higher heaviness ratings. With respect to positional effects, we expected vowel formants in V1 position to affect ratings to a greater extent than in V2 position and acoustic correlates of consonant properties for both C1 and C2 positions to influence weight ratings (see Figure 2B; Supplementary Table 7).

#### Arousal

Higher exciting ratings, relative to calming ratings, were expected to be related to higher F0 values, lower intensity values, and shorter durations, perhaps reflecting front vowels, stops, and fricatives, while higher calming ratings were expected to be related to lower F0, higher intensity values, and longer durations, perhaps reflecting back vowels and sonorants (Ma & Thompson, 2015; Shadle, 1985; see Figures 1B and 4B; Supplementary Tables 4 and 10) We also expected higher F1/lower F2 (low back vowels) to be associated with calming ratings and lower F1/higher F2 (high front vowels) to be associated with higher exciting ratings. Higher F3 reflecting lip rounding was expected to be associated with higher calming ratings. With respect to positional effects, in general, we expected contributions to hardness and softness ratings from both consonants and the first vowel position (see Figure 2B; Supplementary Table 7).

#### Brightness

For the brightness domain, we expected associations between higher brightness ratings and higher F0 values, higher intensity, and shorter durations and associations between higher darkness ratings and lower F0 values, lower intensity, and longer durations (see Marks, 1987; Figure 4B; Supplementary Table 10), reflecting high front vowels and voiceless consonants in bright pseudowords and the lower back vowels and voiced consonants in dark pseudowords (see Figure 1B; Supplementary Table 4). We also expected higher F1 and lower F2 (lower vowel height and backness) to be associated with darkness ratings and lower F1 and higher F2 (higher vowel height and frontness) to be associated with higher brightness ratings. Lower F3, reflecting lip rounding, was expected to be associated with higher darkness ratings. With respect to positional effects, based on the phoneme analysis (see Figure 2B; Supplementary Table 7), we expected vowel formants in V1 position to affect ratings to a greater extent than in V2 position and acoustic correlates of consonant properties in C2 positions to influence brightness ratings to a greater extent than consonant properties in C1.

**Figure 4B:**
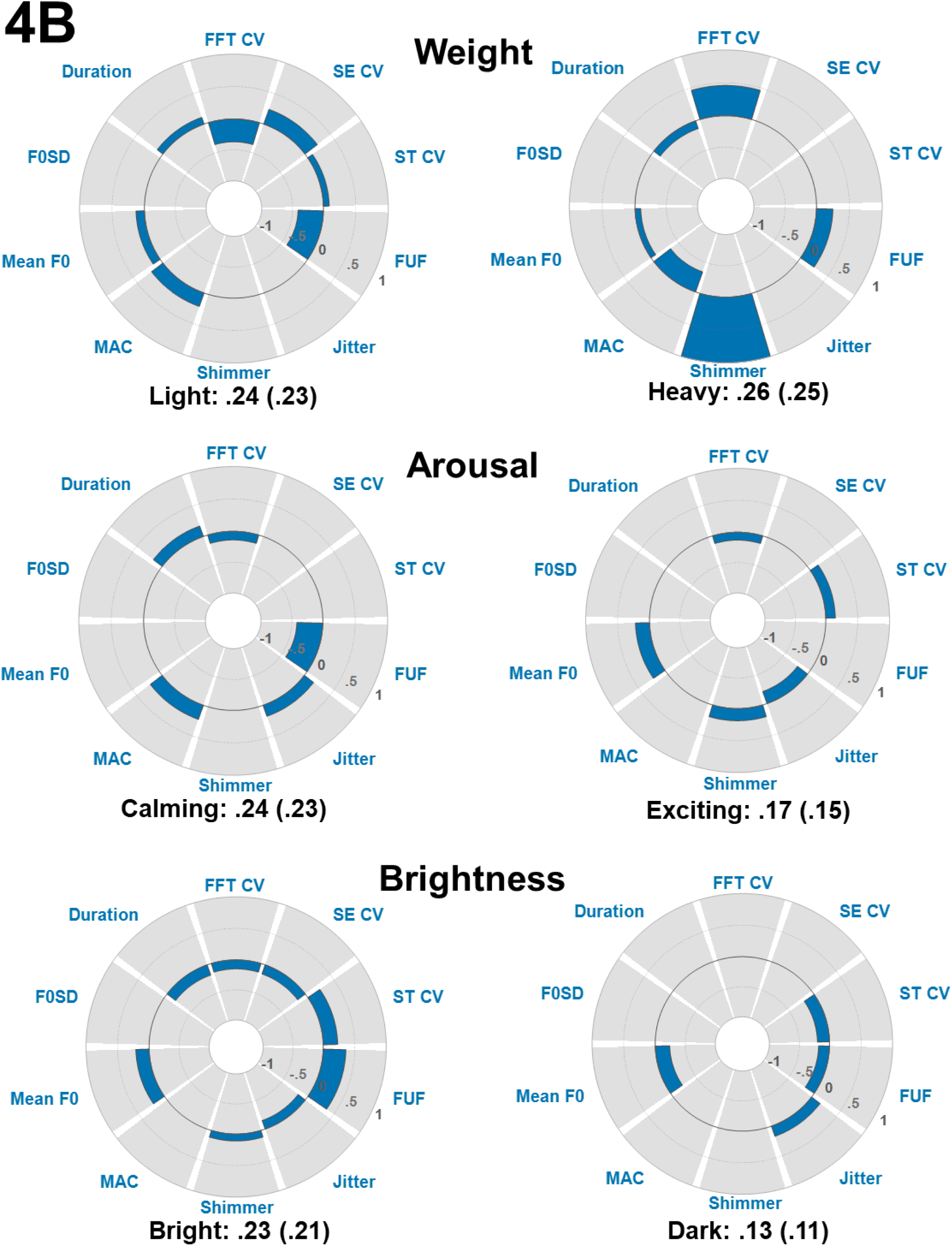
Whole-item acoustic regression results for the opposing scales of the weight, arousal, and brightness domains. Interpretation as for Fig. 1A. For the complete results, see Supplementary Table 10.

#### Valence

We expected associations between higher F0 values and goodness ratings and lower F0 values and badness ratings, reflecting the high front vowels that are associated with goodness and the lower back vowels associated with badness (see Figure 1C; Supplementary Table 4). Based on the whole word acoustic analysis, we did not expect overall associations between pseudoword duration and goodness or badness ratings (Figure 4C; Supplementary Table 10). Because goodness ratings were associated with sonorants and badness with stops and fricatives, we expected consonant segment duration values to be longer and intensity values to be higher for goodness ratings relative to badness ratings. In contrast, because goodness ratings were associated with high front vowels and badness with rounded back vowels, we expected vowel segment duration values to be shorter for goodness ratings relative to badness ratings. We also expected higher F1 and lower F2 (lower vowel height and backness) to be associated with badness ratings and lower F1 and higher F2 (higher vowel height and frontness) to be associated with higher goodness ratings. Lower F3 reflecting lip rounding was expected to be associated with higher badness ratings. With respect to positional effects, we expected vowel formants in V1 position to affect ratings to a greater extent than in V2 position and acoustic correlates of consonant properties for both C1 and C2 positions to influence valence ratings (see Figure 2C; Supplementary Table 7).

**Figure 4C:**
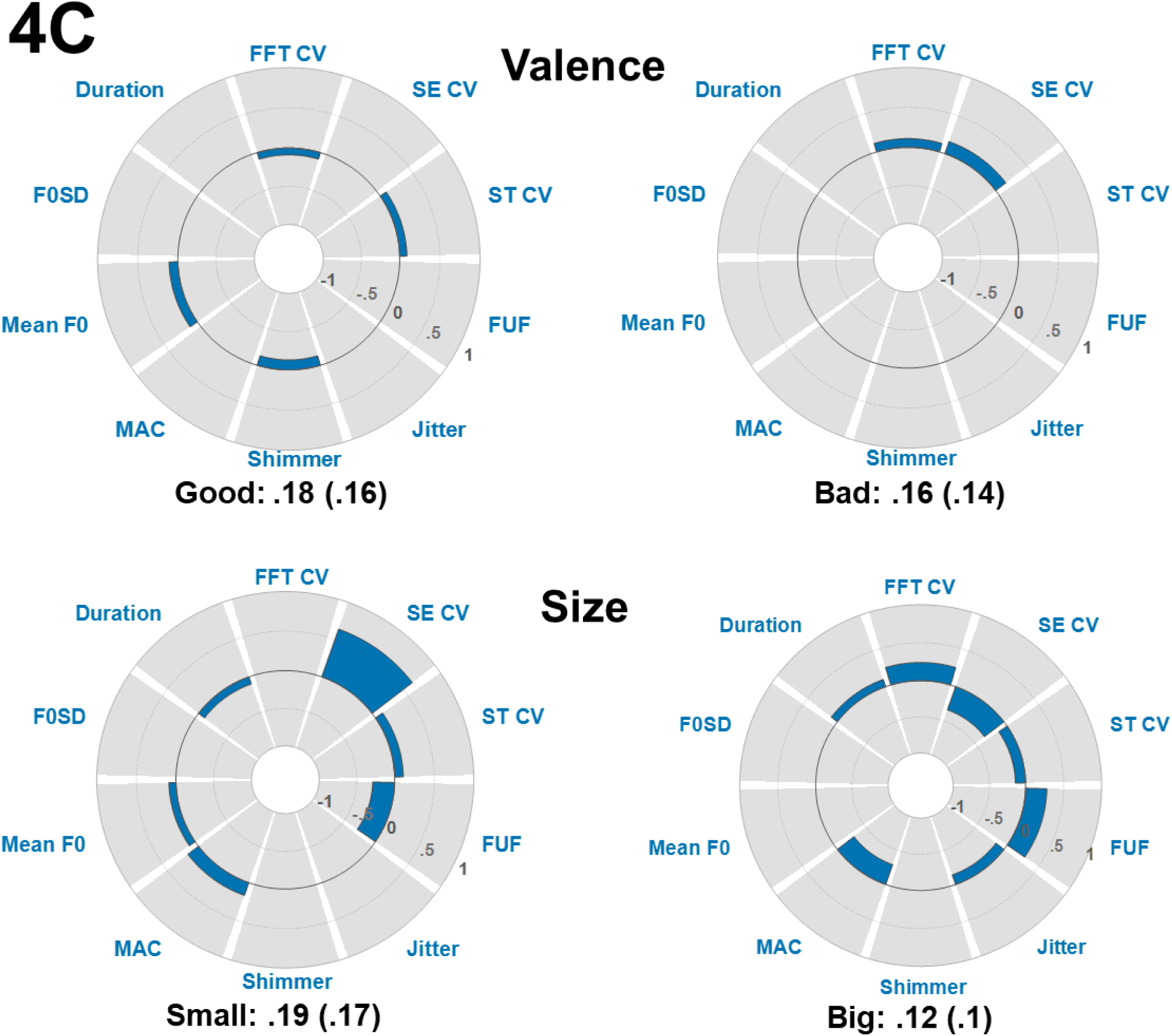
Whole-item acoustic regression results for the opposing scales of the valence and size domains. Interpretation as for Fig. 1A. For the complete results, see Supplementary Table 10.

#### Size

Higher bigness ratings were expected to be related to lower F0 and higher intensity values, and relatively longer durations, perhaps reflecting both the greater presence of back rounded vowels stops and fricatives (Umeda, 1977; van Son & van Santen, 2005; Figures 1C and 4C; Supplementary Tables 4 and 10), relative to smallness ratings, which exhibited fewer associations overall with phonetic features or particular phonemes. We also expected higher F1/lower F2 (low back vowels) to be associated with bigness ratings. Higher F3 reflecting lip rounding was also expected to be associated with higher bigness ratings. With respect to positional effects, in general, we expected larger contributions to bigness and smallness ratings from the second consonant and the first and second vowel positions, based on findings the phoneme analysis (see Figure 2C; Supplementary Table 7).

#### Summary of domain-specific predictions

To summarize, we expected that pseudowords rated as pointed, light, exciting, bright, good, and small would be associated with segments having a higher F0 than those rated as rounded, heavy, calming, dark, bad, and big. Pseudowords rated as rounded, smooth, soft, light, exciting, bright, good, and big should exhibit higher intensity values, whereas those rated as pointed, rough, hard, heavy, calming, dark, bad, and small should exhibit lower intensity values. For duration, we expected that shorter durations would be associated with ratings of rounded, smooth, hard, light, bright, and small and that longer durations would be associated with ratings of pointed, rough, soft, heavy, dark, and big, depending on the relative contributions of consonant and vowel durations. With respect to associations with formant frequencies, we expected higher F1 and lower F2 (and lower F3) to be associated with ratings of rounded, smooth, soft, heavy, calming, dark, bad, and big. Lower F1 and higher F2 (and higher F3) were expected to be associated with ratings of pointed, rough, hard, light, exciting, bright, good, and small. Finally, with respect to positional effects, we expected contributions to meaning ratings to be primarily driven by the first vowel and the second consonant.

### Methods

In order to determine if the acoustic correlates of individual CVCV segments comprising the auditory pseudowords better accounted for listeners’ judgments of iconicity across meaning domains relative to other characterizations of pseudoword structure, each auditory pseudoword was segmented and analyzed in Praat (Boersma & Weenink, 2006) using the textgrid function. For each pseudoword, portions of the speech signal associated with each of the four component phonemes (C1, V1, C2, and V2) were identified by a single coder and acoustic features of those segments were calculated. To determine segmentation for the acoustic analysis, both the spectrogram and waveform representations of each pseudoword were inspected. Segmental boundaries for each syllable were operationalized as the approximate timepoint when the vocalic portion of each syllable became steady state. Likewise, the boundary between syllables, V1-C2 was coded as the approximate timepoint when voicing ceased or a change in formant structure occurred. Because the vowels occurred in a variety of consonant contexts, criteria for determining vowel onset and offset were taken from Munson and Solomon (2004).

Once each of the four phonetic segments were identified, acoustic measurements were averaged across each segment using the voice report function in Praat. Mean fundamental frequency (F0; Hz), intensity (db SPL) and duration (ms) were calculated for each consonant and vowel segment. Because the presence or absence of clear formant structure for the consonants would be unreliable, first, second, and third formant center frequencies (F1, F2, F3) were calculated for the vowels only.

### Statistical analysis

We performed a multiple regression analysis using the acoustic parameters for each phoneme position, C1 (F0, intensity, duration), V1 (F0, intensity, duration, F1, F2, F3), C2 (F0, intensity, duration), and V2 (F0, intensity, duration, F1, F2, F3), as predictors of ratings for each meaning domain (e.g., rounded, pointed). Other aspects of the analysis are as described above for the phonetic category analysis.

Multicollinearity testing for the full model described above showed that F2 in both the first and second vowel positions had tolerance values < .2 (Menard, 1995) but since their VIF values were < 10 (Myers, 1990), we opted to retain these predictors rather than exclude them. Thus, observed tolerance values ranged from .14 to .98, and VIF values ranged from 1.02 to 7.26, below the threshold value of 10 recommended by Myers (1990: Supplementary Table 11). In addition, the mean VIF of 2.14 was close to 1, as recommended by Bowerman & O’Connell (1990). Thus, we concluded that multicollinearity was not a problem in this model. There were no Durbin-Watson test values < 1 or > 3 (Field, 2018), ranging from 1.85 to 2.14 (Supplementary Table 12), thus the assumption of independent errors was also met. Visual inspection of histograms and Q-Q plots for standardized residuals indicated that these were normally distributed for all scales. Plots of standardized residuals against standardized predicted values indicated that the assumption of homoscedasticity was also met for all scales.

Figures 5A-C and Supplementary Table 13 report R^2^ values for each meaning domain with accompanying β values and changes to the model as a function of the addition of each predictor. Positive and negative β again indicated that the predictor and ratings were either positively or negatively related.

**Figure 5A:**
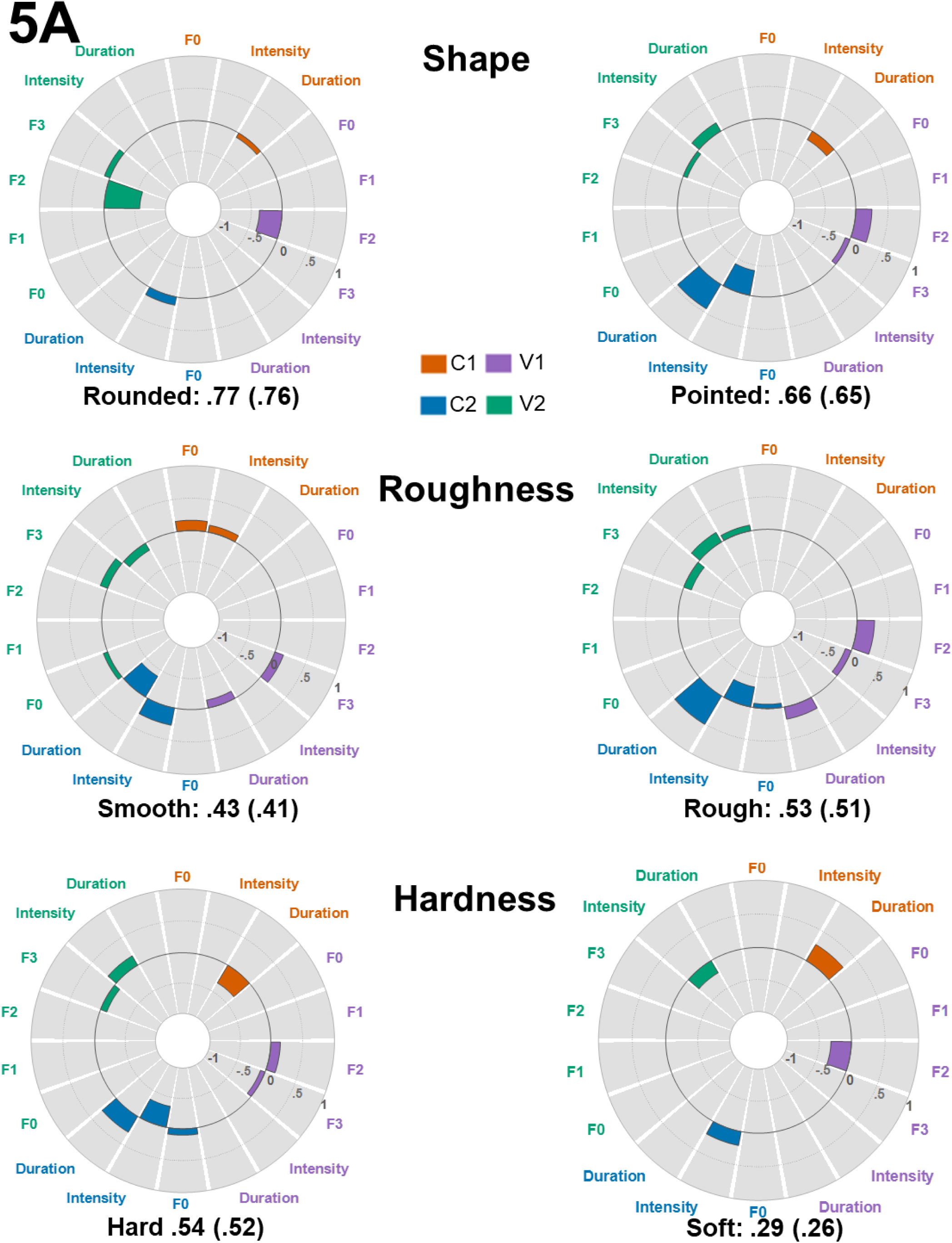
Segmental acoustic regression results for the opposing scales of the shape, roughness, and hardness domains. Interpretation as for Figure 1A. For the complete results, see Supplementary Table 13.

**Figure 5B:**
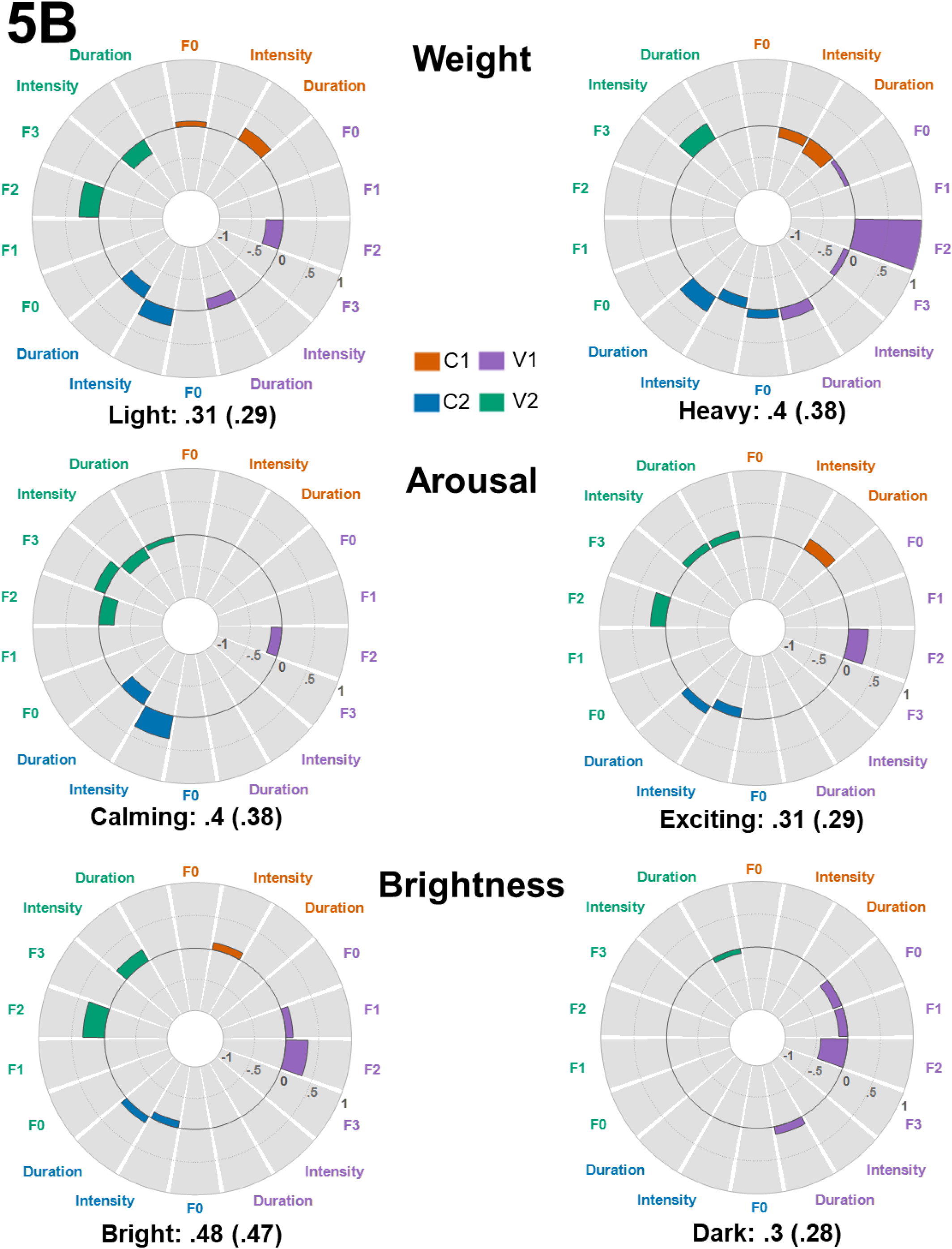
Segmental acoustic regression results for the opposing scales of the weight, arousal, and brightness domains. Interpretation as for Fig. 1A. For the complete results, see Supplementary Table 13.

**Figure 5C:**
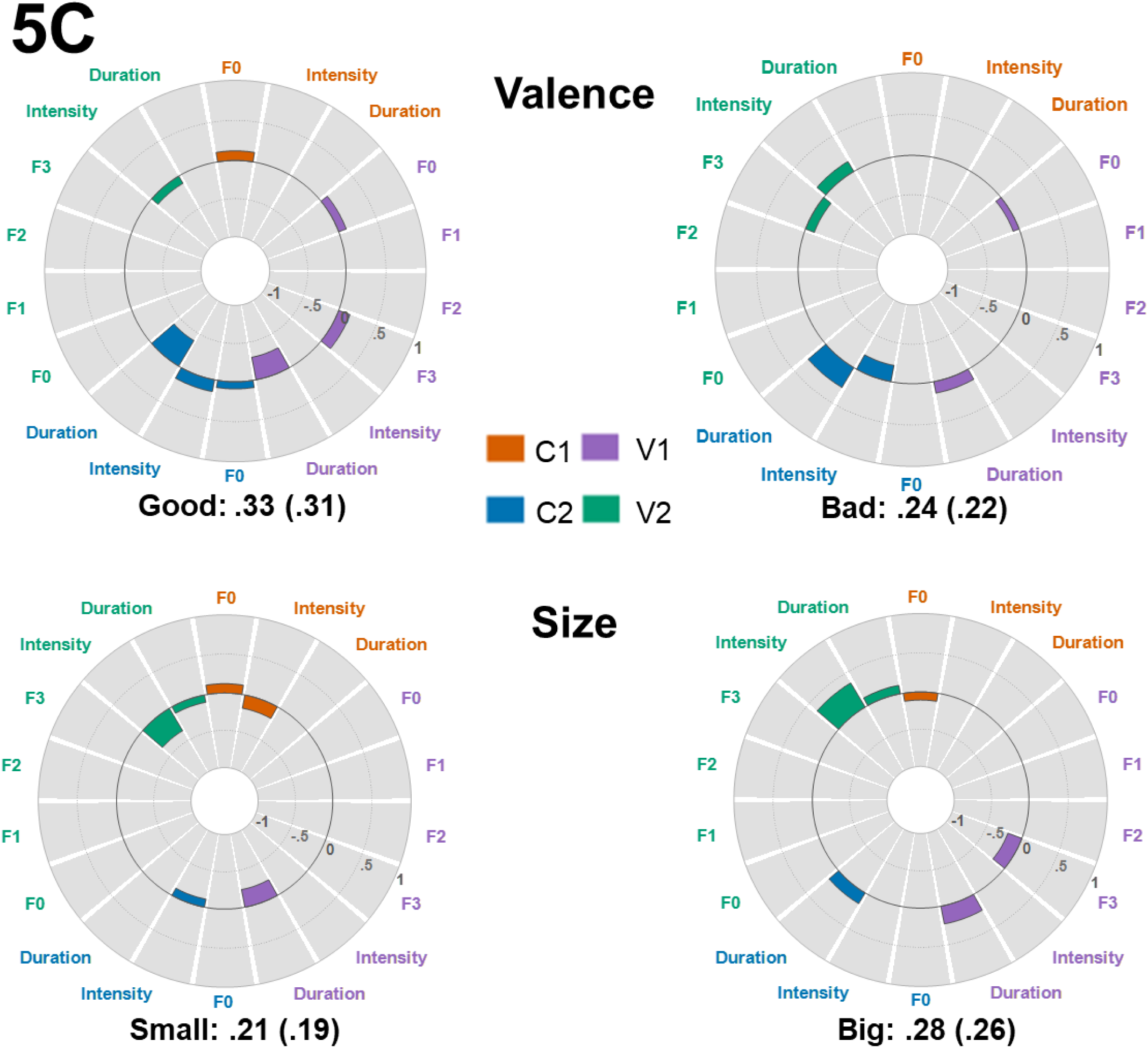
Segmental acoustic regression results for the opposing scales of the valence and size domains. Interpretation as for Fig. 1A. For the complete results, see Supplementary Table 13.

### Results

#### Shape

As for the other types of regression models, the shape domain had the best model fit between ratings and the acoustic parameters associated with particular segments, with R^2^ of .77 for the rounded dimension and .66 for the pointed dimension (Figure 5A; Supplementary Table 13).

In general, although acoustic properties of both the consonants and vowels across positions contributed to shape ratings, properties of the consonants were somewhat more associated with pointed ratings while properties of the vowels contributed more to rounded ratings. In particular, the duration of the initial consonant contributed to both roundedness and pointedness: as the duration increased, roundedness and pointedness ratings both decreased, although to a greater degree for pointed ratings. However, this pattern reversed for the second consonant such that as the duration increased, higher pointedness ratings also increased. This increase may reflect longer closure durations in intervocalic voiceless stops (Davis & Van Summers, 1989) and longer characteristic durations for stops and fricatives in pointed pseudowords relative to durations for sonorants in rounded pseudowords (Umeda, 1977). This juxtaposition of duration effects highlights the complex changes in the realization of acoustic-phonetic forms as a function of phonetic context. Intensity of the second consonant was associated with both pointedness and roundedness. As the intensity increased, rounded ratings increased and pointed ratings decreased, with the latter accounting for a relatively large proportion of the variance in ratings. The presence of lower intensity segments such as unvoiced stops in pointed pseudowords and higher intensity segments such as sonorants in rounded pseudowords would be consistent with this finding (Ohala, 1992).

For the vowels, the second and third formants were the largest contributors to shape ratings, while there was no significant effect of the first formant in either vowel position. In the first vowel, the second formant frequency distinguished between roundedness and pointedness with increases in F2 frequency associated with decreases in rounded ratings and increases in pointed ratings respectively. Lower F2 values are associated with vowel backness, which in turn are associated with roundedness. Conversely, higher F2 values are associated with high, front vowels, which in turn are associated with higher pointed ratings. The contribution of F2 values to roundedness judgments was found for the second vowel as well, with lower F2 values associated with higher roundedness ratings. Contrary to our predictions, lower F3 values were associated with higher pointedness ratings at both V1 and V2, perhaps due to the presence of surrounding velar consonants for pointed pseudowords (Hillenbrand et al., 2001), and higher F3 values were associated with higher roundedness ratings at V2. Intensity of the second vowel, but not the first, was positively associated with pointed ratings. Contrary to our predictions, F0 was not predictive of shape ratings across either vowel or consonant segments.

#### Roughness

Roughness had the next best model fit with R^2^ of .43 for the smooth dimension and .52 for the rough dimension (Figure 5A; Supplementary Table 13).

In general, acoustic properties of both the consonants and vowels contributed to roughness ratings, with the largest contributions for properties of vowels at both vowel positions, and for the second consonant. For the initial consonant, higher F0 was associated with higher smoothness ratings and lower roughness ratings, likely reflecting the interaction between the consonant formant transitions and the following vowel. As predicted, for the second consonant, duration and intensity were contributors to both smooth and rough ratings. As intensity increased, ratings of smoothness increased and ratings of roughness decreased, while the converse occurred for duration: as duration increased, ratings of smoothness decreased and roughness increased. The association between lower intensity and longer duration with higher roughness ratings may be related, as noted above, to the acoustic characteristics of the stops and fricatives (lower intensity and longer duration) associated with rough pseudowords relative to the sonorants associated with smooth pseudowords.

Contributions to roughness ratings varied for the first and second vowels. For the first vowel, F2 contributed to roughness ratings with increases in F2 associated with higher roughness ratings, while higher F3 values were associated with higher smoothness ratings and lower roughness ratings. Longer duration vowels were associated with lower smoothness ratings and higher roughness ratings. Although it is unclear why F3 would be related to roughness judgments in this way, the combination of F2 and duration is consistent with the contribution of the tense, fronted vowel /i/ to higher roughness judgments and fewer lax, back rounded vowels to higher smoothness judgements (see Figure 2A; Supplementary Table 7). For the second vowel, there was no significant contribution of F2 values to roughness judgments; however, the pattern for F3 was similar to contributions for the first vowel. In addition, for the second vowel, higher intensity values contributed to higher roughness ratings and lower smoothness ratings. F0 of the vowels contributed to smoothness, but not roughness, ratings and only for the second vowel. There was no significant effect of F1. Similar effects of duration were found for the second vowel as for the first, with longer duration vowels associated with higher roughness ratings, perhaps reflecting longer duration tense vowel segments in the rough pseudowords. For the vowels, F0 made a contribution to smoothness ratings only at V2, but not to roughness ratings in either vowel position. There was no significant effect of F1 at all.

#### Hardness

Overall, the model fit for hardness was somewhat lower than for roughness with R^2^ of .54 for the hardness dimension and .29 for the softness dimension (Figure 5A; Supplementary Table 13).

Acoustic characteristics of segments at each position contributed to the hardness aspect of texture, with the largest contributions overall for intensity and duration. For the initial consonant, duration was associated with ratings for both the hardness and softness dimensions. As duration increased, ratings of softness increased and ratings of hardness decreased. This is inconsistent with our overall predictions for consonant segment duration but may reflect longer durations for labial consonants for soft pseudowords, particularly in the initial position (Crystal & House, 1988). Intensity and F0 of the initial consonant did not influence either hardness or softness ratings. For the second consonant, duration and intensity were contributors to both hard and soft ratings. As expected, as intensity increased, ratings of softness increased and ratings of hardness decreased. As duration and F0 increased, ratings of hardness increased, but there were no significant effects of these variables on softness ratings. The association between lower intensity and longer duration consonants with higher hardness ratings may be related, as noted above for roughness, to longer durations for the stops and fricatives found in hard pseudowords relative to the voiced segments, particularly sonorants, in soft pseudowords.

Contributions to hardness ratings varied for the first and second vowels. For the first vowel, F2 contributed primarily to softness ratings with increases in F2 associated with lower softness ratings and to some degree higher hardness ratings. Lower F3 values were associated with lower hardness ratings, but F3 did not affect softness ratings. The association with F2 is consistent with the contribution of high, front vowels such as /i/ to higher hardness judgments and low, back rounded vowels to higher softness judgements. F0, F1, intensity, and duration did not contribute to either hardness or softness ratings for the first vowel. For the second vowel, the pattern for F3 was similar to contributions for the first vowel but there was no significant contribution of F0, F1, F2, or duration to listeners’ ratings. Primarily, for the second vowel, there was a strong contribution for intensity with higher intensity values contributing to higher hardness ratings and lower softness ratings. It is unclear what phonemic properties might be related to these intensity differences; these associations may reflect additional stress on the second syllable in pseudowords rated as hard rather than soft (Cutler & Jesse, 2021).

#### Weight

Consistent with patterns observed for the other regression model approaches, model fit values were lower for the weight domain than for those discussed above with R^2^ of .31 for the lightness dimension and .40 for the heaviness dimension (Figure 5B; Supplementary Table 13).

Acoustic properties of both consonants and vowels contributed to weight ratings, but the exact nature of the contribution varied with respect to position, pointing to phonetic context and positional effects in how segments are realized acoustically. In particular, the duration of the initial consonant contributed to both lightness and heaviness ratings: as the duration increased, lightness ratings increased and heaviness ratings decreased. However, this pattern reversed for the second consonant such that as the duration increased, heaviness ratings increased and lightness ratings decreased, perhaps reflecting longer closure durations in intervocalic voiceless stops (Davis & Van Summers, 1989), and the realization of acoustic-phonetic properties associated with other phoneme classes changing as a function of phonetic context. Increases in intensity of the initial consonant were associated with decreases in heaviness ratings, albeit without an effect on lightness ratings. This is consistent with generally lower intensity for obstruents relative to the sonorants, which were associated with lightness. For the second consonant, as intensity increased, lightness ratings increased and heaviness ratings decreased. The presence of lower intensity segments such as stops in pseudowords rated as heavy, and higher intensity segments such as fricatives and sonorants in pseudowords rated as light, would be consistent with this finding. Consonant F0 was not influential except for a small negative effect for the initial consonant on heaviness, but not lightness, ratings.

For the first vowel, F2 frequency contributed to both lightness and heaviness ratings with increases in F2 associated with decreases in lightness ratings and increases in heaviness ratings respectively. Higher F2 values are typically associated with front vowels, which were not found to be strongly predictive of lightness ratings. Lower F2 values are associated with vowel backness and lip rounding, which are in turn associated with heaviness ratings, so it is unclear why we find higher F2 associated with higher heaviness ratings, at least in the first syllable. However, longer duration vowels were associated with higher ratings of heaviness and shorter durations with lower ratings of lightness, which is consistent with the characteristic durations of back and front vowels associated with heaviness and lightness, respectively. F0 and F3 both had small negative association with heaviness ratings, while F1 and intensity had no influence. For the second vowel, the contribution of F2 values to lightness judgments was reversed, with higher F2 values associated with higher lightness ratings, but without affecting heaviness ratings. Intensity of the second vowel was negatively associated with lightness ratings and positively associated with heaviness ratings. The remaining parameters of the second vowel (F0, F1, F3, and duration) did not contribute to ratings. Here, it appears that combination of formant structure and intensity reflect the differing vowel composition of the second syllables (e.g., six vowels in V1 and 2 vowels in V2) and perhaps mechanisms of vowel reduction that impact vowel realization (Ladefoged & Johnson, 2011). For example, the association of higher vowel intensity with higher heaviness ratings and lower lightness ratings could result from less vowel reduction for segments associated with higher heaviness ratings than for lightness ratings in pseudoword-final position (e.g., Moon & Lindblom, 1994).

#### Arousal

For the arousal domain, R^2^ was .40 for the calming dimension and .31 for the exciting dimension (Figure 5B; Supplementary Table 13).

As for the preceding domains, acoustic properties of both the consonants and vowels contributed to arousal ratings. For the initial consonant, duration contributed to excitingness ratings; as duration increased, exciting ratings increased. For the second consonant, longer durations were associated with lower calmingness ratings and again, with higher excitingness ratings. Both findings may be related to the longer durations for the stops and fricatives associated with higher excitingness ratings. Higher intensity ratings were associated with the opposite pattern: higher calmingness, and lower excitingness, ratings. This pattern may again reflect the presence of stops in pseudowords rated as exciting, with lower intensity values relative to the voiced sonorant segments in pseudowords rated as calming.

Contributions to arousal ratings were more prominent for the second than for the first vowel. For both vowels, increases in F2 were associated with higher excitingness ratings as well as with lower calmingness ratings. The contribution of F2 is consistent with the association of high, front vowels such as /i/ with higher exciting ratings and back rounded vowels with calming ratings in other analyses (see Figures 1B and 2B; Supplementary Tables 4 and 7). Higher F3 values of the second vowel were associated with higher calmingness ratings, which is inconsistent with the presence of lip rounding for vowels in pseudowords rated as calming. F3 did not influence excitingness ratings. Higher intensity and duration values contributed to higher excitingness ratings and lower calmingness ratings. As with hardness (see above), it is unclear what vowel properties might be related to these intensity and duration differences; these associations may reflect additional stress on the second syllable in pseudowords rated as exciting rather than calming (Cutler & Jesse, 2021).

#### Brightness

For the brightness domain, R^2^ was .48 for the brightness dimension and .30 for the darkness dimension (Figure 5B; Supplementary Table 13).

The acoustic properties of the vowels were primary contributors to the brightness ratings, whereas few features of the consonants were associated with brightness. For the initial consonant, as intensity increased, brightness ratings increased. For the second consonant, that pattern reversed with higher intensity associated with lower brightness ratings. In addition, longer durations were associated with higher brightness ratings. This pattern may reflect the presence of longer closure durations for voiceless stops in intervocalic positions affecting brightness judgments (Davis & Van Summers, 1989). Note that these effects sizes were small, and darkness ratings were not impacted by these predictors. The remaining consonant predictors did not produce significant effects.

Contributions to brightness ratings were largely consistent for the first and second vowels. For the first vowel, increases in F1 and particularly increases in F2 were associated with higher brightness ratings as well as with lower darkness ratings. Lower F0 was associated with higher darkness ratings, without affecting brightness ratings. These patterns are consistent with lower frequencies overall being associated with darkness as opposed to brightness, and the contribution of F2 specifically is consistent with the association of high, front vowels such as /i/ with higher brightness ratings in other analyses (see Figures 1B and 2B; Supplementary Tables 4 and 7). Longer duration vowels were associated with higher darkness ratings but did not influence brightness ratings, perhaps reflecting the longer duration back vowels. F2 and intensity of the second vowel positively influenced brightness ratings without impacting darkness ratings, likely again reflecting the association of high front vowels and potentially increased stress with brightness ratings, while duration was negatively related to darkness ratings but had no effect on brightness ratings, again potentially an indicator of reduced stress on the second syllable of pseudowords with higher darkness ratings (Cutler & Jesse, 2021). None of the other vowel parameters were influential.

#### Valence

The valence domain again had the second worst model fit with R^2^ of .33 for the goodness dimension and .24 for the badness dimension (Figure 5C; Supplementary Table 13).

The acoustic properties of both consonants and vowels contributed to valence ratings. For the initial consonant, as F0 increased, goodness ratings increased, perhaps reflecting production of initial sonorants with higher F0 and/or coarticulatory effects with the following vowel (Whalen & Levitt, 1995; and see below); however, badness ratings were not affected. For the second consonant, the same pattern was found for F0. In addition, higher intensity values were associated with higher goodness ratings and lower badness ratings and longer durations were associated with lower goodness ratings and higher badness ratings. This pattern may reflect the presence of higher-intensity and shorter-duration sonorants associated with goodness judgements and lower-intensity and longer-duration fricatives associated with badness ratings.

For the initial vowel, as F0 increased, goodness ratings increased and badness ratings decreased, perhaps reflecting the association of high vowels with higher F0 (Whalen & Levitt, 1995). There was also a positive relationship between F3 values and goodness ratings, but no significant association with badness ratings. As lower F3 values can reflect lip rounding, this association likely reflects the association of goodness with unrounded vowels. Longer durations of the first vowel were associated with lower goodness, and higher badness, ratings, again perhaps reflecting the presence of shorter duration front vowels. For the second vowel, higher intensity values were associated with higher badness and lower goodness ratings, while F3 was negatively related to badness ratings, without affecting goodness ratings, reflecting the back rounded vowels associated with higher badness ratings. None of the remaining predictors was significant in the regression.

#### Size

As for the other regression analyses, the model fit for size was the poorest with R^2^ of .21 for the smallness dimension and .28 for the bigness dimension (Figure 5C; Supplementary Table 13).

The acoustic properties of both consonants and vowels contributed to size ratings. For the initial consonant, as F0 increased, small ratings increased and big ratings decreased, possibly reflecting the associations of higher frequencies with small and lower frequencies with large objects (Spence, 2011) and/or the influence of the high front vowels associated with ratings of smallness and the low back vowels associated with ratings of bigness. Higher intensity values were also associated with lower smallness ratings, again perhaps reflecting overall associations between the level of acoustic energy and size and/or the influence of high, front vowels, but bigness ratings were unaffected by this parameter. For the second consonant, the same pattern was found for intensity, and longer duration values were associated with higher bigness ratings, consistent with both the association of stops and fricatives with higher ratings of bigness and general non-linguistic cross-modal associations with size (Birngruber & Ulrich, 2019; Spence, 2011: although these parameters only influenced one of the two opposing scales).

For the initial vowel, there was a contribution of F3 values to bigness ratings with increases in F3 associated with lower bigness ratings; smallness ratings were not impacted. Because back rounded vowels tend to appear in pseudowords rated as big, this association likely reflects the association between F3 and lip-rounding. Longer durations of the first vowel were associated with higher bigness ratings and lower smallness ratings, reflecting well-established associations between size and duration (e.g., Birngruber & Ulrich, 2019) and the characteristic durations of shorter high front vowels (smallness) and longer low back vowels (bigness). For the second vowel, higher intensity and longer duration values were associated with higher bigness and lower smallness ratings. Here, the relationship between intensity and duration with size may reflect general cross-modal associations (Koch et al., 2025), and the role of stress in the production of the second syllable (Cutler & Jesse, 2021). The remaining parameters had no significant effects on ratings.

### Discussion

The associations between the acoustic segmental properties of the pseudowords and ratings across domains were largely consistent with the previous phonetic category, phoneme, and whole item acoustic analyses. The patterns of acoustic parameters across consonants and vowels and across positions were again distinct across domains with collections of properties and segmental positions predicting ratings in a domain-specific way. More specifically, the segmental acoustic analysis, along with the phoneme analysis, allowed us to capture positional effects and this analysis provides information as well about interactions between phoneme properties and details of their acoustic-phonetic realization in context.

#### Fundamental Frequency

Across domains and across segment type (consonant versus vowel) and position, F0 *generally* did not contribute significantly to the ratings of meaning. However, higher F0 values for C1 were associated with higher smoothness (and lower roughness), lightness, goodness and smallness (and lower bigness) ratings. Higher F0 values for V1 were associated with lower heaviness, darkness, and badness (and higher goodness) ratings. Higher F0 values for C2 were associated with lower roughness, higher hardness, and higher heaviness ratings. Higher F0 values for V2 were associated with higher smoothness ratings. These associations are largely consistent with perceived correspondences between F0 and multiple domains of meaning (Sidhu & Pexman, 2018) and likely reflect the acoustic-phonetic properties of the component segments, such as the higher intrinsic F0 of high front versus low back vowels (Shadle, 1985).

#### Intensity and Duration

Intensity and duration values were particularly impactful for consonants in the second position and were also associated with the vowels in both positions of nearly every meaning domain. However, whether intensity or duration was associated with ratings and in what position varied across meaning domains. For C1, longer duration consonant segments were associated with higher ratings of softness (and lower hardness), lightness (and lower heaviness), and excitingness. Higher intensity consonant segments in C1 were associated with higher ratings of smoothness and brightness and lower ratings of heaviness and smallness. For V1, longer duration consonant segments were associated with higher ratings of roughness (and lower smoothness), heaviness (and lower lightness), badness (and lower goodness), and bigness (and lower smallness). There were no associations between intensity of V1 and ratings.

For consonants in second position, the predictive value of intensity and duration parameters were more prevalent and mirrored one another and, with higher intensity and shorter duration segments such as sonorants associated with certain meanings and lower intensity and longer duration segments such as stops and fricatives associated with opposite meanings. For C2, longer duration consonant segments were associated with higher ratings of pointiness, roughness (and lower softness), hardness, heaviness (and lower lightness), excitingness (and lower calmness), brightness, badness (and lower goodness), and bigness. Higher intensity consonant segments in C2 were associated with higher ratings of roundedness (and lower pointedness), smoothness (and lower roughness), softness (and lower hardness), lightness (and lower heaviness), calmingness (and lower excitingness), and goodness (and lower badness), as well as lower brightness and smallness ratings.

There was a larger number of associations between intensity values and meaning ratings for vowels in second position than for vowels in first position. In contrast, there was a smaller number of associations between duration values and meaning ratings for vowels in second position than for vowels in first position, For V2, longer duration segments were associated with higher ratings of roughness, excitingness (and lower calmness), and bigness (and lower smallness) as well as lower darkness ratings. Higher intensity segments in V2 were associated with higher ratings of pointedness, roughness (and lower smoothness), hardness (and lower softness), heaviness (and lower lightness), excitingness (and lower calmness), brightness, badness (and lower goodness), and bigness (and lower smallness). The associations between intensity values and particular meanings (e.g., pointedness) do not appear to reflect vowel quality (higher intensities are associated with low back vowels and lower intensities with high front vowels; Ohala, 1992). Rather, the pattern of intensity to meaning mappings may reflect the acoustic correlates of increased stress (or emphasis) on the second syllable for certain meanings, perhaps consistent with the magnitude relations within a domain (e.g., Cutler & Jesse, 2021; Spence, 2011).

#### Formant frequencies (F1, F2, F3)

The values of the formant frequencies, particularly F2, were consistent with the types of vowels that are typically associated with each meaning domain and that were found to be associated with each meaning in the phonetic category and phoneme analyses. In general, F1 values were not associated with meaning ratings, with the exception of higher and lower F1 values for pseudowords rated as bright and dark, respectively. F2 values, reflecting vowel backness, were associated with meaning ratings in both vowel positions, with stronger and more consistent relationships in V1. Higher F2 values were associated with higher ratings of pointedness (and lower roundedness), roughness, hardness (and lower softness), heaviness (and lower lightness), excitingness (and lower calmness), and brightness (and lower darkness). There were also associations in V2 positions, in which higher F2 values were associated with higher ratings of lightness, excitingness, and brightness and lower ratings of roundedness and calmingness. These associations are consistent with the presence of high front vowels in meanings associated with higher F2 values (e.g., pointedness) and low back vowels in meanings associated with lower F2 values (e.g., calmingness). The more prevalent associations with F2 in the first vowel position may be a consequence of typical vowel reduction in unstressed second syllables, with less distinctiveness of vowel formant structure (Barbes, 2006; Lindblom, 1963). These findings highlight the impact of positional effects with some segments (e.g., V1) contributing more to ratings of iconicity than other segments (e.g., V2) because of coarticulatory and other processes of the acoustic-phonetic realization of phonemes.

F3 was associated with ratings across meaning domains and across positions. For V1 vowels, higher F3 values were associated with higher ratings of smoothness and goodness and lower ratings of pointedness, roughness, hardness, heaviness, and bigness. For V2 vowels, higher F3 values were associated with higher ratings of roundedness, smoothness, and calmingness and lower ratings of pointedness, roughness, hardness, and badness. These findings reveal mappings that were somewhat inconsistent with results of the previous analyses. F3 typically reflects lip rounding with higher F3 indicating less rounding and lower F3 indicating more rounding. Lip rounding in our pseudoword stimuli and in American English is correlated with vowel quality such that high front vowels are unrounded and low back vowels are rounded. The associations between F3 values and ratings of meaning were both consistent (e.g., the presence of unrounded high front vowels in pseudowords rated as good) and inconsistent (e.g., the presence of rounded low back vowels in pseudowords rated as rounded) with the types of vowels we would expect in pseudowords with particular meaning ratings. The contradictory findings may stem in part from coarticulatory processes and their consequences for acoustic-phonetic realization: F3 values for vowels can be influenced by the place of articulation of surrounding consonants (Hillebrand, Clark & Nearey, 2001; Recasens, 1985) and these processes would be reflected in the acoustic instantiation of vowel formant structure, particularly given that mean formant values in this study were averaged across the duration of each vowel segment. These findings highlight the potential role of context effects, both in terms of surrounding segments and in terms of segmental position, in the instantiation of acoustic-phonetic form and its associations with ratings of iconicity.

## GENERAL DISCUSSION

With few exceptions, studies of sound iconicity have concentrated on a single mapping of sound to meaning, predominantly that of the rounded-pointed dimension of shape. Those studies that have addressed multiple domains of meaning have generally involved a trade-off between either a large number of domains but relatively few real/pseudowords (e.g., Miron, 1961; Johansson et al., 2020; Sidhu et al., 2022) or relatively few domains compared to very large sets of either real words (Winter et al., 2017; see also Tzeng et al., 2017) or pseudowords (Westbury et al., 2018). Here, we obtained ratings across eight domains for a large set of 537 pseudowords as a reasonable compromise; these ratings were used as the outcome variables in four separate multiple regression analyses to determine the associations of each domain with phonetic categories, specific phonemes, and selected acoustic characteristics of both the whole pseudoword and the component consonant and vowel segments.

Although the multi-domain studies referred to above were designed to answer particular questions, the approach we took here has some advantages over these earlier studies. Firstly, we collected dimensional ratings of sound-to-meaning correspondence (Miron, 1961; Sidhu et al., 2022) which allow for graded responses across the entire stimulus set, rather than forced choice assessments of meaning (Tzeng et al., 2017) or yes/no category membership decisions (Westbury et al., 2018) which could obscure the granular detail achieved here. Secondly, we extensively sampled the phonological space and ensured that a wide variety of phonetic categories were both adequately and equally represented (see McCormick et al., 2015, for details); for example, affricates were not included in the phonetic inventory of Sidhu et al. (2022). The resulting pseudowords were controlled for the number and structure of syllables, for example they contained either voiced or unvoiced consonants but not both (though this might arguably be a limitation given that such instances occur in real words). In common with Johansson et al. (2020) and Westbury et al. (2018), we modelled both manner and place of articulation; the latter was not accounted for by Sidhu et al. (2022). Finally, rating the same pseudowords for multiple domains enables meaningful cross-domain comparisons, as opposed to comparing different domains across different studies that might have different phonetic inventories, stimulus set size, measurements, etc. This approach to both ratings and phonological space allowed us to directly assess the relative contribution of particular phonetic categories, specific phonemes, and both whole-item and segmental acoustic characteristics, across multiple domains of meaning.

Since the same pseudowords were rated across eight different domains, we examined the ten highest-rated pseudowords for each of the two dimensions for each domain (since both scales were used in all analyses) to see how often pseudowords repeated across domains. It was clear that these highly-rated pseudowords fell into two distinct groups that shared phonetic categories. The ten highest-rated pseudowords for the rounded, smooth, soft, heavy, calming, dark, good, and big dimensions were largely composed of sonorants, voiced stops and af/fricatives, and back rounded vowels (Table 3A). For the pointed, rough, hard, light, exciting, bright, bad, and small dimensions, however, the highest-rated pseudowords were largely composed of unvoiced stops and af/fricatives, and front unrounded vowels (Table 3B). Despite the fact that the pseudowords in each group were very similar phonetically across domains, over 60% of the pseudowords in each group did not repeat across domains.

### Domain-general vs domain-specific accounts of sound iconicity

The ratings and the four regression analyses argue against domain-general accounts of sound iconicity based on, for example, mappings to higher-order or abstract dimensions such as arousal (Aryani et al., 2020) or valence (for example, Osgood, 1969; but see Tzeng et al., 2017, for commentary). The claim that sound and meaning are connected via arousal (Aryani et al., 2020) is undermined by the fact that only the shape domain was tested in this study when, if true, the effect should generalize to other domains. The present results allow a test of this because a strong prediction of this account would be that arousal ratings for the pseudowords would be related to their ratings for any other domain. In other words, the opposing dimensions of every domain would be assigned to either high or low arousal – for example, brightness and darkness, respectively – and we would expect that high/low arousal ratings would be significantly positively correlated with one and negatively with the other. For the shape, roughness, hardness, and brightness domains, this was, in fact, the case (Table 2), but it was not so for the other domains. Thus, the results suggest that arousal is unlikely to be a domain-general factor underlying all sound-iconicity mappings. Similarly, valence ratings showed fully opposing correlations for the roughness, hardness, weight, and brightness domains but none of the others (Table 2). The lack of evidence for a domain-general account based on the ratings data is consistent with Tzeng et al. (2017) who analyzed their ratings data for semantic relatedness to a range of possible higher-order factors but found no convincing evidence for a domain-general factor underlying the sound-iconic mappings tested.

Domain-general accounts are also contradicted by the finding that different domains were underpinned by different patterns of phonetic categories (Table 4). Even where there were cross-domain correlations for ratings, the phonetic patterns were not necessarily related: some correlated domains shared similar phonetic categories (sometimes very similar), but many did not. For an arousal account in particular, ratings for exciting (high arousal) pseudowords were significantly associated with obstruents which was also true for ratings on other high-arousal scales – for example, pointed, hard, and heavy – but not true for the high-arousal bright scale. In the same way, exciting ratings were significantly associated with unvoiced consonants, as were pointed and bright ratings, but ratings for the high-arousal dimension of heaviness were associated with voiced consonants. Similarly, the patterns of specific phonemes, and both whole-item and segmental acoustic parameters, were highly domain-specific (Tables 5–7).

### Limitations

A general caveat is that the pseudowords were constructed using phonemes that had established associations to the rounded-pointed dimension of shape (McCormick et al., 2015) and thus might be said to be optimized for the shape domain in preference to the others tested here. Against that, as the survey of prior literature shows (see Introduction), none of these phonemes are *exclusively* associated with shape but, instead, have connections to several other domains. Nonetheless, the pseudowords only employ a subset of all English phonemes. In particular, /a/, /ӕ/ and /ʌ/, which have established associations to the size domain (reviewed in Ekström, 2022), are absent, which may help explain the poor model fit for size (Table 4). In other words, it may be that the pseudowords are not so much optimal for shape but rather sub-optimal for size. In fact, Ekström (2022) suggests that there is no settled account of size iconicity. In any case, a more complete account of the phonetic bases for sound iconicity in different domains may emerge from either more targeted sets of phonemes or, given that phonemic associations are likely not exclusive, from a pseudoword set that samples the complete phonetic space with each phoneme equally represented (or in proportion to their frequency in the relevant language).

Another general limitation is that the pseudowords were recorded from a single speaker and that the present findings might not generalize to other speakers, i.e. a ‘speaker-as-fixed-effect’ fallacy (see Winter & Grice, 2021, and Clark, 1973). Not only do listeners draw different inferences from different speakers (see Creel & Bregman, 2011, for a review), but they are also sensitive to the distinctive features of a particular speaker’s voice (Nygaard & Pisoni, 1998; Nygaard et al., 1994). Certainly, future work could address the effect of employing multiple speakers in the same stimulus set, but this also raises the question whether such distinctive features could influence perception of sound iconicity. For example, recent evidence suggests that, for American English speakers, regional accents that produce differences in vowel fronting also produce differences in iconic size judgments, at least for pseudowords (Hoffmann, 2025). Voice quality might also affect iconicity judgments; for example, listening to a speaker whose voice is naturally ‘gravelly’ i.e., tends to pulse phonation, might produce more pronounced perception of iconic pointedness. A recent study of Japanese ideophones suggests that, for example, fast motion was best conveyed by a falsetto voice (Akita & Kawahara, 2025); no effects were found for ‘creaky voice’, i.e., pulse phonation, but perhaps this was because the study concentrated on motion-related ideophones and did not address iconic texture words.

We excluded a relatively large number of participants (almost 20% of those who completed the study) because they were bilingual or spoke a second language to varying degrees of proficiency. The reason for doing this was to reduce the possibility that any pseudoword was actually a real word in another language that was not familiar to the author group. Whilst this reduced the possibility that semantic knowledge from another language influenced ratings, it did have the effect of rendering the sample more linguistically homogenous than the society from which they were drawn and thus may limit the generalizability of the results (see Yarkoni, 2022).

Relatedly, an important limitation is that the pseudowords were constructed according to the phonology and phonotactics of English and thus might not generalize beyond English speakers. Although many of the phonemes in the current study undoubtedly occur in many languages other than English, iconic mappings can be language-specific as well as shared (Saji et al., 2019). Certainly, kiki/bubu’ (Styles & Gawne, 2017) and ‘maluma/takete’ (Rogers & Ross, 1975) sound-to-shape mappings have not been replicated in speakers of some other languages, likely because the test items were not phonetically or phonotactically ‘legal’ in the target language (Styles & Gawne, 2017).

One limitation that might be raised for the shape domain in particular is that the speaker who recorded the pseudowords might have unconsciously pronounced them differently according to their expectations about the pseudowords’ roundedness/pointedness (see also Lacey et al., 2020). This is unlikely because the speaker was instructed to employ a neutral intonation and two independent judges rejected items that did not sound both neutral and consistent with the other recordings. Furthermore, it would be difficult to manipulate complex voice parameters like jitter or shimmer in such a way as to produce systematic effects, particularly over more than 500 items when these were recorded in random order. For the remaining seven domains reported here, the speaker did not know that their recordings would be used for other domains in the future; therefore, they could not have been affected by prior knowledge or expectations about those domains.

Another limitation specific to a domain is that the model fits for the size domain were generally not as good as the other domains and this may be because the pseudowords were optimized for the shape, rather than the size, domain as noted above. This did not seem to prevent participants making reasonably organized ratings and certainly non-optimization was also true of all the other non-shape domains in which meaningful relationships were found. However, a different set of pseudowords, using the same phonemes as here but with the addition of the vowels /ɑ/, /ʌ/, and /ӕ/, that have sound-iconic associations to size (reviewed in Ekstrom, 2022), does show a better model fit for phonetic categories than the current results, together with significant relationships for some voice quality parameters (Nayak, 2024; Nayak et al., 2023, 2024), thus the size domain requires further examination.

In terms of the trade-off between the number of domains and pseudowords, studies using a small pseudoword set may be at a disadvantage: the smaller the pseudoword set, the more items repeat across domains because participants have a restricted choice. For the set of 537 pseudowords used here, we observed more than 60% unique items in the 10 highest-rated items for both dimensions in 8 domains. For a smaller set of 40 pseudowords, Sidhu et al. (2022, Figure 4) provide information for 4 of the 24 domains: of the 8 highest-rated items for the opposing dimensions of each domain, less than 25% are unique (and even in the 4 highest-rated items, unique items still constitute less than 50%). By contrast, for a large set of 7996 pseudowords reported in Westbury et al., (2018: Tables 4, 7, 9), the 10 items most likely to be assigned to 3 domains were 100% unique. (There may be many reasons for this last result aside from the very large set size: for example, pseudowords could consist of more than two syllables and were not restricted to a CVCV sequence since they could start with a vowel and end with a consonant, and could also include biphones like /br/ and /mb/.)

Finally, much may depend on the definition of the domain in terms of aspect and the related dimensions. Even shape, for which the rounded/pointed dimensions might seem foundational, could also be characterized as symmetric/asymmetric. An interesting question here is whether (a)symmetry is iconically represented at the whole-item level, rather than single phonemes; for example, /keke/ might be considered symmetric because the two syllables are the same, /kepe/ less so because only the vowels are the same, and /vʊdʒo/ as asymmetric because the two syllables are completely different. As noted above, we might have found different results if arousal had been defined as restful vs stressful rather than calming vs exciting, with potentially a greater role for jitter, shimmer, and variability in pitch (Van Puyvelde et al., 2018; Kliper et al., 2016). Similarly, good vs bad is a rather ‘all-purpose’ conceptualization of valence and doesn’t necessarily have a more specific moral dimension, e.g. we could have tested virtuous/wicked or honest/dishonest (or even rude/polite given the iconic associations recently reported for profanity [Lev-Ari et al., 2023]); we could also have tested emotional valence using a sad/happy scale (see Warriner et al., 2013). These different aspects and dimensions of valence may have different iconicity profiles. For example, motivationally positive emotions are vocalized with both higher pitch and greater amplitude than motivationally negative emotions, while esthetically positive emotions were vocalized with both *lower* pitch and amplitude than their negative counterparts (Belyk & Brown, 2014). Furthermore, morally positive emotions are vocalized with higher pitch but lower amplitude than morally negative emotions (Belyk & Brown, 2014). These examples show that any domain may comprise multiple related aspects and dimensions with different acoustic profiles. Furthermore, domains may not be orthogonal to each other: as noted above, although the calming and exciting dimensions differed in the level of arousal, they were potentially confounded with valence to some extent because they were both positively valenced. These issues are worth exploring in relation to how different speech sounds differentiate between related concepts.

## CONCLUSIONS

The present findings suggest that the contributions and relative weightings of different phonetic and acoustic features are largely specific to each domain and thus argue against a domain-general account in which sound-meaning mappings are driven by a single common factor. The results also suggest that some iconic mappings may be related to each other in ways that reflect relationships that are either literal (i.e., physical and perceptual) or metaphorical. This might be a more parsimonious account than grouping domains together under over-arching factors such as activity or potency – which are essentially themselves ‘super’-domains – and would reflect two basic modes of expression: literal and figurative language. Further work is needed to explore this more fully.

## ACKNOWLEDGMENTS

This work was supported by grants to KS and LCN from the National Eye Institute at the NIH (R01EY025978) and the Emory University Research Council, and by institutional funds provided to KS by Penn State College of Medicine. Earlier versions of these data were presented at the 2022 meeting of the Cognitive Neuroscience Society (Lacey et al., 2022) and were available as preprints at *bioRxiv*, doi: 10.1101/2024.09.03.610970 (phonetic analyses) and doi: 10.1101/2024.09.03.610973 (acoustic analyses). We thank Ana Maria Hoffmann for assistance with data collation, and Saachi Nayak for the use of data on the rough-smooth dimension of texture from her MS thesis under the direction of KS and SL (Nayak, 2024). We also thank Nancy Bliwise and Yuk Fai Cheong, Department of Psychology, Emory University, for statistical advice.

## Author contributions

SL, KS and LCN designed the study; KLM collected the data; KLM, AS and SL analyzed the data; and SL, KLM, AS, KS and LCN wrote the paper.

## DATA AVAILABILITY

The set of 537 pseudowords is available at https://osf.io/ekpgh/ and the rating data for all domains are available at https://osf.io/y9zjc/.

## Footnotes

1 Sound iconicity in spoken language is often referred to as ‘sound symbolism’ but, while this term has a long history, and is in frequent use, it has been criticized as being out of step with current thinking in linguistics, semiotics, and related disciplines (as well as excluding non-spoken languages). Essentially, the objection is that these disciplines use ‘iconic’ to refer to a relationship between form and meaning based on resemblance, but ‘symbolic’ to refer to relationships based on convention (see Ćwiek et al., 2025; also Ahlner & Zlatev, 2010, and Winter et al., in press, among others). Thus, we use the terms ‘iconicity’ or ‘sound iconicity’ to refer to instances in which the sounds in an utterance resemble its meaning.

2 Only three of these were constructed, as here, from two opposing dimensions: shape: round/sharp, size: large/small, and gender: masculine/feminine. The remaining two, concrete/abstract and high/low valence, were constructed by concatenating multiple categories which either did not have a formal opposite dimension, e.g. fungus, or omitted it, e.g., sadness was included but not happiness.

3 Broadly speaking, vowels can be identified by the relative frequencies of their formants – the resonance frequencies of the vocal tract when producing the vowel sound – and to a lesser extent by their characteristic fundamental frequency (F0). The first three formants, F1-F3, are the most informative about vowel identity, while higher formants are informative about speaker identity (Knoeferle et al., 2017).

4 As will become clear, some meaning domains can be defined in different ways, and we distinguish between domain (the ‘headline’ concept, e.g., shape), aspect (a specific attribute, e.g., angularity), and dimensions (the component, usually opposing, parts of the attribute, e.g., rounded vs pointed). Another aspect of the shape domain could be symmetry, the dimensions of which would be symmetric vs asymmetric.

5 Sharpness refers to the high frequency content of a sound and can be thought of as the “bass/treble ratio” (Villegas et al., 2023, p2); sharper sounds have a greater proportion of high frequencies (Fastl & Zwicker, 2007).

6 Note that shimmer and jitter shimmer only apply to voiced segments and that the Praat algorithm identifies these before making these measurements.

## REFERENCES

Adelman, J.S., Estes, Z. & Cossu, M. (2018). Emotional sound symbolism: languages rapidly signal valence via phonemes. Cognition, 175:122–130.

Ahlner, F., & Zlatev, J. (2010). Cross-modal iconicity: A cognitive semiotic approach to sound symbolism. Sign Systems Studies, 38:298–348.

Aiken, S. J., & Picton, T. W. (2008). Human cortical responses to the speech envelope. Ear & Hearing, 29:139–157.

Akita, K. (2021). Phonation types matter in sound symbolism. Cognitive Science, 45:e12982.

Akita, K. (2025). Voice quality has robust visual associations in English and Japanese speakers. Open Linguistics, 11:20250069.

Akita, K. & Kawahara, S. (2025). Iconic prosody enhances the depictive power of ideophones. Language & Cognition, 17:e77.

Anselme, R., Pellegrino, F. & Dediu, D. (2025). Not just the alveolar trill, but “r-like” sounds are associated with roughness across languages, pointing to a more general link between sound and touch. Scientific Reports, 15:12930, doi:10.1038/s41598-025-94850-0

Anwyl-Irvine, A. L., Massonnié, J., Flitton, A., Kirkham, N., & Evershed, J. K. (2020). Gorilla in our midst: An online behavioral experiment builder. Behavior Research Methods, 52:388–407.

Aryani, A., Conrad, M., Schmidtke, D. & Jacobs, A. (2018). Why ‘piss’ is ruder than ‘pee’? the role of sound in affective meaning. PLoS ONE, 13:e0198430, doi: 10.1371/journal.pone.0198430

Aryani, A., Isbilen, E.S & Christiansen, M.H. (2020). Affective arousal links sound to meaning.Psychological Science, 31:978–986.

Auracher, J. (2017). Sound iconicity of abstract concepts: Place of articulation is implicitly associated with abstract concepts of size and social dominance. PLoS ONE, 12:e0187196, doi: 10.1371/journal.pone.0187196

Barnes, J. (2006). Strength and weakness at the interface: positional neutralization in phonetics and phonology. (Phonology and Phonetics 10.) Berlin & New York: Mouton de Gruyter.

Belyk, M. & Brown, S. (2014). The acoustic correlates of valence depend on emotion family. Journal of Voice, 28:523.e9–523.e18.

Bender, E.M. & Friedman, B. (2018). Data statements for natural language processing: toward mitigating system bias and enabling better science. Transactions of the Association for Computational Linguistics, 6:587–604.

Birngruber, T. & Ulrich, R. (2019). Perceived duration increases not only with physical, but also with implicit size. *Journal of Experimental Psychology: Learning*, Memory, and Cognition, 45:969–979.

Blasi, D. E., Wichmann, S., Hammarström, H., Stadler, P. F., & Christiansen, M. H. (2016). Sound–meaning association biases evidenced across thousands of languages. Proceedings of the National Academy of Sciences, 113, 10818–10823.

Boersma, P. & Weenink, D. (2012). Praat: doing phonetics by computer. Accessed at http://www.praat.org/.

Bonett, D.G. (2002). Sample size requirements for testing and estimating coefficient alpha. Journal of Educational & Behavioral Statistics, 27:335–340.

Bowerman, B.L. & O’Connell, R.T. (1990). Linear Statistical Models: An Applied Approach, 2^nd^ edn. Duxbury Press, Belmont CA.

Brockmann, M., Drinnan, M.J., Storck, C. & Carding, P.N. (2011). Reliable jitter and shimmer measurements in voice clinics: the relevance of vowel, gender, vocal intensity, and fundamental frequency effects in a typical clinical task. Journal of Voice, 25:44–53.

Brockmann-Bauser, M., Bohlender, J.E. & Mehta, D.D. (2018). Acoustic perturbation measures improve with increasing vocal intensity in individuals with and without voice disorders. Journal of Voice, 32:162–168.

Catricalà, M. & Guidi, A. (2015). Onomatopoeias: a new perspective around space, image schemas and phoneme clusters. Cognitive Processing, 16, Supplement 1:S175–S178.

Cauldwell, R. Hewings, M. (1996). Intonation rules in ELT textbooks. ELT Journal, 50:327–334.

Cheng, L. S. P., Burgess, D., Vernooij, N., Solis-Barroso, C., McDermott, A., & Namboodiripad, S. (2021). The problematic concept of native speaker in psycholinguistics: replacing vague and harmful terminology with inclusive and accurate measures. Frontiers in Psychology, 12:715843, doi: 10.3389/fpsyg.2021.715843

Chodroff, E. & Wilson, C. (2014). Burst spectrum as a cue for the stop voicing contrast in American English. Journal of the Acoustical Society of America, 136:2762–2772.

Clark, H.H. (1973). The language-as-fixed-effect fallacy: A critique of language statistics in psychological research. Journal of Verbal Learning and Verbal Behavior, 12:335–359.

Connell, L., & Lynott, D. (2012). Strength of perceptual experience predicts word processing performance better than concreteness or imageability. Cognition, 125:452–465.

Creel, S. C., & Bregman, M. R. (2011). How talker identity relates to language processing. Language and Linguistics Compass, 5:190–204.

Crystal, T.H., & House, A.S. (1982). Segmental durations in connected speech signals: preliminary results. Journal of the Acoustical Society of America, 72:705–716.

Crystal, T.H., & House, A.S. (1988). The duration of American-English stop consonants: an overview. Journal of Phonetics, 16:285–294.

Cuskley, C., Simner, J., & Kirby, S. (2017). Phonological and orthographic influences in the bouba–kiki effect. Psychological Research, 81, 119–130.

Cutler, A., & Jesse, A. (2021). Word stress in speech perception. In J. S. Pardo, L. C. Nygaard, R. E. Remez, & D. B. Pisoni (Eds.), The Handbook of Speech Perception, (2nd ed.), pp. 239–265. New York: Wiley.

Ćwiek, A., Anselme, R., Dediu, D., Fuchs, S., Kawahara, S., et al. (2024). The alveolar trill is perceived as jagged/rough by speakers of different languages. Journal of the Acoustical Society of America, 156:3468–3479.

Ćwiek, A., Kreiman, J. & Fuchs, S. (2025). Introduction to the special issue on iconicity and sound symbolism. Journal of the Acoustical Society of America, 157:3806–3813.

Davis, S. & Van Summers, W. (1989). Vowel length and closure duration in word-medial VC sequences. Journal of Phonetics, 17:339–353.

de Saussure, F. (1916/2009). Course in General Linguistics. Open Court Classics: Peru, IL, USA.

D’Onofrio, A. (2014). Phonetic detail and dimensionality in sound-shape correspondences: Refining the bouba-kiki paradigm. Language and Speech, 57:367–393.

Ekström, A.G. (2022). What’s next for size-sound symbolism? Frontiers in Language Sciences, 1:1046637, doi: 10.3389/flang.2022.1046637

Emmorey, K. (2023). Ten things you should know about sign languages. Current Directions in Psychological Science, 32:387–394.

Ferrand, C. T. (2002). Harmonics-to-noise ratio: An index of vocal aging. Journal of Voice, 16:480–487.

Field, A. (2018). Discovering Statistics Using IBM SPSS Statistics, 5^th^ edition. Sage Publications: Thousand Oaks, CA USA.

Fort, M., Martin, A. & Peperkamp, S. (2015). Consonants are more important than vowels in the Bouba-Kiki effect. Language & Speech, 58, 247–266.

Fort, M. & Schwartz, J.-L. (2022). Resolving the bouba-kiki effect enigma by rooting iconic sound symbolism in physical properties of round and spiky objects. Scientific Reports, 12:19172.

Greenberg, J.H. & Jenkins, J.J. (1966). Studies in the psychological correlates of the sound system of American English. Word, 22:207–242.

Guzman-Martinez, E., Ortega, L., Grabowecky, M., Mossbridge, J. & Suzuki, S. (2012). Interactive coding of visual spatial frequency and auditory amplitude-modulation rate. Current Biology, 22:383–388.

Hair, J.F., Hult, G.T.M., Ringle, C.M. & Sarstedt, M. (2022). A Primer on Partial Least Squares Structural Equation Modeling (PLS-SEM), 3^rd^ edition. Sage: Thousand Oaks, CA.

Hillenbrand, J.M., Clark, M.J., & Nearey, T.M. (2001). Effects of consonant environment on vowel formant patterns. Journal of the Acoustical Society of America, 109:748–763.

Hillenbrand, J., Getty, L. A., Clark, M. J., & Wheeler, K. (1995). Acoustic characteristics of American English vowels. Journal of the Acoustical Society of America, 97:3099–3111.

Hirata, S., Ukita, J. & Kita, S. (2011). Implicit phonetic symbolism in voicing of consonants and visual lightness using Garner’s speeded classification task. Perceptual & Motor Skills, 113:929–940.

Hockett, C. (1959). Animal “languages” and human language. Human Biology, 31:32–39.

Hoffmann, A.M. (2025). Bouba, kiki, and beyond: exploring the flexibility of sound symbolism across language users. Unpublished doctoral dissertation, Emory University, Atlanta GA, USA.

Hollien, H., Girard, G. T., & Coleman, R. F. (1977). Vocal fold vibratory patterns of pulse register phonation. Folia Phoniatrica et Logopaedica, 29:200–205.

Hollins, M., Bensmaia, S., Karlof, K. & Young, F. (2000). Individual differences in perceptual space for tactile textures: evidence from multidimensional scaling. Perception & Psychophysics, 62:1534–1544.

Hornibrook, J., Ormond, T. & Maclagan, M. (2018). Creaky voice or extreme vocal fry in young women. New Zealand Medical Journal, 131:36–40.

Ishi, C.T., Sakakibara, K.-I., Ishiguro, H. & Hagita, N. (2008). A method for automatic detection of vocal fry. *IEEE Transactions on Audio, Speech*, & Language Processing, 16:47–56.

Johansson, N.E., Anikin, A., Carling, G. & Holmer, A. (2020). The typology of sound symbolism: Defining macro-concepts via their semantic and phonetic features. Linguistic Typology, 24:253–310.

Jongman, A., Wayland, R. & Wong, S. (2000). Acoustic characteristics of English fricatives. Journal of the Acoustical Society of America, 108:1252–1263.

Klatzky, R.L., Lederman, S. & Reed, C. (1987). There’s more to touch than meets the eye: the salience of object attributes for haptics with and without vision. Journal of Experimental Psychology: General, 116:356–369.

Klink, R.R. & Wu, L. (2014). The role of position, type, and combination of sound symbolism imbeds in brand names. Marketing Letters, 25:13–24.

Kliper, R., Portuguese, S., & Weinshall, D. (2016). Prosodic analysis of speech and the underlying mental state. In S. Serino et al., (eds.) Pervasive Computing Paradigms for Mental Health: MindCare 2015 Selected Papers, pp52–62. Springer, Switzerland.

Kluender, K.R., Diehl, R. L. & Wright, B.A. (1988). Vowel-length differences before voiced and voiceless consonants: an auditory explanation. Journal of Phonetics, 16:153–169.

Knoeferle, K., Li, J., Maggioni, E., & Spence, C. (2017). What drives sound symbolism? Different acoustic cues underlie sound-size and sound-shape mappings. Scientific Reports, 7, 5562. doi: 10.1038/s41598-017-05965-y.

Koch, S., Schubert, T. & Blankenberger, S. (2025). Large sounds and loud numbers? Investigating the bidirectionality and automaticity of cross-modal loudness-number interactions. Quarterly Journal of Experimental Psychology, 78:2741–2757.

Köhler, W. (1929). Gestalt Psychology. New York: Liveright Publishing Corporation.

Köhler, W. (1947). Gestalt Psychology: An Introduction to New Concepts in Modern Psychology. Liveright: New York, NY.

Körner, A. & Rummer, R. (2022). Articulation contributes to valence sound symbolism. Journal of Experimental Psychology: General, 151:1107–1114.

Kumar, G.V., Lacey, S., Nygaard, L.C. & Sathian, K. (2025). Acoustic parameter combinations underlying mapping of pseudoword sounds to multiple domains of meaning: representational similarity analyses and machine-learning models. Journal of the Acoustical Society of America, 158:4243–4267.

Lacey, S., Jamal, Y., List, S.M., McCormick, K., Sathian, K. & Nygaard, L.C. (2020). Stimulus parameters underlying sound-symbolic mapping of auditory pseudowords to visual shapes. Cognitive Science, 44:e12883.

Lacey, S., Martinez, M., McCormick, K. & Sathian, K. (2016). Synesthesia strengthens sound-symbolic cross-modal correspondences. European Journal of Neuroscience, 44:2716–2721.

Lacey, S., Matthews, K.L., Hoffmann, A.M., Sathian, K. & Nygaard, L.C. (2022). Perceptual and metaphorical clustering in sound-symbolic ratings of pseudowords. Poster, Cognitive Neuroscience Society, San Francisco, April 23–26, 2022.

Lacey, S., Matthews, K., Hoffmann, A.M., Sathian, K. & Nygaard, L.C. (2024). Differential contributions of the spectro-temporal and vocal characteristics of auditory pseudowords to multiple domains of meaning. Preprint, bioRxiv, BIORXIV/2024/610973.

Ladefoged, P. & Johnson, K. (2011). A Course in Phonetics, 6^th^ edition. Wadsworth: Boston MA.

Leung, K. K., Jongman, A., Wang, M. D., & Sereno, J. A. (2016). Acoustic characteristics of clearly spoken English tense and lax vowels. Journal of Phonetics, 57: 1–13.

Lev-Ari, S. & McKay, R. (2023). The sound of swearing: Are there universal patterns in profanity? Psychonomic Bulletin & Review, 30:1103–1114.

Lindblom, B. (1963). Spectrographic study of vowel reduction. Journal of the Acoustical Society of America, 35:1773–1781.

Ma, W., & Thompson, W.F. (2015). Human emotions track changes in the acoustic environment. Proceedings of the National Academy of Sciences, 112:14563–14568.

Marks, L.E. (1987). On cross-modal similarity: auditory-visual interactions in speeded discrimination. Journal of Experimental Psychology: Human Perception & Performance, 13:384–394.

Maurer, D., Pathman, T., & Mondloch, C. J. (2006). The shape of boubas: Sound-shape correspondences in toddlers and adults. Developmental Science, 9:316–322.

McCormick, K., Kim, J. Y., List, S., & Nygaard, L. C. (2015). Sound to meaning mappings in the bouba-kiki effect. In D. C. Noelle, R. Dale, A. S. Warlaumont, J. Yoshimi, T. Matlock, C. D. Jennings, & P. P. Maglio (Eds.), Proceedings 37th Annual Meeting Cognitive Science Society (pp. 1565–1570). Austin TX, USA: Cognitive Science Society.

Menard, S. (1995). Applied Logistic Regression Analysis. Sage University paper Series on Quantitative Applications in the Social Sciences, 07-106. Sage, Thousand Oaks CA.

Miller J. L. & Volaitis L. E. (1989). Effect of speaking rate on the perceptual structure of a phonetic category. Perception & Psychophysics, 46:505–512.

Miron, M.S. (1961). A cross-linguistic investigation of phonetic symbolism. Journal of Abnormal & Social Psychology, 62:623–630.

Monaghan, P., & Fletcher, M. (2019). Do sound symbolism effects for written words relate to individual phonemes or to phoneme features? Language and Cognition, 11(2), 235–255.

Moon, S-J., & Lindblom, B. (1994). Interaction between duration, context, and speaking style in English stressed vowels. Journal of the Acoustical Society of America, 96:40–55.

Mooshammer, C., Bobeck, D., Hornecker, H., Meinhardt, K., Olina, O. et al. (2023). Does Orkish sound evil? Perception of fantasy languages and their phonetic and phonological characteristics. Language & Speech, 67:961–1000.

Motamedi, Y., Little, H., Nielsen, A. & Sulik, J. (2019). The iconicity toolbox: empirical approaches to measuring iconicity. Language & Cognition, 11:188–207.

Munson, B. & Solomon, N.P. (2004). The effect of phonological neighborhood density on vowel articulation. *Journal of Speech, Language*, & Hearing Research, 47:1048–1058.

Myers, R. (1990). Classical and Modern Regression with Applications, 2^nd^ edn. Duxbury Press, Boston MA.

Nayak, S. (2024). Phonetic and acoustic analyses of sound-symbolic associations of a large pseudoword set across multiple domains of meaning. Unpublished master’s thesis, Pennsylvania State University, College of Medicine, Hershey, PA, USA.

Nayak, S., Lacey, S., Nygaard, L. & Sathian, K. (2023). Differing contributions of acoustic parameters to sound-symbolic associations for shape- and size-optimized pseudoword sets. Poster, Society for Neuroscience, Washington, DC, October 11–15, 2023.

Nayak, S., Lacey, S., Nygaard, L.C. & Sathian, K. (2024). Varying sound-symbolic contributions of acoustic parameters in shape- and size-optimized pseudoword sets across multiple domains of meaning. Poster, Cognitive Neuroscience Society, Toronto, April 13–16, 2024.

Newman, S.S. (1933). Further experiments in phonetic symbolism. American Journal of Psychology, 45:53–75.

Nielsen, A., & Rendall, D. (2011). The sound of round: Evaluating the sound-symbolic role of consonants in the classic Takete-Maluma phenomenon. Canadian Journal of Experimental Psychology, 65, 115–124.

Nuckolls, J.B. (1999). The case for sound symbolism. Annual Review of Anthropology, 28:225–252.

Nygaard, L.C. & Pisoni, D.B. (1998). Talker-specific learning in speech perception. Perception & Psychophysics, 60:355–376.

Nygaard, L.C., Sommers, M.S. & Pisoni, D.B. (1994). Speech perception as a talker-contingent process. Psychological Science, 5:42–46.

Ohala, J. J. (1992). Alternatives to the Sonority Hierarchy for Explaining Segmental Sequential Constraints. In Papers from the Parasession on the Syllable (pp. 319–338). Chicago: Chicago Linguistic Society.

Osgood, C.E. (1969). On the whys and wherefores of E, P, and A. Journal of Personality and Social Psychology, 12:194–199.

Osgood, C.E., Suci, G.J. & Tannenbaum, P.H. (1957). The Measurement of Meaning. University of Illinois Press: Urbana, IL, USA.

Parise, C.V. & Pavani, F. (2011). Evidence of sound symbolism in simple vocalizations. Experimental Brain Research, 214:373–380.

Peer, E., Brandimarte, L., Samat, S. & Acquisti, A. (2017). Beyond the Turk: Alternative platforms for crowdsourcing behavioral research. Journal of Experimental Social Psychology, 70:153–163.

Perlman, M. & Lupyan, G. (2018). People can create iconic vocalizations to communicate various meanings to naïve listeners. Scientific Reports, 8:2634, doi: 10.1038/s41598-018-20961-6

Peterson, G. E., & Lehiste, I. (1960). Duration of syllable nuclei in English. Journal of the Acoustical Society of America, 32:693–703.

Preziosi, M.A. & Coane, J.H. (2017). Remembering that big things sound big: sound symbolism and associative memory. Cognitive Research: Principles & Implications, 2:10, doi: 10.1186/s41235-016-0047-y

Ramachandran, V.S. & Hubbard, E.M. (2001). Synaesthesia – a window into perception, thought and language. Journal of Consciousness Studies, 8:3–34.

Recasens, D. (1985). Coarticulatory patterns and degrees of coarticulatory resistance in Catalan CV sequences. Language & Speech, 28: 97–114.

Reetz, H. & Jongman, A. (2020). Phonetics: Transcription, Production, Acoustics, and Perception, 2^nd^ edition. John Wiley & Sons, Inc., Hoboken NJ.

Rogers, S.K. & Ross, A.S. (1975). A cross-cultural test of the Maluma-Takete phenomenon. Perception, 4:105–106.

Saji, N., Akita, K., Kantartzis, K., Kita, S. & Imai, M. (2019). Cros-linguistically shared and language-specific sound symbolism in novel words elicited by locomotion videos in Japanese and English. PLoS ONE, 14:e0218707.

Sakamoto, M. & Watanabe, J. (2018). Bouba/kiki in touch: Associations between tactile perceptual qualities and Japanese phonemes. Frontiers in Psychology, 9:295, doi:10.3389/fpsyg.2018.00295

Sapir, E. (1929). A study in phonetic symbolism. Journal of Experimental Psychology, 12, 225–239.

Schmidtke, D., Körner, A., Glim, S. & Rummer, R. (2025). Valence sound symbolism facilitates classification of vowels and emotional facial expressions. Journal of Experimental Psychology: Learning, Memory, & Cognition, 51:661–667.

Shadle, C.H. (1985). Intrinsic fundamental frequency of vowels in sentence context. Journal of Acoustical Society of America, 78:1562–1567.

Shinohara, K., & Kawahara, S. (2010). A cross-linguistic study of sound symbolism: The images of size. Annual Meeting of the Berkeley Linguistics Society, 36:396–410.

Sidhu, D.M. & Pexman, P.M. (2018). Five mechanisms of sound symbolic association. Psychonomic Bulletin & Review, 25:1619–1643.

Sidhu, D.M., Vigliocco, G. & Pexman, P.M. (2022). Higher order factors of sound symbolism. Journal of Memory & Language, 125:104323.

Spence, C. (2011). Crossmodal correspondences: A tutorial review. *Attention*, Perception, and Psychophysics, 73:971–995.

Srinivasan, M.A. & LaMotte, R.H. (1995). Tactual discrimination of softness. Journal of Neurophysiology, 73:88–101.

Styles, S.J. & Gawne, L. (2017). When does Maluma/Takete fail? Two key failures and a meta-analysis suggest that phonology and phonotactics matter. i-Perception, 8:2041669517724807.

Sučević, J., Savić, A.M., Popović, M.B., Styles, S.J. & Ković, V. (2015). Balloons and bavoons versus spikes and shikes: ERPs reveal shared neural processes for shape-sound-meaning congruence in words, and shape-sound congruence in pseudowords. Brain & Language, 145/146, 11–22.

Svantesson, J.-O. (2017). Sound symbolism: the role of word sound in meaning. WIREs Cognitive Sci. 8:e1441, doi:10.1002/wcs.1441

Teixeira, J. P., & Fernandes, P. O. (2014). Jitter, shimmer and HNR classification within gender, tones and vowels in healthy voices. Procedia Technology, 16:1228–1237.

Thompson, P.D. & Estes, Z. (2011). Sound symbolic naming of novel objects is a graded function. Quarterly Journal of Experimental Psychology, 64, 2392–2404.

Tzeng, C.T., Duan, J., Namy, L.L. & Nygaard, L.C. (2018). Prosody in speech as a source of referential information. *Language*, Cognition & Neuroscience, 33:512–526.

Tzeng, C. Y., Nygaard, L. C., & Namy, L. L. (2017). The Specificity of Sound Symbolic Correspondences in Spoken Language. Cognitive Science, 41, 2191–2220.

Umeda, N. (1975). Vowel duration in American English. Journal of the Acoustical Society of America, 58:434–445.

Umeda, N. (1977). Consonant duration in American English. Journal of the Acoustical Society of America, 61:846–858.

Uno, R., Shinohara, K., Hosokawa, Y., Atsumi, N., Kumagai, G. et al. (2020). What’s in a villain’s name? Sound symbolic values of voiced obstruents and bilabial consonants. Review of Cognitive Linguistics, 18:428–457.

Valchev, N., Tidoni, E., Hamilton, A., Gazzola, V. & Avenanti, A. (2017). Primary somatosensory cortex necessary for the perception of weight from other people’s actions: a continuous theta-burst TMS experiment. NeuroImage, 152:195–206.

Van Puyvelde, M., Neyt, X., McGlone, F. & Pattyn, N. (2018). Voice stress analysis: A new framework for voice and effort in human performance. Frontiers in Psychology, 9:1994, doi:10.3389/fpsyg.2018.01994

van Son, R.J.J.H., & van Santen, J.P.H. (2005). Duration and spectral balance of intervocalic consonants: A case for efficient communication. Speech Communication, 47:100–123.

Villegas, J., Akita, K. & Kawahara, S. (2023). Psychoacoustic features explain subjective size and shape ratings of pseudo-words. Proceedings of Forum Acusticum, the 10^th^ Convention of the European Acoustics Association, Turin, Italy, September 2023.

Walker, P. & Parameswaran, C.R. (2019). Cross-sensory correspondences in language: vowel sounds can symbolize the felt heaviness of objects. Journal of Experimental Psychology: Learning, Memory, & Cognition, 45:246–252.

Warriner, A.B., Kuperman, V. & Brysbaert, M. (2013). Norms of valence, arousal, and dominance for 13,915 English lemmas. Behavior Research Methods, 45:1191–1207.

Westbury, C., Hollis, G., Sidhu, D.M. & Pexman, P.M. (2018). Weighing up the evidence for sound symbolism: distributional properties predict cue strength. Journal of Memory & Language, 99, 125–150.

Whalen, D.H. & Levitt, A.G. (1995). The universality of intrinsic F0 of vowels. Journal of Phonetics, 23:349–366.

Whitehead, R.L., Metz, D.E. & Whitehead, B.H. (1984). Vibratory patterns of the vocal folds during pulse register phonation. Journal of the Acoustical Society of America, 75:1293–1297.

Wicker, F.W. (1968). Mapping the intersensory regions of perceptual space. American Journal of Psychology, 81:178–188.

Winter, B., & Grice, M. (2021). Independence and generalizability in linguistics. Linguistics, 59(5), 1251–1277.

Winter, B., Lupyan, G., Perry, L.K., Dingemanse, M. & Perlman, M. (2023). Iconicity ratings for 14000+ English words. Behavior Research Methods, 56:1640–1655.

Winter, B. & Perlman, M. (2021). Size sound symbolism in the English lexicon. Glossa, 6:1646. doi: 10.5334/gjgl.1646

Winter, B., Perlman, M., Perry, L.K. & Lupyan, G. (2017). Which words are most iconic? Iconicity in English sensory words. Interaction Studies, 18:430–451.

Winter, B., Sóskuthy, M., Perlman, M. & Dingemanse, M. (2022). Trilled /r/ is associated with roughness, linking sound and touch across spoken languages. Scientific Reports, 12:1035, doi:10.1038/s41598-021-04311-7

Winter, B., Woodin, G. & Perlman, M. (in press). Defining iconicity for the cognitive sciences. In O. Fischer, K. Akita & P. Pernis (Eds.) The Oxford Handbook of Iconicity in Language, Oxford University Press: Oxford, UK.

Wong, L.S., Kwon, J., Zheng, Z., Styles, S.J., Sakamoto, M. et al. (2022). Japanese sound-symbolic words for representing the hardness of an object are judged similarly by Japanese and English speakers. Frontiers in Psychology, 13:830306, doi:10.3389/fpsyg.2022.830306

Woods, K. J., Siegel, M. H., Traer, J., & McDermott, J. H. (2017). Headphone screening to facilitate web-based auditory experiments. *Attention, Perception*, & Psychophysics, 79:2064–2072.

Xu, Y. & Xu, A. (2021). Consonantal F0 perturbation in American English involves multiple mechanisms. Journal of the Acoustical Society of America, 149:2877–2895.

Yarkoni, T. (2022). The generalizability crisis. Behavioral and Brain Sciences, 45 e1:1–78.

Yu, C.S.-P., McBeath, M.K. & Glenberg, A.M. (2021). The gleam-glum effect: /i:/ versus /˄/ phonemes generically carry emotional valence. Journal of Experimental Psychology: Learning, Memory, & Cognition, 47:1173–1185.

