## Supplementary Material for "Domain-specificity in phonological and acoustic characteristics of pseudoword iconicity across multiple meaning domains"

### DOMAIN-SPECIFICITY OF ICONIC PHONOLOGICAL AND ACOUSTIC ASSOCIATIONS FOR PSEUDOWORDS ACROSS MULTIPLE MEANING DOMAINS.

Departments of Neurology[1], Neuroscience & Experimental Therapeutics[2], Penn State College of Medicine, Hershey, PA 17033, USA; Department of Psychology[3], Penn State University, College of Liberal Arts, State College, PA 16802, USA;
Department of Psychology[4], Emory University, Atlanta, GA 30322, USA;
[5]Present address: Department of Psychological & Brain Sciences, Washington University in St. Louis, St. Louis MO 63130

Corresponding authors:

**Supplementary Methods**

*Computation of speech envelope, spectral tilt, and FFT, and their coefficients of variation*

The speech envelope was calculated using the root mean square of the amplitude and a sliding window length of 120 data points; the window was centered over each data point and moved one data point at a time along the vector of 10077 data points, thus giving 10077 measurements per pseudoword. The coefficient of variation (CV) for the speech envelope of each pseudoword was obtained by dividing the standard deviation of the 10077 measurement samples by the mean of those samples.

The calculation of any spectral parameter is limited by the Nyquist frequency (0.5 x sampling rate) to avoid aliasing, or false signals (Reetz & Jongman, 2020). To calculate spectral tilt, we first generated the power spectral density function over the 10077 data points of the normalized pseudoword duration. The resulting power spectral density is mirror-symmetrical along the full vector of 10077 datapoints: applying the Nyquist frequency limit reduces this to a vector with length 5038 and removes the symmetry. Over the vector of 5038 datapoints, we used a sliding window of 600 data points with overlap of 15, i.e, after the first 600 data points, the window moves forward 585 data points until it goes past the end of the vector. The final, incomplete window was discarded. Within each window, we fitted a polynomial to the 600 data points and estimated the slope from the polynomial coefficient of the periodogram (power across frequency). This provided 8 measurement windows, corresponding to 8 slopes, providing a measure of spectral tilt over time. Note that at the end of the vector, the final, incomplete window was 343 data points; discarding this represents a loss of 6.8% of the data. The CV for the spectral tilt of each pseudoword was calculated by dividing the standard deviation of the 8 slope values by their mean.

The FFT was calculated with a sliding time window of 100 data points with an overlap of 80, and a frequency window of 100 discrete Fourier transform points from 1 to 11 kHz. The output of the spectrogram function in MATLAB was a vector of length 499 for each frequency; since we only used a limited range of the frequencies, i.e., 1-11 kHz, the vectors were concatenated, resulting in 5489 measurements per pseudoword. The CV for the FFT of each pseudoword was obtained by dividing the standard deviation of the 5489 measurement samples by their mean.

**Supplementary Table 1:** Anchor words for the categorical opposites for each scale.

| Domain | Categorical opposites | Anchor words |
|---|---|---|
| Shape | Rounded | rounded; bloblike; amoeboid |
| | Pointed | pointed; angular; jagged |
| Roughness | Smooth | smooth; sleek; polished |
| | Rough | rough; coarse; bristly |
| Hardness | Hard | hard; unyielding; firm |
| | Soft | soft; squishy; cushioned |
| Weight | Light | light; weightless; airy |
| | Heavy | heavy; weighty; hefty |
| Size | Small | small; tiny; little |
| | Big | big; large; huge |
| Brightness | Bright | bright; light; luminous |
| | Dark | dark; dim; unlit |
| Arousal | Calming | calming; tranquil; passive |
| | Exciting | exciting; exhilarating; active |
| Valence | Good | good; nice; excellent |
| | Bad | bad; nasty; terrible |

**Supplementary Table 2:** Phonetic feature regression: multicollinearity statistics; VIF = variance inflation factor.

| Predictor | Tolerance | VIF |
|---|---|---|
| Stops | .39 | 2.54 |
| Af/fricatives | .39 | 2.55 |
| Unvoiced | .84 | 1.19 |
| Alveolar 1 | .72 | 1.39 |
| Alveolar 2 | .74 | 1.35 |
| Post-alveolar/velar 1 | .71 | 1.41 |
| Post-alveolar/velar 2 | .72 | 1.38 |
| Back rounded | .95 | 1.05 |
| High 1 | .94 | 1.06 |
| High 2 | .98 | 1.02 |

**Supplementary Table 3:** Phonetic feature regression: Durbin-Watson test values.

| **Domain** | **Scale** | **Test value** |
|---|---|---|
| Shape | Rounded | 2.23 |
| | Pointed | 2.17 |
| Roughness | Smooth | 2.03 |
| | Rough | 1.94 |
| Hardness | Hard | 1.89 |
| | Soft | 1.84 |
| Weight | Light | 1.94 |
| | Heavy | 1.81 |
| Arousal | Calming | 1.79 |
| | Exciting | 1.90 |
| Brightness | Bright | 2.10 |
| | Dark | 2.01 |
| Valence | Good | 1.93 |
| | Bad | 2.02 |
| Size | Small | 1.89 |
| | Big | 1.89 |

**Supplementary Figures 1a-h:** Homoscedasticity assessment for the phonetic category regression: histograms (top row) and Q-Q plots (middle row) for standardized residuals and scatterplots (bottom row) for standardized residuals against standardized predicted values.

Supplementary Figure 1a: Shape – rounded (left) & pointed (right)

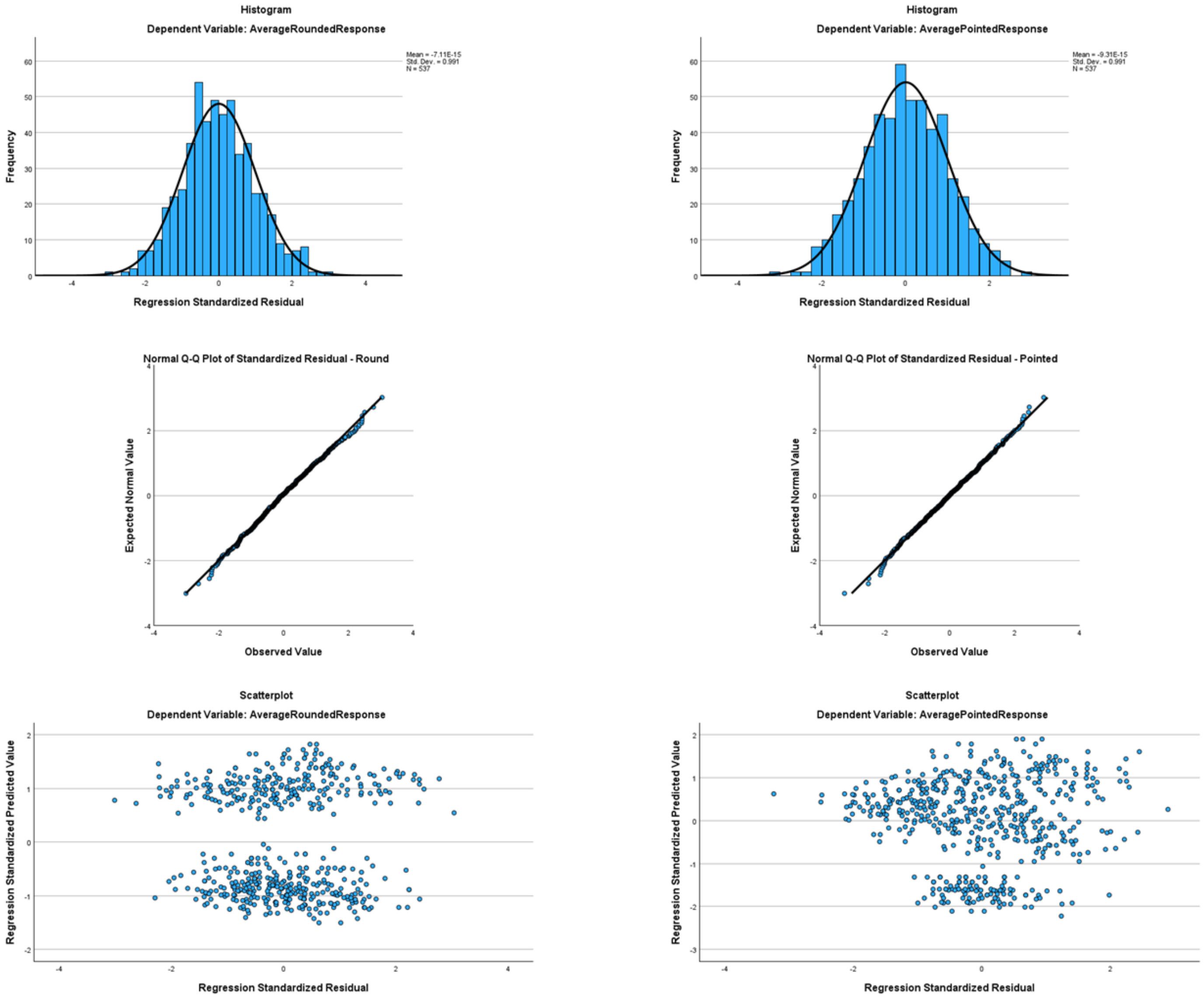

### Supplementary Figure 1b: Roughness – smooth (left) & rough (right)

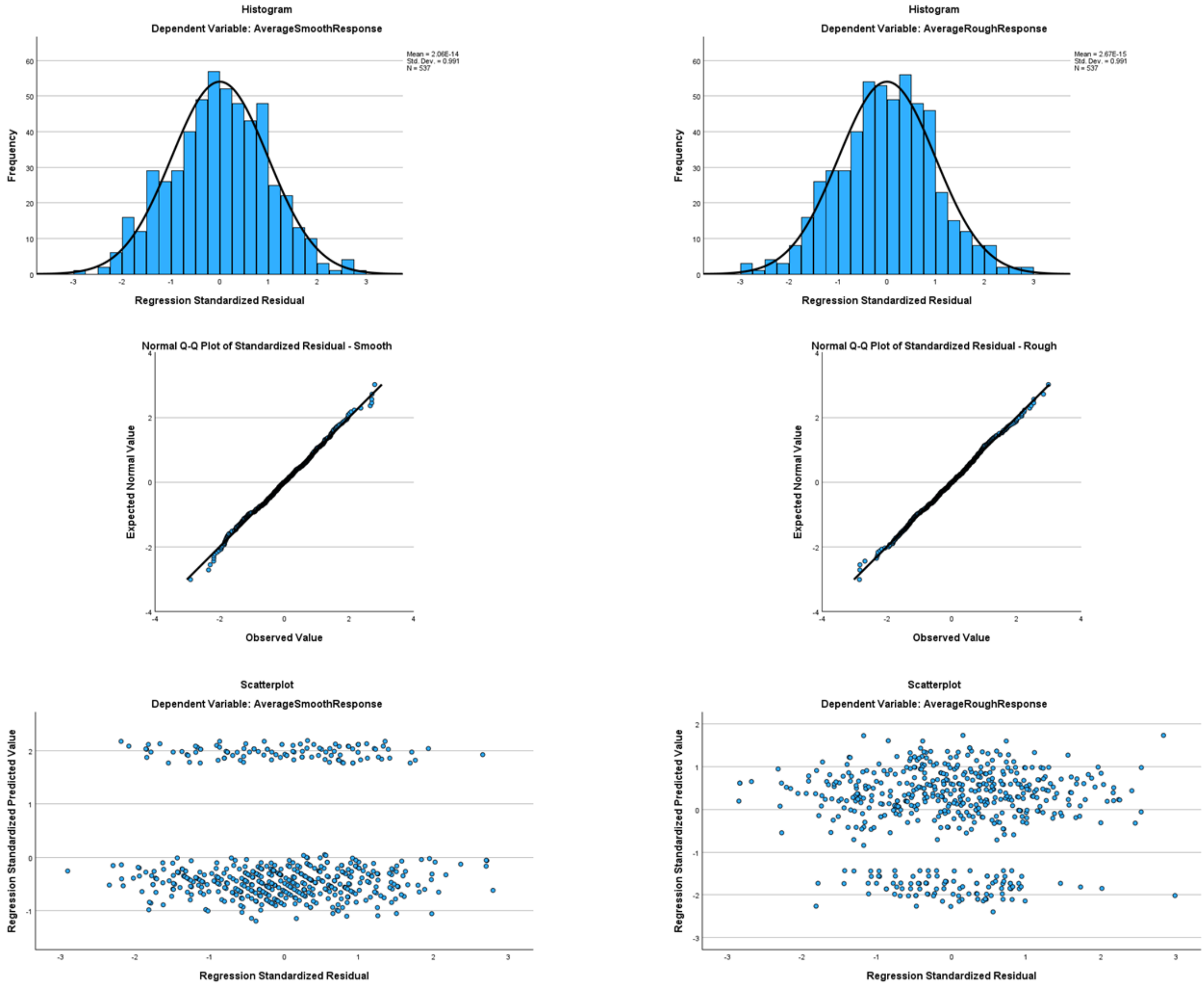

#### Supplementary Figure 1c: Hardness – hard (left) & soft (right)

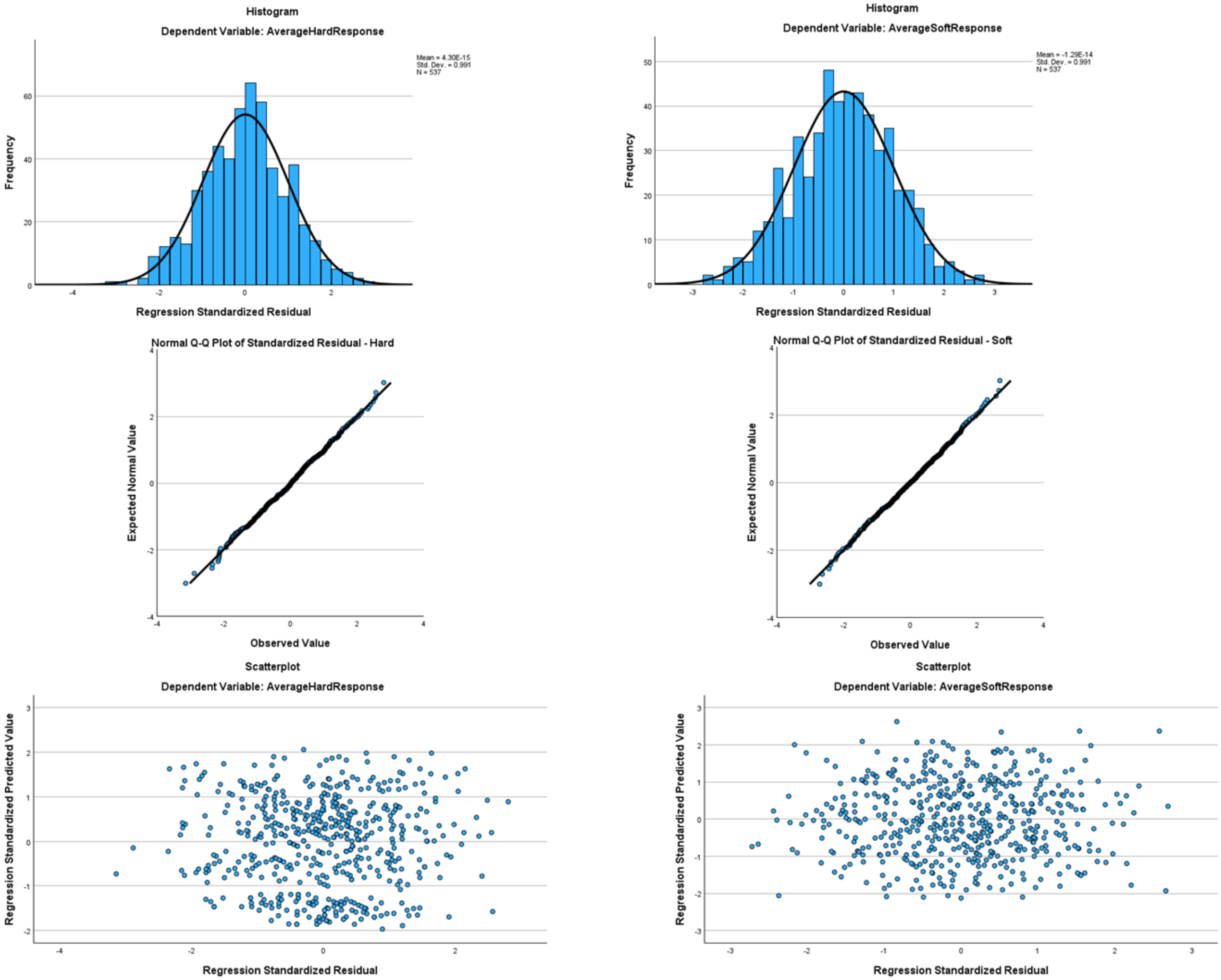

### Supplementary Figure 1d: Weight – light (left) & heavy (right)

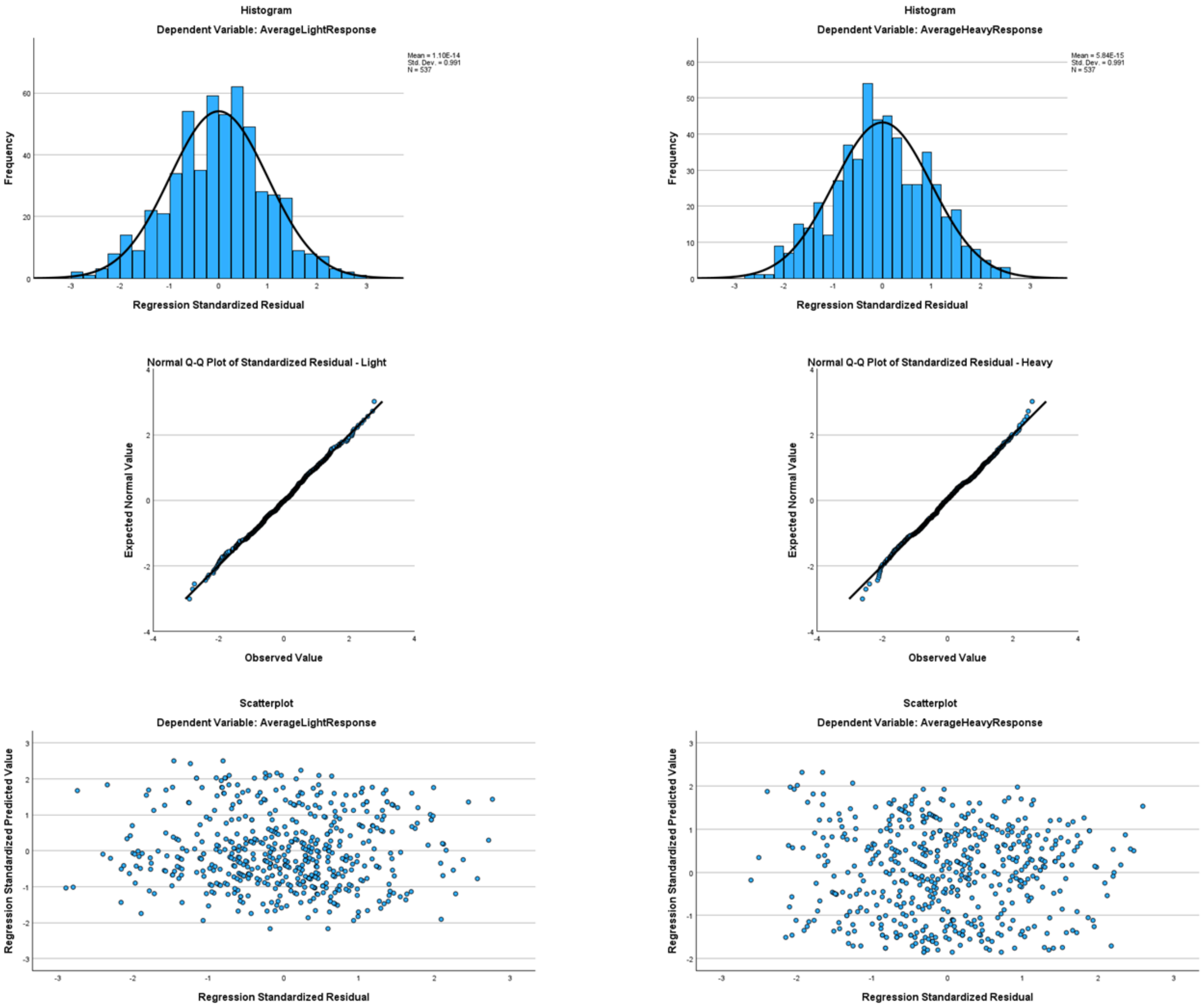

### Supplementary Figure 1e: Arousal – calming (left) & exciting (right)

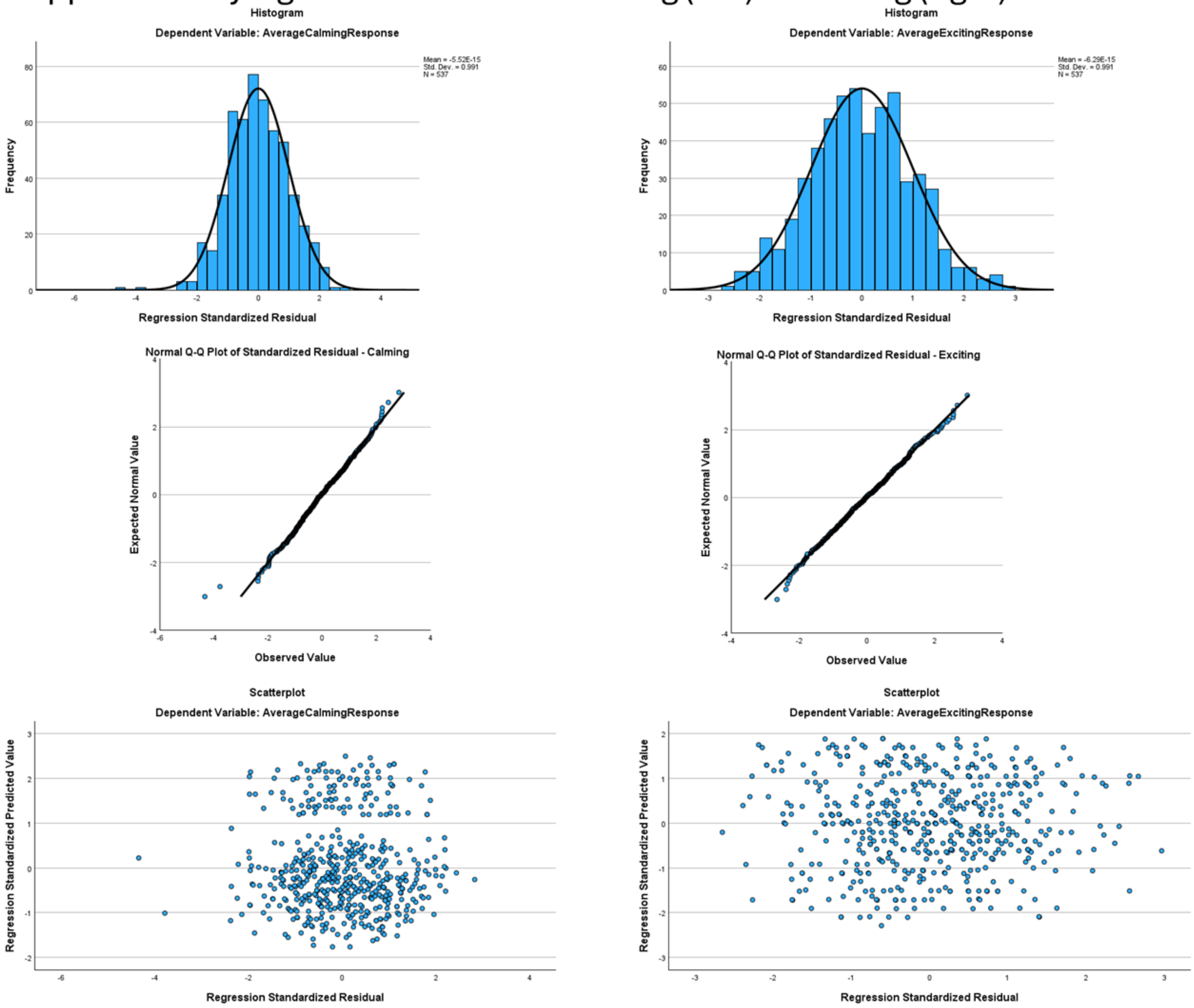

Supplementary Figure 1f: Brightness – bright (left) & dark (right)

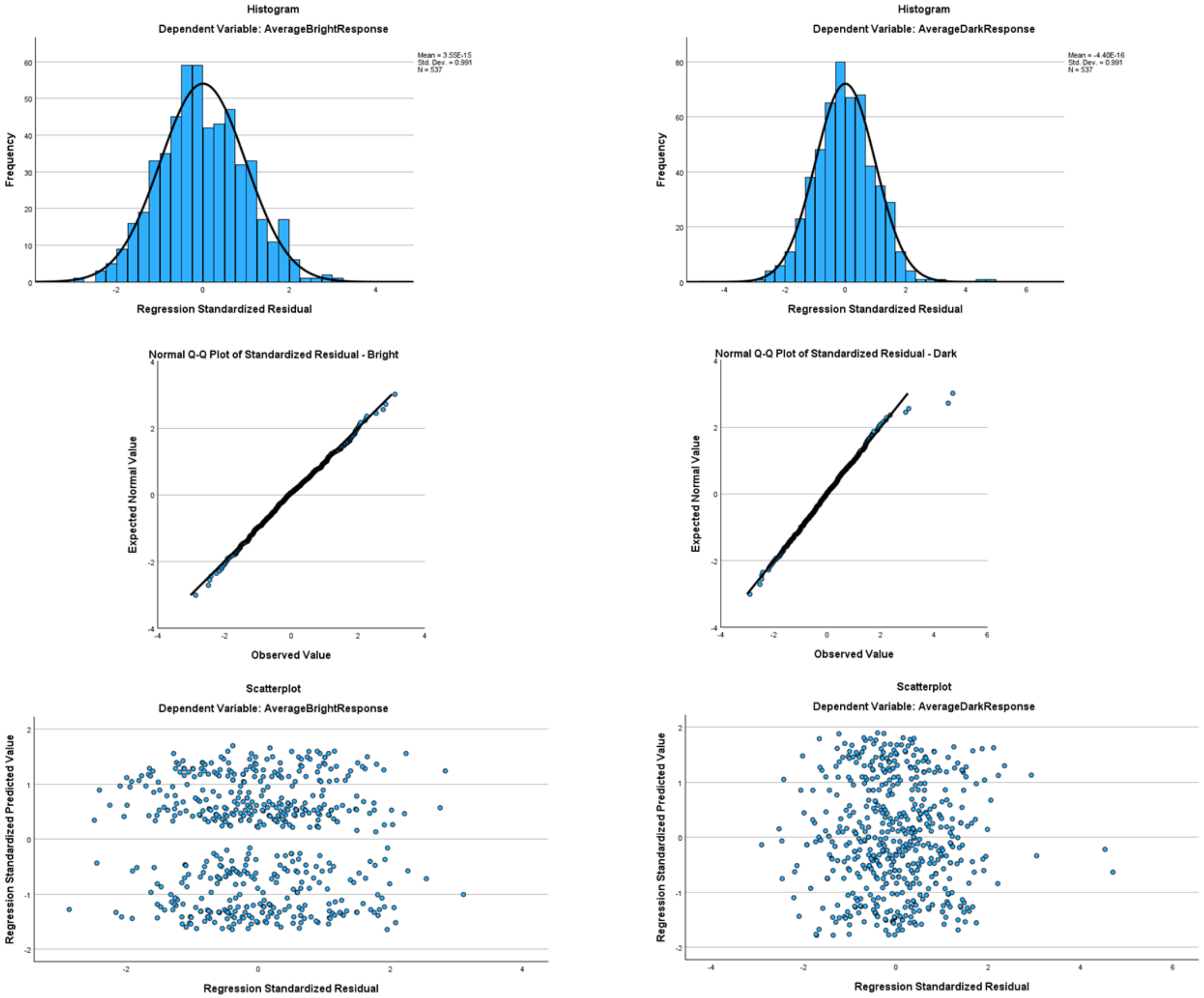

Supplementary Figure 1g: Valence – good (left) & bad (right)

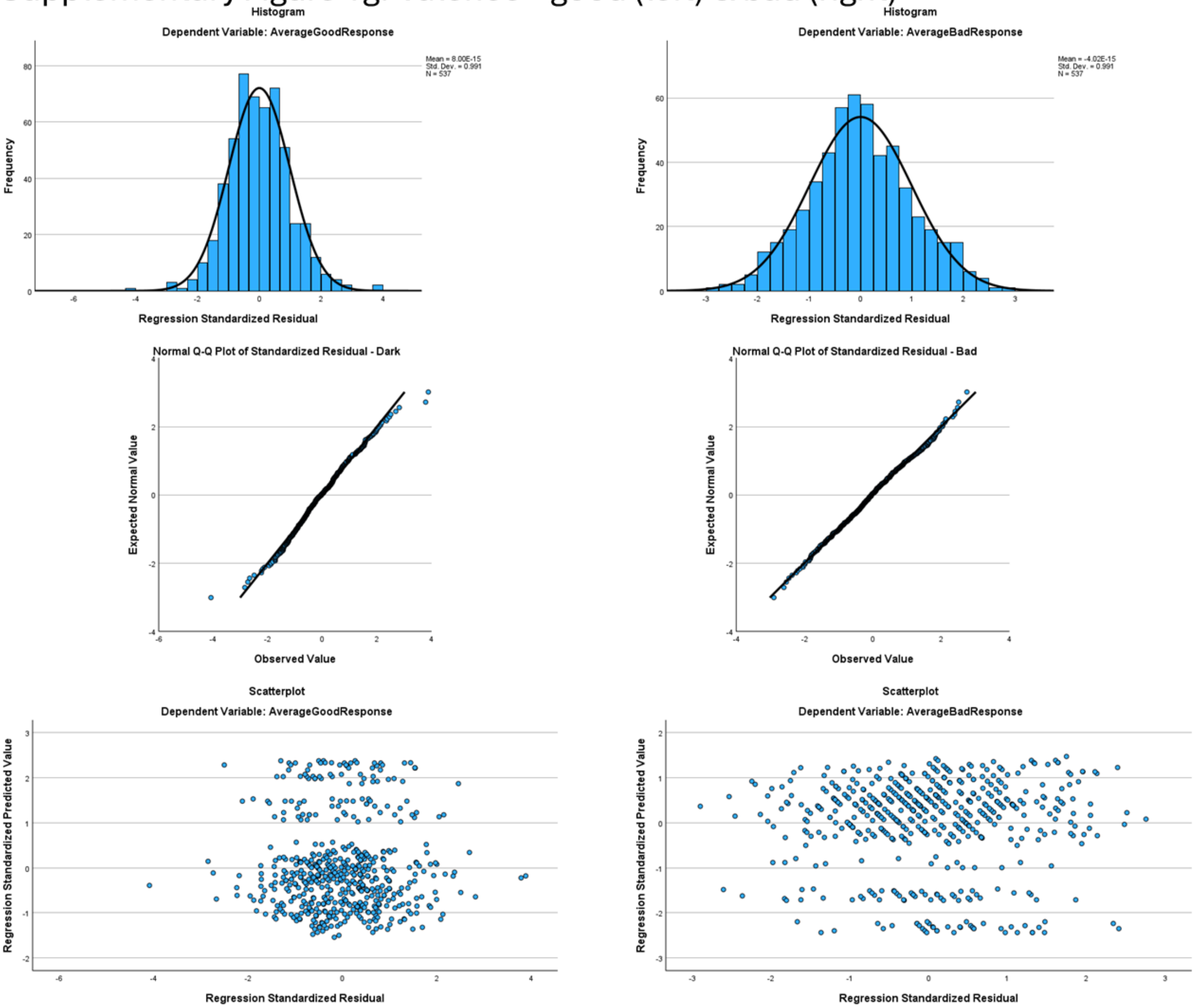

### Supplementary Figure 1h: Size – small (left) & big (right)

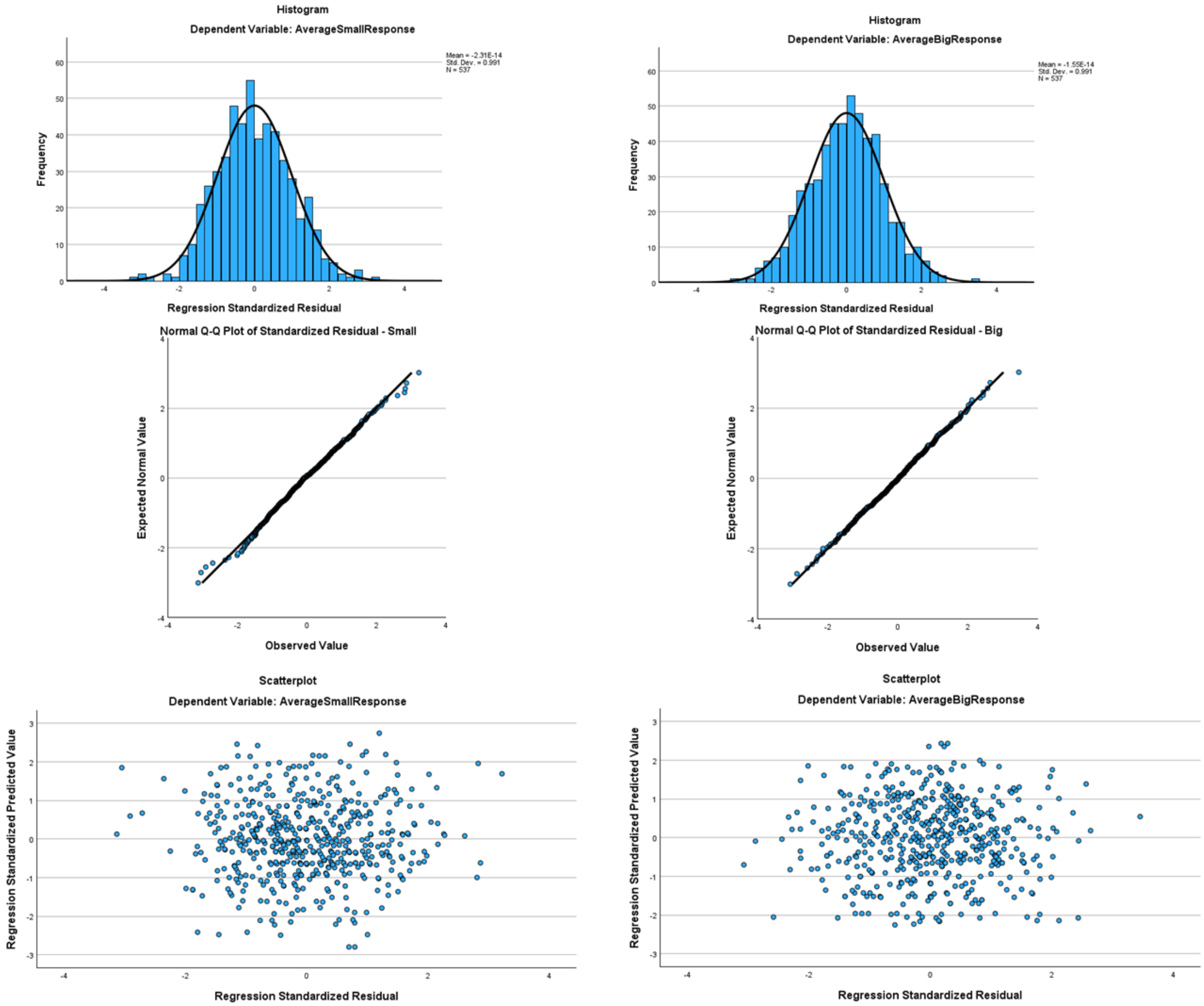

**Supplementary Table 4:** Phonetic feature regression analysis for each domain. $R^2$, adjusted $R^2$ (Adj $R^2$), standardized beta coefficients (β), and the change in $R^2$ ($R^2Δ$) attributable to each predictor tested against the full model (significant changes only). The dummy-coded reference predictors are italicized; (a) manner and (b) place of articulation for consonants; (c) manner and (d) place of articulation for vowels; 1,2, first or second consonant or vowel position; all F-values $p < .001$; * $p < .05$, ** $p < .01$, *** $p < .001$.

| | | **Shape** | | | | **Roughness** | | | | **Hardness** | | | | **Weight** | | | |
|---|---|---|---|---|---|---|---|---|---|---|---|---|---|---|---|---|---|
| | | Rounded | | Pointed | | Smooth | | Rough | | Hard | | Soft | | Light | | Heavy | |
| | **$R^2$ (Adj $R^2$)** | .88 (.87) | | .76 (.75) | | .57 (.56) | | .77 (.77) | | .68 (.67) | | .33 (.31) | | .43 (.42) | | .53 (.52) | |
| | **Consonants** | **β** | **$R^2Δ$** | **β** | **$R^2Δ$** | **β** | **$R^2Δ$** | **β** | **$R^2Δ$** | **β** | **$R^2Δ$** | **β** | **$R^2Δ$** | **β** | **$R^2Δ$** | **β** | **$R^2Δ$** |
| (a) | *Sonorants* | | | | | | | | | | | | | | | | |
| | Stops | -.18*** | .01 | .69*** | .17 | -.87*** | .29 | .67*** | .18 | .81*** | .25 | -.55*** | .12 | -.74*** | .21 | .82*** | .26 |
| | Af/fricatives | -.23*** | .02 | .50*** | .10 | -.82*** | .26 | .85*** | .28 | .36*** | .05 | -.24*** | .02 | -.50*** | .10 | .45*** | .08 |
| | *Voiced* | | | | | | | | | | | | | | | | |
| | Unvoiced | -.07*** | <.01 | .38*** | .12 | .02 | | .17*** | .02 | .06* | <.01 | .01 | | .15*** | .02 | -.25*** | .05 |
| (b) | *Bilabial/labiodental* | | | | | | | | | | | | | | | | |
| | Alveolar 1 | -.12*** | .01 | .09*** | .01 | <-.01 | | .11*** | .01 | .04 | | -.16*** | .02 | -.13*** | .01 | -.04 | |
| | Alveolar 2 | -.08*** | <.01 | .13*** | .01 | -.02 | | .12*** | .01 | .13*** | .01 | -.21*** | .03 | -.15*** | .02 | .03 | |
| | Post-alveolar/velar 1 | -.02 | | .20*** | .03 | -.11*** | .01 | .24*** | .04 | .20*** | .03 | -.23*** | .04 | -.19*** | .02 | .13*** | .01 |
| | Post-alveolar/velar 2 | -.12*** | .01 | .28*** | .05 | -.12*** | .01 | .29*** | .06 | .36*** | .09 | -.25*** | .04 | -.29*** | .06 | .23*** | .04 |
| | **Vowels** | | | | | | | | | | | | | | | | |
| (c) | *Front unrounded* | | | | | | | | | | | | | | | | |
| | Back rounded | .91*** | .78 | -.12*** | .01 | -.02 | | -.13*** | .02 | -.05 | | .19*** | .03 | -.14*** | .02 | .12*** | .01 |
| (d) | *Middle* | | | | | | | | | | | | | | | | |
| | High 1 | -.04* | <.01 | .05* | <.01 | -.03 | | .06* | <.01 | -.03 | | -.08* | .01 | -.03 | | .02 | |
| | High 2 | -.09*** | .01 | < .01 | | -.08** | .01 | <-.01 | | -.07** | <.01 | -.01 | | .09* | .01 | -.09** | .01 |

| | | **Arousal** | | | | **Brightness** | | | | **Valence** | | | | **Size** | | | |
|---|---|---|---|---|---|---|---|---|---|---|---|---|---|---|---|---|---|
| | | Calming | | Exciting | | Bright | | Dark | | Good | | Bad | | Small | | Big | |
| | **$R^2$ (Adj $R^2$)** | .48 (.47) | | .36 (.35) | | .53 (.52) | | .35 (.34) | | .4 (.39) | | .28 (.26) | | .12 (.11) | | .29 (.27) | |
| | **Consonants** | **β** | **$R^2Δ$** | **β** | **$R^2Δ$** | **β** | **$R^2Δ$** | **β** | **$R^2Δ$** | **β** | **$R^2Δ$** | **β** | **$R^2Δ$** | **β** | **$R^2Δ$** | **β** | **$R^2Δ$** |
| (a) | *Sonorants* | | | | | | | | | | | | | | | | |
| | Stops | -.62*** | .15 | .18** | .01 | .05 | | .12* | <.01 | -.66*** | .17 | .5*** | .10 | .17** | .01 | .41*** | .06 |
| | Af/fricatives | -.55*** | .12 | .43*** | .07 | .01 | | .17** | .01 | -.74*** | .21 | .52*** | .10 | -.05 | | .43*** | .07 |
| | *Voiced* | | | | | | | | | | | | | | | | |
| | Unvoiced | -.09** | .01 | .13*** | .01 | .25*** | .05 | -.32*** | .09 | .1** | .01 | .04 | | .18*** | .03 | -.2*** | .03 |
| (b) | *Bilabial/labiodental* | | | | | | | | | | | | | | | | |
| | Alveolar 1 | -.06 | | .06 | | .03 | | -.03 | | .04 | | -.19*** | .02 | -.05 | | .05 | |
| | Alveolar 2 | -.11** | .01 | .06 | | .08* | <.01 | <-.01 | | -.02 | | -.01 | | -.08 | | .01 | |

| | | | | | | | | | | | | | | | | | |
|---|---|---|---|---|---|---|---|---|---|---|---|---|---|---|---|---|---|
| | Post-alveolar/velar 1 | -.14*** | .01 | .02 | | .1** | .01 | -.09* | .01 | -.02 | | | | -.23*** | .04 | .20*** | .03 |
| | Post-alveolar/velar 2 | -.28*** | .06 | .05 | | .06 | | <-.01 | | -.05 | | | | -.13** | .01 | .15*** | .02 |
| | **Vowels** | | | | | | | | | | | | | | | | |
| (c) | *Front unrounded* | | | | | | | | | | | | | | | | |
| | Back rounded | .22*** | .05 | -.45*** | .19 | -.67*** | .43 | .50*** | .24 | -.27*** | .07 | .19*** | .03 | -.06 | | .28*** | .07 |
| (d) | *Middle* | | | | | | | | | | | | | | | | |
| | High 1 | -.01 | | .11** | .01 | .04 | | -.03 | | .01 | | -.02 | | -.05 | | -.02 | |
| | High 2 | .05 | | <.01 | | .04 | | .07* | <.01 | -.1** | .01 | .03 | | .05 | | -.14*** | .02 |

**Supplementary Table 5:** Phoneme regression: multicollinearity statistics; VIF = variance inflation factor.

| Predictor | Tolerance | VIF |
|---|---|---|
| **C1** | | |
| [m] | .55 | 1.81 |
| [l] | .53 | 1.89 |
| [d] | .51 | 1.97 |
| [g] | .50 | 2.00 |
| [p] | .55 | 1.82 |
| [t] | .54 | 1.83 |
| [v] | .29 | 3.50 |
| [z] | .30 | 3.37 |
| [ʤ] | .31 | 3.22 |
| [s] | .51 | 1.86 |
| [ʧ] | .53 | 1.88 |
| **V1** | | |
| [u] | .57 | 1.76 |
| [ʊ] | .54 | 1.84 |
| [e] | .42 | 2.37 |
| [ɛ] | .42 | 2.39 |
| [i] | .47 | 2.15 |
| [ɪ] | .43 | 2.30 |
| **C2** | | |
| [m] | .29 | 3.40 |
| [n] | .31 | 3.28 |
| [l] | .32 | 3.14 |
| [d] | .55 | 1.82 |
| [g] | .52 | 1.91 |
| [p] | .32 | 3.11 |
| [t] | .32 | 3.13 |
| [k] | .29 | 3.39 |
| [z] | .54 | 1.84 |
| [ʤ] | .54 | 1.85 |
| [f] | .28 | 3.56 |
| [s] | .30 | 3.29 |
| [ʧ] | .30 | 3.37 |
| **V2** | | |
| [u] | .63 | 1.59 |
| [i] | .70 | 1.42 |

**Supplementary Table 6:** Phoneme regression: Durbin-Watson test values.

| Domain | Scale | Test value |
|---|---|---|
| Shape | Rounded | 2.35 |
| | Pointed | 1.98 |
| Roughness | Smooth | 1.98 |
| | Rough | 1.99 |
| Hardness | Hard | 1.92 |
| | Soft | 1.93 |
| Weight | Light | 2.04 |
| | Heavy | 1.82 |
| Arousal | Calming | 1.89 |
| | Exciting | 1.93 |
| Brightness | Bright | 2.05 |
| | Dark | 2.05 |
| Valence | Good | 1.90 |
| | Bad | 2.06 |
| Size | Small | 1.93 |
| | Big | 1.91 |

**Supplementary Figures 2a-h:** Homoscedasticity assessment for the phonemic regression: histograms (top row) and Q-Q plots (middle row) for standardized residuals and scatterplots (bottom row) for standardized residuals against standardized predicted values.

Supplementary Figure 2a: Shape – rounded (left) & pointed (right)

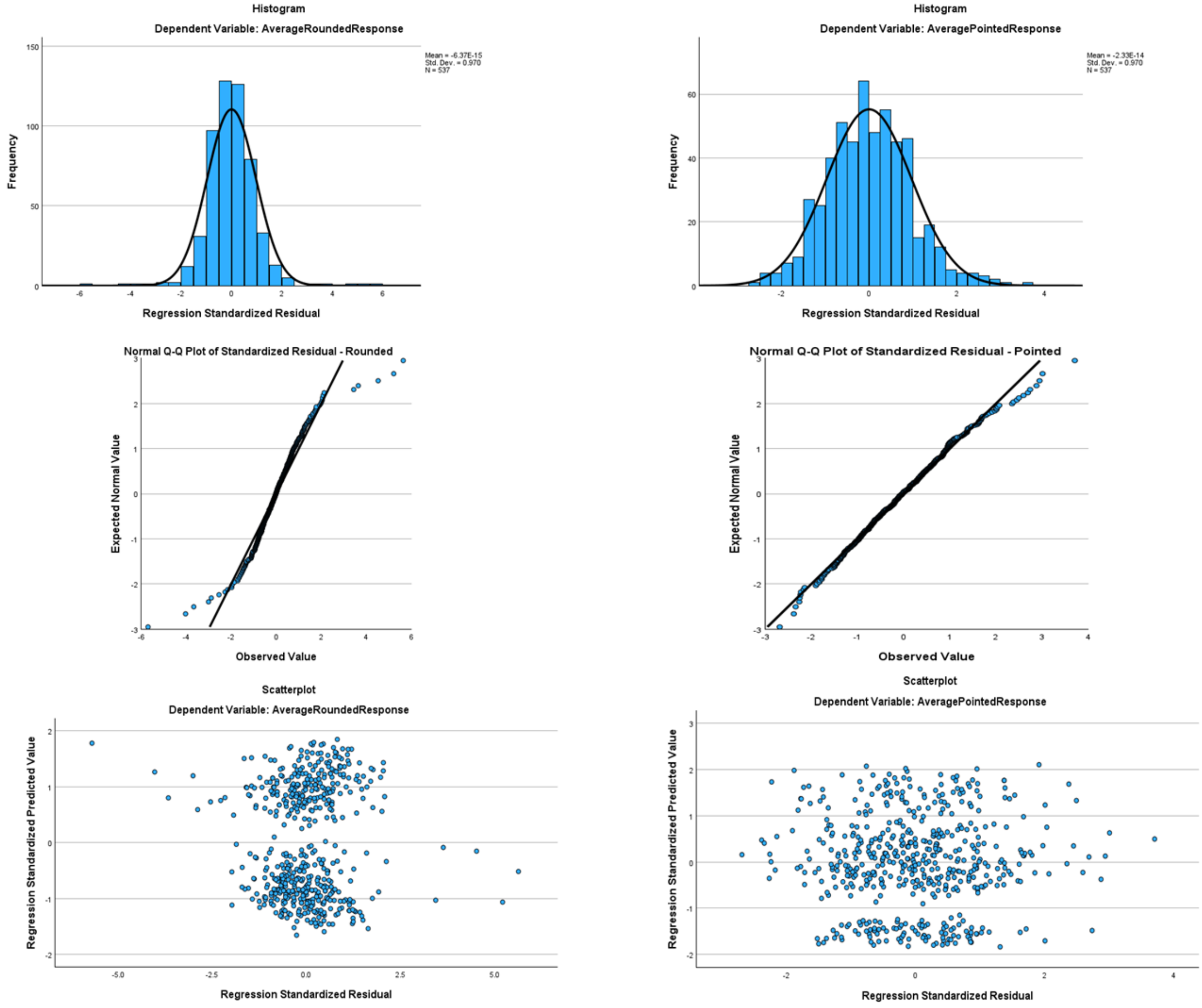

#### Supplementary Figure 2b: Roughness – smooth (left) & rough (right)

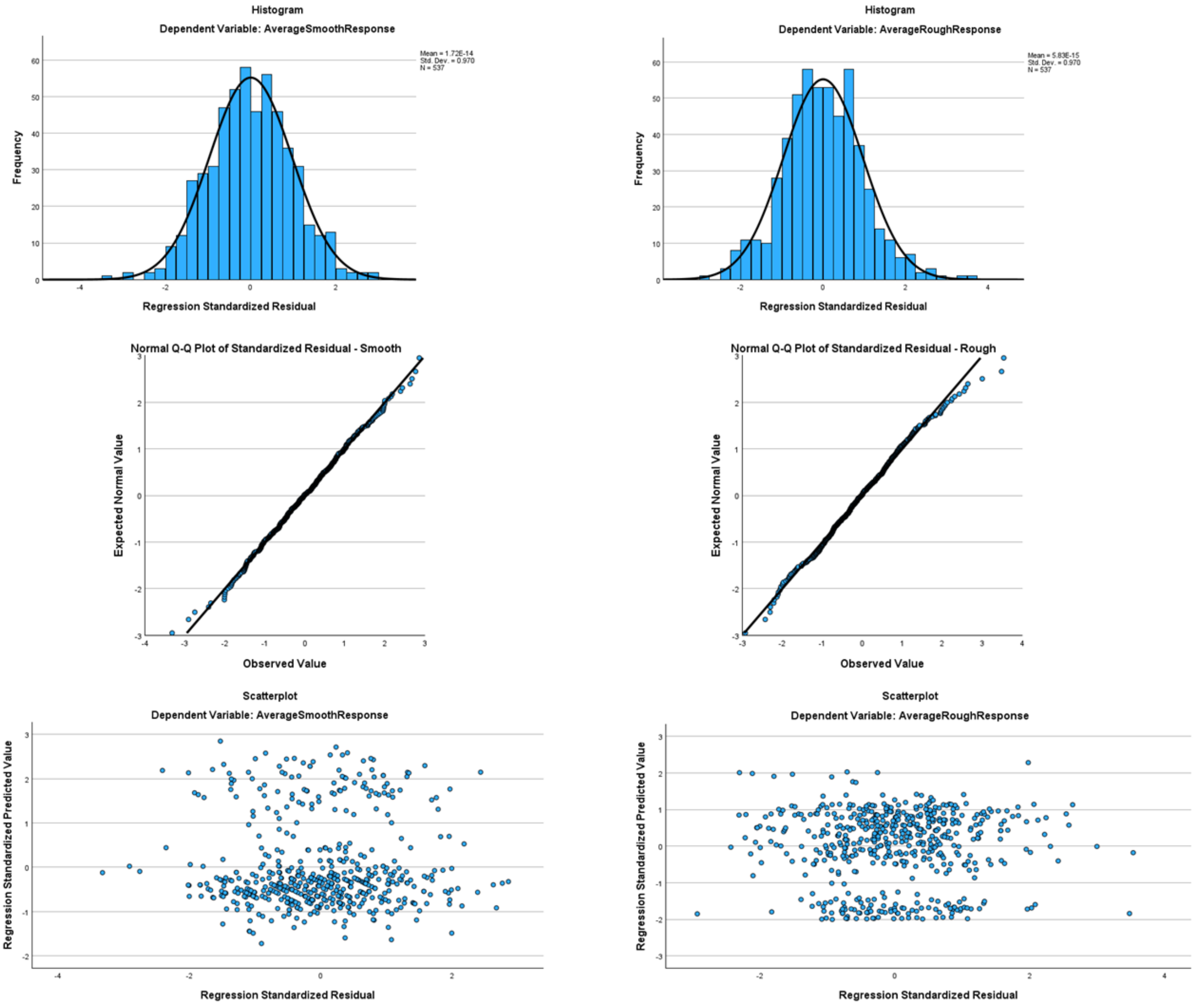

Supplementary Figure 2c: Hardness – hard (left) & soft (right)

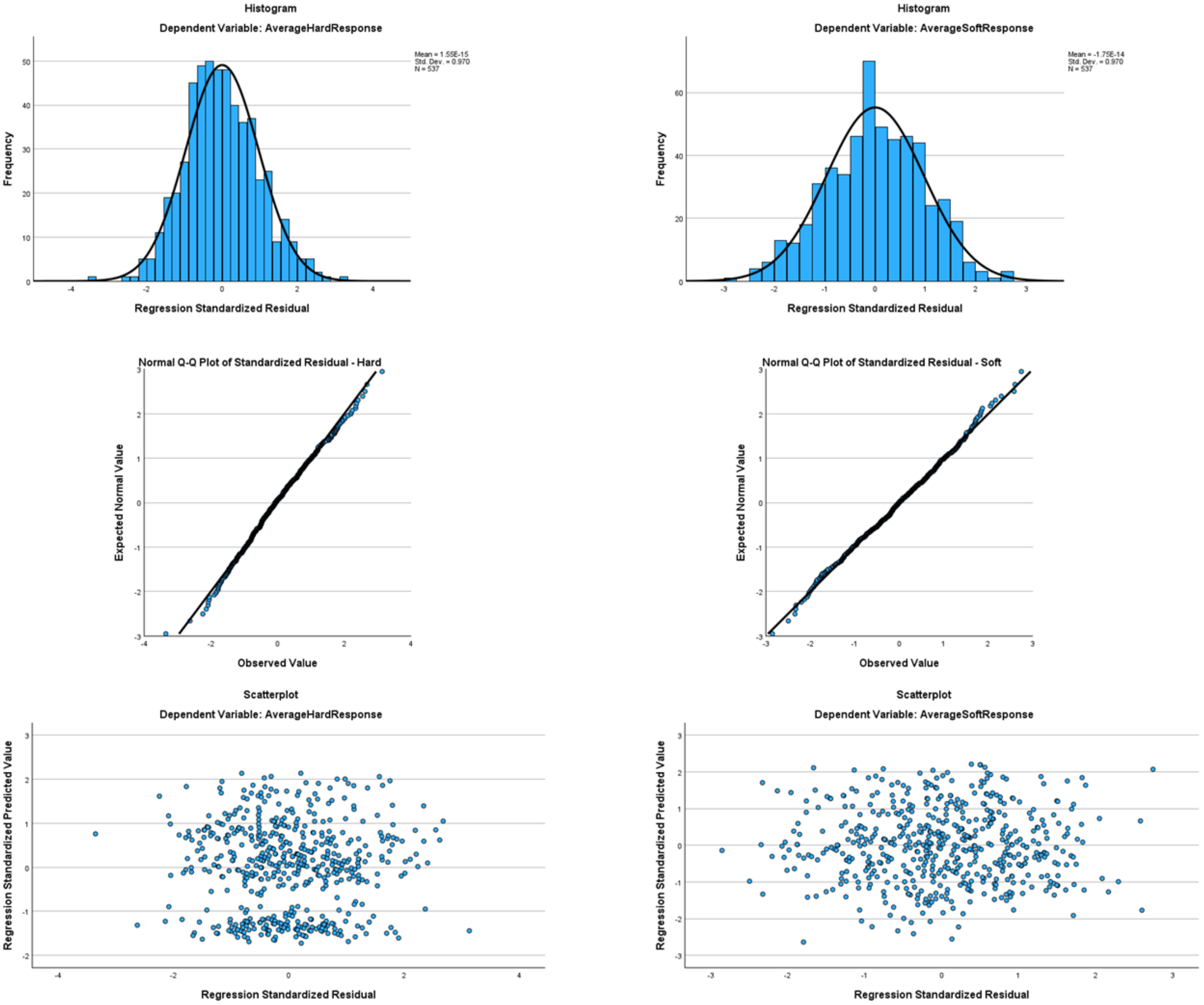

Supplementary Figure 2d: Weight – light (left) & heavy (right)

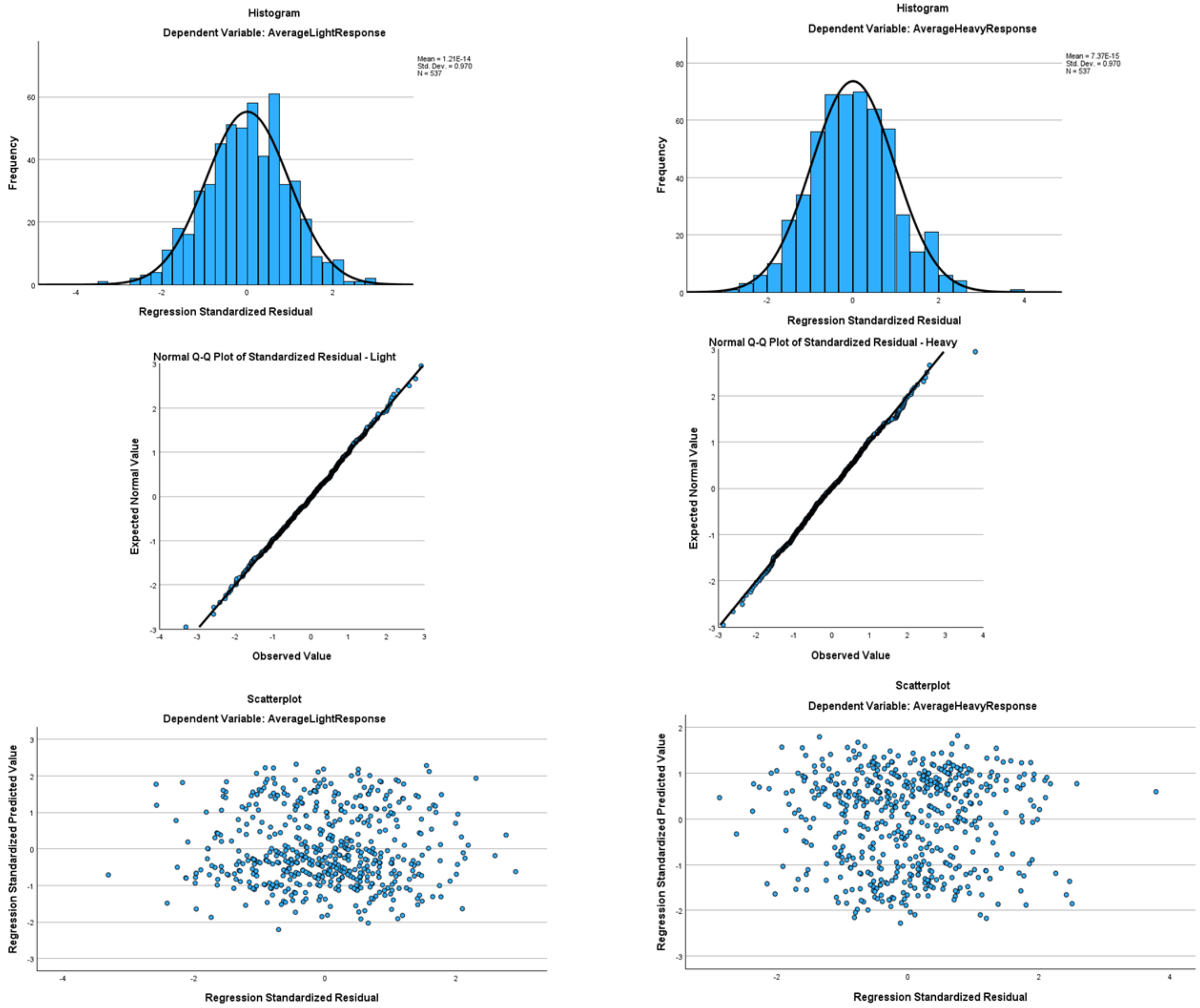

Supplementary Figure 2e: Arousal – calming (left) & exciting (right)

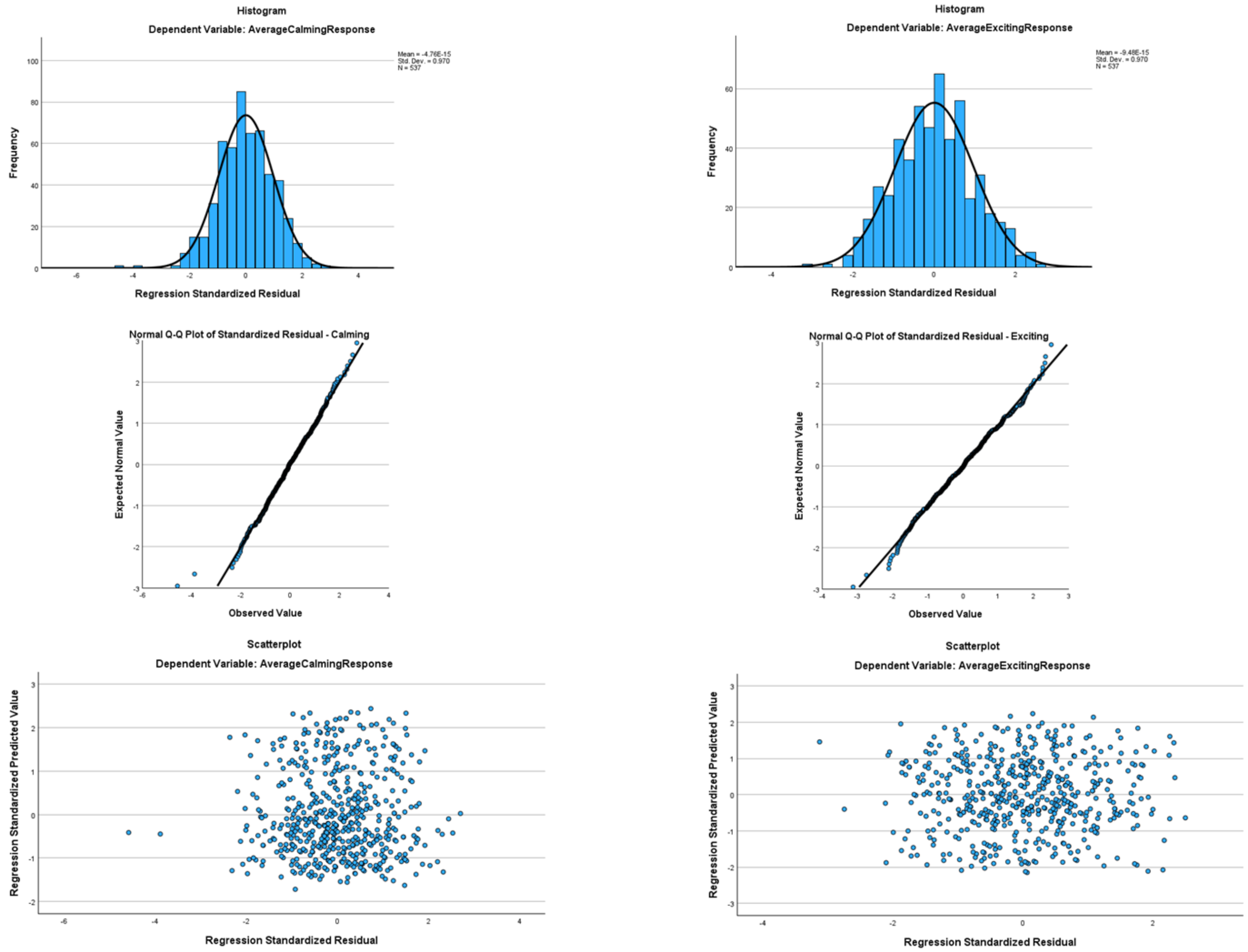

### Supplementary Figure 2f: Brightness – bright (left) & dark (right)

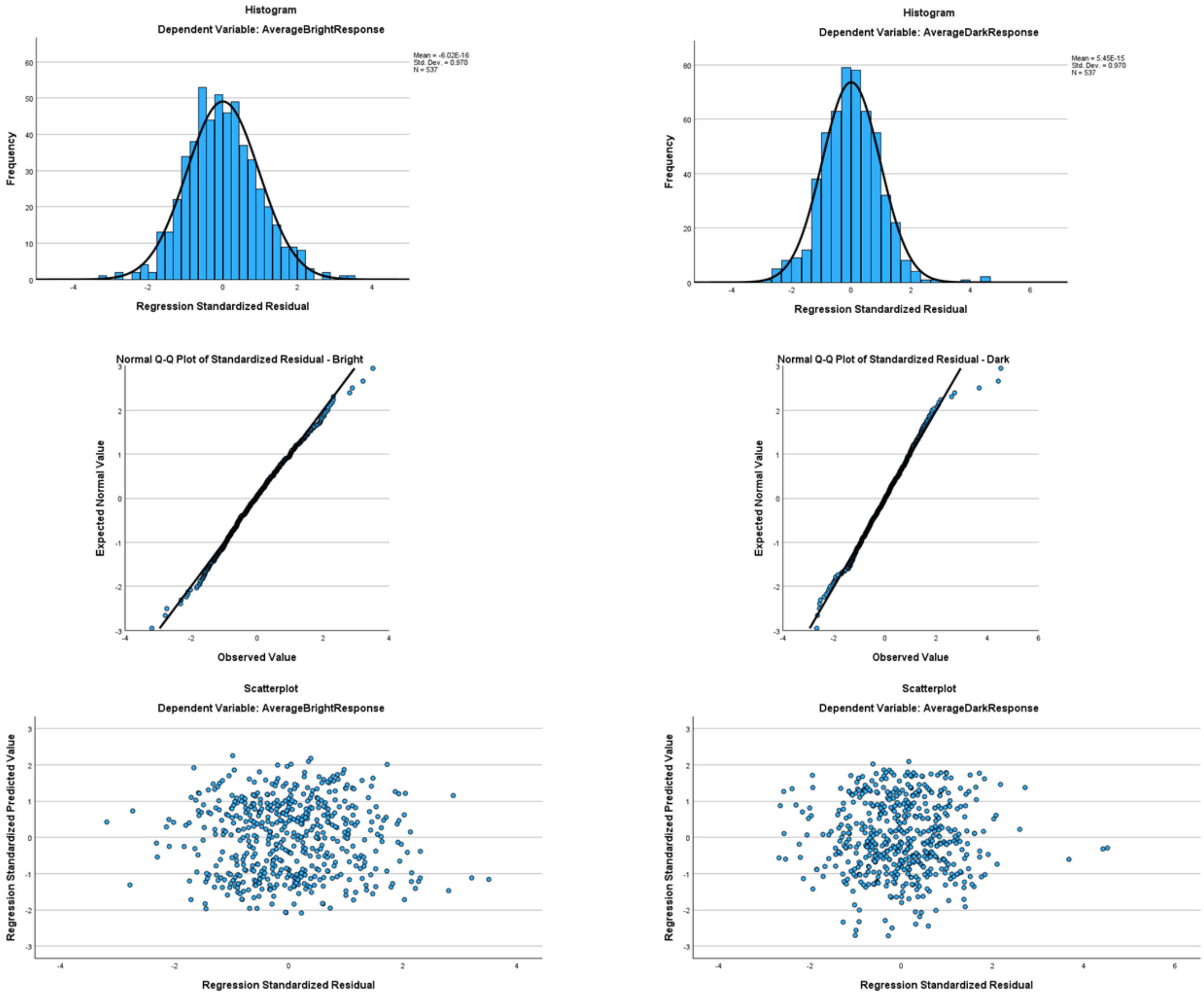

### Supplementary Figure 2g: Valence – good (left) & bad (right)

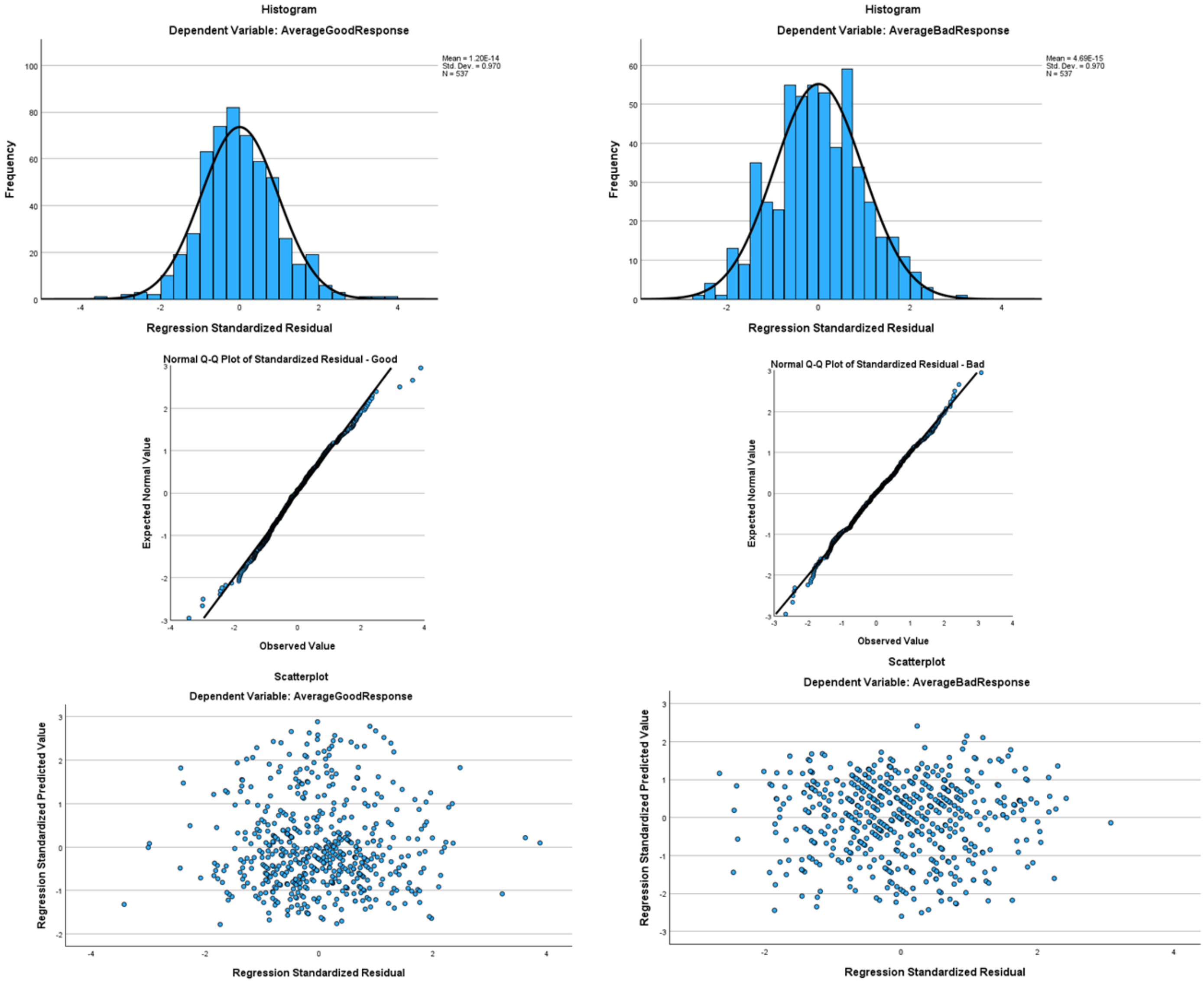

### Supplementary Figure 2h: Size – small (left) & big (right)

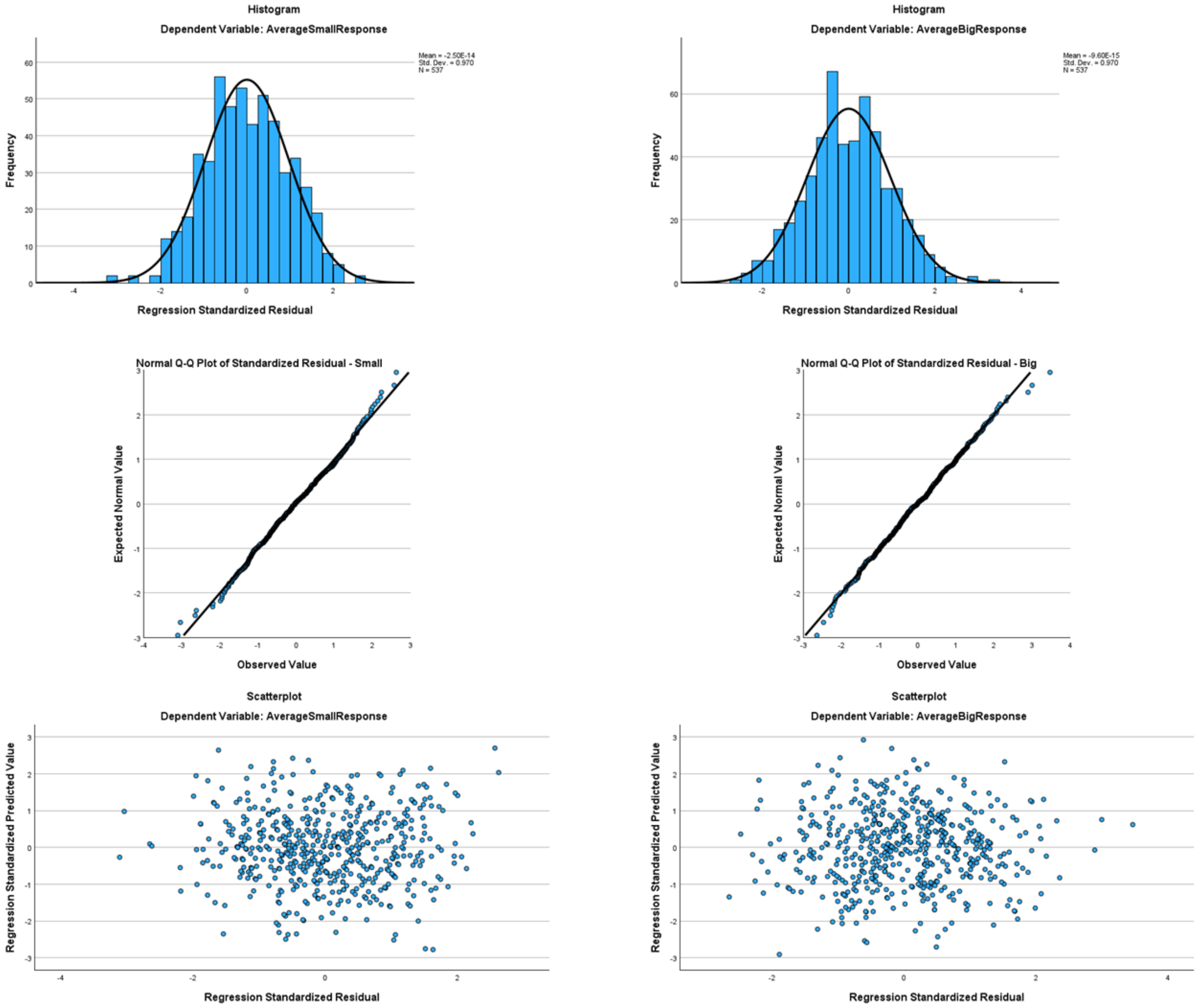

**Supplementary Table 7:** Phonemic regression analysis for each domain: $R^2$, Adj $R^2$, $\beta$, and $R^2\Delta$ as for Table 3. C1/2, V1/2: first and second consonant and vowel positions, respectively. Voiced consonants are in **bold** type. All F-values $p < .001$; p-values as for Supplementary Table 4.

| | **Shape** | | | | **Roughness** | | | | **Hardness** | | | | **Weight** | | | |
|---|---|---|---|---|---|---|---|---|---|---|---|---|---|---|---|---|
| | Rounded | | Pointed | | Smooth | | Rough | | Hard | | Soft | | Light | | Heavy | |
| **$R^2$ (Adj $R^2$)** | .83 (.82) | | .9 (.9) | | .66 (.63) | | .86 (.85) | | .8 (.79) | | .45 (.41) | | .52 (.48) | | .61 (.58) | |
| **C1** | **β** | **$R^2\Delta$** | **β** | **$R^2\Delta$** | **β** | **$R^2\Delta$** | **β** | **$R^2\Delta$** | **β** | **$R^2\Delta$** | **β** | **$R^2\Delta$** | **β** | **$R^2\Delta$** | **β** | **$R^2\Delta$** |
| *Sonorants* | | | | | | | | | | | | | | | | |
| **[m]** | .03 | | -.01 | | .06 | | <.01 | | .02 | | .09 | | .03 | | .04 | |
| **[l]** | -.06* | <.01 | .01 | | .09** | <.01 | .03 | | -.02 | | .01 | | .11* | .01 | -.07 | |
| *Stops* | | | | | | | | | | | | | | | | |
| **[d]** | -.12*** | .01 | .07*** | <.01 | <.01 | | .09*** | <.01 | .07* | <.01 | -.13** | .01 | -.15*** | .01 | .08 | |
| **[g]** | -.02 | | .11*** | <.01 | .03 | | .12*** | .01 | .10*** | <.01 | -.09* | <.01 | -.12** | .01 | .12** | .01 |
| [p] | .07** | <.01 | -.05** | <.01 | .03 | | -.08** | <.01 | -.07** | <.01 | .09* | <.01 | <.01 | | -.01 | |
| [t] | -.03 | | .06** | <.01 | <.01 | | -.02 | | -.05 | | .05 | | -.05 | | -.06 | |
| *Af/fricatives* | | | | | | | | | | | | | | | | |
| **[v]** | -.17*** | .01 | .09** | <.01 | -.03 | | .18*** | .01 | -.08* | <.01 | .03 | | .09 | | -.11* | <.01 |
| **[z]** | -.22*** | .02 | .12*** | <.01 | -.02 | | .29*** | .02 | <.01 | | -.11 | | -.01 | | -.09 | |
| **[ʤ]** | -.12*** | <.01 | .17*** | .01 | -.05 | | .28*** | .02 | .01 | | -.06 | | -.05 | | -.02 | |
| [s] | -.01 | | <.01 | | .05 | | .02 | | -.01 | | -.01 | | -.07 | | -.07 | |
| [ʧ] | -.05 | | .22*** | .03 | -.14*** | .01 | .27*** | .04 | .22*** | .03 | -.26*** | .04 | -.22*** | .03 | .10** | <.01 |
| **V1** | | | | | | | | | | | | | | | | |
| *Rounded* | | | | | | | | | | | | | | | | |
| [u] | -.02 | | -.01 | | -.03 | | <.01 | | -.04 | | .02 | | <.01 | | -.06 | |
| [ʊ] | -.05* | <.01 | .04* | <.01 | -.16*** | .01 | .05* | <.01 | -.02 | | -.04 | | -.08 | | .05 | |
| *Unrounded* | | | | | | | | | | | | | | | | |
| [e] | -.60*** | .15 | .06** | <.01 | -.04 | | .09*** | <.01 | .03 | | -.05 | | .12** | .01 | -.08 | |
| [ɛ] | -.60*** | .15 | .04 | | -.04 | | .08** | <.01 | .02 | | -.07 | | .10* | <.01 | -.11** | <.01 |
| [i] | -.60*** | .17 | .11*** | <.01 | .05 | | .17*** | .01 | -.01 | | -.26*** | .03 | .02 | | -.02 | |
| [ɪ] | -.64*** | .18 | .08*** | <.01 | -.07 | | .09*** | <.01 | .02 | | -.03 | | .18*** | .01 | -.13** | <0.1 |
| **C2** | | | | | | | | | | | | | | | | |
| *Sonorants* | | | | | | | | | | | | | | | | |
| **[m]** | -.01 | | -.22*** | .01 | .38*** | .04 | -.23*** | .02 | -.27*** | .02 | .14* | .01 | .18** | .01 | -.30*** | .02 |
| **[n]** | -.05 | | -.18*** | .01 | .28*** | .02 | -.20*** | .01 | -.27*** | .02 | .02 | | .18** | .01 | -.31*** | .03 |
| **[l]** | -.07* | <.01 | -.19*** | .01 | .44*** | .06 | -.21*** | .02 | -.28*** | .03 | .09 | | .18** | .01 | -.37*** | .04 |
| *Stops* | | | | | | | | | | | | | | | | |
| **[d]** | -.08*** | <.01 | .07*** | <.01 | -.03 | | .07** | <.01 | .10*** | .01 | -.13** | .01 | -.10* | .01 | .06 | |
| **[g]** | -.10*** | <.01 | .10*** | <.01 | -.05 | | .14*** | .01 | .15*** | .01 | -.08 | | -.17*** | .02 | .06 | |

| | | | | | | | | | | | | | | | | |
|---|---|---|---|---|---|---|---|---|---|---|---|---|---|---|---|---|
| [p] | -.15*** | .01 | .40*** | .05 | -.03 | | .31*** | .03 | .26*** | .02 | -.17** | .01 | -.10 | | .08 | |
| [t] | -.24*** | .02 | .51*** | .08 | -.16*** | .01 | .35*** | .04 | .36*** | .04 | -.30*** | .03 | -.12* | <.01 | .06 | |
| [k] | -.26*** | .02 | .55*** | .09 | -.05 | | .44*** | .06 | .48*** | .07 | -.35*** | .04 | -.11 | | .12* | <.01 |
| *Af/fricatives* | | | | | | | | | | | | | | | | |
| **[z]** | -.03 | | .08*** | <.01 | .04 | | .13*** | .01 | .08** | <.01 | -.06 | | -.10* | <.01 | .03 | |
| **[ʤ]** | -.04 | | .13*** | .01 | -.07 | | .16*** | .01 | .18*** | .02 | -.02 | | -.17*** | .02 | .18*** | .02 |
| [f] | -.11** | <.01 | .04 | | .06 | | .12*** | <.01 | -.26*** | .02 | .25*** | .02 | .30*** | .03 | -.36*** | .04 |
| [s] | -.15*** | .01 | .09*** | <.01 | .11* | <.01 | .18*** | .01 | -.18*** | .01 | .12* | <.01 | .10 | | -.25*** | .02 |
| [ʧ] | -.20*** | .01 | .31*** | .03 | -.08 | | .39*** | .05 | .06 | | -.03 | | -.04 | | -.07 | |
| **V2** | | | | | | | | | | | | | | | | |
| *Rounded* | | | | | | | | | | | | | | | | |
| [u] | -.07** | <.01 | -.03 | | -.03 | | <.01 | | -.06* | <.01 | .02 | | .12** | .01 | -.08* | <.01 |
| *Unrounded* | | | | | | | | | | | | | | | | |
| [i] | -.09*** | <.01 | .03 | | -.09** | .01 | <-.01 | | -.05* | <.01 | -.04 | | .03 | | -.08* | <.01 |

| | **Arousal** | | | | **Brightness** | | | | **Valence** | | | | **Size** | | | |
|---|---|---|---|---|---|---|---|---|---|---|---|---|---|---|---|---|
| | Calming | | Exciting | | Bright | | Dark | | Good | | Bad | | Small | | Big | |
| **$R^2$ (Adj $R^2$)** | .56 (.53) | | .52 (.49) | | .54 (.51) | | .36 (.32) | | .49 (.46) | | .38 (.34) | | .25 (.2) | | .36 (.32) | |
| **C1** | **β** | **$R^2Δ$** | **β** | **$R^2Δ$** | **β** | **$R^2Δ$** | **β** | **$R^2Δ$** | **β** | **$R^2Δ$** | **β** | **$R^2Δ$** | **β** | **$R^2Δ$** | **β** | **$R^2Δ$** |
| *Sonorants* | | | | | | | | | | | | | | | | |
| **[m]** | .06 | | -.02 | | .02 | | -.02 | | .18*** | .02 | -.08 | | -.09 | | .08 | |
| **[l]** | .09* | <.01 | .13** | .01 | .17*** | .02 | -.08 | | .14** | .01 | -.05 | | -.12* | .01 | .02 | |
| *Stops* | | | | | | | | | | | | | | | | |
| **[d]** | -.15*** | .01 | -.09* | <.01 | -.04 | | .05 | | -.03 | | .07 | | -.03 | | .01 | |
| **[g]** | -.18*** | .02 | .02 | | -.05 | | -.01 | | -.02 | | .07 | | -.08 | | .18*** | .02 |
| [p] | -.02 | | -.07 | | -.08 | | .15** | .01 | -.17*** | .02 | .21*** | .02 | .09 | | -.07 | |
| [t] | -.01 | | -.08 | | -.09* | <.01 | .10* | .01 | -.07 | | .03 | | -.02 | | -.02 | |
| *Af/fricatives* | | | | | | | | | | | | | | | | |
| **[v]** | -.20*** | .01 | .25** | .02 | -.01 | | -.03 | | -.04 | | .21** | .01 | .09 | | -.06 | |
| **[z]** | -.21*** | .01 | .30*** | .03 | -.02 | | -.05 | | .02 | | -.05 | | .01 | | .05 | |
| **[ʤ]** | -.23*** | .02 | .12* | <.01 | .03 | | -.04 | | -.10 | | .13* | <.01 | -.15* | .01 | .11 | |
| [s] | .03 | | .09* | <.01 | .06 | | .01 | | .13** | .01 | -.19*** | .02 | .04 | | .04 | |
| [ʧ] | -.13** | .01 | .04 | | .13** | .01 | <.01 | | -.03 | | -.06 | | -.15** | .01 | .12* | .01 |
| **V1** | | | | | | | | | | | | | | | | |
| *Rounded* | | | | | | | | | | | | | | | | |
| [u] | .03 | | .05 | | -.03 | | <.01 | | .01 | | -.01 | | -.01 | | <.01 | |
| [ʊ] | -.07 | | .07 | | .04 | | -.09 | | -.03 | | .06 | | -.04 | | .05 | |
| *Unrounded* | | | | | | | | | | | | | | | | |
| [e] | -.18*** | .01 | .31*** | .04 | .37*** | .06 | -.38*** | .06 | .11* | <.01 | -.11 | | .06 | | -.13* | .01 |
| [ɛ] | -.18*** | .01 | .30*** | .04 | .32*** | .04 | -.36*** | .05 | .18*** | .01 | <.01 | | .14* | .01 | -.19*** | .02 |

| | | | | | | | | | | | | | | | | |
|---|---|---|---|---|---|---|---|---|---|---|---|---|---|---|---|---|
| [i] | -.19*** | .02 | .39*** | .07 | .42*** | .08 | -.36*** | .06 | .20*** | .02 | -.15** | .01 | -.07 | | -.09 | |
| [ɪ] | -.15*** | .01 | .39*** | .07 | .33*** | .05 | -.38*** | .06 | .15** | .01 | -.04 | | .18** | .01 | -.30*** | .04 |
| **C2** | | | | | | | | | | | | | | | | |
| *Sonorants* | | | | | | | | | | | | | | | | |
| [**m**] | .06 | | -.11 | | -.12* | <.01 | -.03 | | .23*** | .02 | -.17** | .01 | -.03 | | -.13* | <.01 |
| [**n**] | .01 | | -.11 | | -.14* | .01 | .04 | | .14* | .01 | -.12 | | .03 | | -.25*** | .02 |
| [**l**] | .06 | | -.05 | | -.03 | | -.05 | | .19** | .01 | -.10 | | -.04 | | -.20** | .01 |
| *Stops* | | | | | | | | | | | | | | | | |
| [**d**] | -.11** | .01 | -.06 | | .01 | | .04 | | -.03 | | .06 | | -.04 | | .05 | |
| [**g**] | -.23*** | .03 | -.05 | | .01 | | -.01 | | -.13** | .01 | .12* | .01 | -.11* | .01 | .15** | .01 |
| [p] | -.34*** | .04 | .18*** | .01 | .19*** | .01 | -.19** | .01 | <.01 | | .11 | | .01 | | .01 | |
| [t] | -.35*** | .04 | .24*** | .02 | .22*** | .01 | -.28*** | .03 | .11 | | <.01 | | .09 | | .03 | |
| [k] | -.39*** | .04 | .28*** | .02 | .25*** | .02 | -.35*** | .04 | .15* | .01 | .02 | | .08 | | <.01 | |
| *Af/fricatives* | | | | | | | | | | | | | | | | |
| [**z**] | -.07 | | .07 | | .05 | | .01 | | -.05 | | -.01 | | -.16** | .01 | .04 | |
| [**ʤ**] | -.16*** | .01 | .01 | | .02 | | .07 | | -.08 | | .04 | | -.16** | .01 | .14** | .01 |
| [f] | -.12* | <.01 | .07 | | -.03 | | -.14* | .01 | -.10 | | .27*** | .02 | .01 | | -.06 | |
| [s] | -.17** | .01 | .12* | <.01 | .06 | | -.14* | .01 | -.06 | | .20** | .01 | -.11 | | -.03 | |
| [ʧ] | -.34*** | .03 | .13* | <.01 | .04 | | -.10 | | -.11 | | .20** | .01 | -.12 | | .06 | |
| **V2** | | | | | | | | | | | | | | | | |
| *Rounded* | | | | | | | | | | | | | | | | |
| [u] | .04 | | .01 | | -.07 | | .02 | | -.10* | .01 | .08 | | .11* | .01 | -.12** | .01 |
| *Unrounded* | | | | | | | | | | | | | | | | |
| [i] | .05 | | -.01 | | .13*** | .01 | .07 | | -.06 | | -.02 | | -.01 | | -.13** | .01 |

**Supplementary Table 8:** Whole-item acoustic regression: multicollinearity statistics; VIF = variance inflation factor.

| Predictor | Tolerance | VIF |
|---|---|---|
| FFT CV | .71 | 1.41 |
| SE CV | .31 | 3.27 |
| ST CV | .78 | 1.28 |
| FUF | .26 | 3.84 |
| Jitter | .36 | 2.81 |
| Shimmer | .51 | 1.97 |
| MAC | .23 | 4.33 |
| Mean F0 | .80 | 1.25 |
| $F0_{SD}$ | .69 | 1.44 |
| Duration | .72 | 1.38 |

**Supplementary Table 9:** Whole-item acoustic regression: Durbin-Watson test values.

| Domain | Scale | Test value |
|---|---|---|
| Shape | Rounded | 1.93 |
| | Pointed | 2.08 |
| Roughness | Smooth | 2.02 |
| | Rough | 2.11 |
| Hardness | Hard | 1.92 |
| | Soft | 1.87 |
| Weight | Light | 1.96 |
| | Heavy | 2.00 |
| Arousal | Calming | 1.85 |
| | Exciting | 1.90 |
| Brightness | Bright | 1.93 |
| | Dark | 2.08 |
| Valence | Good | 2.02 |
| | Bad | 2.07 |
| Size | Small | 2.00 |
| | Big | 2.13 |

**Supplementary Figures 3a-h:** Homoscedasticity assessment for the whole-item acoustic regression: histograms (top row) and Q-Q plots (middle row) for standardized residuals and scatterplots (bottom row) for standardized residuals against standardized predicted values.

Supplementary Figure 3a: Shape – rounded (left & pointed (right)

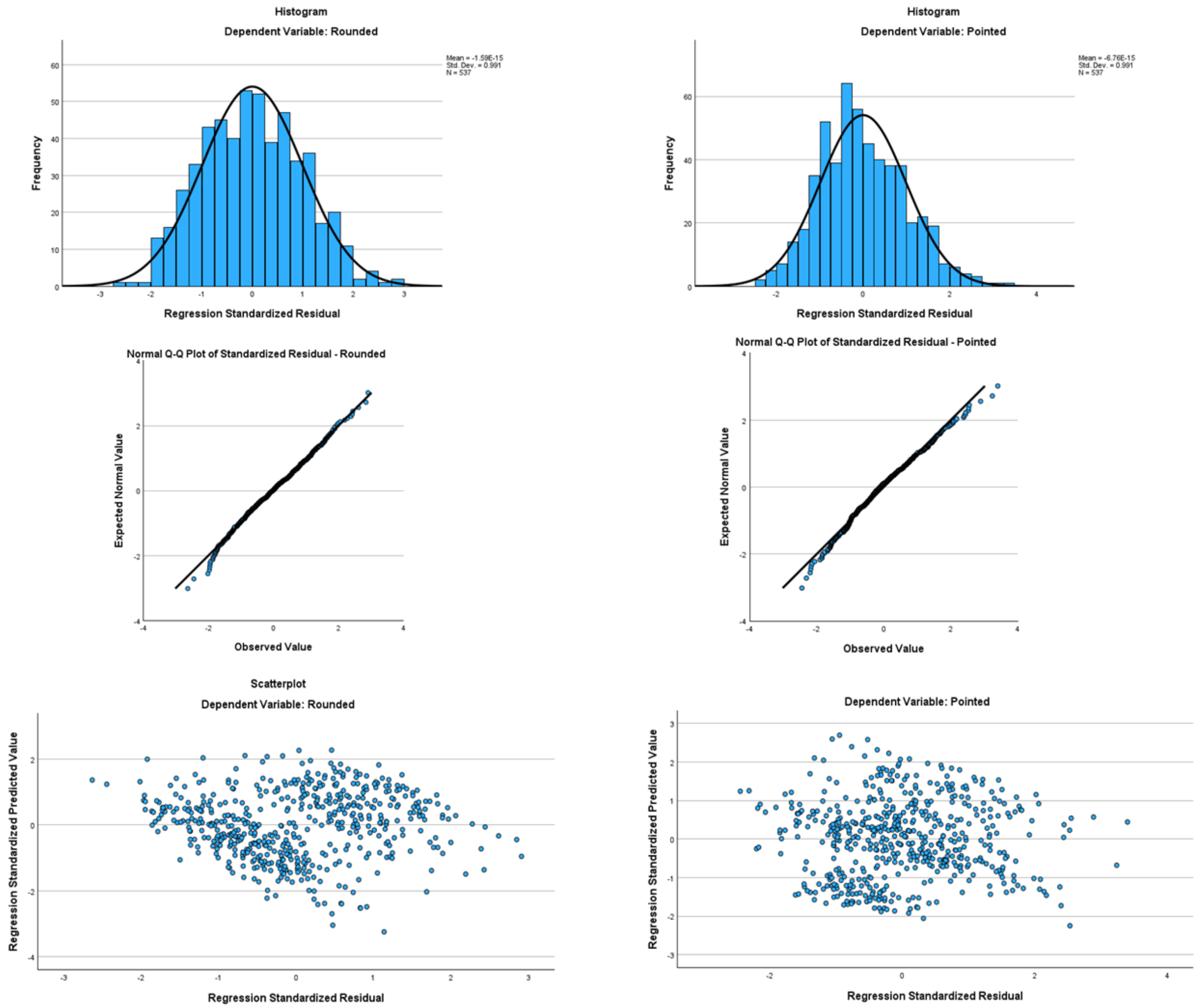

Supplementary Figure 3b: Roughness – smooth (left) & rough (right)

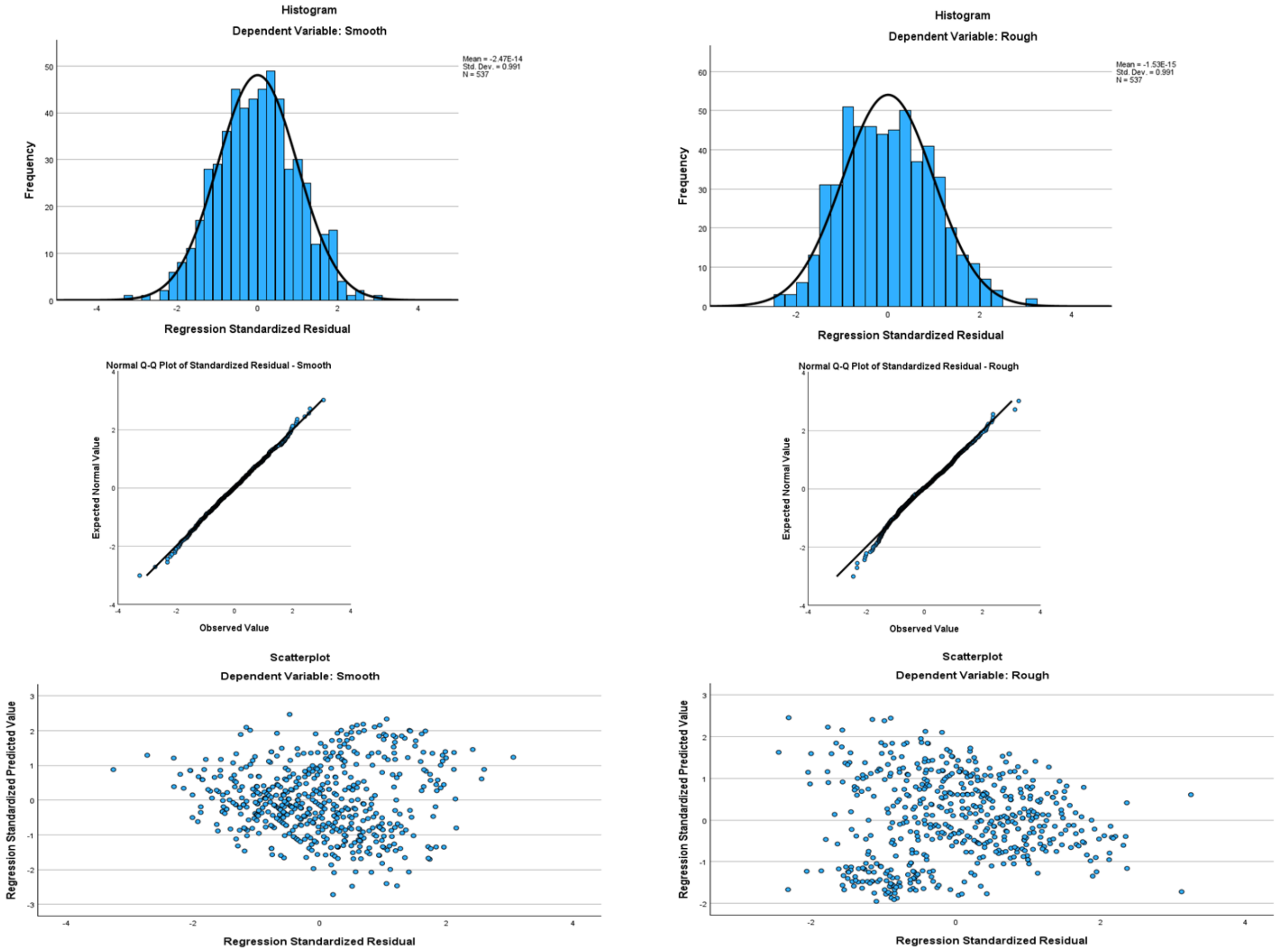

Supplementary Figure 3c: Hardness – hard (left) & soft (right)

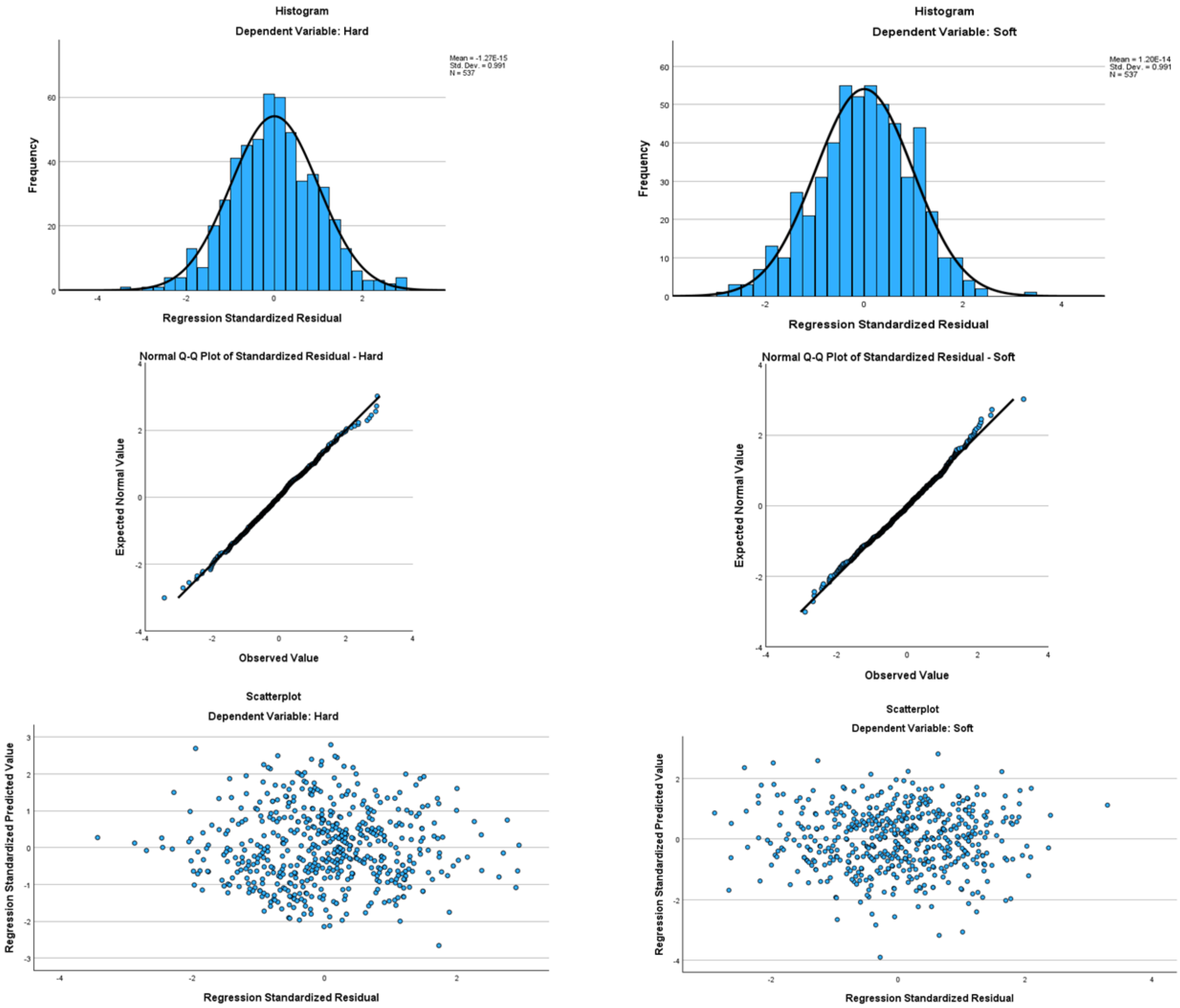

### Supplementary Figure 3d: Weight – light (left) & heavy (right)

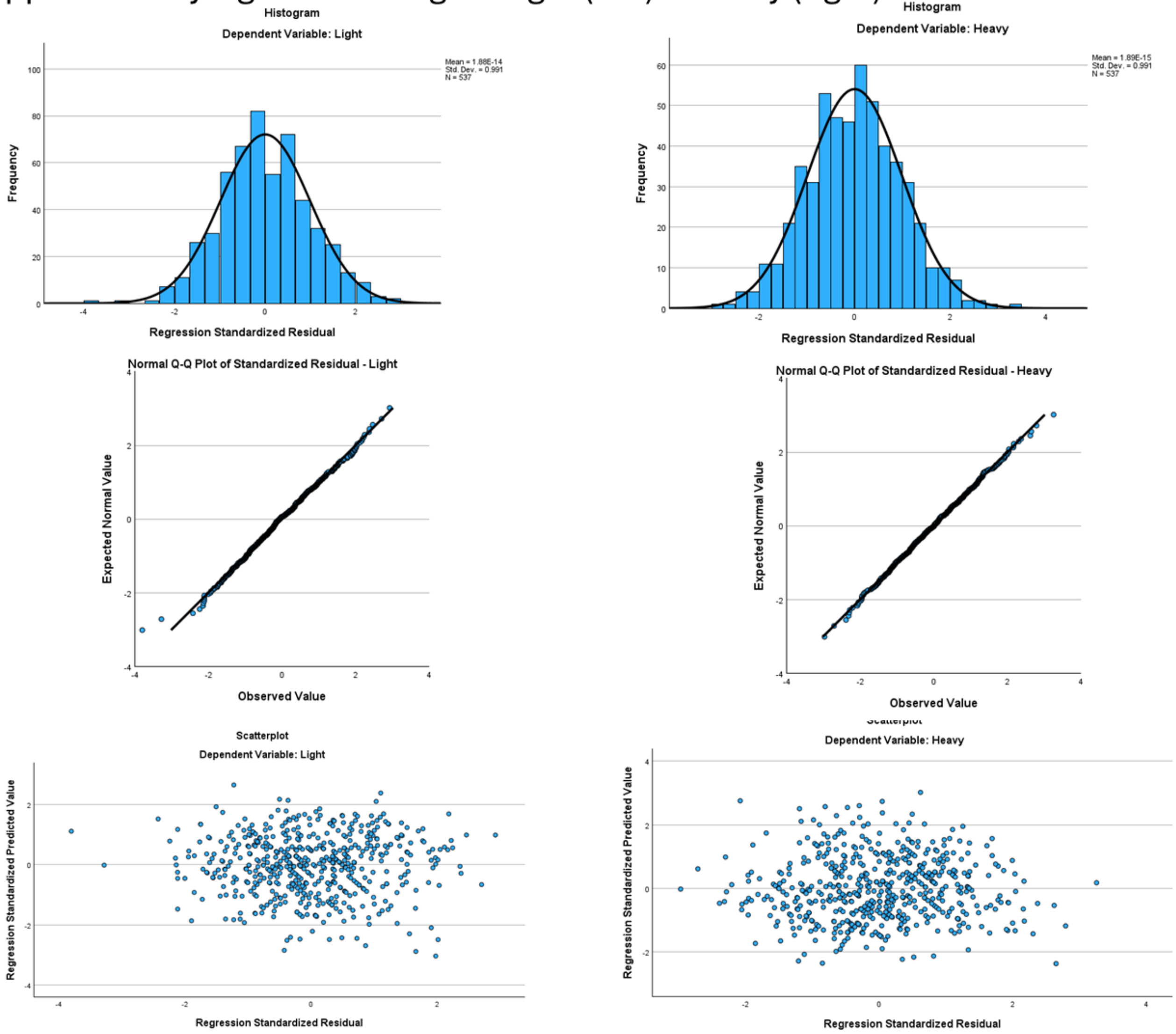

Supplementary Figure 3e: Arousal – calming (left) & exciting (right)

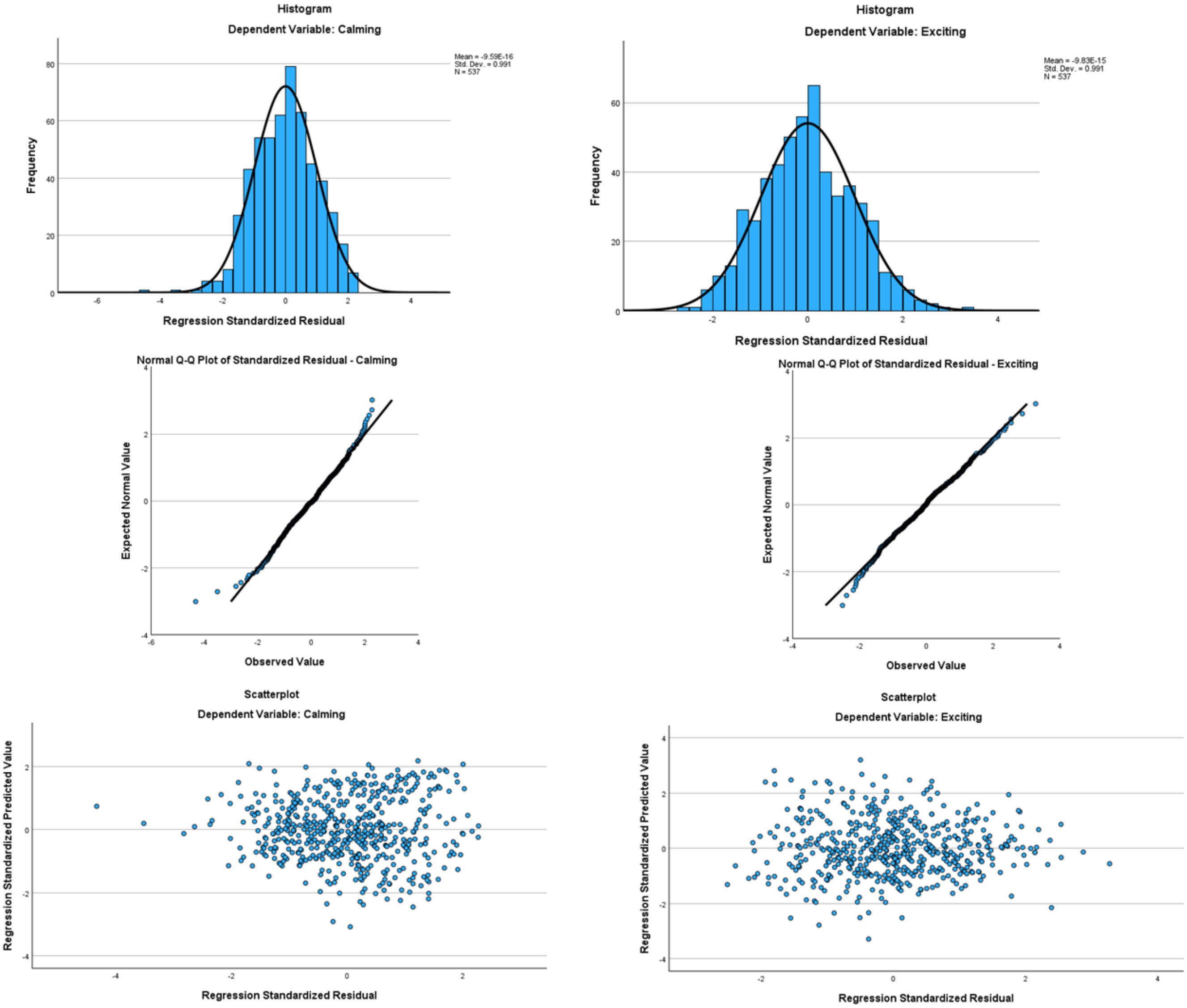

### Supplementary Figure 3f: Brightness – bright (left) & dark (right)

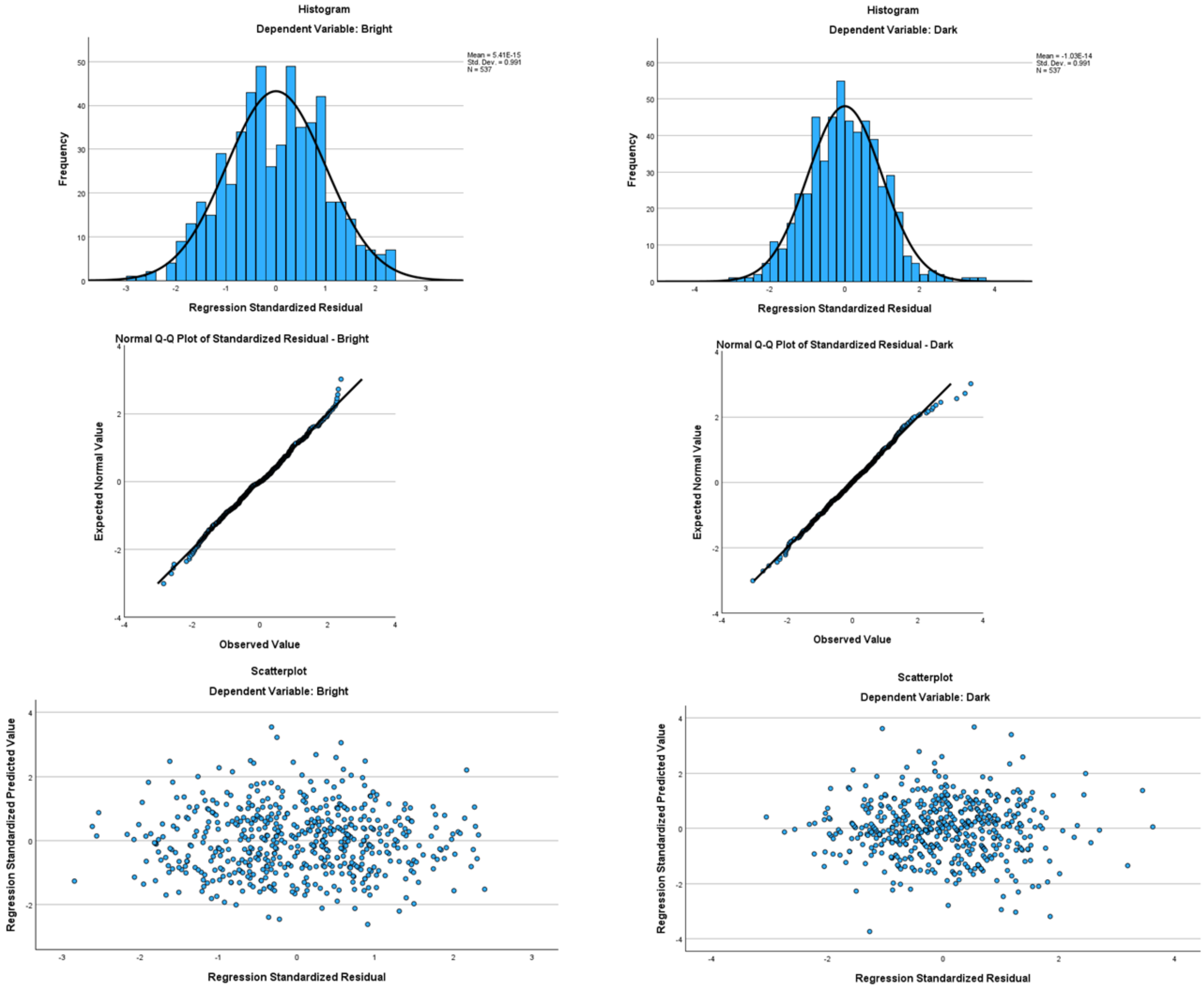

Supplementary Figure 3g: Valence – good (left) & bad (right)

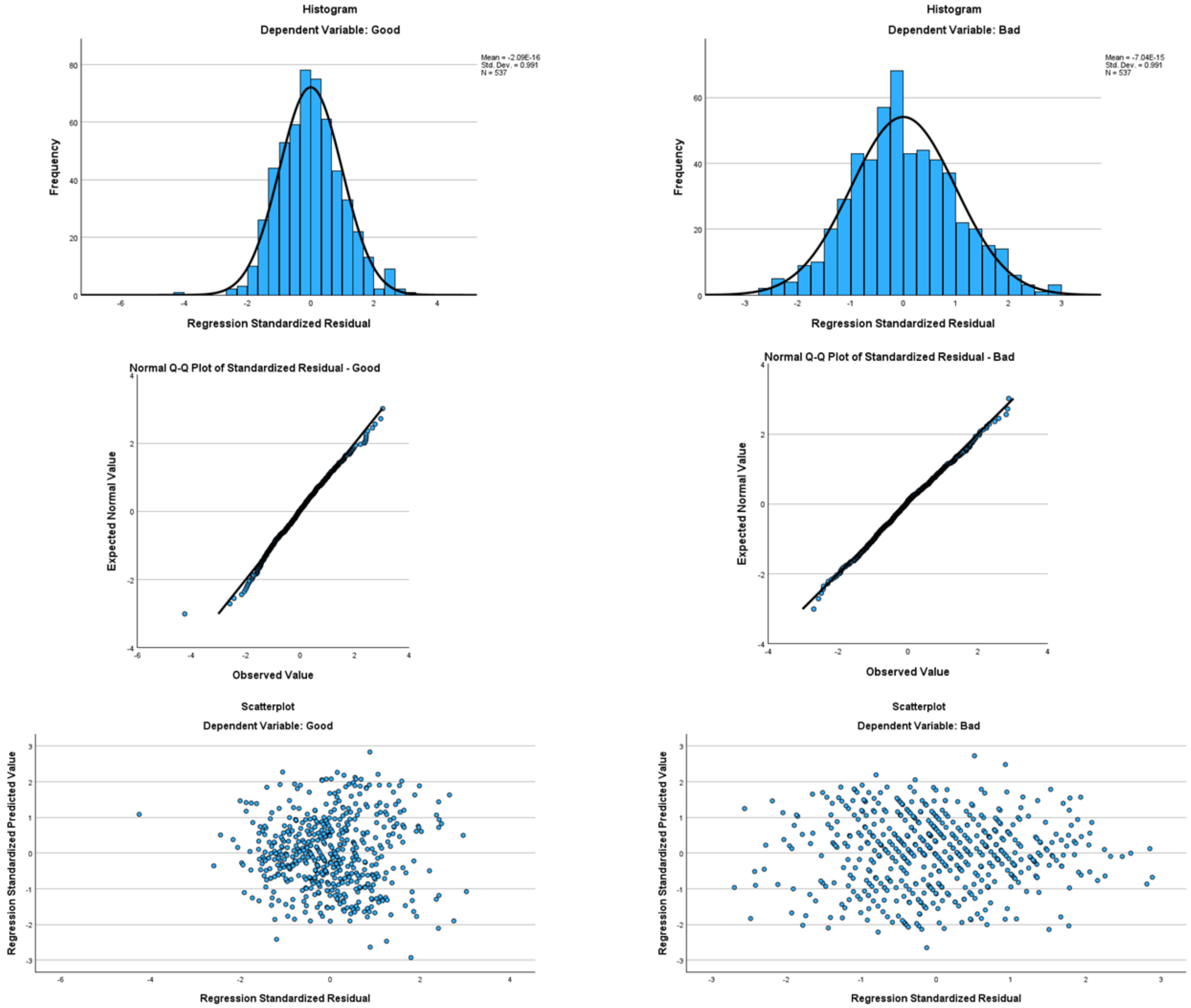

Supplementary Figure 3h: Size – small (left) & big (right)

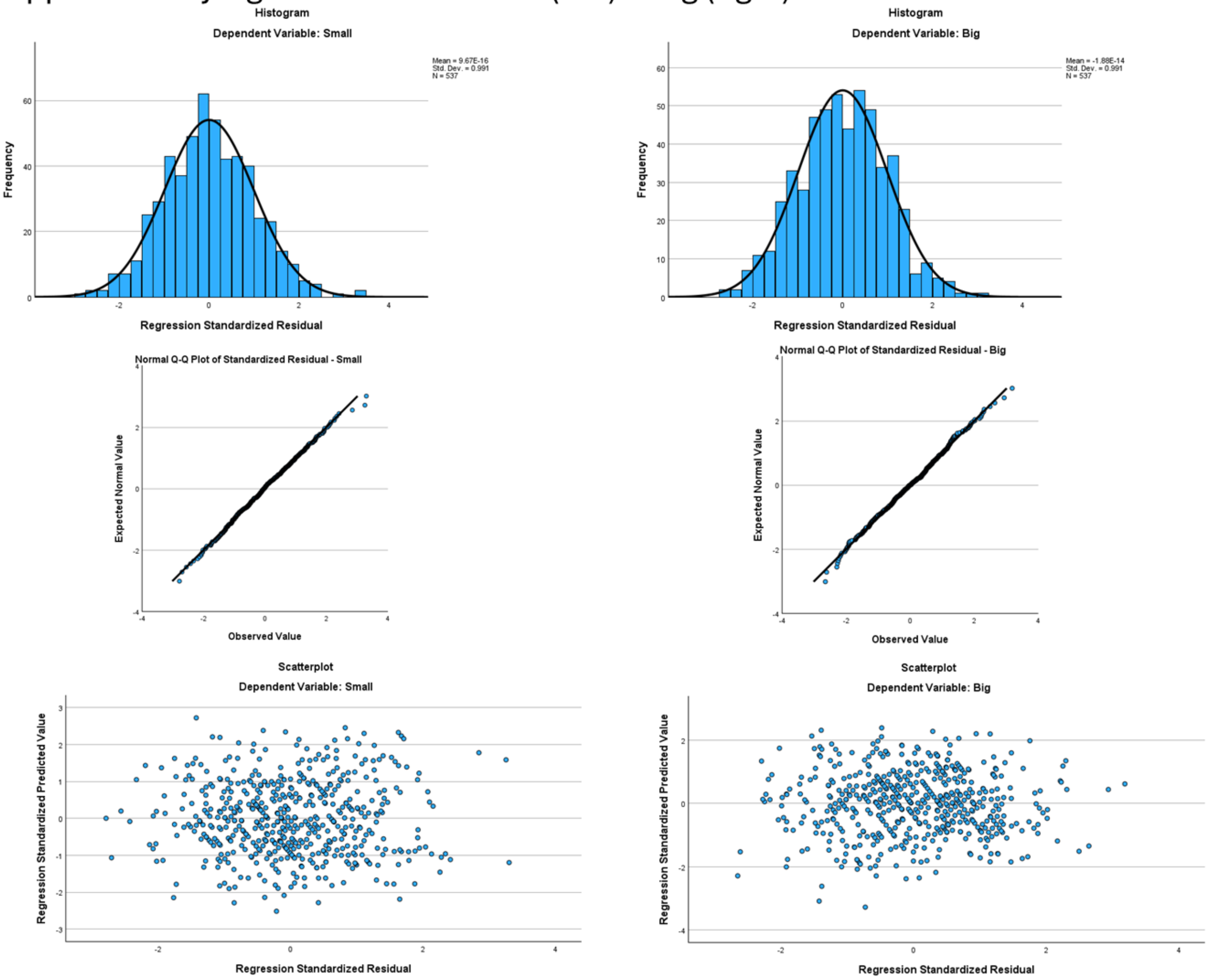

**Supplementary Table 10:** Whole-item acoustic regression analysis for each domain. Coefficient of variation (CV) for fast Fourier transform (FFT), speech envelope (SE), spectral tilt (ST); fraction of unvoiced frames (FUF), jitter, shimmer, mean autocorrelation (MAC), mean fundamental frequency (F0) and its standard deviation ($F0_{SD}$), duration. $R^2$, Adj $R^2$, $\beta$, and $R^2\Delta$ as for Table 3; p-values as for Supplementary Table 4.

| | **Shape** | | | | **Roughness** | | | | **Hardness** | | | | **Weight** | | | |
|---|---|---|---|---|---|---|---|---|---|---|---|---|---|---|---|---|
| | Rounded | | Pointed | | Smooth | | Rough | | Hard | | Soft | | Light | | Heavy | |
| **$R^2$ (Adj $R^2$)** | .3 (.28) | | .56 (.55) | | .35 (.34) | | .42 (.41) | | .45 (.43) | | .18 (.16) | | .24 (.23) | | .26 (.25) | |
| | **$\beta$** | **$R^2\Delta$** | **$\beta$** | **$R^2\Delta$** | **$\beta$** | **$R^2\Delta$** | **$\beta$** | **$R^2\Delta$** | **$\beta$** | **$R^2\Delta$** | **$\beta$** | **$R^2\Delta$** | **$\beta$** | **$R^2\Delta$** | **$\beta$** | **$R^2\Delta$** |
| FFT CV | .34*** | .08 | .27*** | .05 | -.15*** | .02 | .08* | .13 | .42*** | .13 | -.22*** | .03 | -.37*** | .10 | .46*** | .15 |
| SE CV | -.09 | | .08 | | -.16* | .01 | -.05 | | .09 | | .24*** | .02 | .24*** | .02 | -.13 | |
| ST CV | -.17*** | .02 | .01 | | .05 | | -.05 | | -.02 | | <-.01 | | .09* | .01 | <-.01 | |
| FUF | -.06 | | .63*** | .10 | -.22** | .01 | .52*** | .10 | .43* | .05 | -.45*** | .05 | -.41*** | .04 | .25*** | .02 |
| Jitter | .19** | .01 | -.09 | | -.03 | | -.17** | .01 | -.18** | .01 | .02 | | .03 | | -.12 | |
| Shimmer | .08 | | .16*** | .10 | -.15** | .01 | .15*** | .01 | .08 | | <.01 | | -.01 | | <-.01 | |
| MAC | .32*** | .02 | -.05 | | .13 | | - .27*** | .02 | -.22** | .01 | .16 | | .22** | .01 | -.34*** | .03 |
| Mean F0 | -.15*** | .02 | .10** | .01 | .08* | .01 | .01 | | .02 | | <-.01 | | .13** | .01 | -.09* | .01 |
| $F0_{SD}$ | .14** | .01 | .08* | <.01 | -.04 | | -.03 | | -.04 | | .06 | | .02 | | -.05 | |
| Duration | .22*** | .04 | -.21*** | .03 | .11** | .01 | -.03 | | -.26*** | .08 | .26*** | .05 | .11* | .01 | -.13** | .01 |

| | **Arousal** | | | | **Brightness** | | | | **Valence** | | | | **Size** | | | |
|---|---|---|---|---|---|---|---|---|---|---|---|---|---|---|---|---|
| | Calming | | Exciting | | Bright | | Dark | | Good | | Bad | | Small | | Big | |
| **$R^2$ (Adj $R^2$)** | .24 (.23) | | .17 (.15) | | .23 (.21) | | .13 (.11) | | .18 (.16) | | .16 (.14) | | .19 (.17) | | .12 (.1) | |
| | **$\beta$** | **$R^2\Delta$** | **$\beta$** | **$R^2\Delta$** | **$\beta$** | **$R^2\Delta$** | **$\beta$** | **$R^2\Delta$** | **$\beta$** | **$R^2\Delta$** | **$\beta$** | **$R^2\Delta$** | **$\beta$** | **$R^2\Delta$** | **$\beta$** | **$R^2\Delta$** |
| FFT CV | -.14** | .01 | -.13** | .01 | -.15** | .02 | .07 | | -.09* | .01 | .11* | .01 | .04 | | .24*** | .04 |
| SE CV | -.01 | | .07 | | -.16* | .01 | -.04 | | -.08 | | .16* | .01 | .64*** | .12 | -.34*** | .04 |
| ST CV | -.02 | | .15** | .02 | .23*** | .04 | -.20*** | .03 | .09* | .01 | -.07 | | .11* | .01 | -.14** | .02 |
| FUF | -.42*** | .05 | .14 | | .36*** | .03 | -.18* | .01 | -.09 | | .14 | | -.29*** | .02 | .28*** | .02 |
| Jitter | .18** | .01 | -.21** | .02 | -.15* | .01 | .17* | .01 | .01 | | -.04 | | .04 | | -.14* | .01 |
| Shimmer | -.09 | | .19*** | .02 | .11* | .01 | -.09 | | -.13* | .01 | .11 | | .09 | | -.05 | |
| MAC | .23** | .01 | -.09 | | -.13 | | -.03 | | .12 | | -.07 | | .17* | .01 | -.29*** | .02 |
| Mean F0 | -.04 | | .22*** | .04 | .21*** | .04 | -.24*** | .05 | .11* | .01 | -.05 | | .09* | .01 | -.06 | |
| $F0_{SD}$ | .08 | | -.05 | | .04 | | -.09 | | -.02 | | <.01 | | .01 | | <-.01 | |
| Duration | .17*** | .02 | .07 | | -.18*** | .02 | .06 | | -.01 | | -.04 | | -.10* | .01 | .10* | .01 |

**Supplementary Table 11:** Segmental acoustic regression: multicollinearity statistics; VIF = variance inflation factor.

| **Predictor** | **Tolerance** | **VIF** |
|---|---|---|
| **C1** | | |
| F0 | .85 | 1.18 |
| Intensity | .44 | 2.28 |
| Duration | .42 | 2.36 |
| **V1** | | |
| F0 | .91 | 1.10 |
| F1 | .61 | 1.63 |
| F2 | .17 | 5.88 |
| F3 | .72 | 1.39 |
| Intensity | .94 | 1.07 |
| Duration | .48 | 2.10 |
| C2 | | |
| F0 | .86 | 1.16 |
| Intensity | .67 | 1.50 |
| Duration | .38 | 2.63 |
| **V2** | | |
| F0 | .98 | 1.02 |
| F1 | .71 | 1.41 |
| F2 | .14 | 7.26 |
| F3 | .55 | 1.82 |
| Intensity | .74 | 1.36 |
| Duration | .74 | 1.36 |

**Supplementary Table 12:** Segmental acoustic regression: Durbin-Watson test values.

| Domain | Scale | Test value |
|---|---|---|
| Shape | Rounded | 2.11 |
| | Pointed | 2.02 |
| Roughness | Smooth | 1.91 |
| | Rough | 2.14 |
| Hardness | Hard | 1.84 |
| | Soft | 1.85 |
| Weight | Light | 1.96 |
| | Heavy | 1.85 |
| Arousal | Calming | 1.93 |
| | Exciting | 1.91 |
| Brightness | Bright | 2.10 |
| | Dark | 2.04 |
| Valence | Good | 1.91 |
| | Bad | 1.98 |
| Size | Small | 1.96 |
| | Big | 1.96 |

**Supplementary Figures 4a-h:** Homoscedasticity assessment for the segmental acoustic regression: histograms (top row) and Q-Q plots (middle row) for standardized residuals and scatterplots (bottom row) for standardized residuals against standardized predicted values.

Supplementary Figure 4a: Shape – rounded (left) & pointed (right)

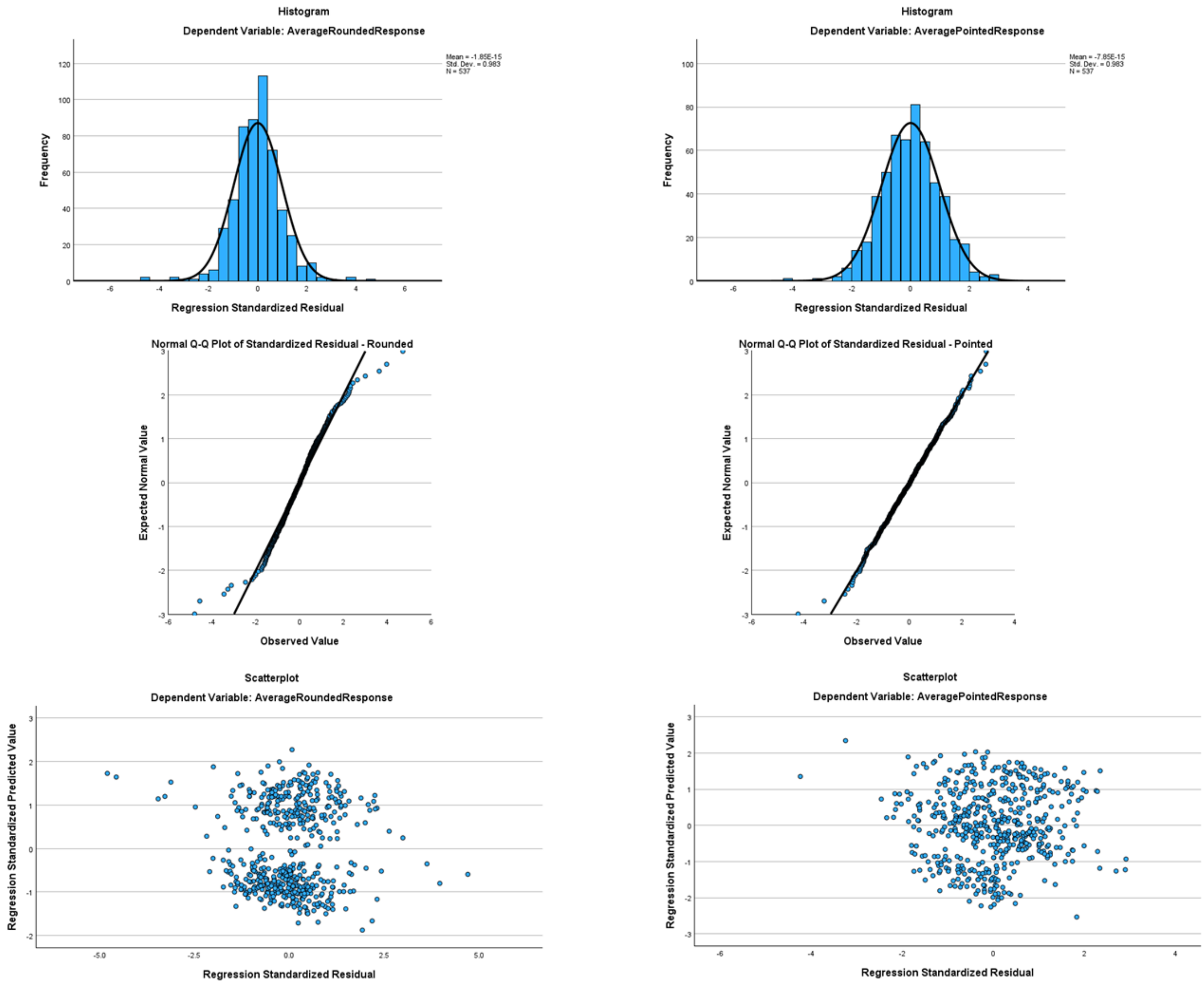

Supplementary Figure 4b: Roughness – smooth (left) & rough (right)

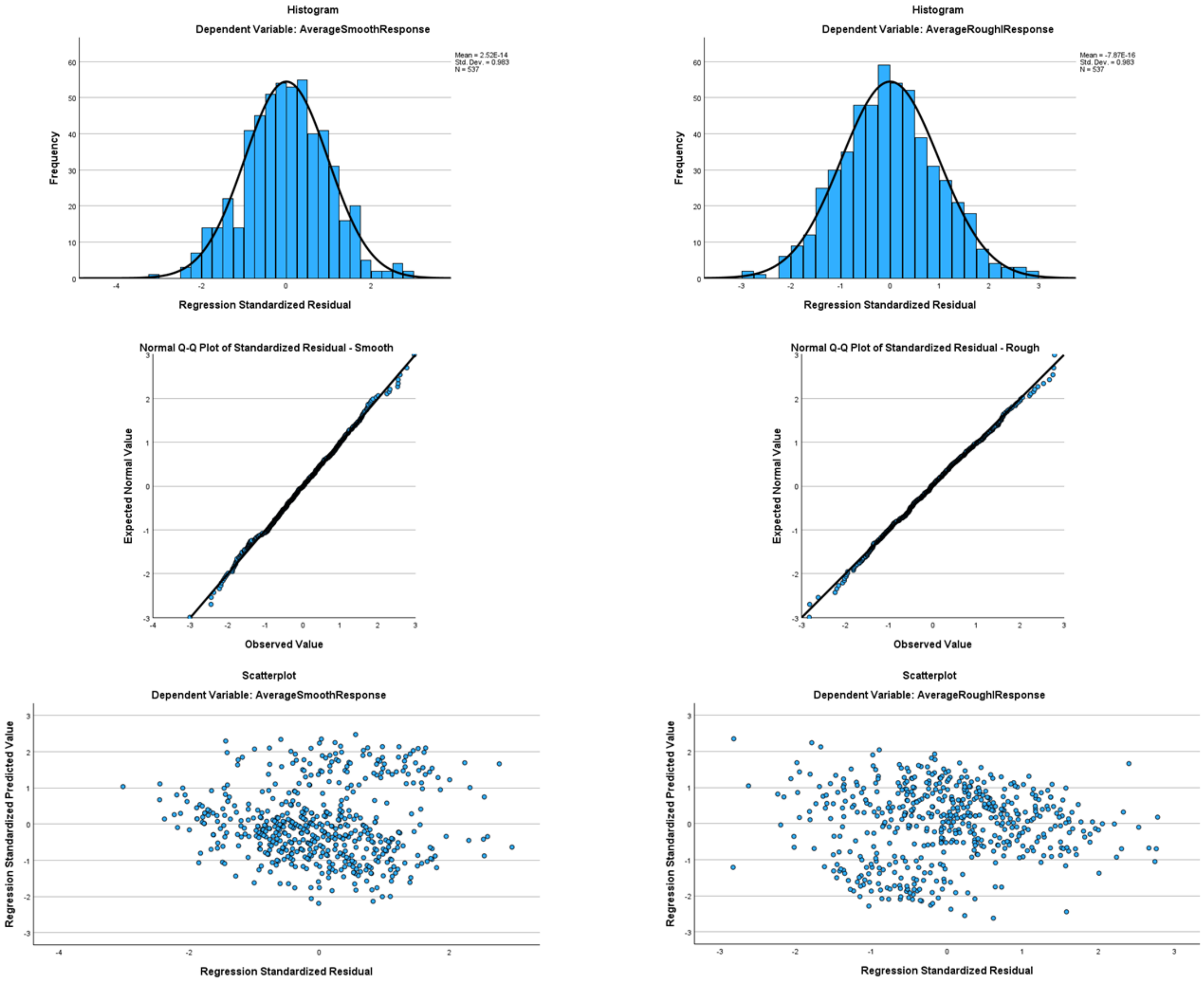

Supplementary Figure 4c: Hardness – hard (left) & soft (right)

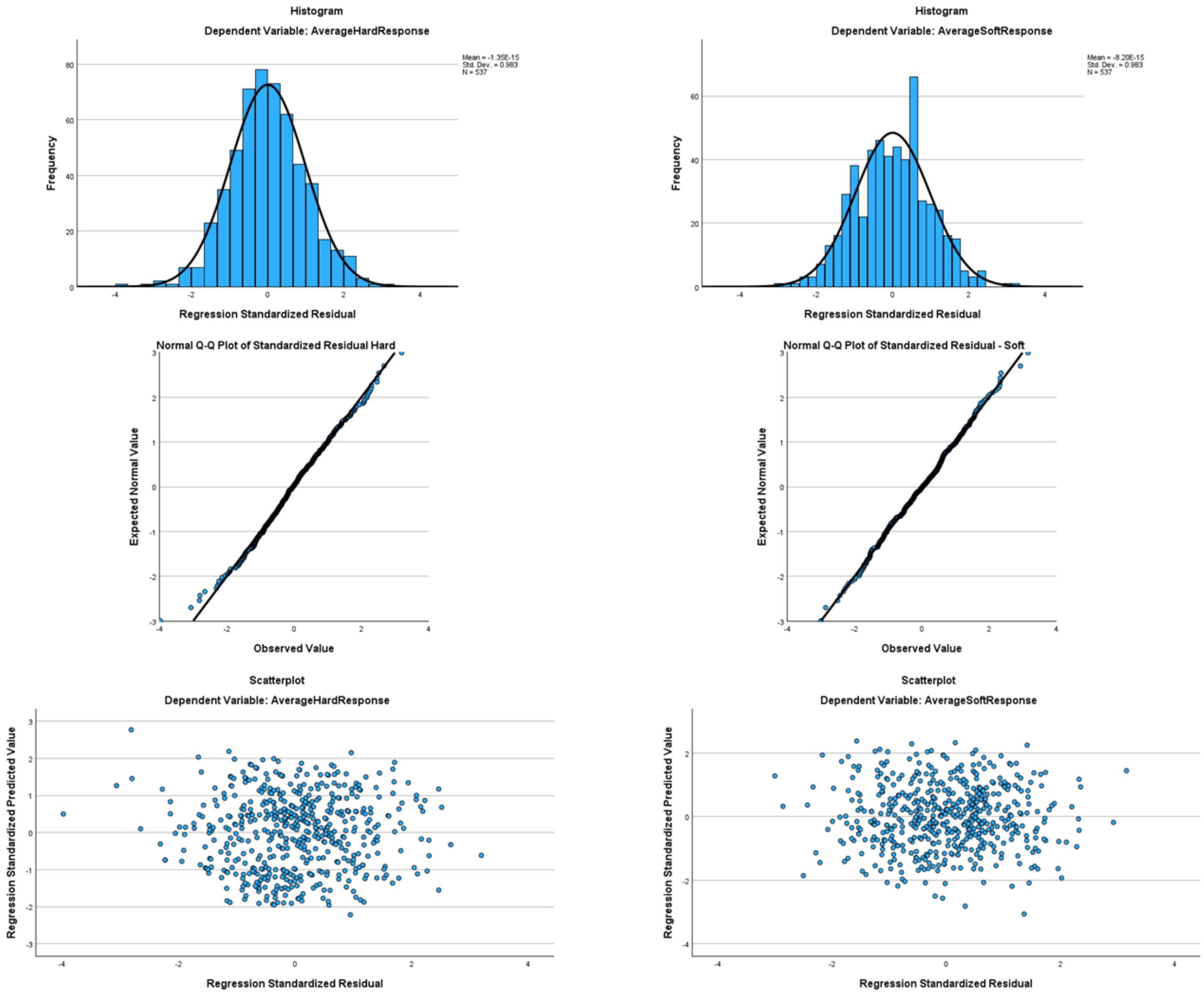

### Supplementary Figure 4d: Weight – light (left) & heavy (right)

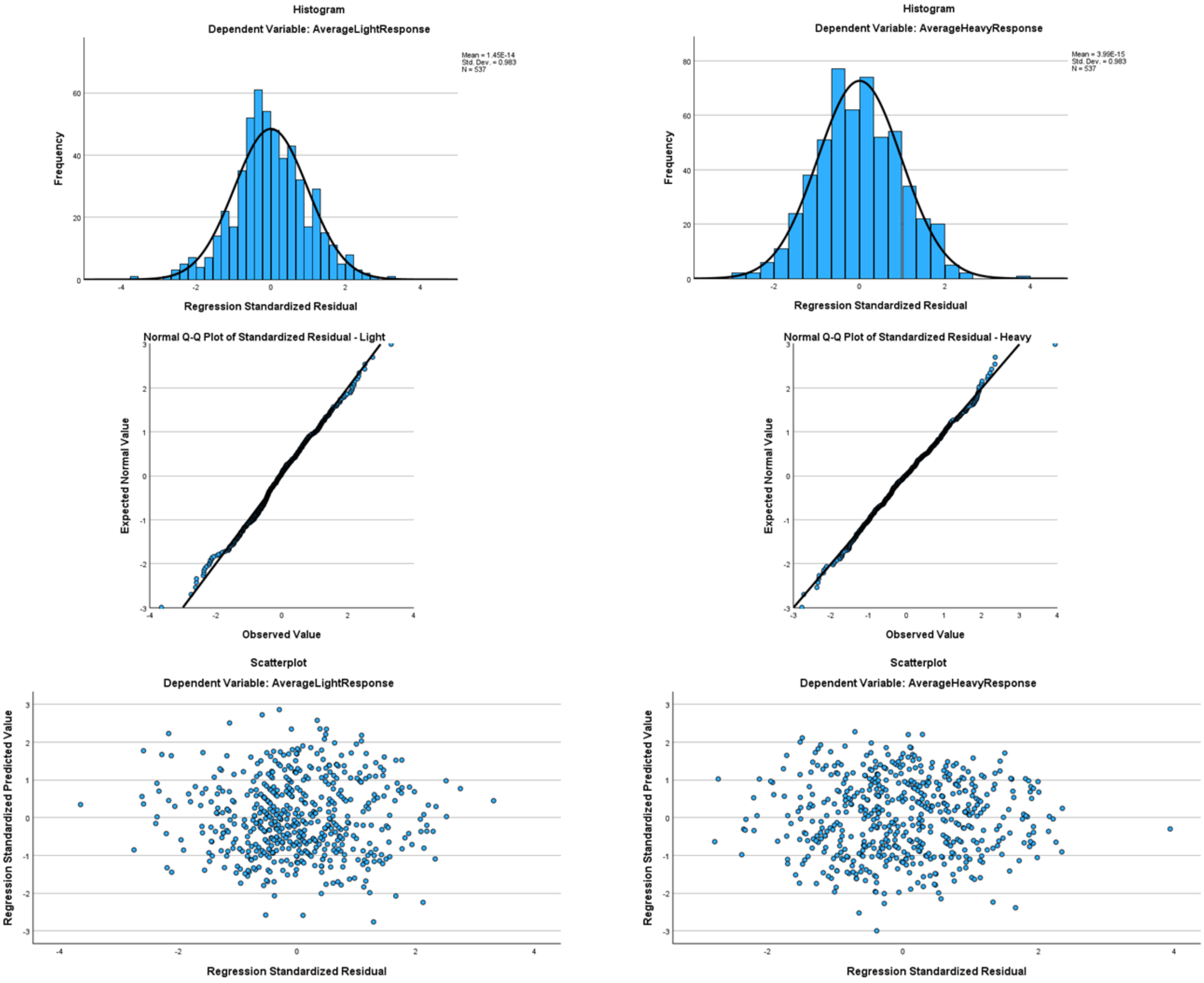

### Supplementary Figure 4e: Arousal – calming (left) & exciting (right)

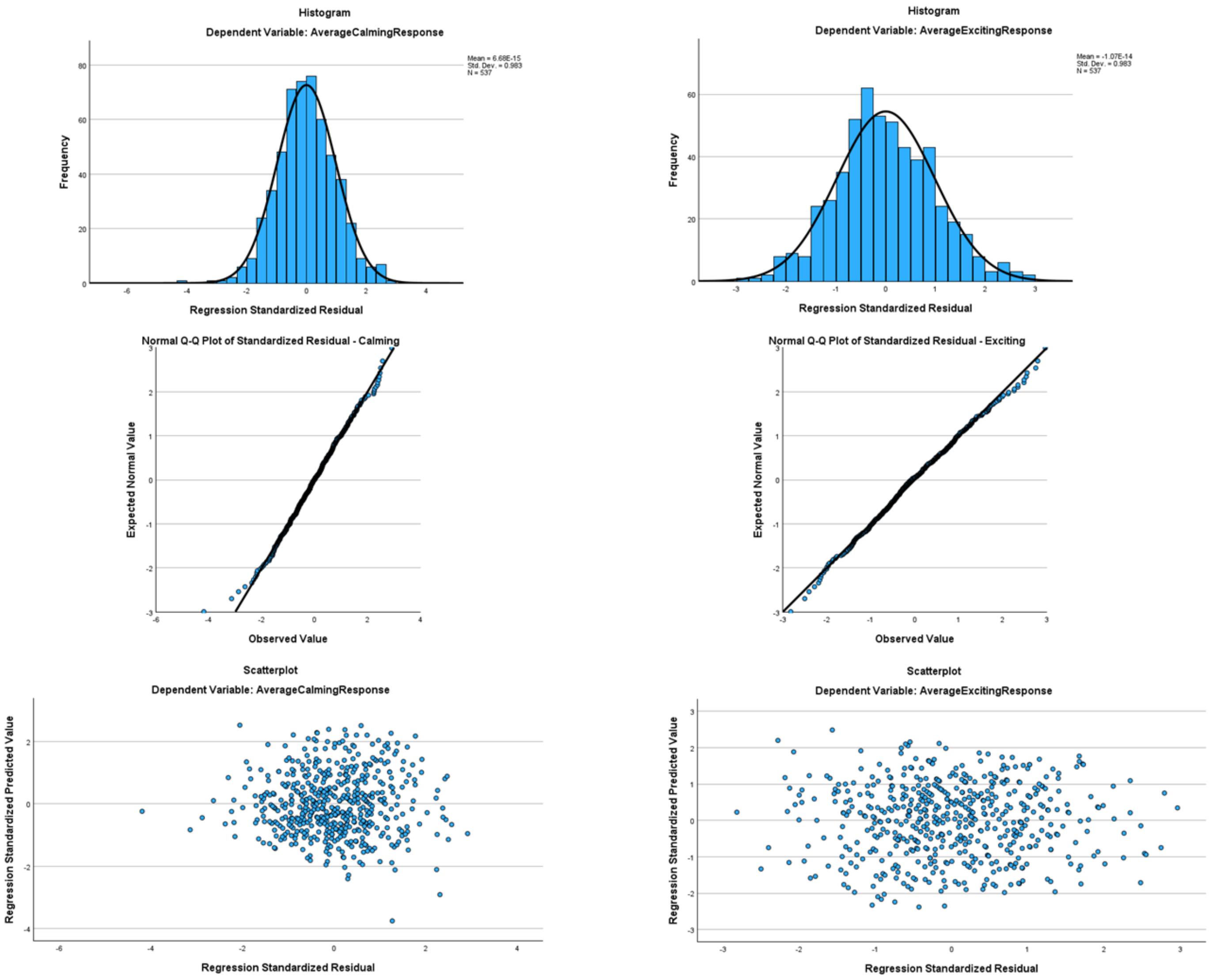

Supplementary Figure 4f: Brightness – bright (left) & dark (right)

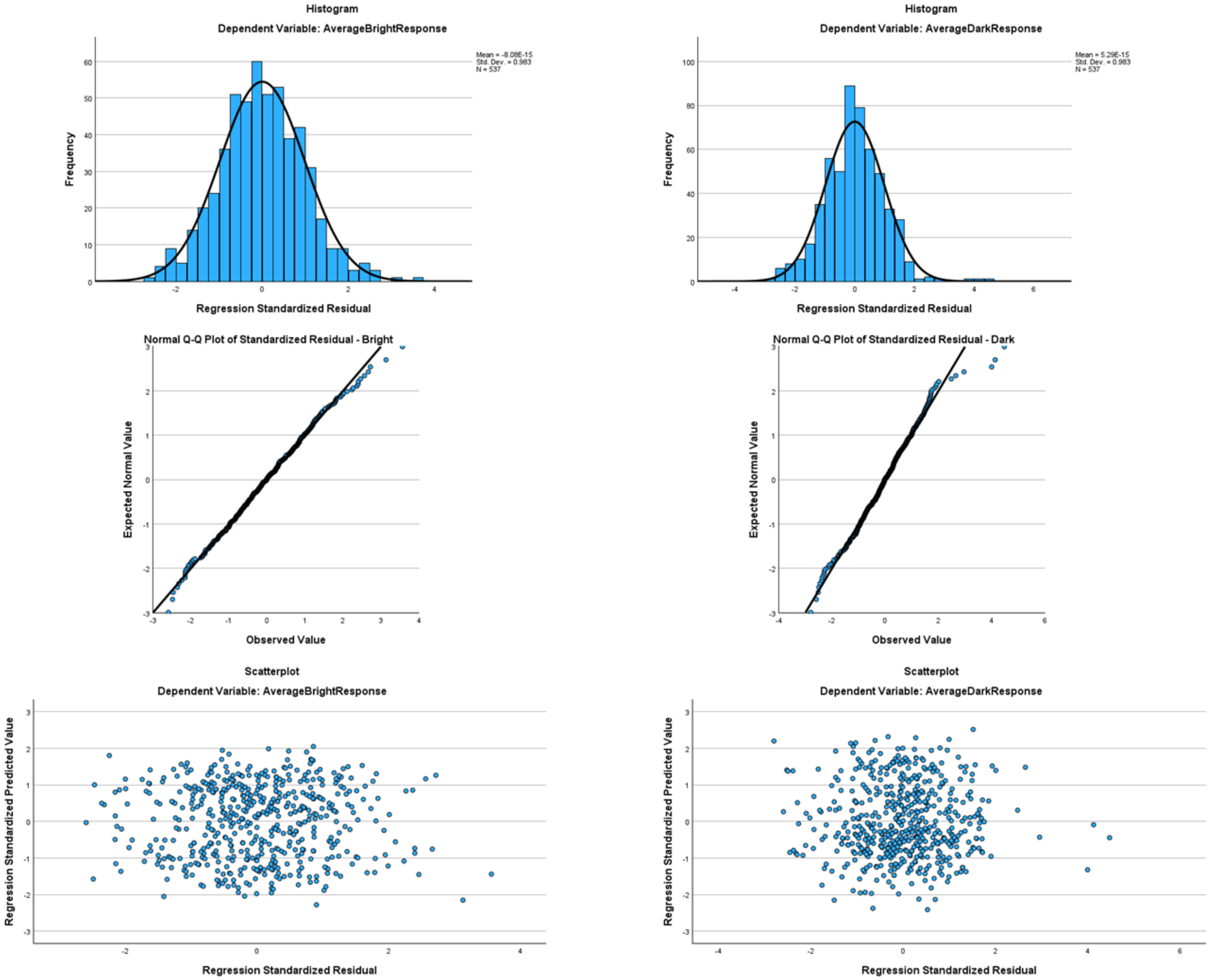

### Supplementary Figure 4g: Valence – good (left) & bad (right)

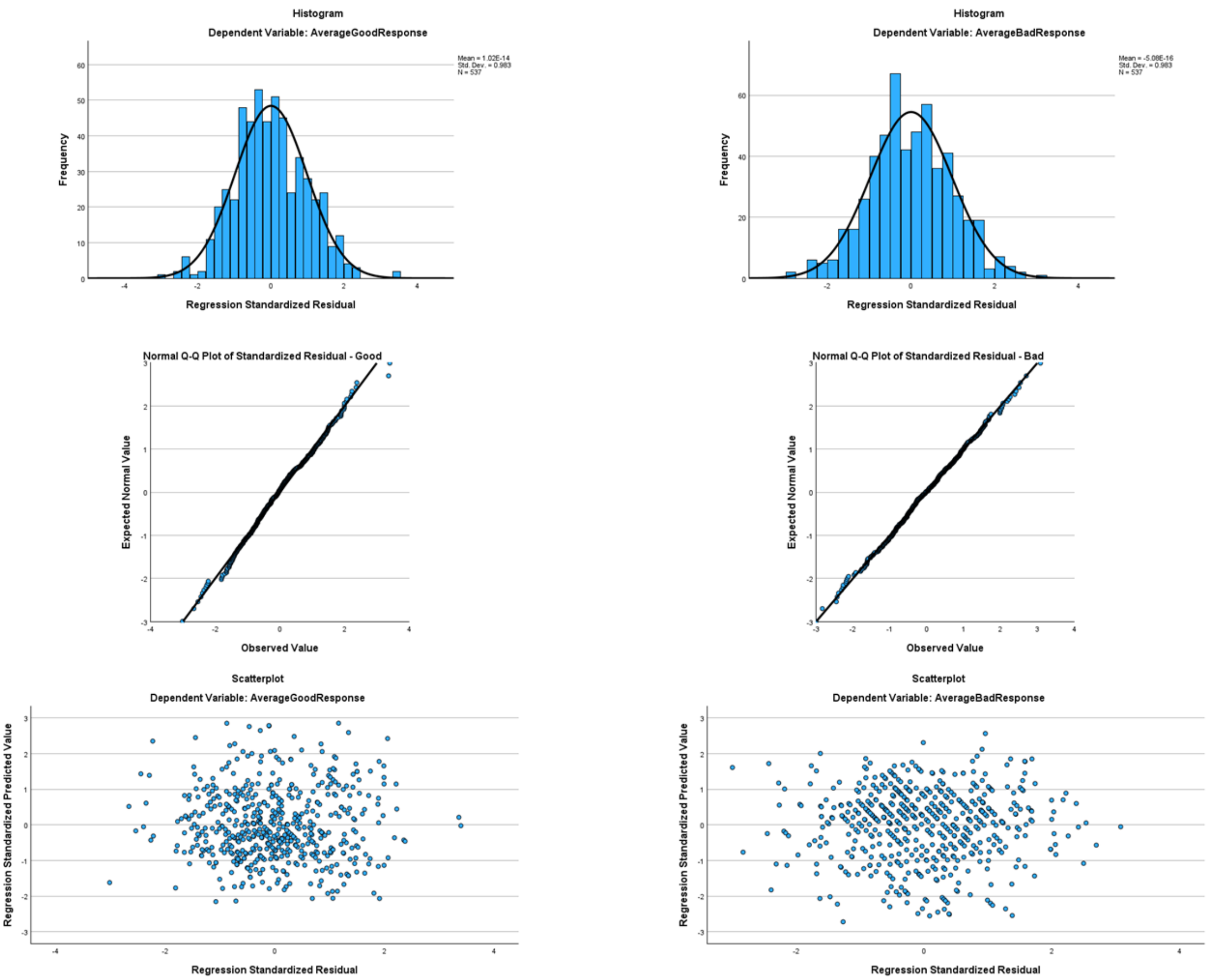

Supplementary Figure 4g: Size – small (left) & big (right)

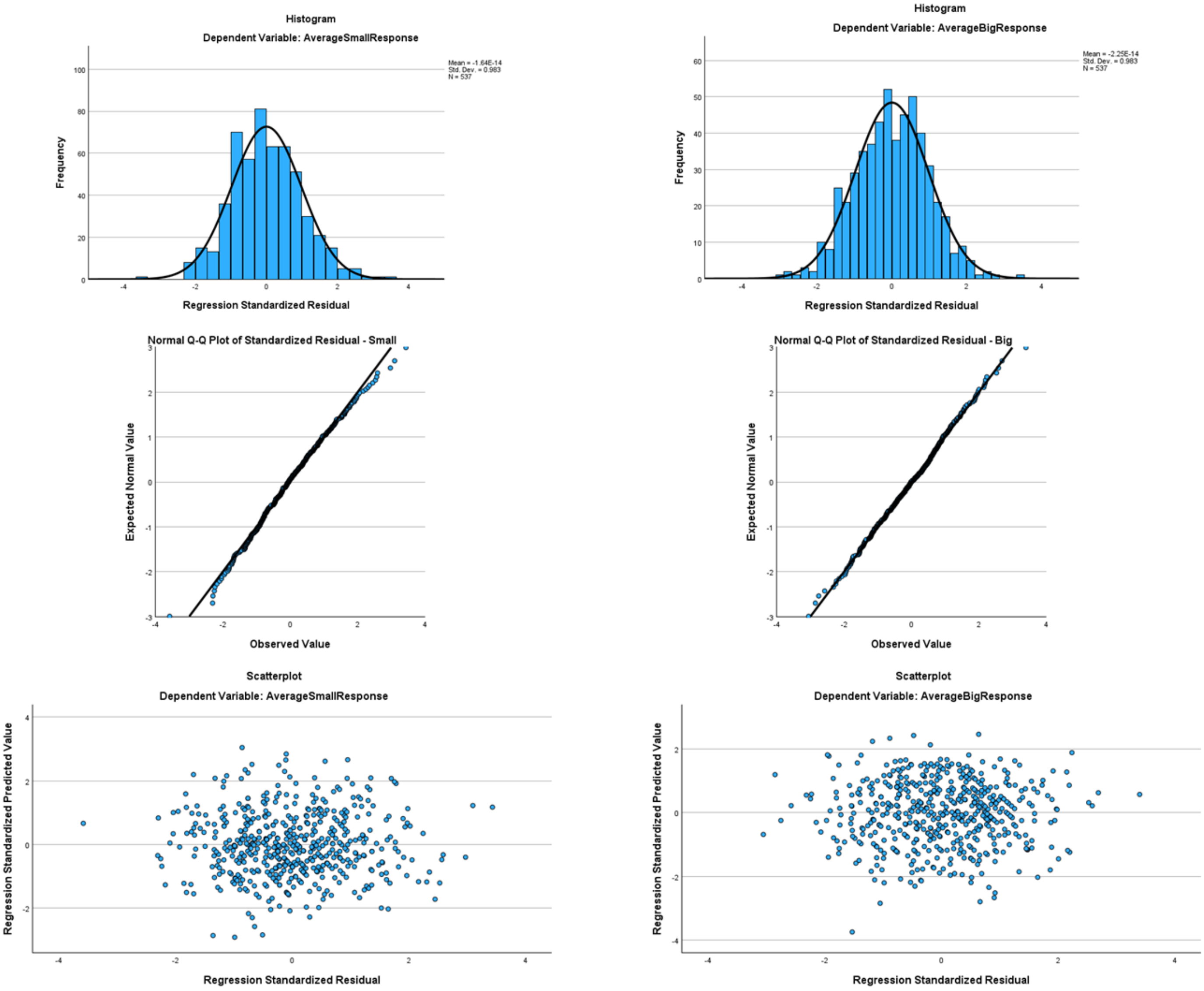

**Supplementary Table 13:** Segmental acoustic regression analysis for each domain. [F0, F1, F2, F3, intensity, duration]. $R^2$, Adj $R^2$, β, and $R^2\Delta$ as for Table 3; C1/2, V1/2 as for Table 4; p-values as for Supplementary Table 4.

| | **Shape** | | | | **Roughness** | | | | **Hardness** | | | | **Weight** | | | |
|---|---|---|---|---|---|---|---|---|---|---|---|---|---|---|---|---|
| | Rounded | | Pointed | | Smooth | | Rough | | Hard | | Soft | | Light | | Heavy | |
| **$R^2$ (Adj $R^2$)** | .77 (.76) | | .66 (.65) | | .43 (.41) | | .52 (.51) | | .54 (.52) | | .29 (.26) | | .31 (.29) | | .40 (.38) | |
| **C1** | **β** | **$R^2\Delta$** | **β** | **$R^2\Delta$** | **β** | **$R^2\Delta$** | **β** | **$R^2\Delta$** | **β** | **$R^2\Delta$** | **β** | **$R^2\Delta$** | **β** | **$R^2\Delta$** | **β** | **$R^2\Delta$** |
| F0 | .01 | | -.02 | | .16*** | .02 | -.08* | .01 | -.02 | | <.01 | | .08* | <.01 | <.01 | |
| Intensity | .01 | | .02 | | .11* | .01 | -.01 | | -.02 | | -.06 | | .07 | | -.16** | .01 |
| Duration | -.08* | <.01 | -.16*** | .01 | .04 | | -.01 | | -.34*** | .05 | .28*** | .03 | .20*** | .02 | -.25*** | .03 |
| **V1** | | | | | | | | | | | | | | | | |
| F0 | < .01 | | .02 | | -.01 | <.01 | -.06 | | <.01 | | .05 | | .04 | | -.07* | <.01 |
| F1 | -.04 | | -.01 | | .02 | | -.05 | | .03 | | -.04 | | -.06 | | .03 | |
| F2 | -.37*** | .02 | .25*** | .01 | -.10 | | .27*** | .01 | .15* | <.01 | -.32*** | .02 | -.27** | .01 | .21* | .01 |
| F3 | .03 | | -.08* | <.01 | .14*** | .01 | -.10** | .01 | -.08* | <.01 | .03 | | .07 | | -.09* | .01 |
| Intensity | -.01 | | -.01 | | .02 | | <.01 | | <.01 | | -.01 | | .01 | | -.06 | |
| Duration | .02 | | -.03 | | -.13** | .01 | .19*** | .02 | <.01 | | .05 | | -.17*** | .01 | .18*** | .02 |
| **C2** | | | | | | | | | | | | | | | | |
| F0 | <.01 | | .03 | | <-.01 | | -.07* | <.01 | .10*** | .01 | -.03 | | <-.01 | | .13*** | .01 |
| Intensity | .13*** | .01 | -.41*** | .11 | .27*** | .05 | -.33*** | .07 | -.35*** | .08 | .21*** | .03 | .25*** | .04 | -.17*** | .02 |
| Duration | -.03 | | .44*** | .07 | -.39*** | .06 | .51*** | .10 | .29*** | .03 | -.08 | | -.21*** | .02 | .26*** | .03 |
| **V2** | | | | | | | | | | | | | | | | |
| F0 | .01 | | <-.01 | | .07* | | -.01 | | -.02 | | <.01 | | <-.01 | | -.02 | |
| F1 | <-.01 | | .02 | | <-.01 | | .03 | | .07 | | .03 | | <.01 | | .01 | |
| F2 | -.58*** | .05 | <.01 | | -.06 | | .05 | | .05 | | .04 | | .30** | .01 | -.17 | |
| F3 | .09** | <.01 | -.07* | <.01 | .12** | | -.13** | .01 | -.11** | .01 | .09 | | .02 | | -.09 | |
| Intensity | <-.01 | | .15*** | .02 | -.13** | .01 | .17*** | .02 | .17*** | .02 | -.22*** | .04 | -.26*** | .05 | .27*** | .05 |
| Duration | .03 | | .03 | | -.02 | | .09* | .01 | .01 | | -.03 | | -.03 | | .06 | |

| | **Arousal** | | | | **Brightness** | | | | **Valence** | | | | **Size** | | | |
|---|---|---|---|---|---|---|---|---|---|---|---|---|---|---|---|---|
| | Calming | | Exciting | | Bright | | Dark | | Good | | Bad | | Small | | Big | |
| **$R^2$ (Adj $R^2$)** | .40 (.38) | | .31 (.29) | | .48 (.47) | | .30 (.28) | | .33 (.31) | | .24 (.22) | | .21 (.19) | | .28 (.26) | |
| **C1** | **β** | **$R^2\Delta$** | **β** | **$R^2\Delta$** | **β** | **$R^2\Delta$** | **β** | **$R^2\Delta$** | **β** | **$R^2\Delta$** | **β** | **$R^2\Delta$** | **β** | **$R^2\Delta$** | **β** | **$R^2\Delta$** |
| F0 | .04 | | -.03 | | -.02 | | -.04 | | .12** | .01 | -.05 | | .12** | .01 | -.11** | .01 |
| Intensity | .08 | | .03 | | .11* | <.01 | -.08 | | .10 | | -.05 | | -.18** | .01 | -.09 | |
| Duration | .05 | | .18** | .01 | -.07 | | .03 | | .10 | | -.08 | | .11 | | .04 | |

| | | | | | | | | | | | | | | | | |
|---|---|---|---|---|---|---|---|---|---|---|---|---|---|---|---|---|
| **V1** | | | | | | | | | | | | | | | | |
| F0 | -.03 | | .05 | | .05 | | -.15*** | .02 | .10** | .01 | -.08* | .01 | .04 | | -.05 | |
| F1 | -.04 | | <-.01 | | .11** | .01 | -.14** | .01 | .09 | | .02 | | -.03 | | .04 | |
| F2 | -.17* | <.01 | .30*** | .02 | .34*** | .02 | -.43*** | .03 | .06 | | -.18 | | -.06 | | .07 | |
| F3 | .02 | | -.01 | | <-.01 | | .03 | | .15*** | .02 | -.07 | | .04 | | -.20*** | .03 |
| Intensity | .02 | | <.01 | | .02 | | <.01 | | .02 | | -.03 | | -.03 | | .02 | |
| Duration | -.02 | | .02 | | -.07 | | .12* | .01 | -.30*** | .04 | .14* | .01 | -.24*** | .03 | .23*** | .03 |
| **C2** | | | | | | | | | | | | | | | | |
| F0 | .01 | | .04 | | <.01 | | -.02 | | .09* | .01 | -.08 | | .08 | | .04 | |
| Intensity | .35*** | .08 | -.15*** | .01 | -.11** | .01 | .03 | | .15*** | .01 | -.21*** | .03 | -.11* | .01 | -.06 | |
| Duration | -.23*** | .02 | .13* | .01 | .11* | <.01 | -.10 | | -.39*** | .06 | .28*** | .03 | -.09 | | .16** | .01 |
| **V2** | | | | | | | | | | | | | | | | |
| F0 | -.01 | | .04 | | .04 | | <-.01 | | .03 | | .04 | | .03 | | .01 | |
| F1 | -.05 | | -.01 | | -.06 | | -.04 | | .06 | | <-.01 | | .04 | | .03 | |
| F2 | -.24* | .01 | .23* | .01 | .33*** | .01 | -.13 | | .09 | | .08 | | .05 | | -.15 | |
| F3 | .17*** | .01 | -.05 | | <-.01 | | -.01 | | .04 | | -.12* | .01 | .03 | | -.08 | |
| Intensity | -.19*** | .02 | .11* | .01 | .20*** | .03 | -.03 | | -.11** | .01 | .14** | .01 | -.35*** | .09 | .36*** | .09 |
| Duration | -.08* | <.01 | .11* | .01 | <.01 | | -.08* | <.01 | -.08 | | .07 | | -.1* | .01 | .11* | .01 |

**Supplementary Figure 5: Shape**

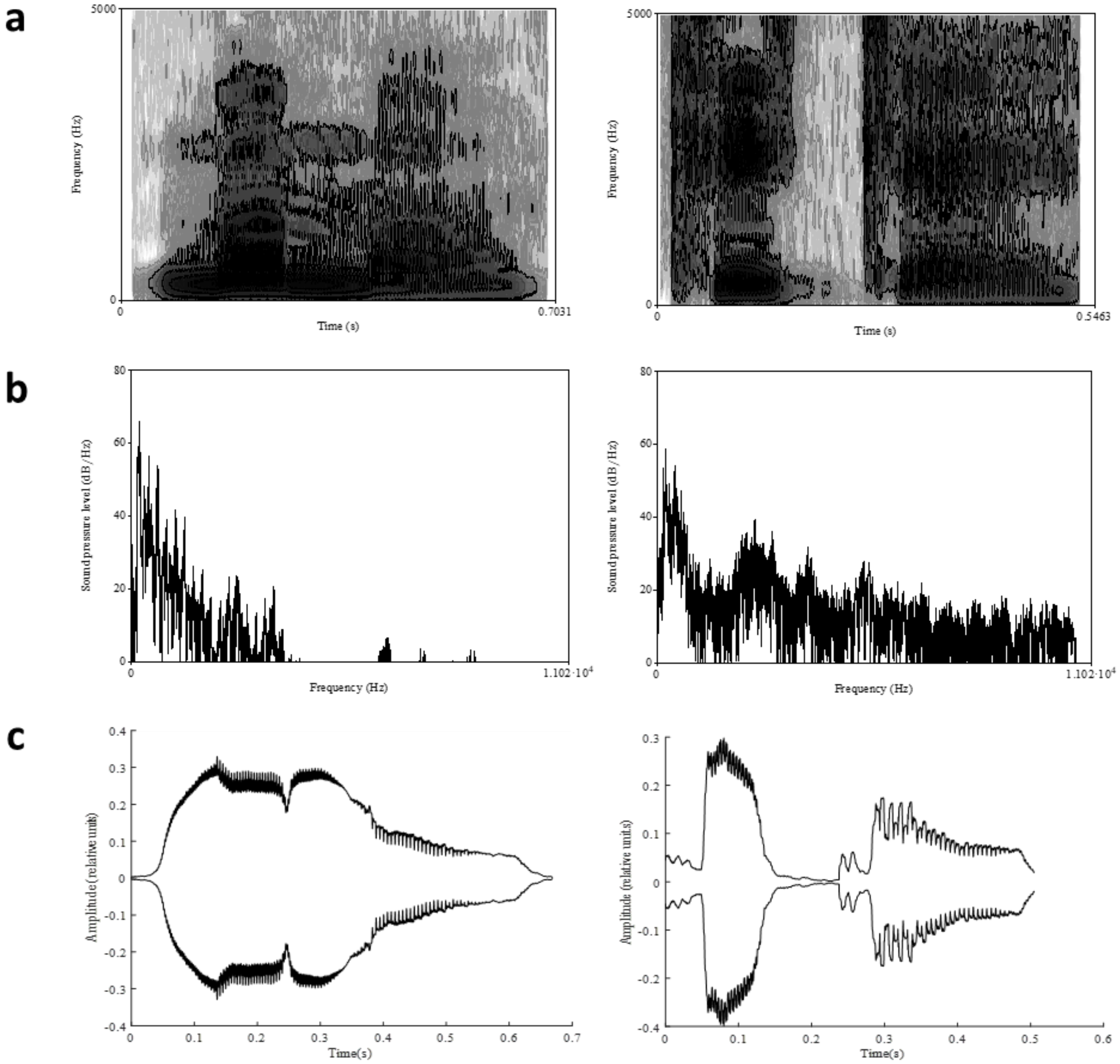

Spectro-temporal properties of representative highly-rated (see main text, Table 3) rounded (/mʊmo/, mean rating 4.9, left column) and pointed (/tike/, mean rating 5.5, right column) pseudowords: (a) spectrogram: frequency is low to high, bottom to top; shading = power, low power is lighter and high power is darker; (b) spectral tilt: frequency is low to high, left to right; (c) speech envelope.

**Supplementary Figure 6: Roughness**

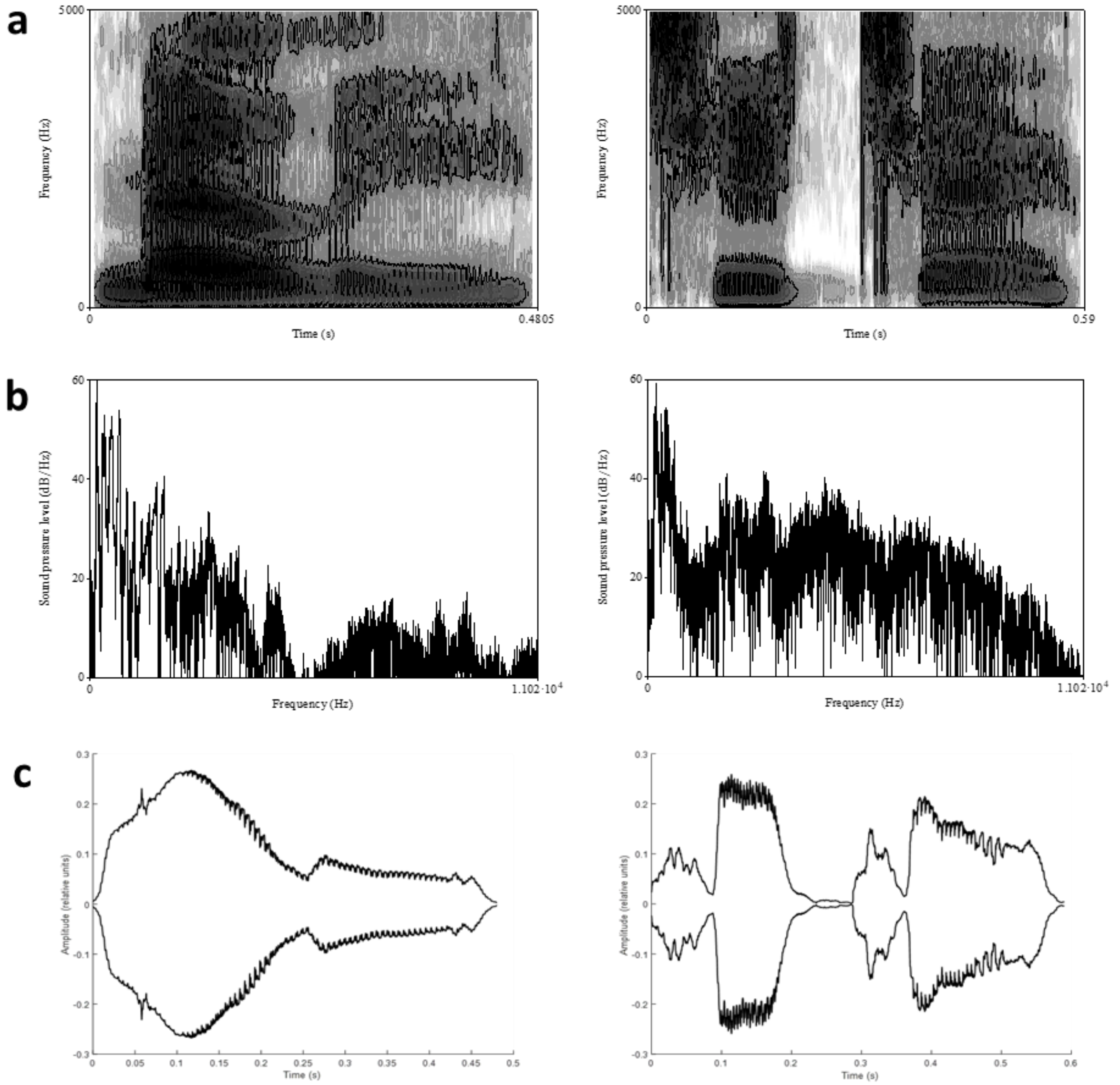

Spectro-temporal properties of representative highly-rated smooth (/mɛli/, mean rating 4.5, left column) and rough (/ʧiʧe/, mean rating 5.4, right column) pseudowords: (a) spectrogram; (b) spectral tilt; (c) speech envelope. Axis labels etc., as for Supplementary Figure 1.

**Supplementary Figure 7: Hardness**

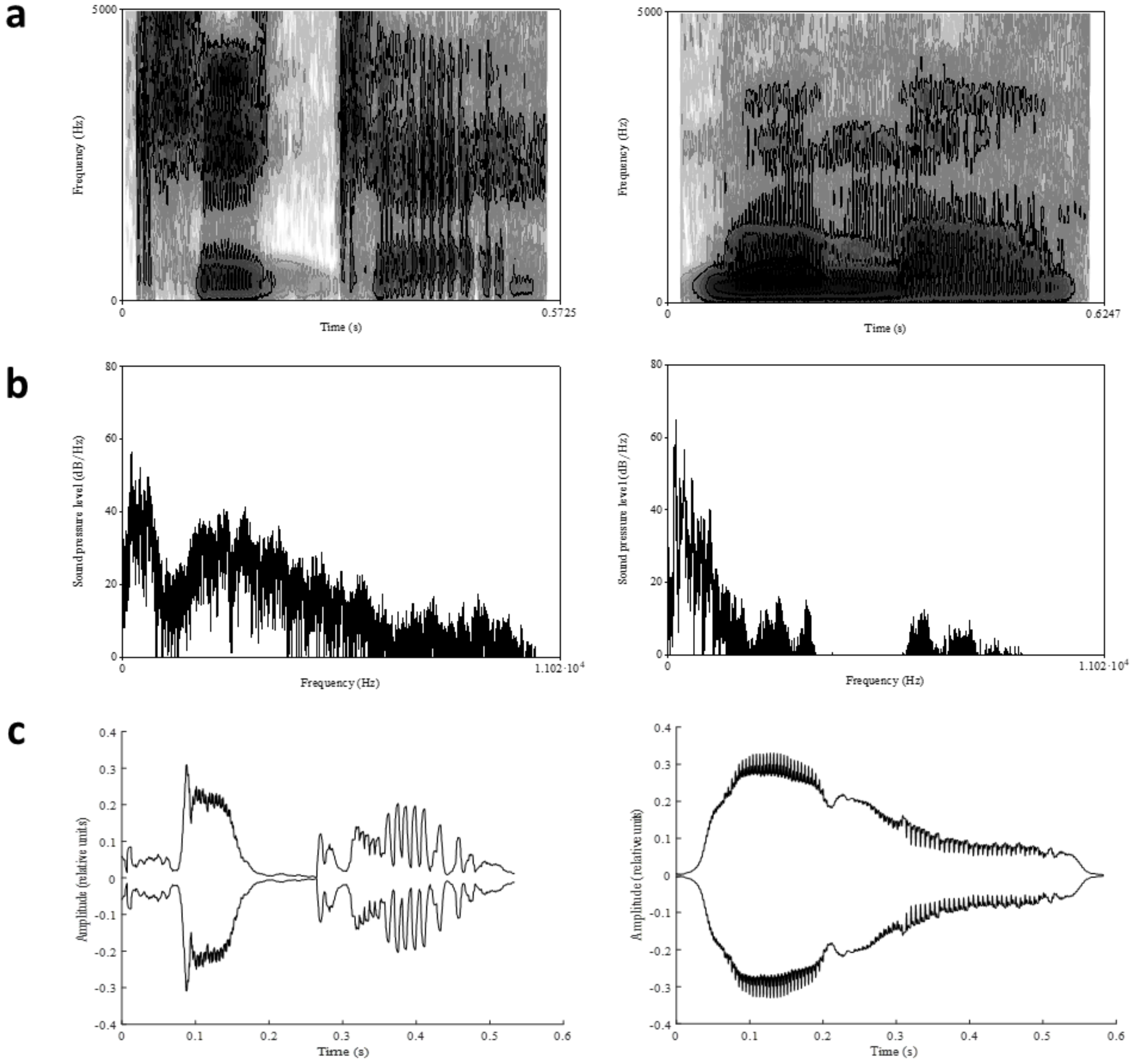

Spectro-temporal properties of representative highly-rated hard (/kike/, mean rating 5.0, left column) and soft (/mumo/, mean rating 5.2, right column) pseudowords: (a) spectrogram; (b) spectral tilt; (c) speech envelope. Axis labels etc., as for Supplementary Figure 1.

**Supplementary Figure 8: Weight**

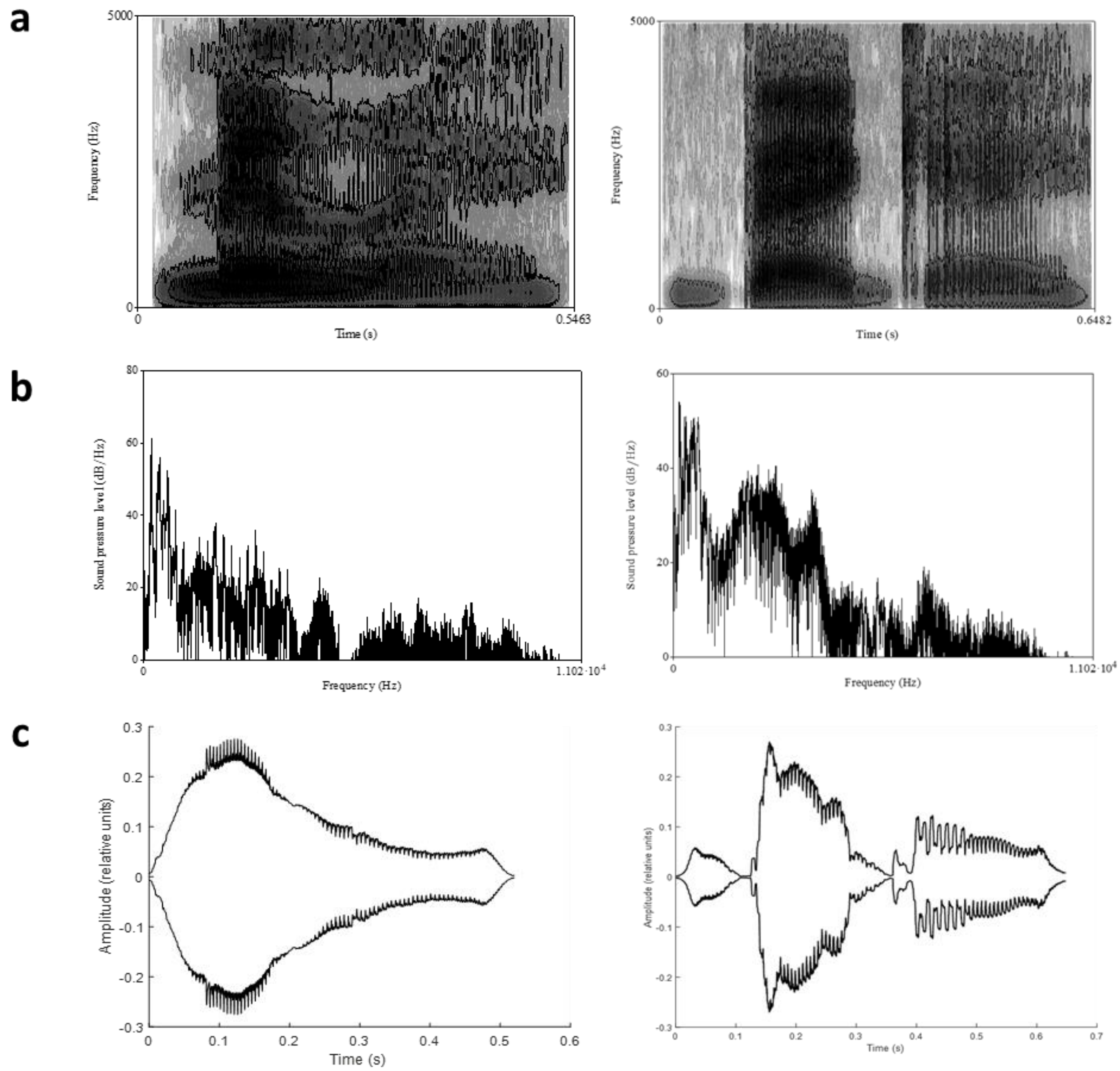

Spectro-temporal properties of representative highly-rated light (/mɪle/, mean rating 4.5, left column) and heavy (/bɛge/, mean rating 5.2, right column) pseudowords: (a) spectrogram; (b) spectral tilt; (c) speech envelope. Axis labels etc., as for Supplementary Figure 1.

**Supplementary Figure 9: Arousal**

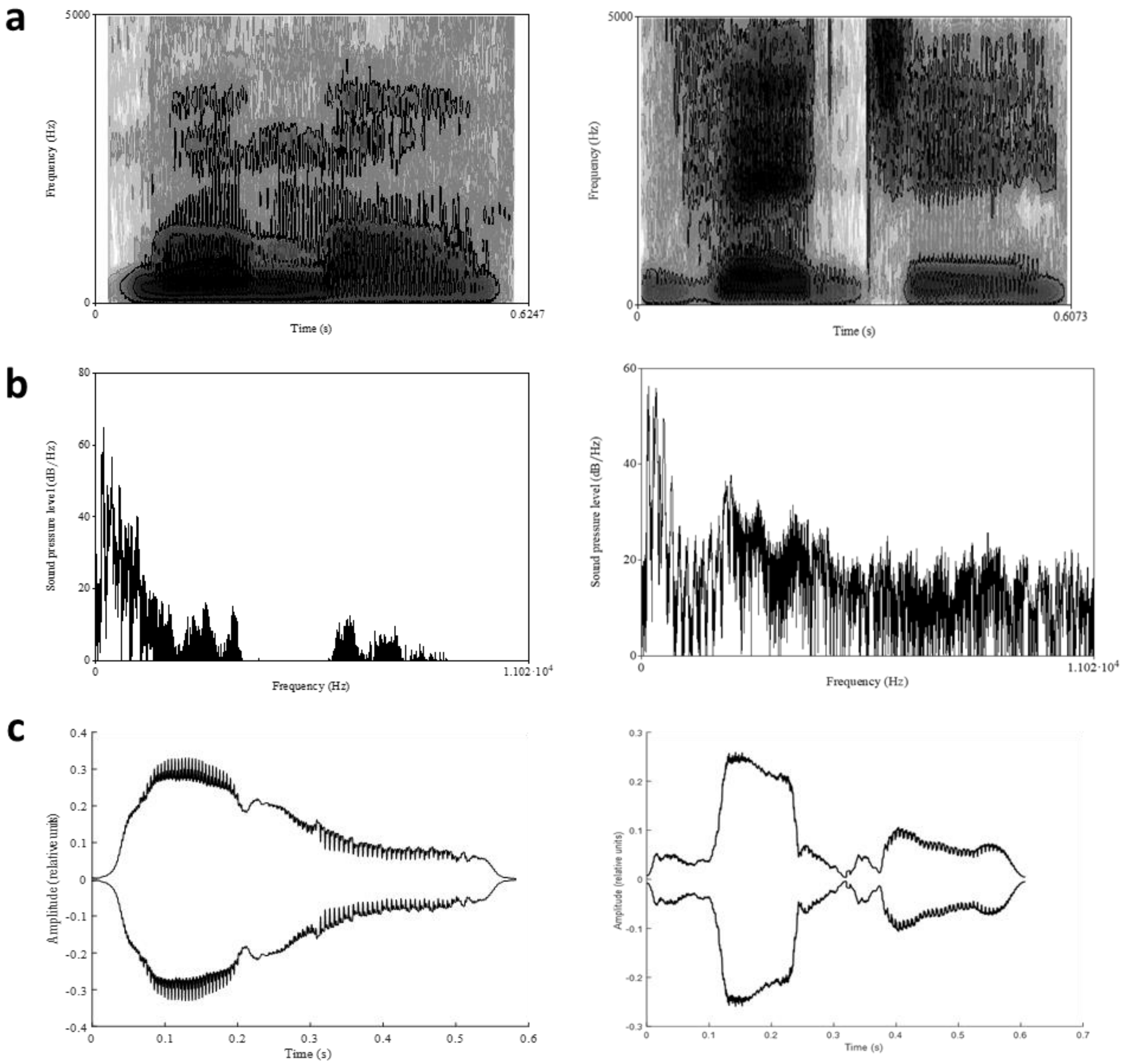

Spectro-temporal properties of representative highly-rated calming (/mumo/, mean rating 5.0, left column) and exciting (/zedʒi/, mean rating 4.7, right column) pseudowords: (a) spectrogram; (b) spectral tilt; (c) speech envelope. Axis labels etc., as for Supplementary Figure 1.

**Supplementary Figure 10: Brightness**

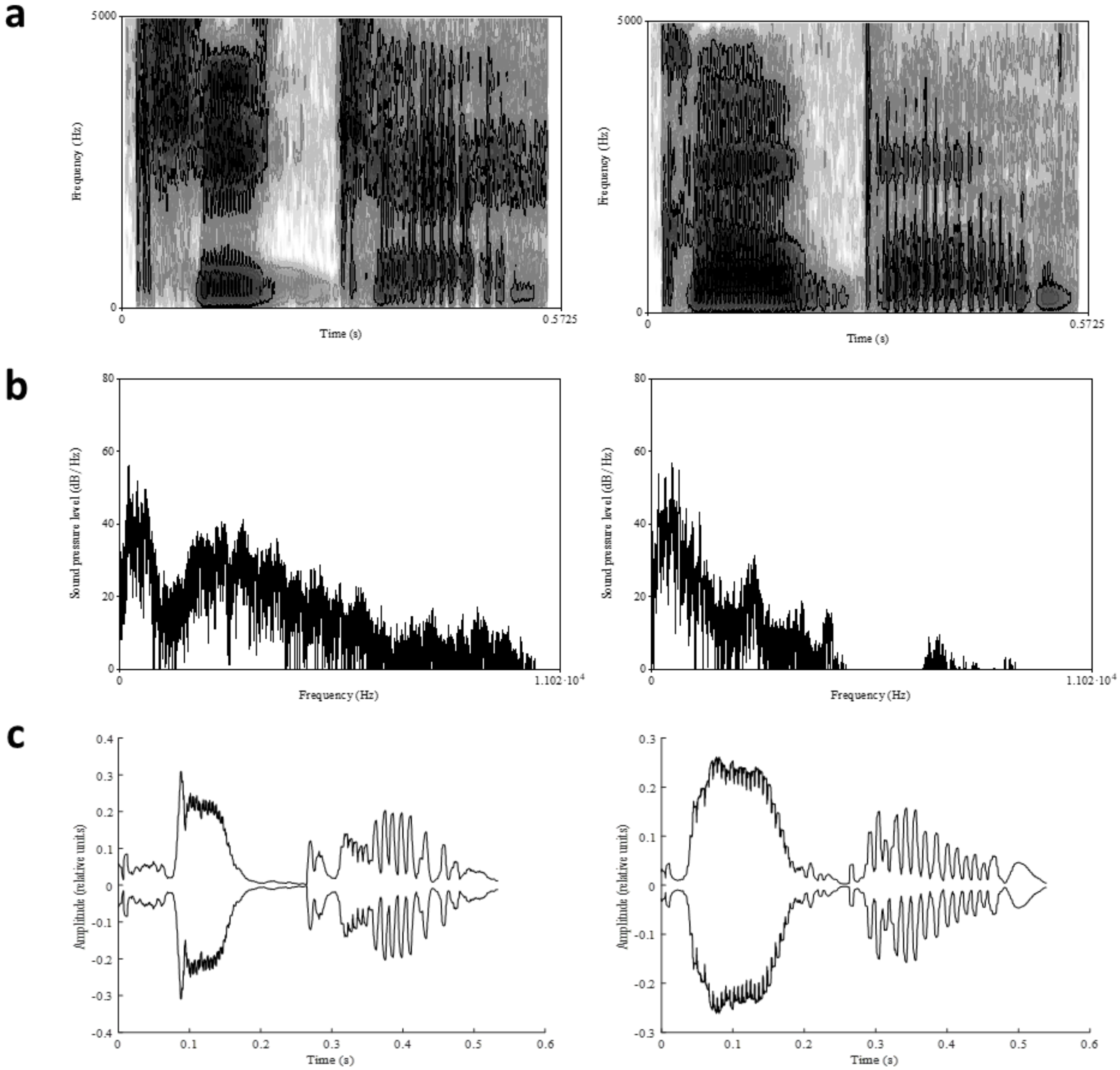

Spectro-temporal properties of representative highly-rated bright (/kike/, mean rating 5.1, left column) and dark (/gobo/, mean rating 4.7, right column) pseudowords: (a) spectrogram; (b) spectral tilt; (c) speech envelope. Axis labels etc., as for Supplementary Figure 1.

**Supplementary Figure 11: Valence**

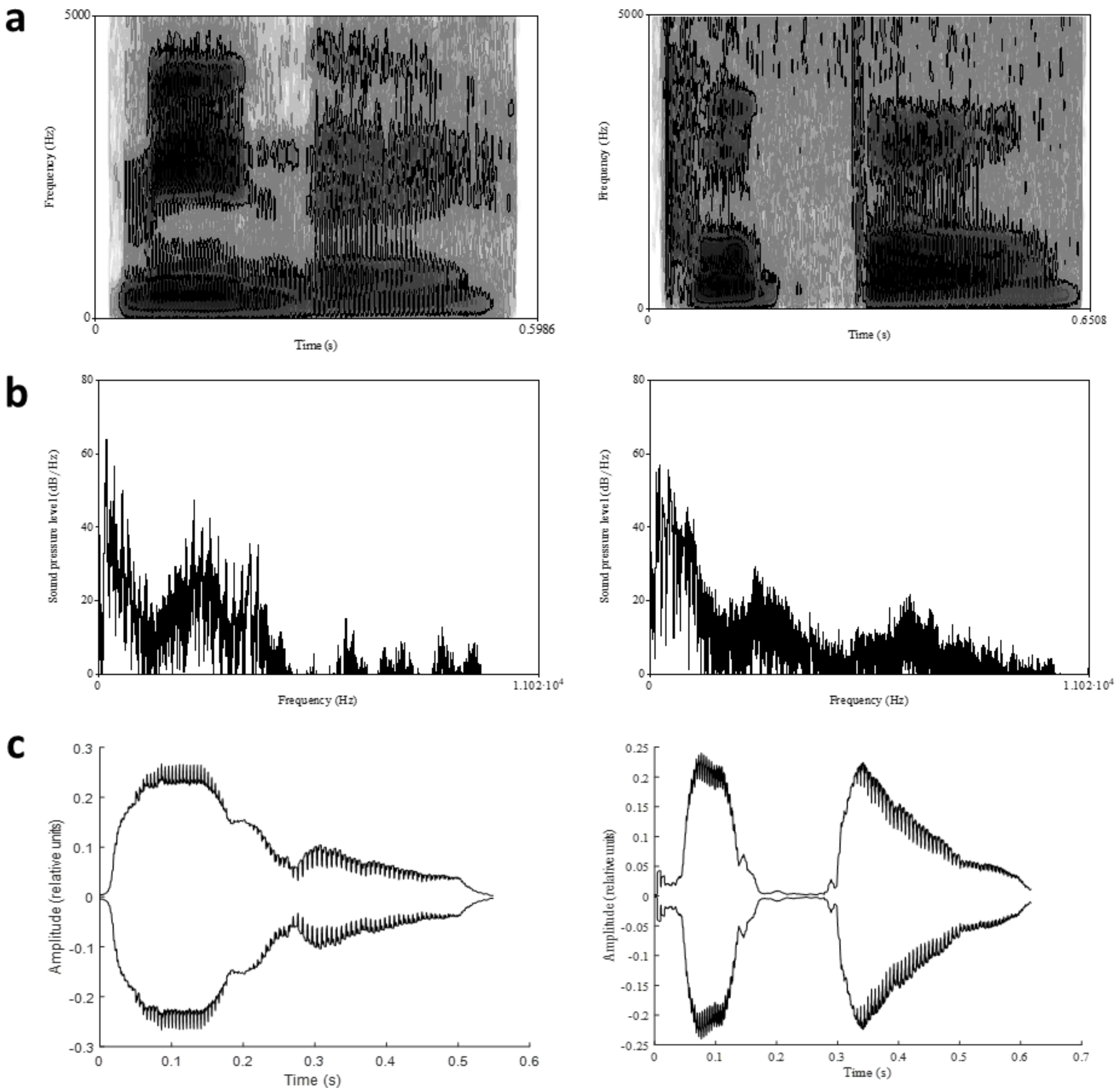

Spectro-temporal properties of representative highly-rated good (/mine/, mean rating 5.2, left column) and bad (/pupo/, mean rating 4.0, right column) pseudowords: (a) spectrogram; (b) spectral tilt; (c) speech envelope. Axis labels etc., as for Supplementary Figure 1.

**Supplementary Figure 12: Size**

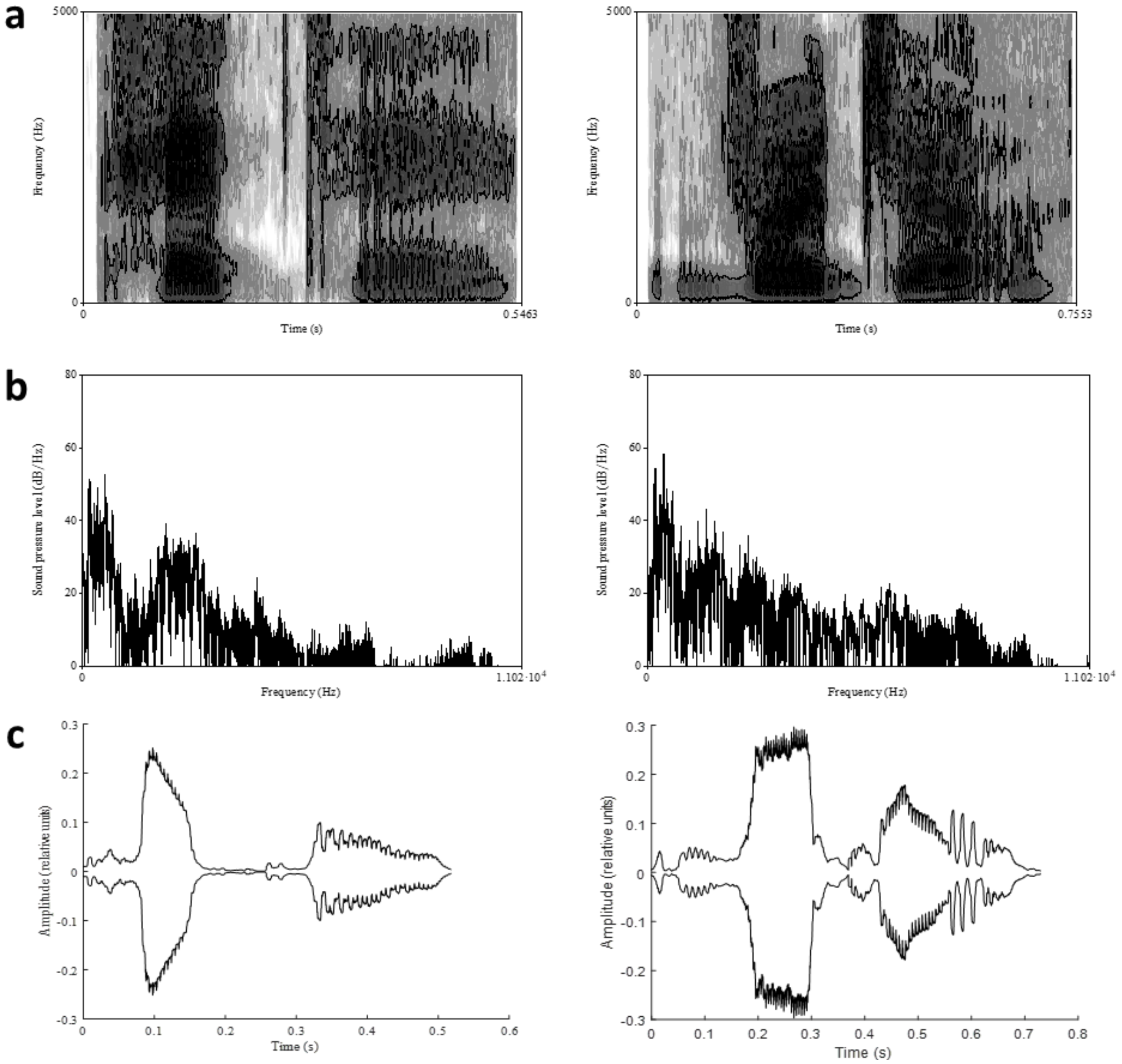

Spectro-temporal properties of representative highly-rated small (/pete/, mean rating 4.5, left column) and big (/vʊdʒo/, mean rating 4.8 right column) pseudowords: (a) spectrogram; (b) spectral tilt; (c) speech envelope. Axis labels etc., as for Supplementary Figure 1.
